# XCP-D: A Robust Pipeline for the post-processing of fMRI data

**DOI:** 10.1101/2023.11.20.567926

**Authors:** Kahini Mehta, Taylor Salo, Thomas Madison, Azeez Adebimpe, Danielle S. Bassett, Max Bertolero, Matthew Cieslak, Sydney Covitz, Audrey Houghton, Arielle S. Keller, Audrey Luo, Oscar Miranda-Dominguez, Steve M. Nelson, Golia Shafiei, Sheila Shanmugan, Russell T. Shinohara, Valerie J. Sydnor, Eric Feczko, Damien A. Fair, Theodore D. Satterthwaite

## Abstract

Functional neuroimaging is an essential tool for neuroscience research. Pre-processing pipelines produce standardized, minimally pre-processed data to support a range of potential analyses. However, post-processing is not similarly standardized. While several options for post-processing exist, they tend not to support output from disparate pre-processing pipelines, may have limited documentation, and may not follow BIDS best practices. Here we present XCP-D, which presents a solution to these issues. XCP-D is a collaborative effort between PennLINC at the University of Pennsylvania and the DCAN lab at the University at Minnesota. XCP-D uses an open development model on GitHub and incorporates continuous integration testing; it is distributed as a Docker container or Singularity image. XCP-D generates denoised BOLD images and functional derivatives from resting-state data in either NifTI or CIFTI files, following pre-processing with fMRIPrep, HCP, and ABCD-BIDS pipelines. Even prior to its official release, XCP-D has been downloaded >3,000 times from DockerHub. Together, XCP-D facilitates robust, scalable, and reproducible post-processing of fMRI data.

## INTRODUCTION

Functional neuroimaging using fMRI is an essential tool for human neuroscience research. Widely used pre-processing pipelines, such as fMRIPrep (Esteban et al., 2018), Human Connectome Project (HCP) pipelines (Glasser et al., 2013), and ABCD-BIDS (Feczko et al., 2021) produce standardized, minimally pre-processed data to support a range of potential analyses. Following pre-processing, investigators typically perform post-processing, which includes critical steps like denoising and generation of derived measures (e.g., functional networks) that are used in hypothesis testing. Unlike the highly standardized software available for pre-processing, there is far more variability in how researchers approach post-processing, for example censoring data to remove high-motion outliers, or despiking data to remove large spikes in images. In general, different approaches towards denoising in the post-processing stage can lead to different results from the same set of data. Prior work has also established that denoising strategies are quite heterogeneous in their effectiveness (Ciric et al., 2017). This may result in findings that cannot be replicated, contradictory results, and other such issues that make it harder for the field to progress. Here we introduce XCP-D: a scalable, robust, and generalizable software package for post-processing resting-state fMRI data.

Widely used pre-processing tools such as fMRIPrep build on the Brain Imaging Data Structure (BIDS) for organizing and describing neuroimaging data (Gorgolewski et al., 2016). As a BIDS App, fMRIPrep builds appropriate pre-processing workflows based on the metadata encoded by BIDS. Following pre-processing with fMRIPrep, many labs use custom workflows for post-processing steps including denoising and generation of derivatives. While such a bespoke approach to post-processing may have advantages – such as being tightly aligned with the needs of a specific study – it leads to the duplication of effort across labs, negatively impacts reproducibility, and may reduce the generalizability of results. One alternative to custom post-processing has been provided by the eXtensible Connectivity Pipelines Engine (XCP; Ciric et al., 2018), a widely used (>6,000 Docker pulls) post-processing package that consumes fMRIPrep output. However, XCP has accumulated substantial technical debt over time, is not compatible with other widely used pre-processing formats (e.g: HCP pipelines), does not support surface-based analyses, and lacks certain advanced denoising features provided by other widely used packages such as ABCD-BIDS.

Here, we introduce XCP-D, a collaborative effort between PennLINC (Pennsylvania Lifespan Informatics and Neuroimaging Center) and DCAN (Developmental Cognition and Neuroimaging Labs) that includes a new Python codebase and important new features. XCP-D focuses on consuming data pre-processed by other widely used tools. Specifically, XCP-D supports post-processing of multiple pre-processed formats, including fMRIPrep, HCP pipelines, and ABCD-BIDS; this allows XCP-D users to apply the same top-performing denoising strategies to datasets that were pre-processed using different software. XCP-D adheres to BIDS derivatives conventions throughout and includes new software engineering features to ensure stability and robustness. These include a refactored and highly modular codebase that is built using NiPype (Gorgolewski et al., 2011) and incorporates extensive continuous integration (CI) testing. Additionally, XCP-D supports CIFTI workflows for surface-based analysis and processing, provides an expanded suite of data quality measures, and includes new visual reports. XCP-D thus allows users to leverage minimally processed data from diverse data resources, apply uniform post-processing, and generate the same derived measures for hypothesis testing. Prior to publication, XCP-D has already been pulled from DockerHub over 3000 times.

## METHODS

### Overview

XCP-D consumes pre-processed resting-state data generated with any of three commonly used pre-processing pipelines: fMRIPrep, HCP, or ABCD-BIDS and implements top-performing denoising strategies (Ciric et al., 2018) for NIfTI or CIFTI timeseries. The pipeline generates resting-state derivatives, including parcellated timeseries and connectivity matrices, using multiple popular atlases. Importantly, XCP-D also calculates additional quality assurance measures. Finally, XCP-D constructs interactive reports that describe the post-processing methods used and facilitate visualization of each step. XCP-D also uses an open, test-driven development model on GitHub, and is distributed as a Docker container or Singularity image.

### Installation procedures

#### Docker

Docker is an open-source platform for developers that makes the distribution of applications easier via packaging of all supporting dependencies into a lightweight, standard form called a “container” (Rad et al., 2017). Docker images create a container that includes the complete operating system and all necessary dependencies. For every new version of XCP-D, continuous integration testing is performed (see **Table 1** for a list of tests implemented in XCP-D). If these tests succeed, a new Docker image is automatically generated and deployed to DockerHub. To run XCP-D via Docker images, Docker Engine must be installed. To pull XCP-D from DockerHub, users must run:

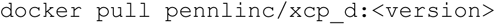

where `<VERSION<` should be replaced with the desired version or tag of XCP-D that users want to download. The image can also be found here: https://registry.hub.docker.com/r/pennlinc/xcp_d XCP-D can be run by interacting directly with the Docker Engine via the `docker run` command.

**Table 1:** Continuous integration tests for different XCP-D stages.

| **XCP-D STEP** | **TESTING** |
| --- | --- |
| Confound selection | Confirming that a loaded confound matrix has the right shape. |
| Removing dummy time | Looping through N=1-10 volumes to be dropped and confirming that the correct number of volumes have been dropped from a BOLD file and the corresponding confounds file. |
| Censoring | Replacing values in the confounds file with values that should be omitted and confirming that the image file and the confounds file have had the same number of volumes dropped. |
| Despiking | Confirming that the maximum value of the data has decreased, and the minimum value of the data has increased after despiking. |
| Confound regression | Confirming that the correlation between a random voxel and the confounds timeseries has decreased after regression. |
| Interpolation | Confirming that the difference between the fast Fourier transform (FFT) of an interpolated file and the original file is less than the difference between the FFT of a file with a spike planted in it and the original file. |
| Filtering | Comparing output of the XCP-D code after filtering to the output of SciPy's code directly in a separate Python notebook. |
| Functional timeseries and connectivity matrices | Confirming that the correlation coefficient of a parcellated timeseries is the same as the connectivity matrix produced. (Parcellations were also performed manually in a separate Python notebook and compared to results from XCP-D.) |
| ReHo | Adding artificial noise to an image and confirming that the mean ReHo value decreases. |
| ALFF | Computing the FFT of a BOLD file, adding to |
|  | the amplitude of its lower frequencies and confirming the ALFF increases. |
| Residual BOLD and resting-state derivatives smoothing | Confirming that smoothness has increased after the module - via AFNI for NIFTIs and via Connectome Workbench for CIFTIs. |
| Quality control | Visually inspecting the quality check reports. |

#### Singularity

Singularity is an open-source software package designed to allow portable computational environments and containers for scientific research (Kurtzer et al., 2017). Many high performance computing (HPC) systems restrict use of Docker, but support Singularity instead. Using Singularity version 2.5 or higher, users can create a Singularity image from a Docker image on DockerHub:

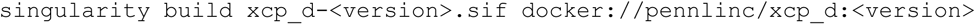

### Design and testing

We used an open-source, test-driven approach in developing XCP-D. To this end, we integrated CircleCI – a web-based continuous integration testing platform – into our development workflow. Each new commit to the software is run through a full suite of CI tests (described in **Table 1**) run on pre-selected datasets during each CircleCI instance. Further, we applied branch protection rules to the development process: namely, any changes to XCP-D must be approved by a reviewer and pass continuous integration testing and full pipeline runs on CircleCI before deployment to the main branch that can be accessed by users. Approximately 81% of the code is covered by our tests according to CodeCov – which determines how much of the codebase is covered by CI testing.

### Workflow

Post-processing in XCP-D involves multiple customizable steps that are widely used: the removal of dummy volumes, despiking, temporal censoring, regression, interpolation, filtering, smoothing, supplemented by the calculation of quality assurance variables, and generation of reports (Satterthwaite et al., 2013; Ciric et al., 2018; see **Figure 1**). Note that XCP-D supports post-processing of fMRI data with a T1 image, a T2 image, or both.

**Figure 1:**
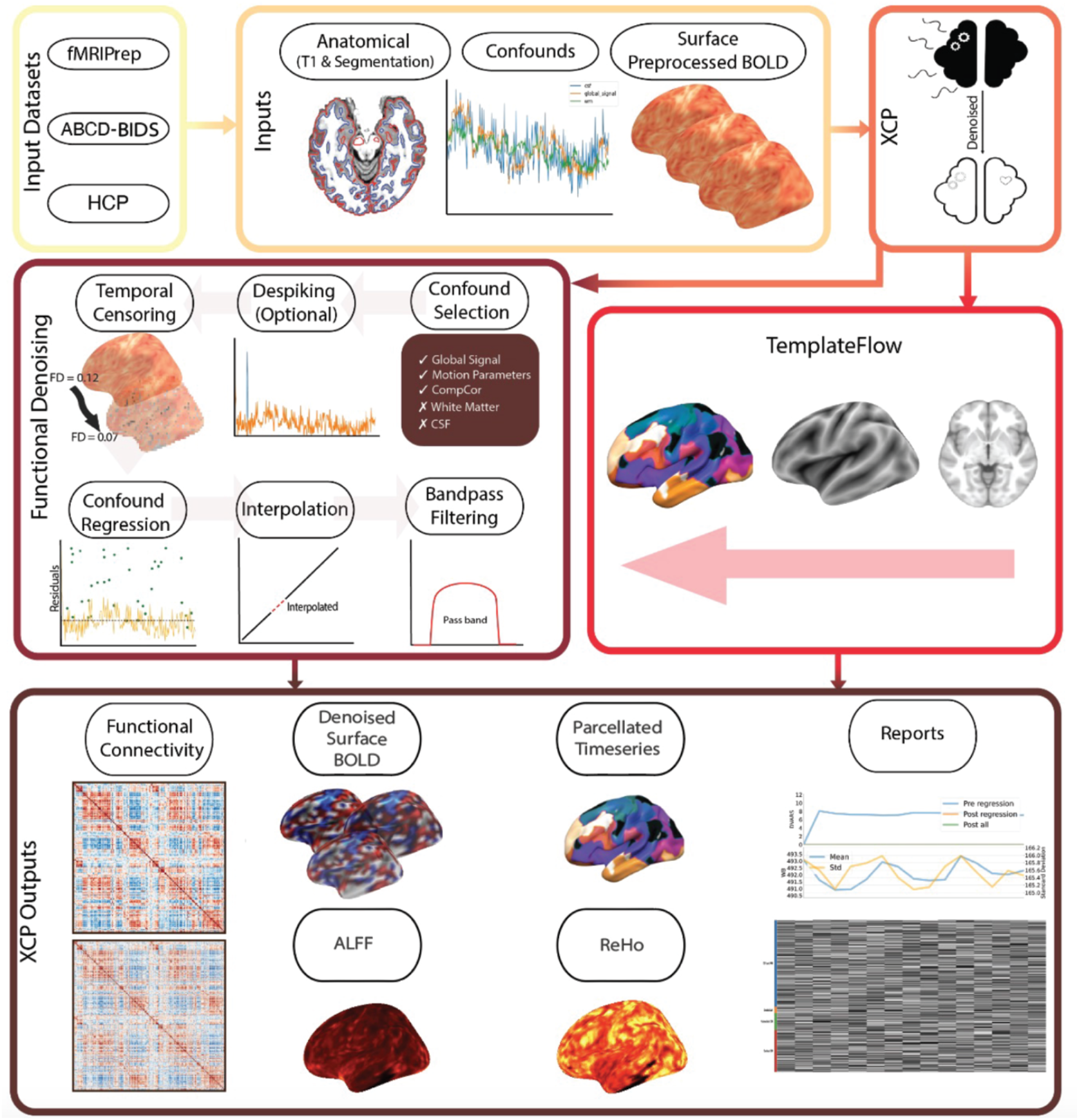
XCP-D Workflow. The XCP-D workflow begins after the pre-processing of fMRI data. XCP-D requires anatomical data, confounds files, and pre-processed BOLD files. It performs functional denoising to produce clean fMRI data and functional derivatives. *ReHo: Regional Homogeneity; ALFF: Amplitude of Low Frequency Fluctuations*.

Through these processes, XCP-D produces multiple functional derivatives, including the dense volumetric and/or surface-based denoised timeseries, parcellated timeseries, correlation matrices, and derived functional metric maps (such as regional homogeneity and fluctuation amplitude). Furthermore, XCP-D also provides detailed quality assurance information regarding both the fMRI data and image registration, as well as interactive graphical reports (see **Table 2** for a list and description of XCP-D outputs).

**Table 2:**
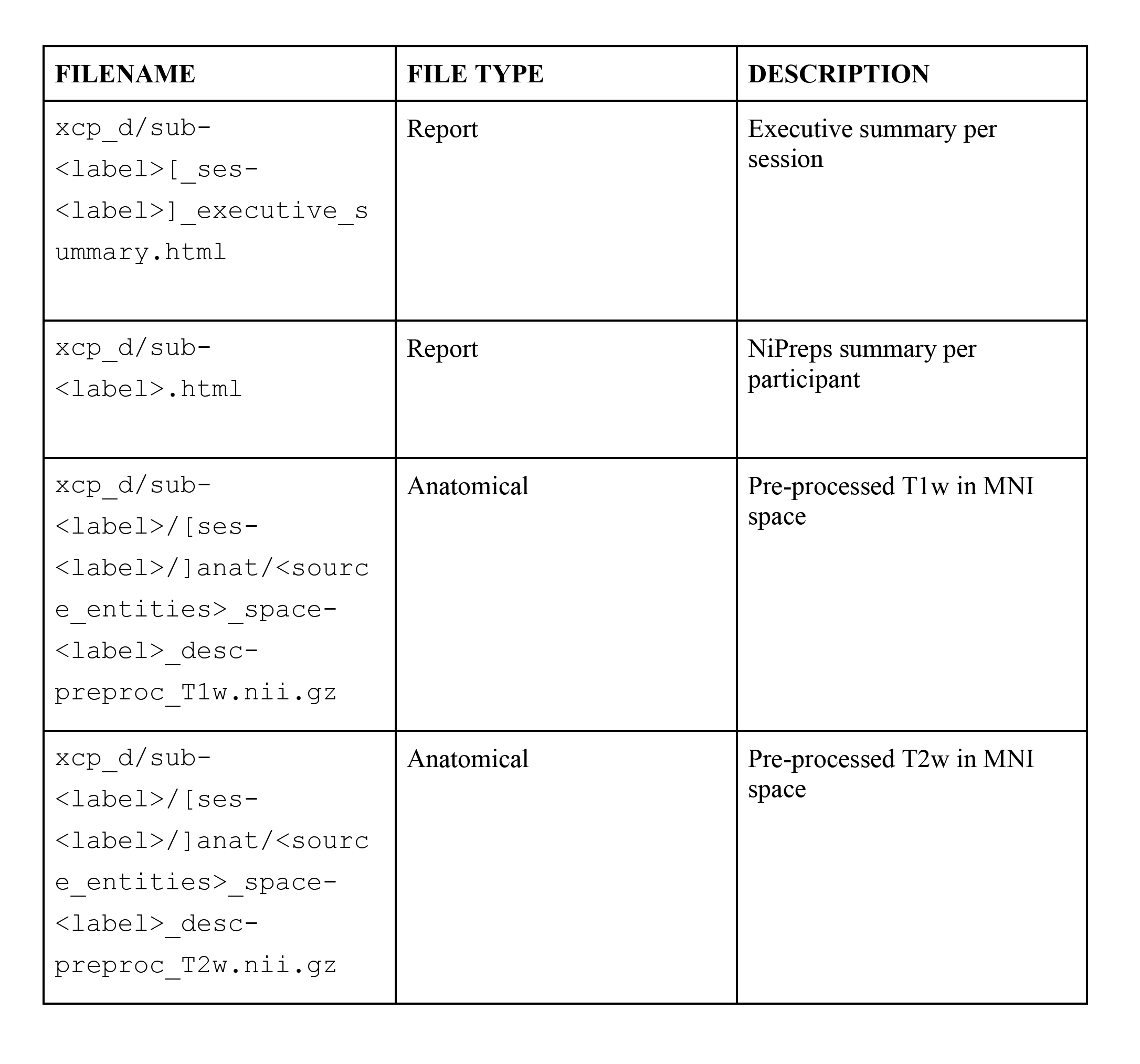
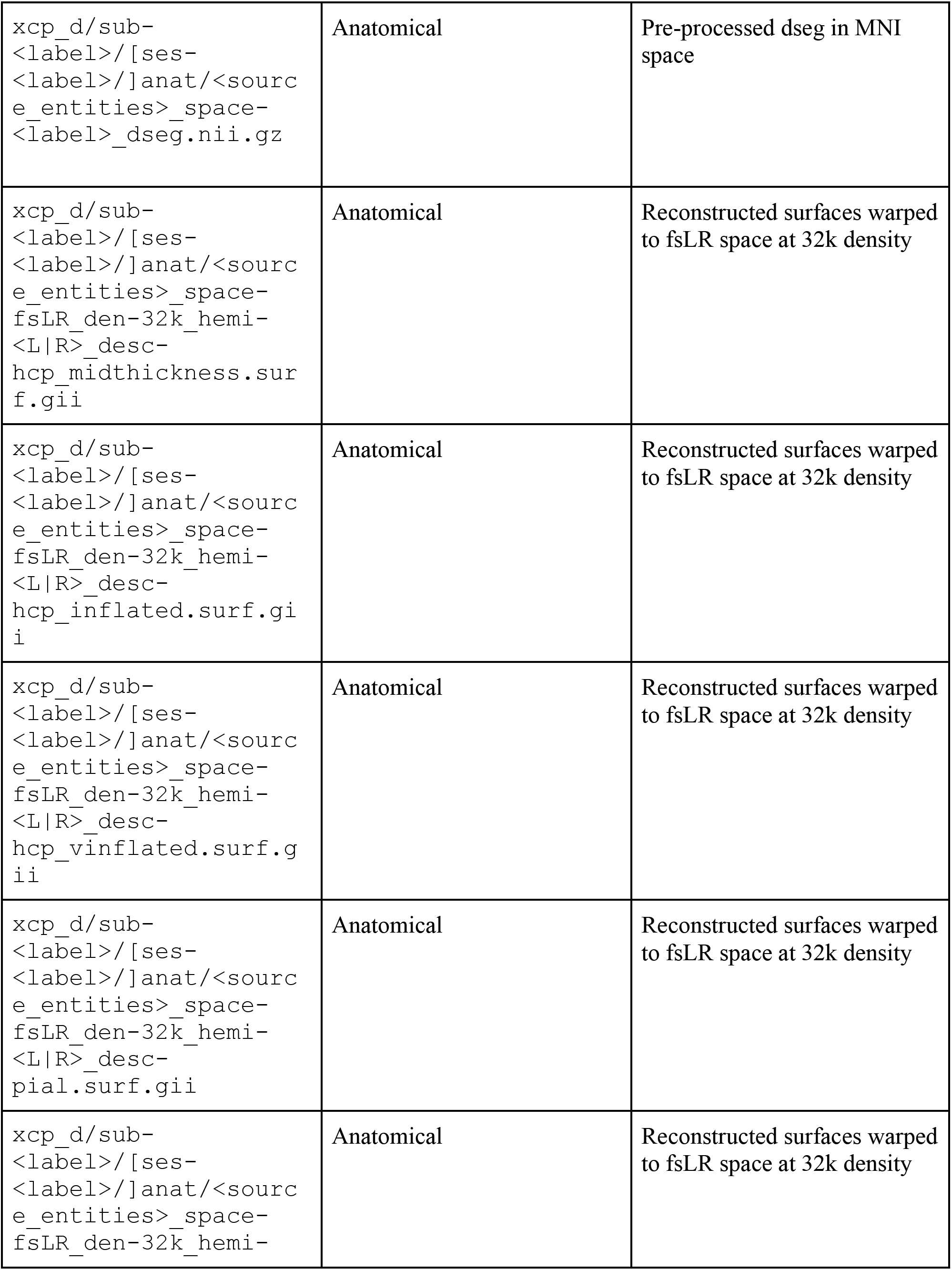
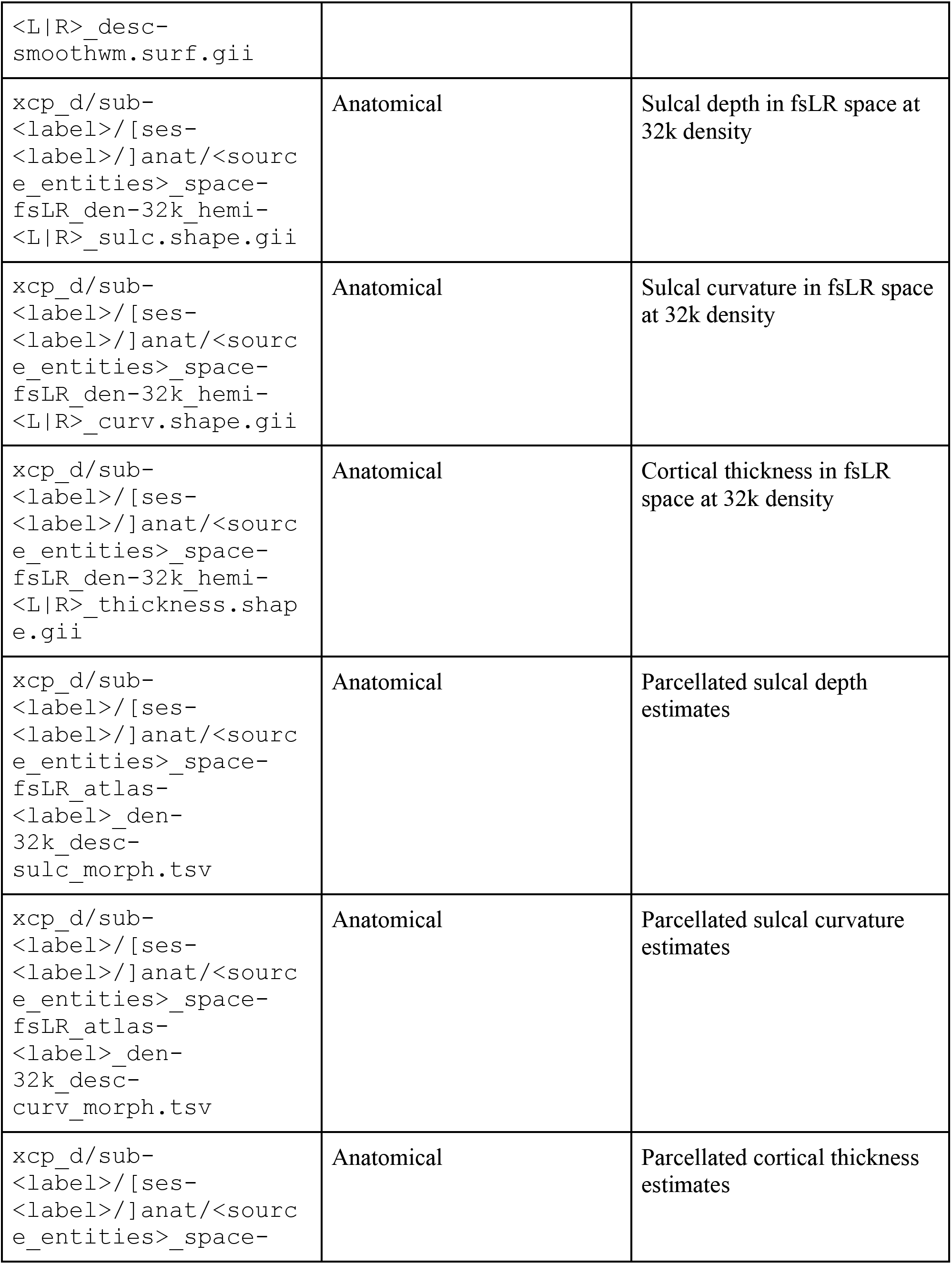
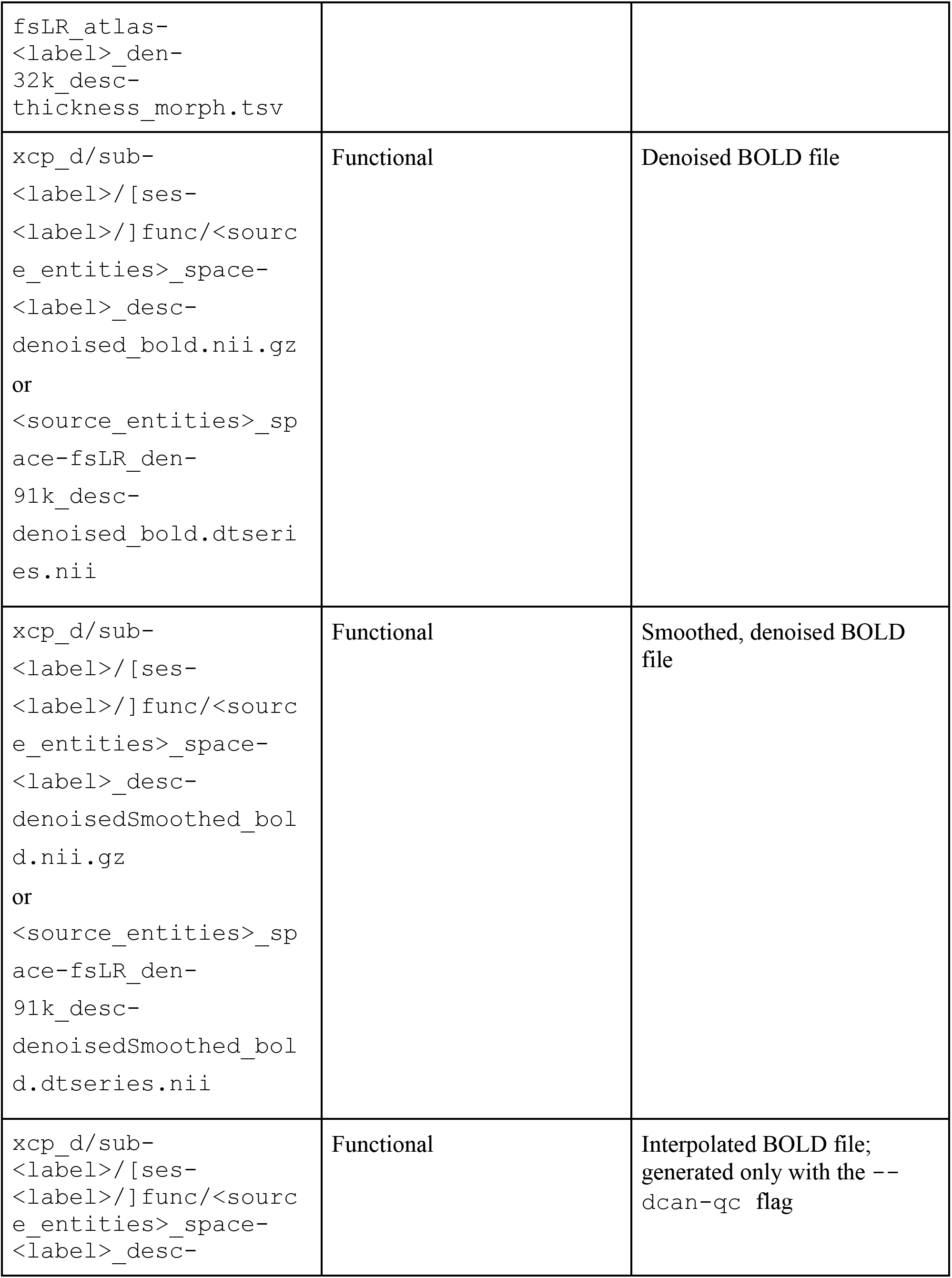
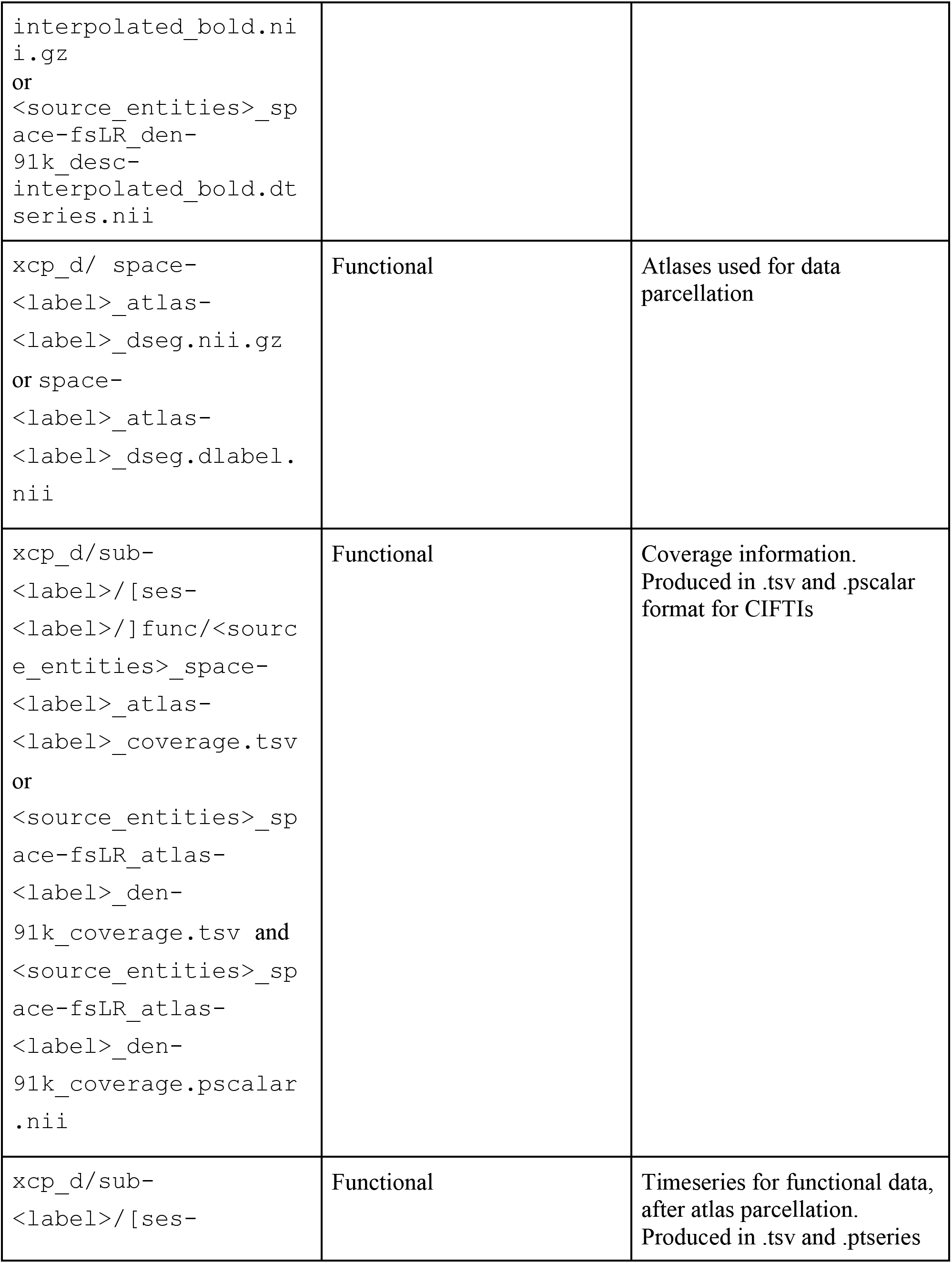
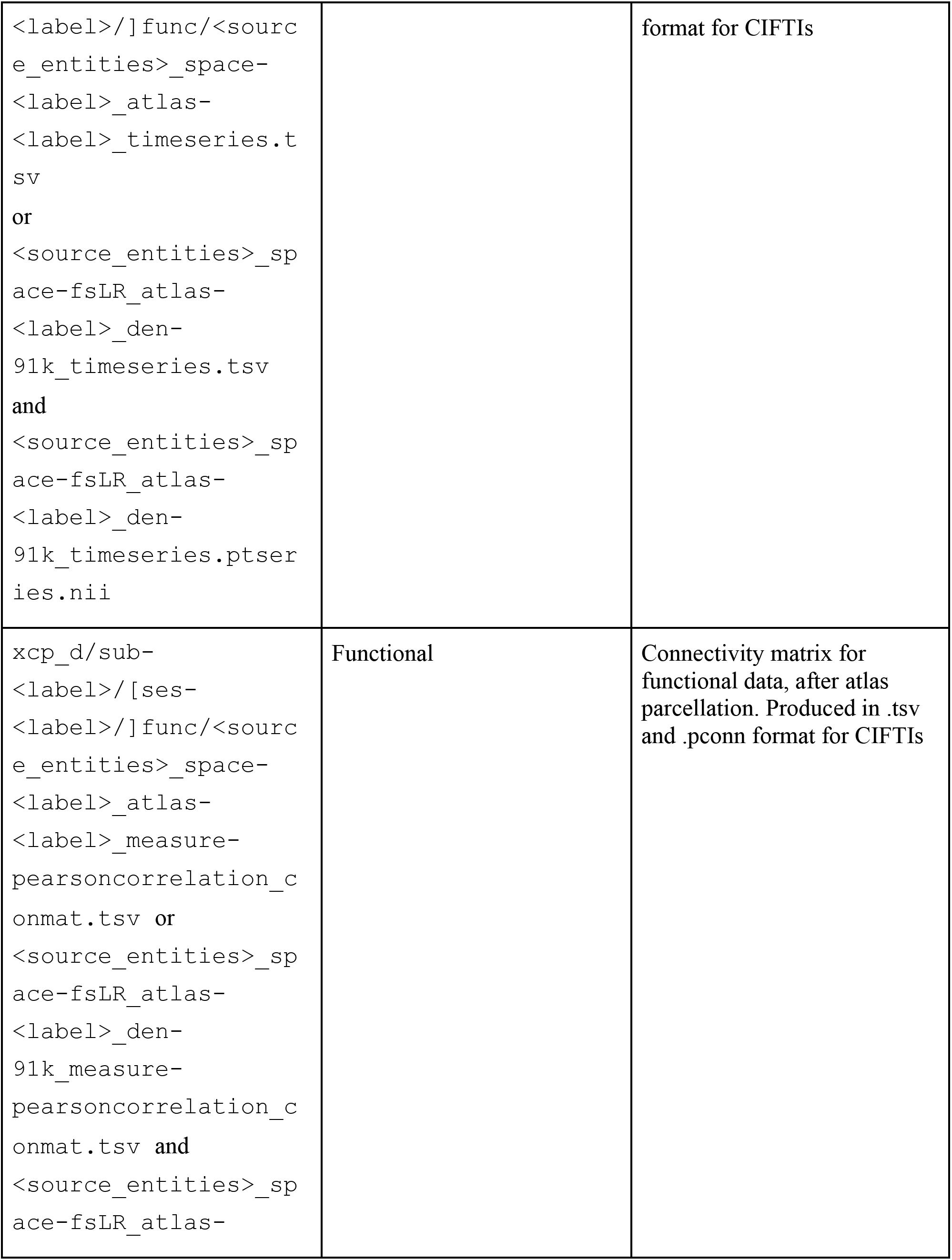
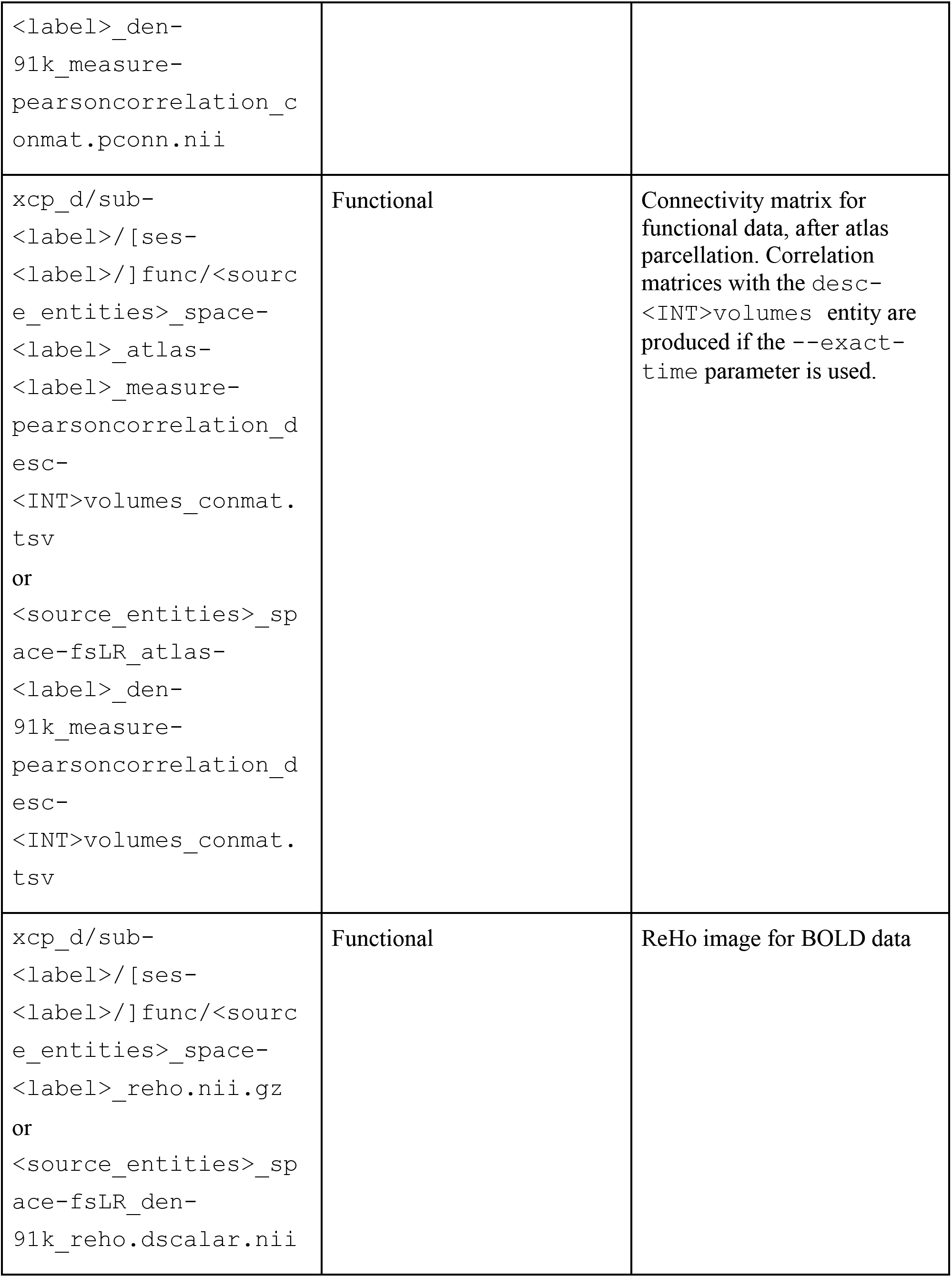
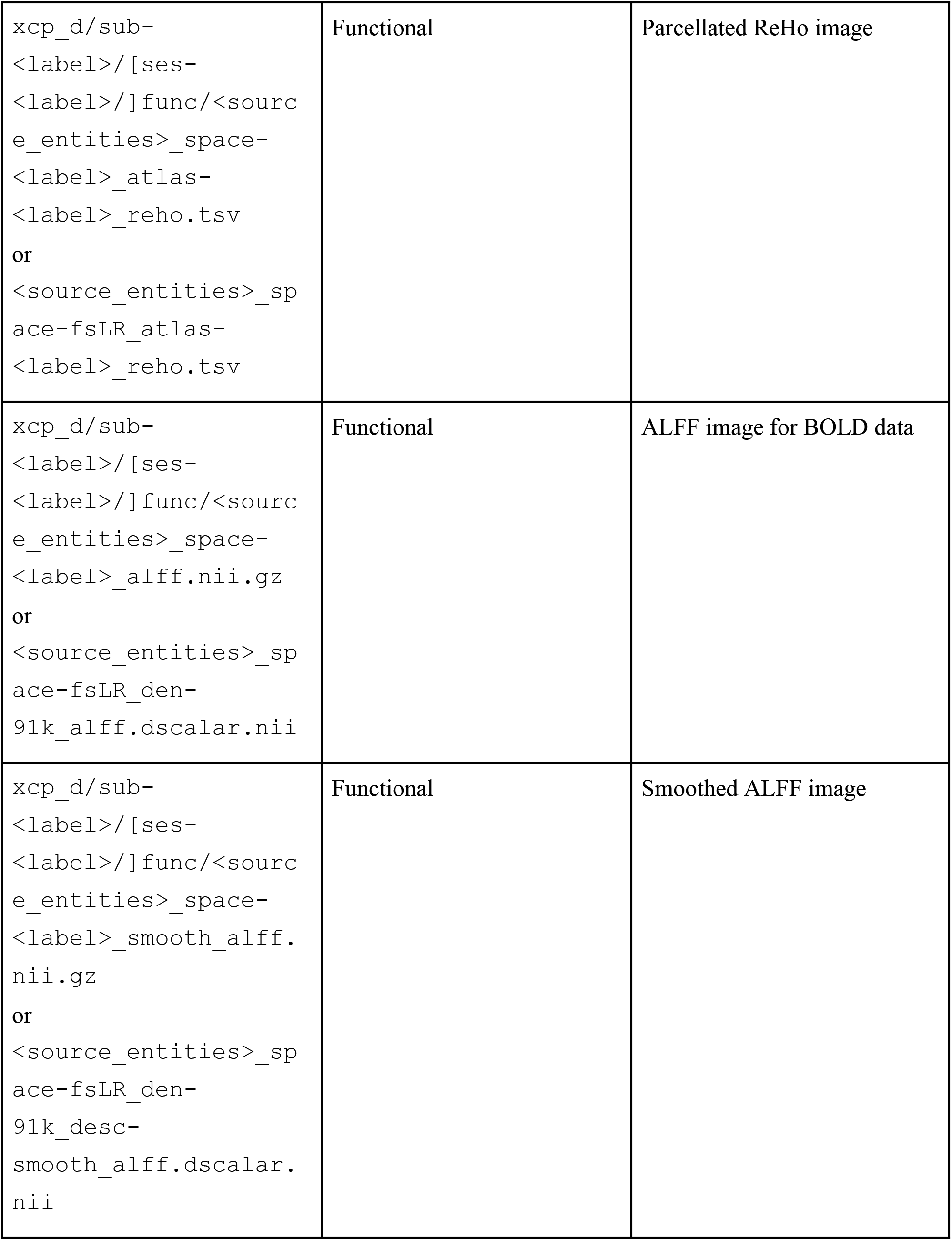
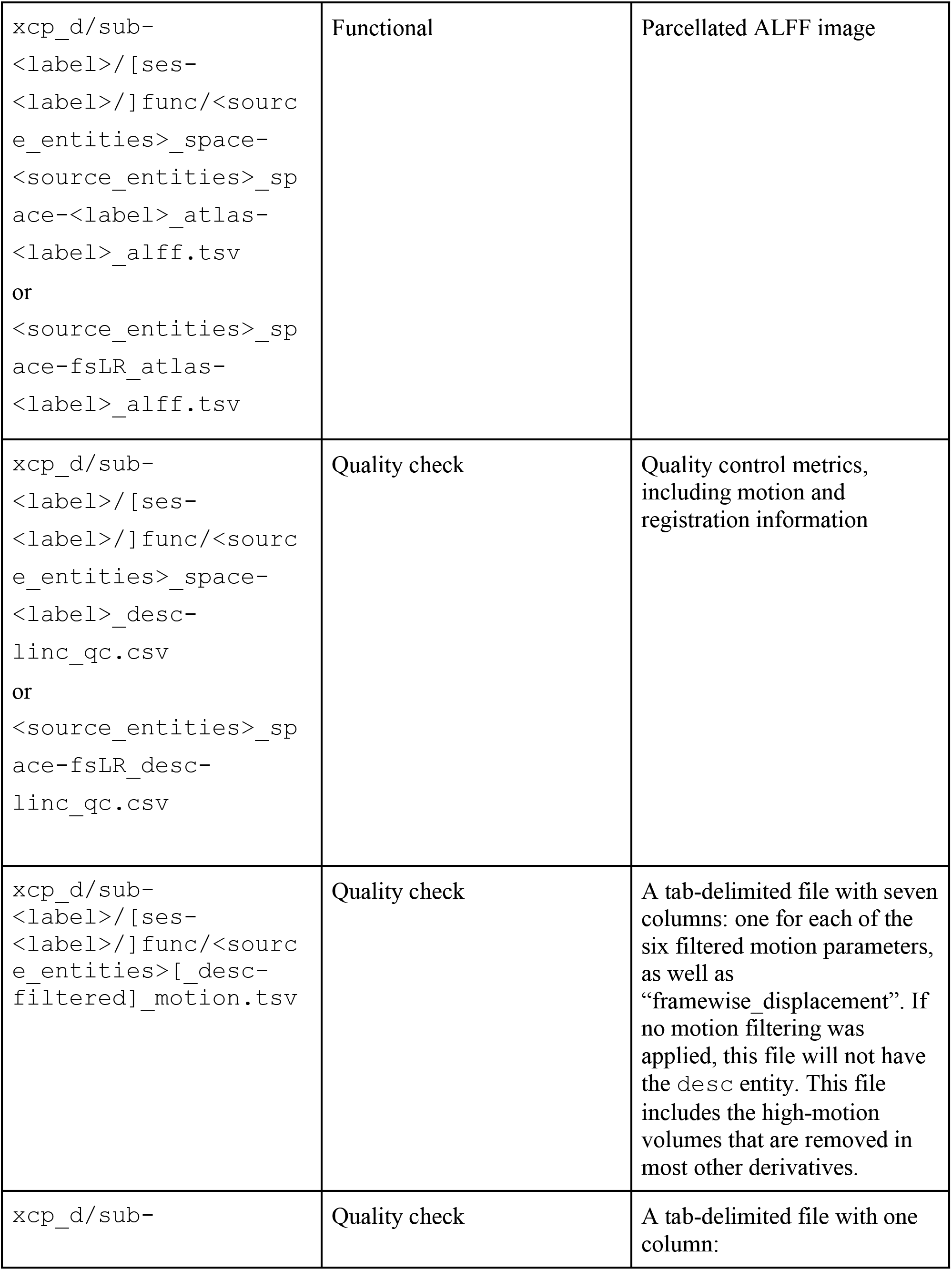
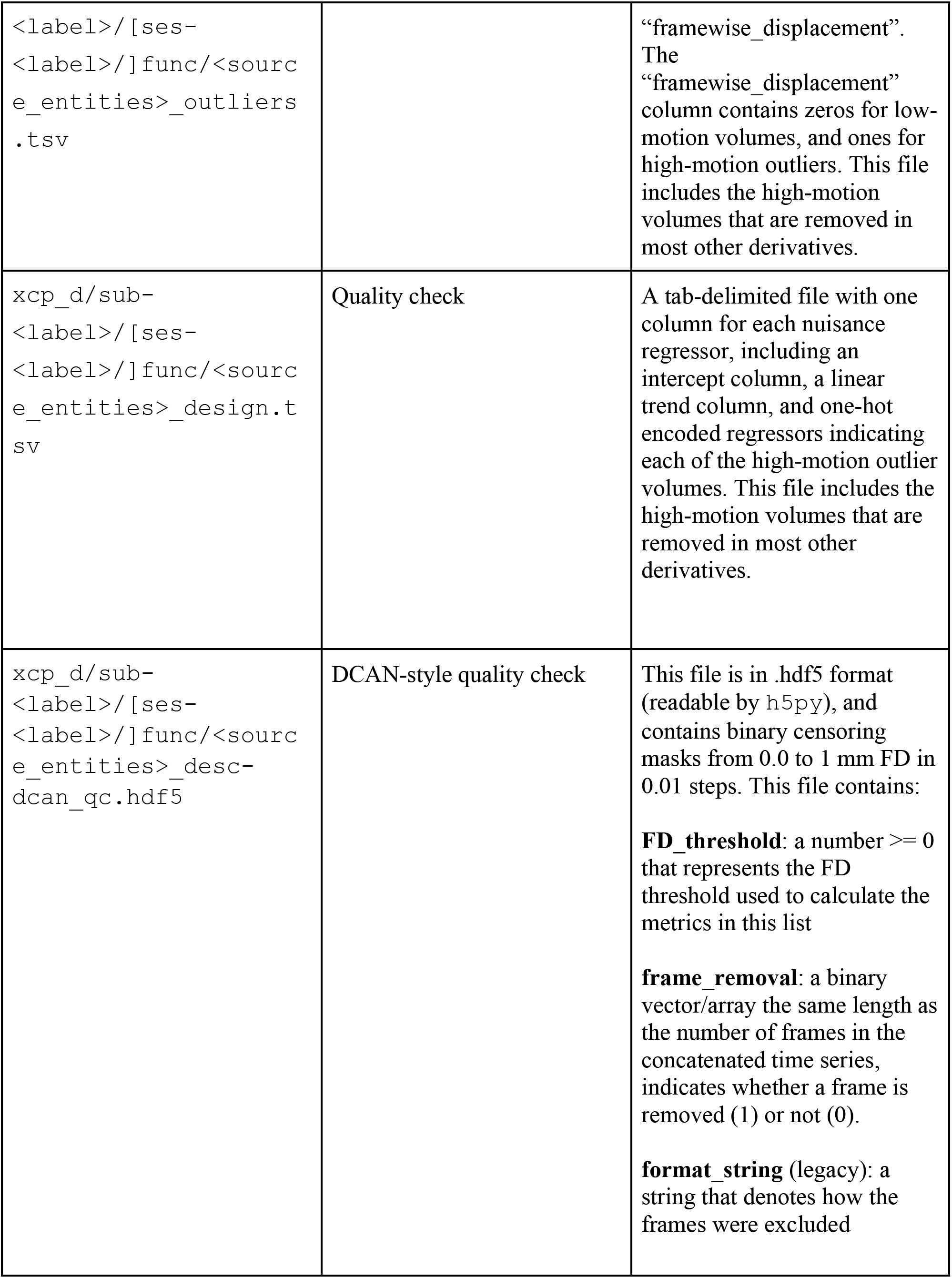
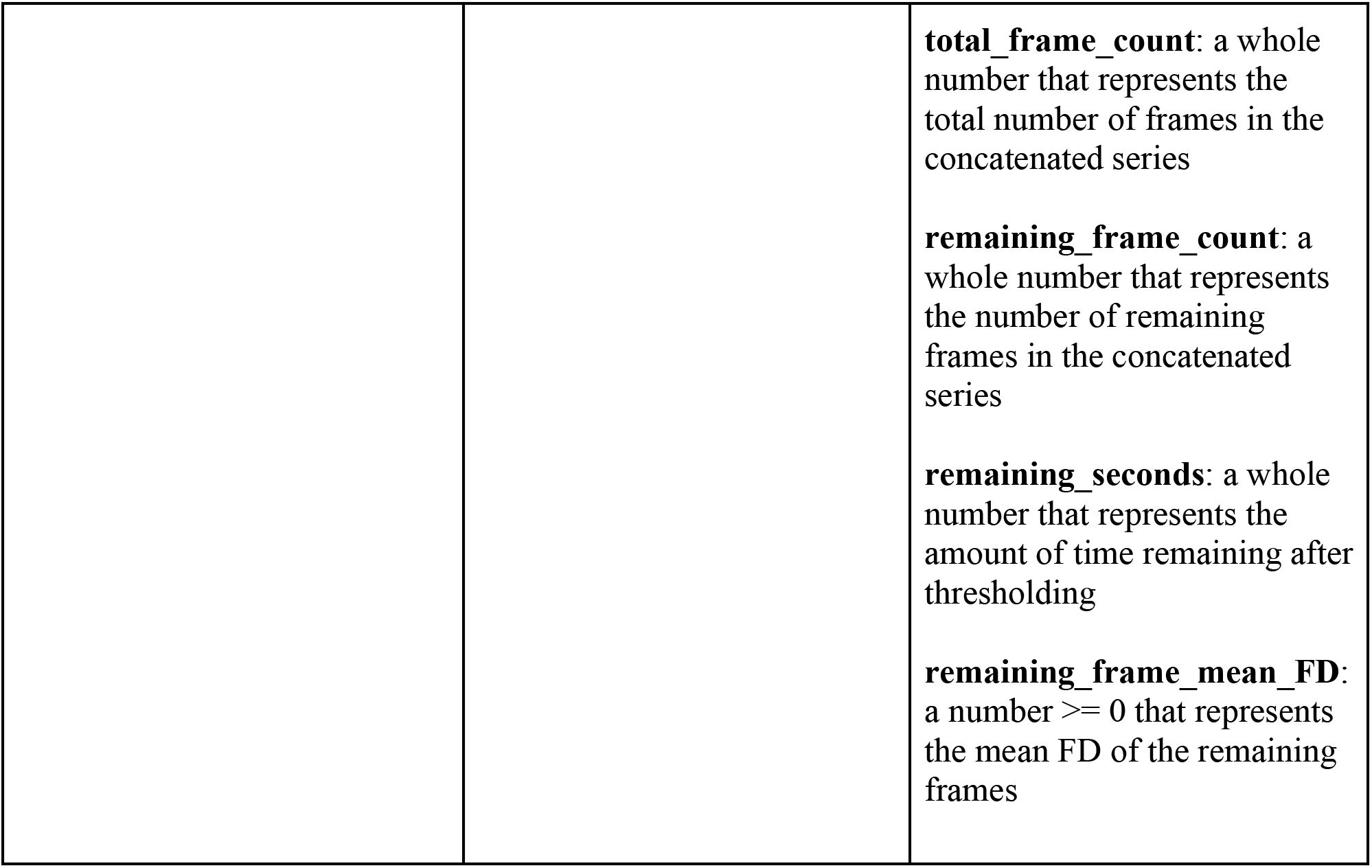
This table describes outputs from a run of XCP-D.

**Table 3:**
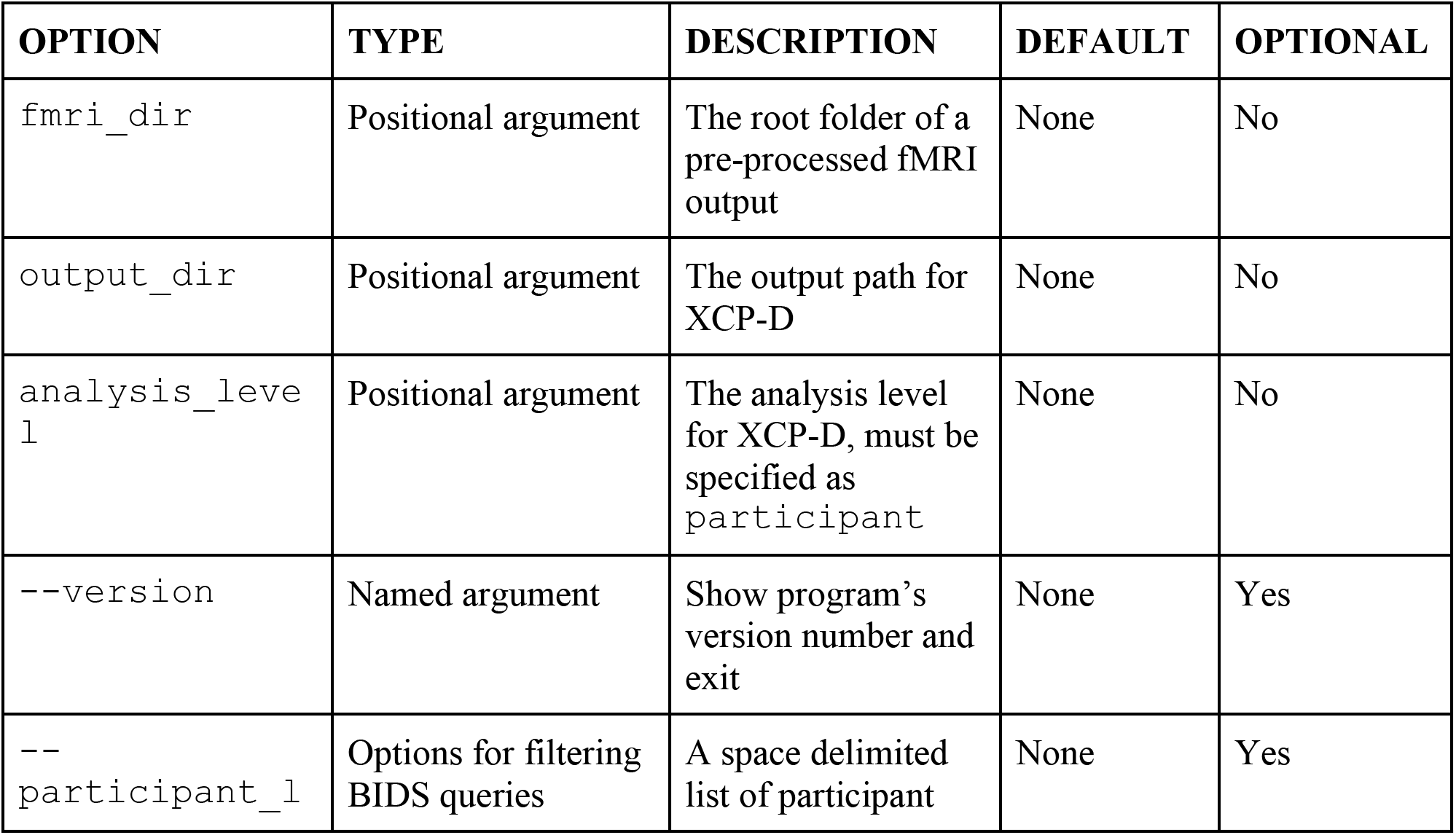
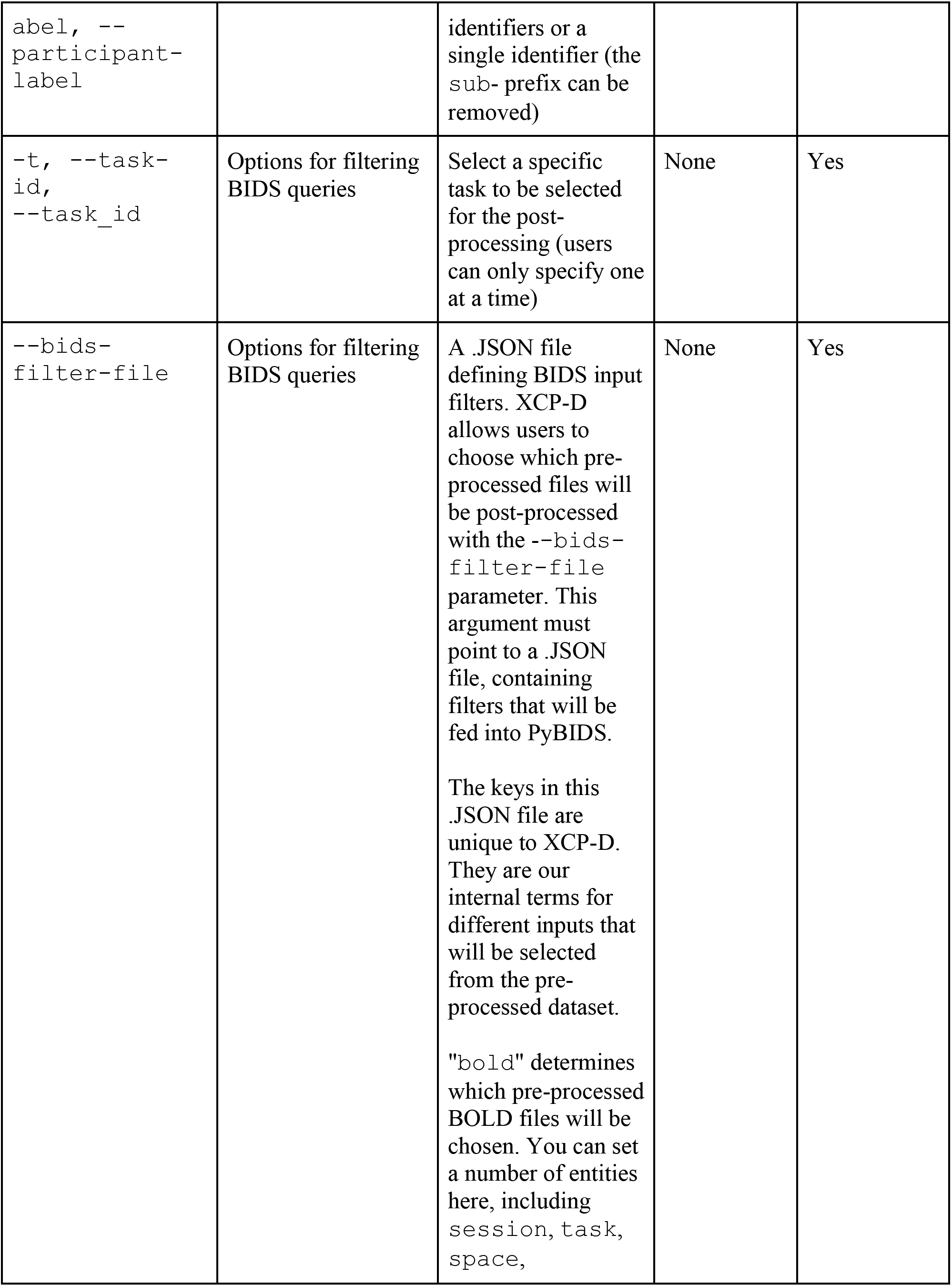
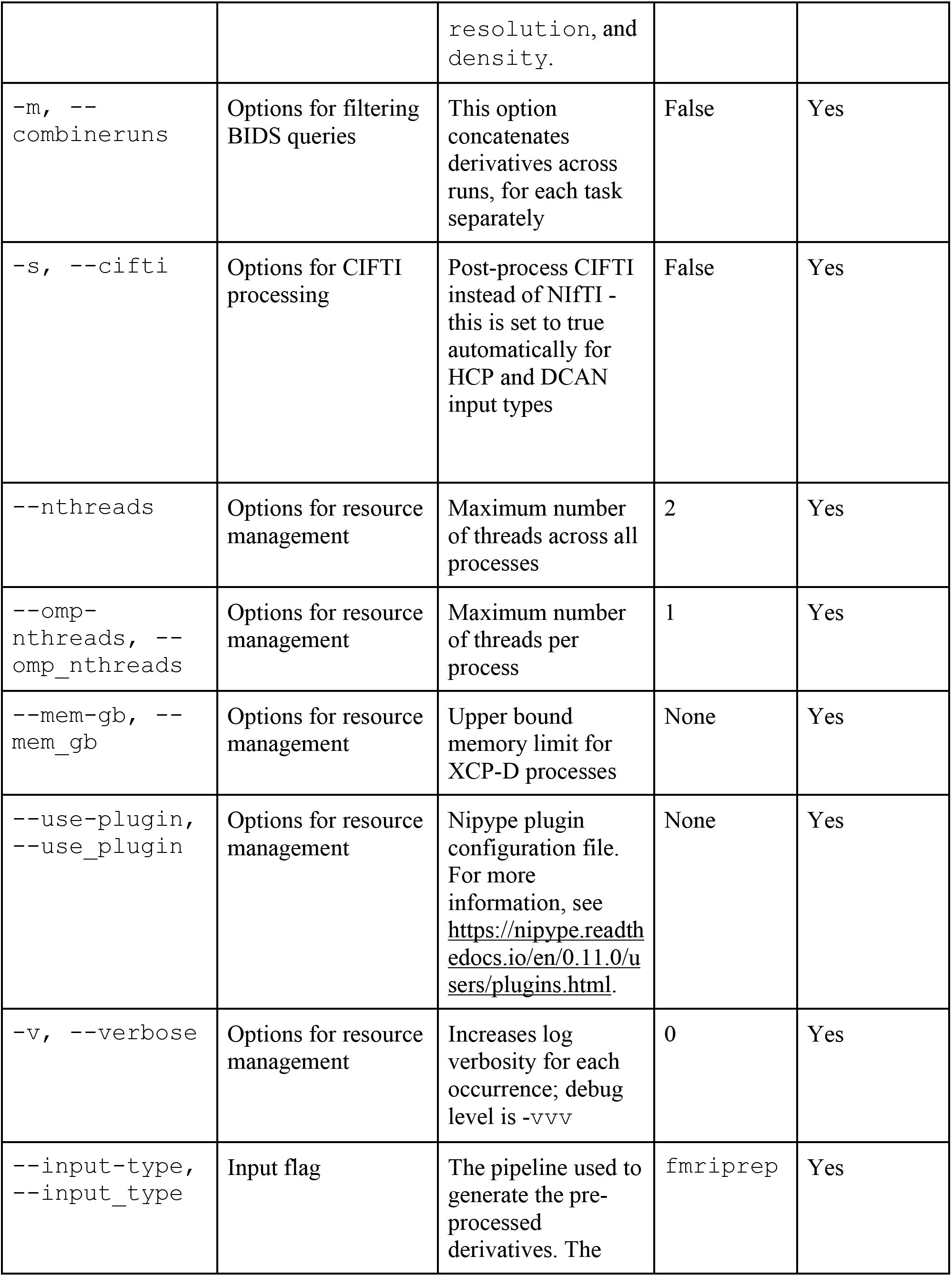
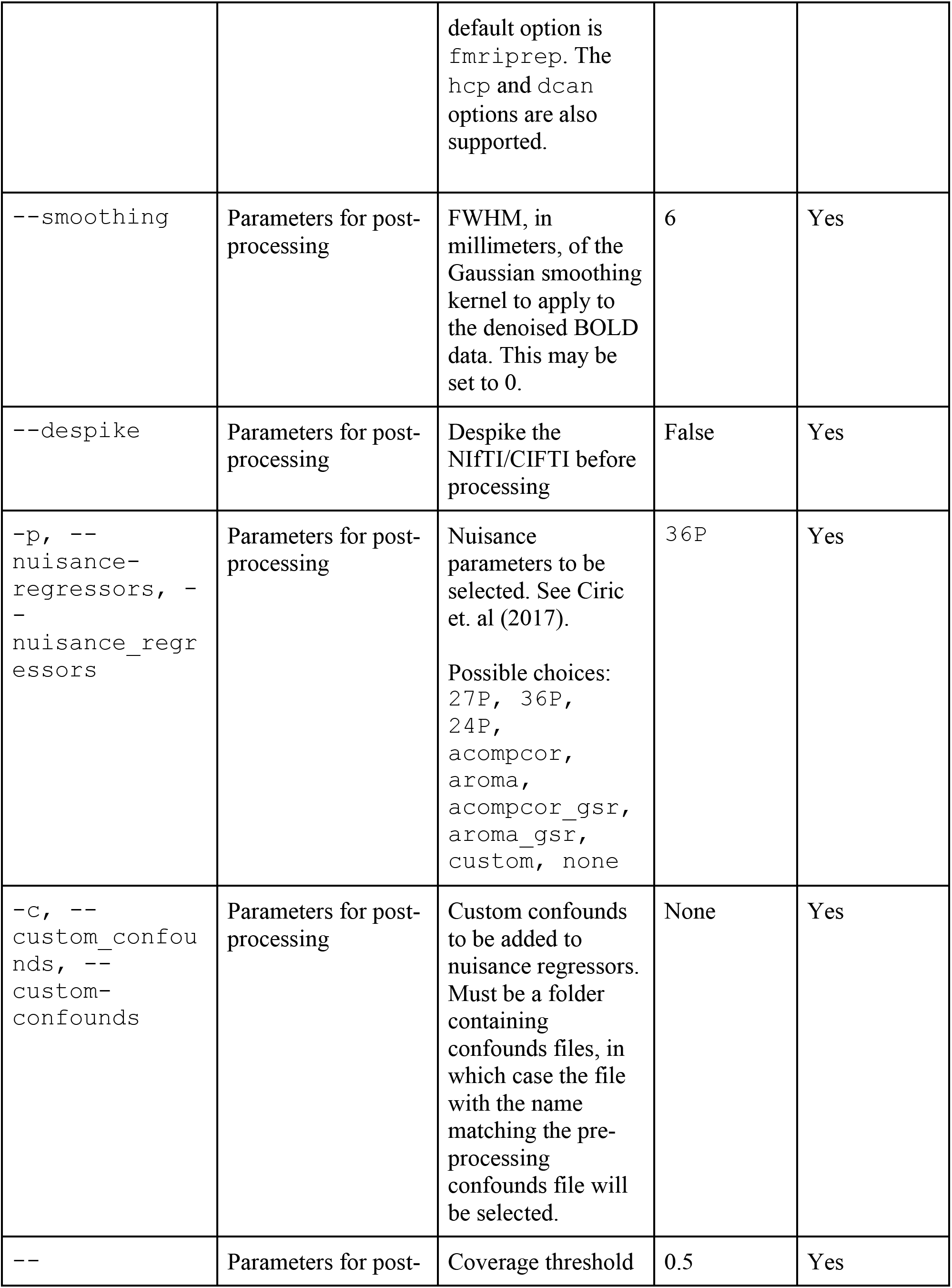
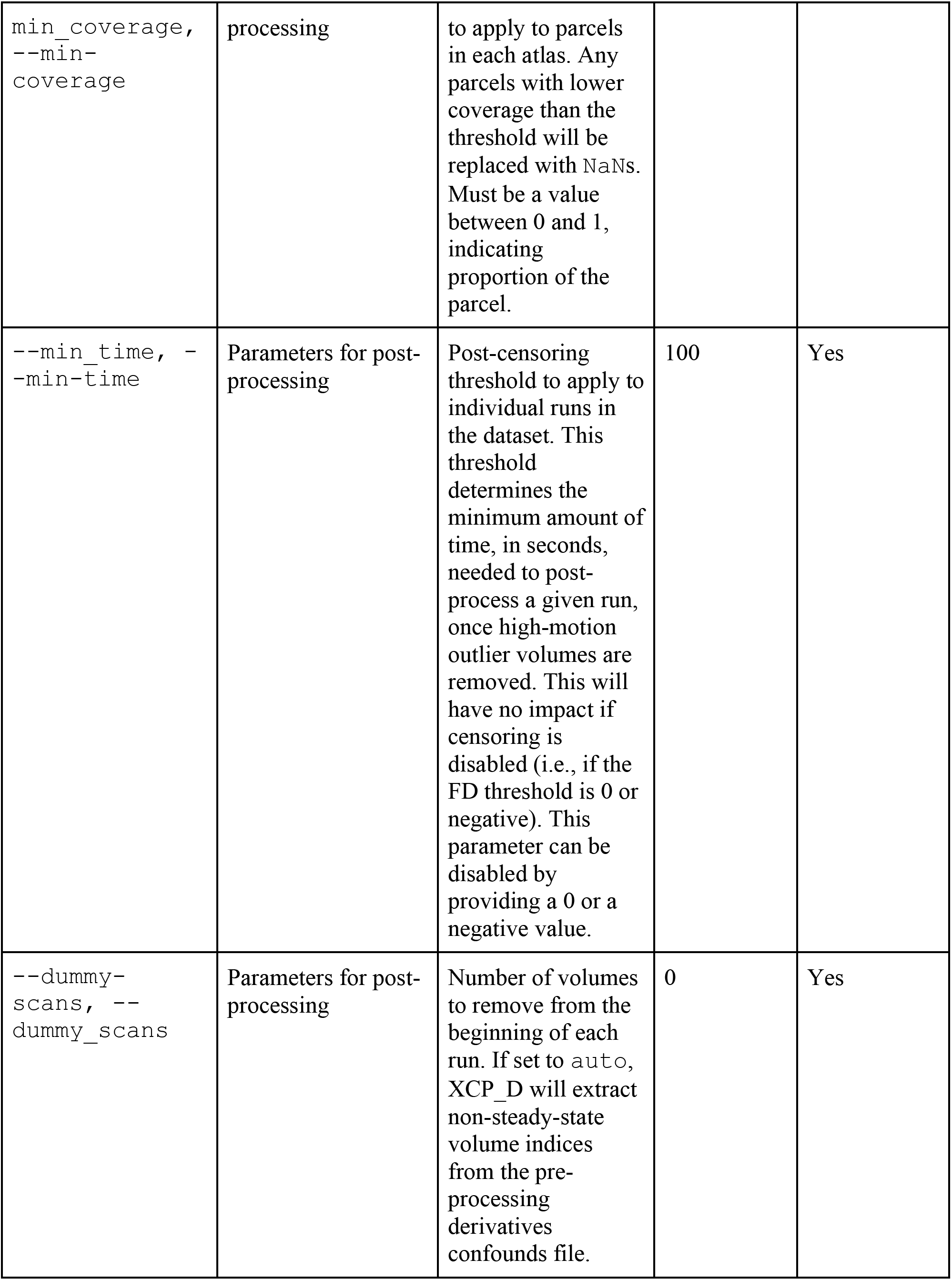
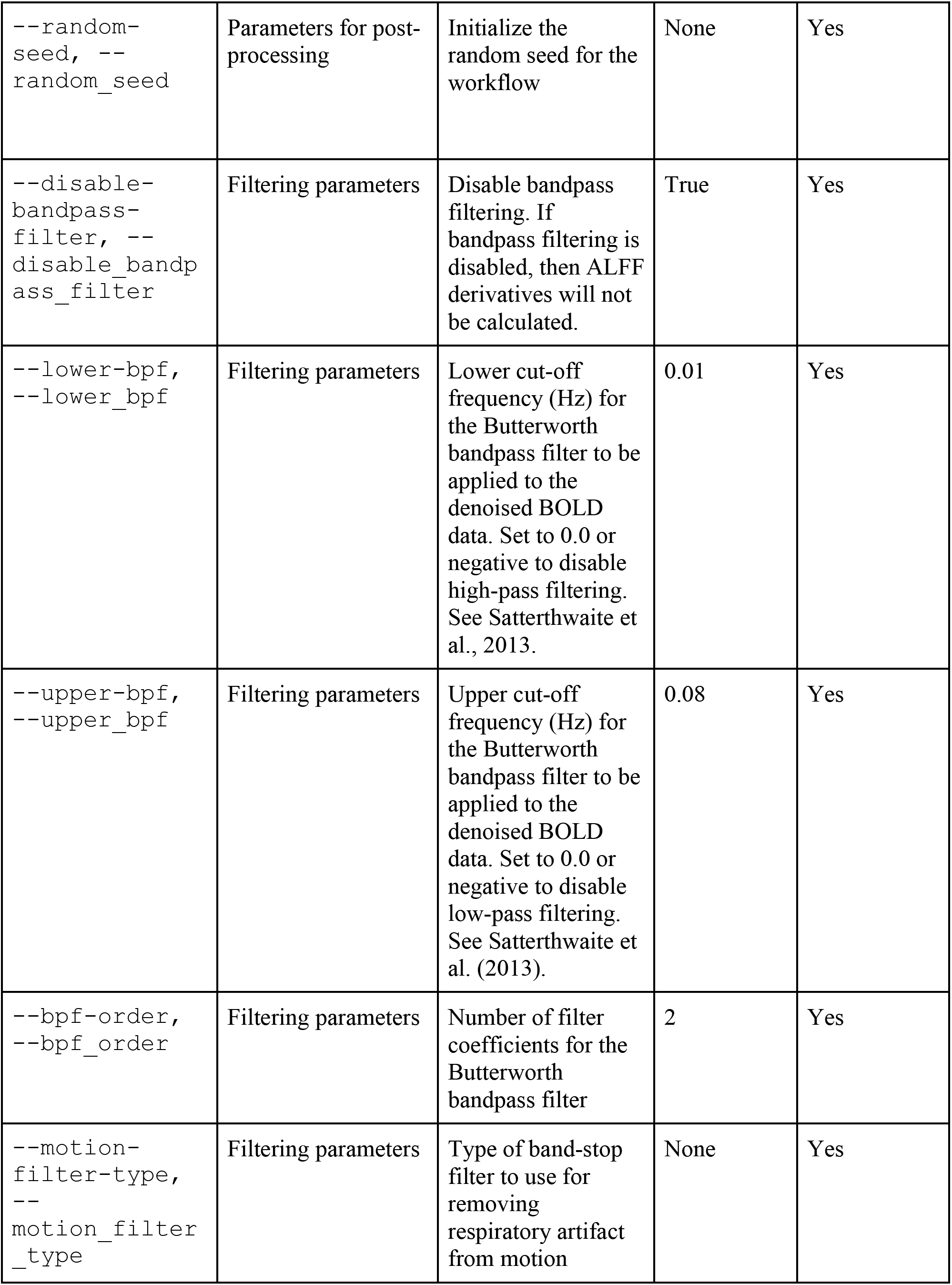
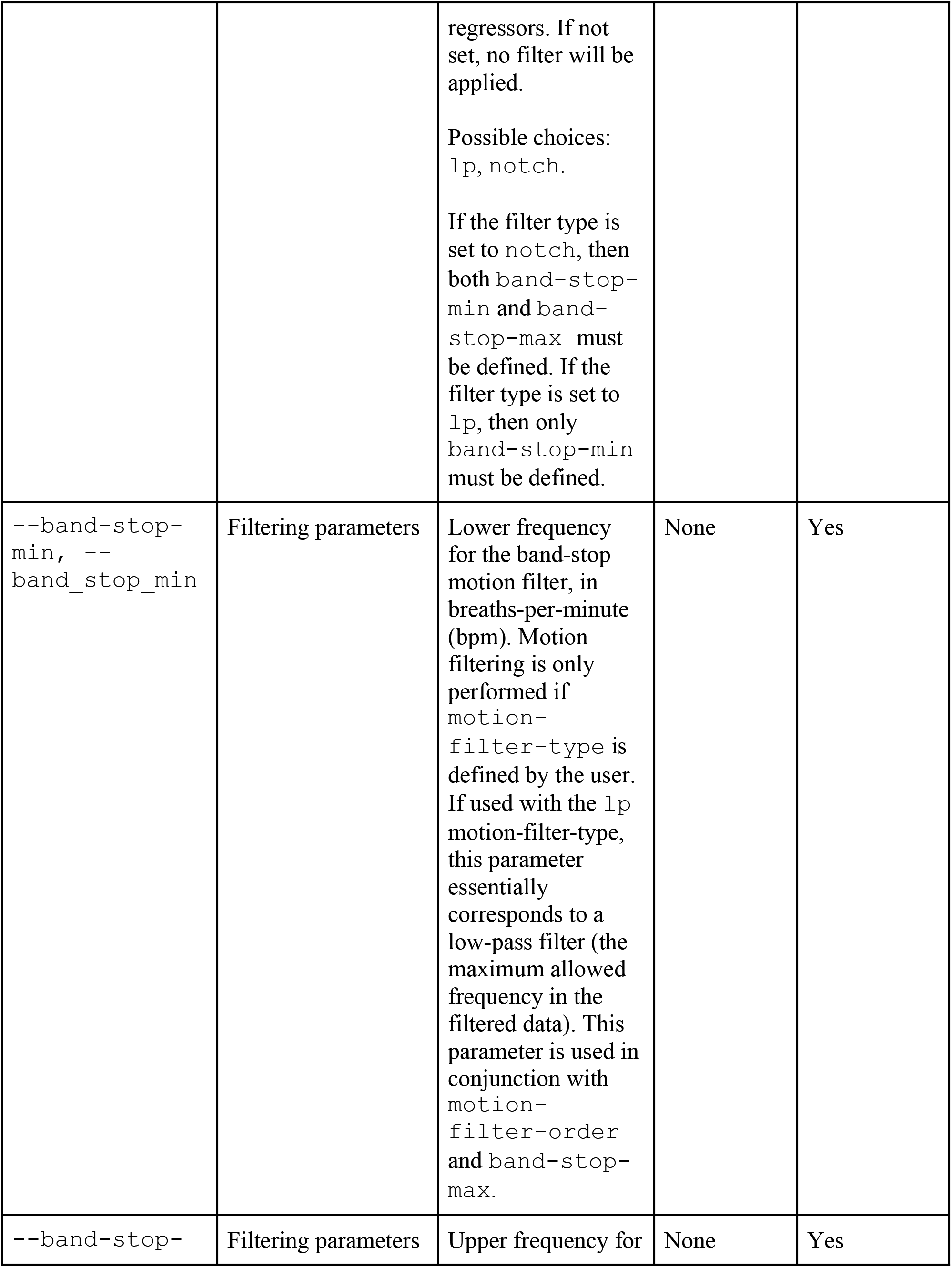
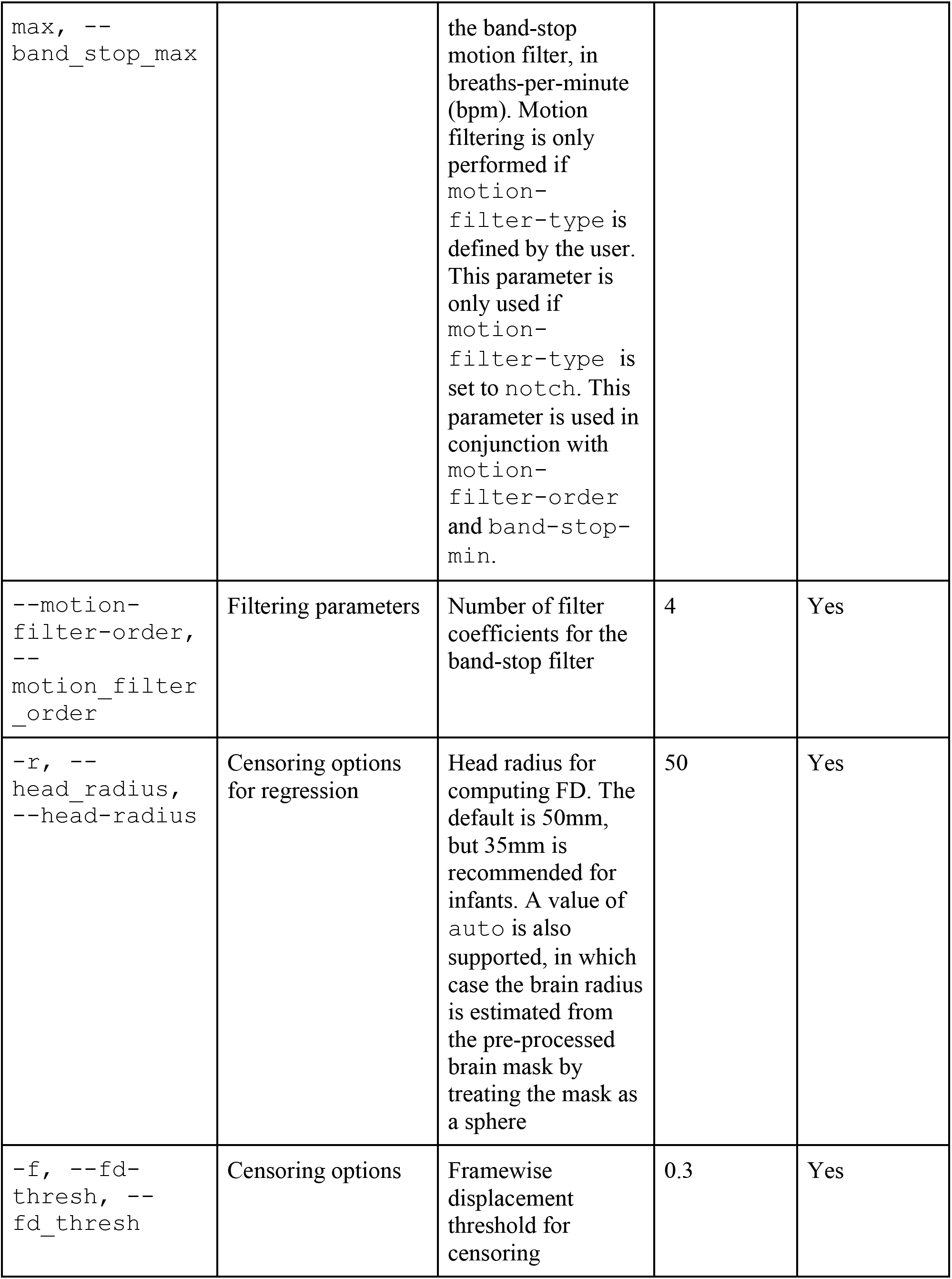
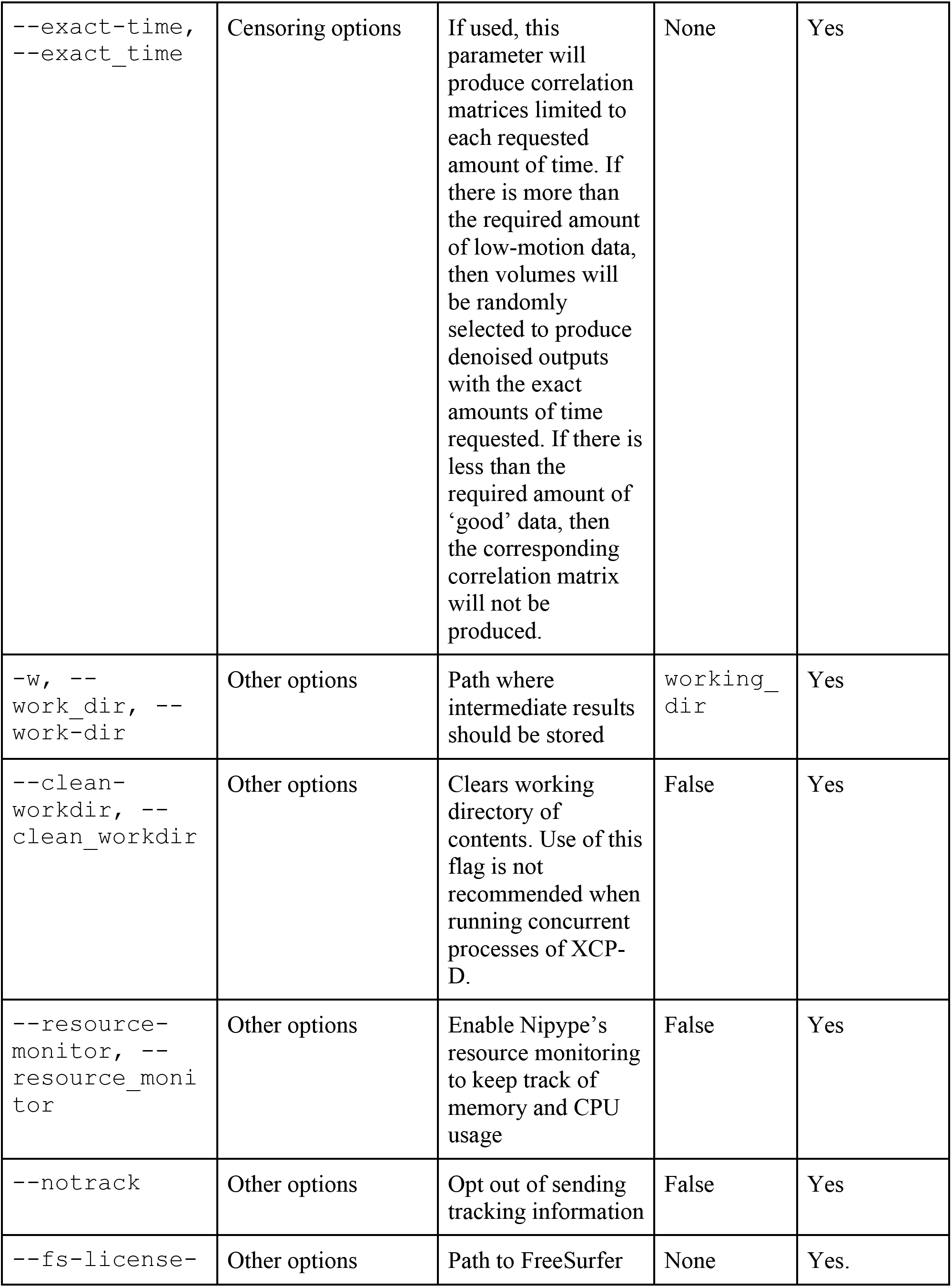
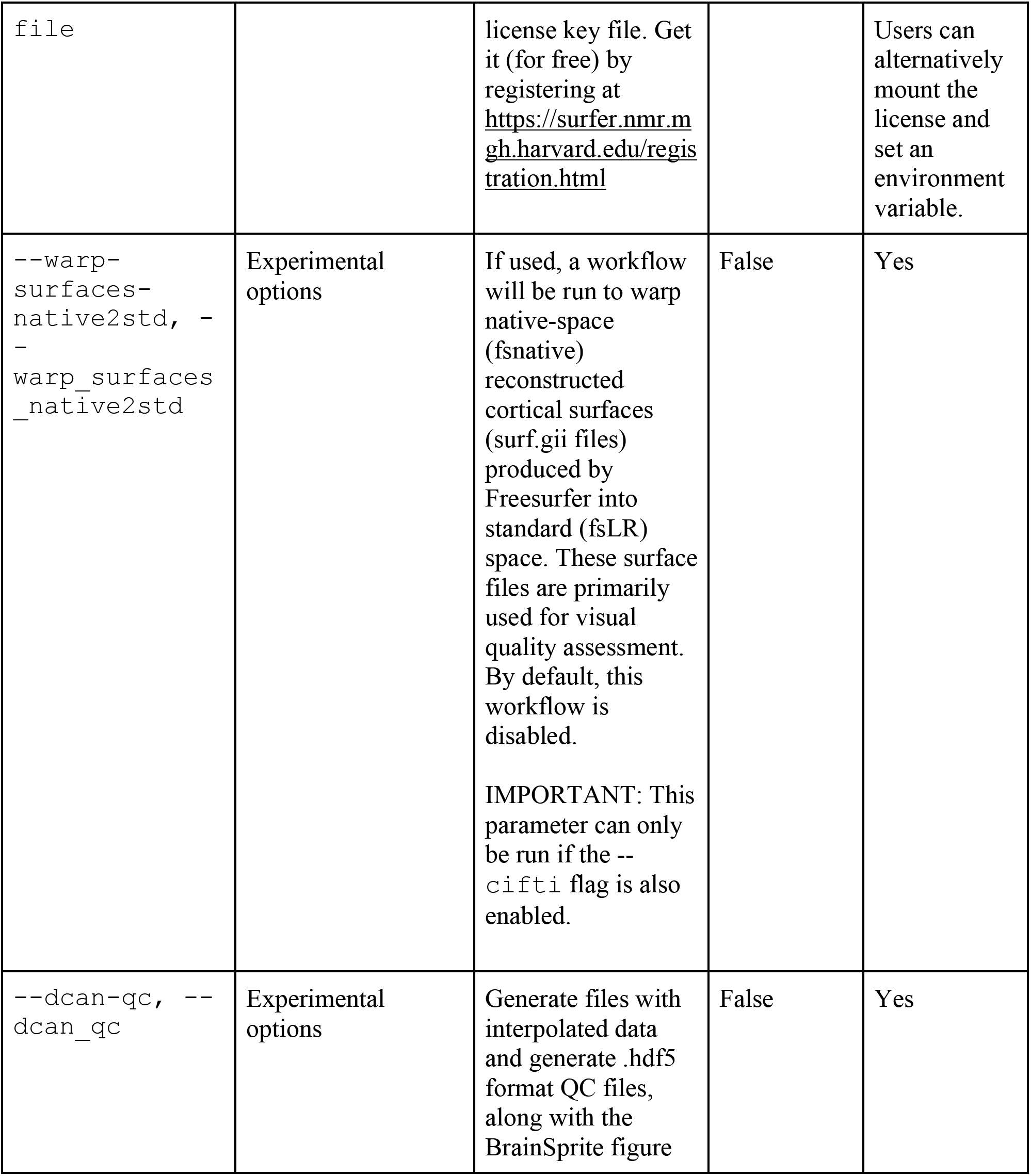
XCP-D command-line options.

Many internal operations of the software use TemplateFlow (Ciric et al. 2022), Nibabel (Brett et al. 2023), numpy (Harris et al. 2020), and scipy (Virtanen et al. 2020). Below, we describe each of the post-processing modules with accompanying command syntax, relevant information, as well as the CI tests for each module.

### Ingression of non-BIDS derivatives

XCP-D supports both BIDS derivatives-compliant pre-processing pipelines (i.e., fMRIPrep) and non-BIDS pipelines (i.e., HCP and ABCD-BIDS). In the latter case, XCP-D indexes the outputs from the pre-processing pipeline and maps the relevant files into a BIDS derivatives-compliant structure in the working directory if the user specifies `--input-type` as `dcan` or `hcp`.

As part of this ingression procedure, XCP-D also extracts minimal confounds. However, this does not fully reproduce the confounds that fMRIPrep creates, which limits the denoising strategies available for these data. Additionally, XCP-D’s anatomical workflow requires that CIFTI surfaces are in fsLR space at 32k density.

### Removal of non-steady state volumes

Some vendors acquire multiple additional volumes at the beginning of a scan to reduce transient T1 signals before a steady state is approached (Jenista et al., 2016). These volumes are often referred to as “dummy scans’’ or “non-steady state volumes’’. Additionally, higher levels of movement at the start of a scan (e.g., startle due to onset of scanner noise) may also lead investigators to remove initial volumes. This is the first post-processing step in XCP-D and occurs optionally. XCP-D allows the first *n* (as supplied by users) number of volumes to be deleted before processing. If set to `auto`, XCP-D will extract non-steady-state volume indices from the pre-processing derivatives confounds file (only included in fMRIPrep confounds files). Removal of dummy volumes is enabled via the `–-dummy-scans` flag and feeds the truncated confounds and image files into the rest of the workflow. This module is tested by evaluating a BOLD file and its corresponding confounds file and specifying a varying number of volumes (1-10) to be removed. The CI test confirms that the correct number of volumes is dropped from both the image and confound timeseries.

### Despiking

Despiking is a process in which large spikes in the BOLD times series are truncated on an adaptive, voxel-specific basis. Despiking limits the amplitude of the large spikes but preserves the data points with an imputed reduced amplitude to minimize the effect of outliers. Notably, despiking is different from temporal censoring as it modifies rather than deletes data – despiking is also performed individually for each voxel whereas temporal censoring removes an entire volume. XCP-D performs despiking via AFNI’s (Cox et al., 1996) 3dDespike using default settings and the `–-NEW` flag, which uses a new fitting algorithm to despike the data rather than AFNI’s original L1 method, due to faster processing speed. For CIFTIs, which are first converted to NIfTIs and back during the despiking process via Connectome Workbench (Marcus et al., 2011), the `–-nomask` flag is used so that the entire volume is despiked. Despiking is performed when the `–-despike` flag is supplied. Despiking is executed before regression, censoring, and filtering to minimize the impact of spikes. Testing for this module involves calculating the maximum and minimum intensity values of the data and ensuring that the range between the two has decreased after despiking - that is, the minimum value of the data has increased, while the maximum value has decreased.

### Filtering of realignment parameters

Recent work has established that respiration can systematically induce fluctuations in the main magnetic field (Fair et al., 2020), which can contaminate estimates of head motion. Such artifacts can be removed via filtering of the realignment parameters using a low-pass filter for single-band images (Gratton et al., 2020) or a notch filter for multiband images (Fair et al., 2020). If users specify a low-pass filter, frequencies above `band_stop_min` (specified in breaths per minute) are removed with a Butterworth filter. If users specify a notch filter (as described in Fair et al., 2020), the frequencies between `band_stop_min` and `band_stop_max` are removed. The notch filter is applied using scipy’s `iirnotch` function, and both filters are applied backwards and forwards using scipy’s `filtfilt` function. Motion parameter filtering will only be enabled if `--motion-filter-type` is provided.

### Temporal censoring

Temporal censoring (also known as motion scrubbing) is a process in which data points with excessive motion are removed from the fMRI timeseries (Power et al., 2012). To aid the fit of the confound regression model, censored data points are removed before regression. The framewise displacement (FD) threshold specified by the user (with a default value of 0.3) is used to identify volumes to be censored. Temporal censoring can be disabled by setting `–-fd-thresh` to 0.

FD is calculated from the (optionally filtered) realignment parameters following the procedure described in Power et al., 2014. The head radius used to calculate FD may be supplied by the user via `–-head-radius`, set to `auto` (which estimates the brain radius based on the pre-processed brain mask), or by defaulting to 50 mm. The FD timeseries and FD threshold are then used to determine the number of high motion outlier volumes. A temporal mask is then generated in .tsv format, with 0s corresponding to volumes that were not flagged for censoring, and 1s indicating high-motion outlier volumes.

For participants with high motion, it is possible that censoring results in a timeseries with few un-censored volumes. XCP-D allows the user to specify a minimum run duration (in seconds) of un-censored data. This minimum time can be specified by the user via `–-min-time` (with a default value of 100, in seconds), which determines the minimum amount of time, in seconds, needed to process a given run once high-motion volumes are removed. This feature can be disabled by providing a 0 or a negative value.

This module is tested by replacing values in the confounds file with values that should be censored and ensuring that the image file and the confounds file have had the same number of volumes dropped after the censoring module.

### Confound selection

Confound selection occurs when a confounds file is supplied from a pre-processing software. A custom confounds file may also accompany or replace this confounds file. The selected nuisance regressors could include realignment parameters, mean timeseries from anatomical compartments (GM, WM, CSF), the global signal (Fox et al., 2009), CompCor components (Behzadi et al., 2007), or independent components from ICA-AROMA (Pruim et al., 2015). Confound configurations can be extracted from these parameters and are then used to remove noise from the BOLD image file during confound regression. Confound configuration preferences may vary across use cases thus XCP-D allows users some flexibility in denoising options (Satterthwaite et al., 2013; Ciric et al., 2017). Note that at present, users cannot apply aCompCor or AROMA nuisance regressors for HCP or ABCD-BIDS inputs; this is a feature that may be added in the future.

The built-in nuisance strategies may be supplemented or replaced with a custom confounds file provided by the user. This functionality allows users to perform more advanced regression strategies. For example, users may convolve task regressors with a hemodynamic response function and provide these regressors in a custom confounds file to regress out task signals and treat the denoised data as pseudo-rest (Fair et al., 2007). If users wish to retain specific signals of interest in the data, they may include those signals in the custom confounds file, with the associated column headers prefixed with “signal ”. This scenario is described in “Confound regression”.

Confound selection is implemented via Nilearn’s (Abraham et al., 2014) `load_confound` functionality. The selected confounds are fed into the beginning of the workflow in .tsv format where dummy time is removed - so it is appropriately truncated, and then passed on throughout the workflow. Pre-configured confound strategies include those described in a prior benchmarking study (Ciric et al., 2018):

- `24P` - six realignment parameters, their squares, derivatives, and squares of the derivatives
- `27P` - the white matter, CSF and global signal parameters in addition to those included in the 24P model
- `36P` - the squares, derivatives, and squares of the derivatives of white matter, CSF and global signal parameters in addition to those included in the 27P model
- `acompcor` - the ACompCor parameters, the six realignment parameters, and their derivatives
- `acompcor_gsr` - the ACompCor parameters, the realignment parameters, their derivatives, and global signal
- `aroma` - the AROMA parameters, realignment parameters, their derivatives, white matter, and CSF
- `AROMA_gsr` - the AROMA parameters, realignment parameters, their derivatives, white matter, CSF, and global signal
- Custom confounds - users provide their own confounds

Confound parameters can be selected by the user via the `-p` flag and corresponding configuration, or `-c` for custom confounds. Nuisance regressors can also be specified as `none` to skip this denoising step. Confound selection is tested by ensuring that the confounds matrix for certain confound configurations and BOLD files have the right number of parameters - for example, 36 parameters if `36P` is selected as the confound configuration.

### Confound regression

Confound regression is used to mitigate motion artifacts in fMRI scans. XCP-D implements denoising via linear least squares regression. First, linear trend and intercept regressors are appended to the selected confounds so that the data is linearly detrended. Next, high-motion outlier volumes are removed from the nuisance regressors and the BOLD data (see section “*Temporal censoring*” above) so that the regression is only performed on low-motion data; the inclusion of very-high motion data that is removed via temporal censoring would reduce the effectiveness of confound regression. Each of the nuisance regressors, except for the intercept, are additionally mean-centered prior to the regression.

In some cases, the selected confounds may be correlated with signals of interest, as in AROMA, where ICA components are labeled as “noise” or “signal.” In these cases, including the “noise” regressors without modification can result in the removal of variance explained by “signal” regressors. To address this issue, XCP-D orthogonalizes all nuisance regressors (except for the linear trend and intercept regressors) with respect to any detected signal regressors. This is done automatically for nuisance regression strategies that include AROMA regressors. For custom confounds derived from spatial ICA components, such as multi-echo denoising with tedana (DuPre, Salo et al., 2021; Kundu et al., 2011; Kundu et al., 2013), users must include “signal” components in their custom confounds file, prefixed with “signal ”. When columns with this prefix are detected in the confounds file, XCP-D will automatically employ this orthogonalization procedure. Then, when the confound regression step is performed, the modified nuisance regressors (i.e., without the signal regressors) will be mean-centered, censored to remove high-motion volumes, and finally regressed out of the fMRI data.

Regression consumes the confounds file and BOLD file to be denoised and produces a residual timeseries for further analysis. Using the user-selected (see above) confounds, regression occurs after despiking and censoring. Confound regression is tested by confirming that the correlation between a random voxel and the confounds timeseries has decreased.

### Interpolation

For accurate bandpass filtering, the original sampling rate of the time series must be retained. Hence, interpolation restores the length of the original timeseries after temporal censoring. It occurs after regression, using the temporal mask generated during censoring to determine which values have been removed during censoring. Then, it uses Nilearn’s interpolation function to interpolate values from high-motion volumes via cubic spline interpolation. Note that interpolation for volumes at the beginning and end of the time series is disabled. Instead, XCP-D propagates the values from the closest low-motion volume. The BOLD timeseries with the interpolated values is then passed to the filtering workflow. Testing of this module involves confirming that the difference between the fast Fourier transform (FFT) of an interpolated file and the original file is less than the difference between the FFT of a file with an artificial spike planted in it and the original file.

### Filtering

Temporal filtering is used in fMRI signal processing to reduce high-frequency and low-frequency artifacts in the timeseries. XCP-D applies a Butterworth bandpass filter to BOLD signals after regression and interpolation. Functional connectivity between regions of interest is typically determined based on synchrony in low-frequency fluctuations (Biswal et al., 1995); therefore, removing higher frequencies using a low-pass filter may effectively remove noise from the timeseries while retaining signal of interest. High-pass filters can be used to remove very-low-frequency drift, which is a form of scanner noise, from an acquisition. Any frequencies below the low-pass cutoff and above the high-pass cutoff will be counted as pass-band frequencies as in the case of our Butterworth filter. These will be retained by the filter when it is applied. High-pass or low-pass only filtering is also supported.

The bandpass filter parameters are set from 0.01 to 0.08 Hz with a filter order of 2 by default, as used in Power et al., 2014. The filter is calculated using scipy’s `butter` functionality to calculate filtering coefficients, and `filtfilt` to apply the filter to the data. The filter uses constant padding with maximum allowed pad length as one less than the total number of volumes. Parameters can be modified in the command line, using the `–-lower-bpf, –-upper-bpf` and `–-bpf-order` flags. This module occurs after regression and before the creation of functional timeseries. It is applied to the unfiltered BOLD file and outputs the filtered image.

Testing of this module involves comparing the output of XCP-D’s Butterworth filtering code to the output of scipy’s code.

After bandpass filtering is performed, the denoised, interpolated, and filtered timeseries is re-censored, so that only low-motion volumes are retained. This occurs as described above in the “*Outlier detection and removal’’* section.

### Parcellated timeseries extraction and calculation of connectivity matrices

Functional connectivity matrices are a widely used approach to examine the coherence in activity between distant brain areas (Hlinka et al., 2011; Biswal et al., 1995). The generation of these matrices involves parcellating the brain into regions determined by atlases and then calculating correlations between regions.

XCP-D extracts voxel-wise timeseries from the denoised BOLD timeseries and outputs parcellated timeseries and correlation matrices for a variety of atlases bundles in the software. The output post-processed BOLD files, parcellated timeseries, and correlation matrices come from censored data. If the user adds the `–-dcan-qc` flag, then the interpolated version of the post-processed data will also be written out, with “desc-interpolated” in the timeseries filename. The local mean timeseries within each brain atlas’s region of interest (ROI) is extracted via Nilearn’s `NiftiLabelsMasker` for NIfTIs, and ConnectomeWorkbench’s `wb_command --cifti-parcellate` function for CIFTIs. Functional connectivity matrices are estimated using the Pearson correlation between all parcels for a given atlas. Before functional connectivity is estimated, a coverage threshold (with a default value of 0.5, or 50% coverage) is applied to parcels in each atlas. Any parcels with lower coverage than the threshold will be replaced with `NaN`s. This may be useful in the case of partial field-of-view acquisition or poor placing of the bounding box during acquisition. Additionally, if the `--exact-time` flag is used, this parameter will produce correlation matrices limited to each requested amount of time (specified in seconds). If there is more than the required amount of low-motion data, then volumes will be randomly selected to produce denoised outputs with the exact amounts of time requested. If there is less than the required amount of ‘good’ data, then the corresponding correlation matrix will not be produced.

The following atlases are implemented in XCP-D: Schaefer 100-1000 (Schaefer et al., 2018), Glasser 360 (Glasser et al., 2016), Gordon 333 (Gordon et al., 2016), the subcortical HCP Atlas (Glasser et al., 2013) and Tian Subcortical Atlas (Tian et al., 2020). Notably, our atlases have been harmonized with QSIPrep (Cieslak et al., 2021) and ASLPrep (Adebimpe et al., 2022) to facilitate multi-modal network analyses. This module is tested by confirming that the correlation coefficient of a parcellated timeseries is the same as in the connectivity matrix produced, when calculated separately in a Python notebook.

### ReHo

Regional Homogeneity (ReHo) is a measure of local temporal uniformity in the BOLD signal computed at each voxel of the processed image. Greater ReHo values correspond to greater synchrony among BOLD activity patterns measured in a local neighborhood of voxels (Zang et al., 2004), with neighborhood size determined by a user-specified radius of voxels. ReHo is calculated as the coefficient of concordance among all voxels in a sphere centered on the target voxel (Zuo et al., 2013).

ReHo is performed on the BOLD file after temporal filtering and the output is written out directly to the XCP-D derivatives folder. For NIfTIs, ReHo is always calculated via AFNI’s `3dReho` with 27 voxels in each neighborhood. For CIFTIs, the left and right hemisphere are extracted into GIFTI format via Connectome Workbench’s `CIFTISeparateMetric`. Next, the mesh adjacency matrix is obtained, and Kendall’s coefficient of concordance (KCC) is calculated (Zhang et al., 2023), with each vertex having four neighbors. For subcortical voxels in the CIFTIs, `3dReho` is used with the same parameters that are used for NIfTIs. This module is tested by adding artificial noise to an image and confirming that the mean ReHo value declines.

### ALFF

The amplitude of low-frequency fluctuations (ALFF) – also called “fluctuation amplitude” – is a measure of regional intensity of BOLD signal fluctuation (Yu-Feng et al., 2006) calculated in each voxel of the processed image. Low-frequency fluctuations are of particular importance because functional connectivity is most typically computed based on synchronous, low frequency fluctuations (Zou et al. 2008).

ALFF is calculated on the BOLD file after filtering and its output can optionally be smoothed (see *Smoothing*). Notably, ALFF is <u>only</u> calculated if bandpass filtering is applied, and motion censoring is disabled. ALFF is computed by transforming the processed BOLD timeseries to the frequency domain using scipy’s `periodogram` function. The power spectrum is computed within the default 0.01-0.08 Hz frequency band (or the band-pass values optionally supplied by the user during filtering), and the mean square root of the power spectrum is calculated at each voxel to yield voxel-wise ALFF measures.

This module is tested by first calculating the ALFF of a BOLD file. Then, the FFT of the BOLD file is calculated. After adding values to the amplitude of its lower frequencies, it is confirmed that the ALFF increases upon being re-computed.

### Spatial smoothing

Noise in the BOLD signal – due to physiological signals or scanner noise – can introduce spurious artifacts in individual voxels (Mikl et al., 2008). The effects of noise-related artifacts can be mitigated by spatial smoothing of the data, which can dramatically increase the signal-to-noise ratio (Mikl et al., 2008). However, spatial smoothing is not without its costs: it effectively reduces volumetric resolution by blurring signals from adjacent voxels (Mikl et al., 2008).

Smoothing optionally occurs after temporal filtering. FWHM smoothing is implemented in XCP-D with a default value of 6.0 mm in volumes and surfaces. Additionally, ALFF maps are also smoothed if the `--smoothing` flag is specified by the user. First, the specified FWHM kernel (specified in mm) is converted to sigma (standard deviation). Smoothing for NIfTIs is performed via Nilearn’s `smooth_img` using a Gaussian filter. For CIFTIs, Connectome Workbench’s `wb_command --cifti-smoothing` is used to smooth each hemisphere and the subcortical volumetric data. This module is tested by confirming smoothness has increased after data has passed through the smoothing workflow, via AFNI for NIfTIs and via Connectome Workbench for CIFTIs.

### Quality control

XCP-D calculates multiple quality control measures. These include estimates of fMRI data quality before and after regression, as well as indices of co-registration and normalization quality. Selected metrics include the following:

● Summary measures of realignment parameters: mean FD, mean and maximum root-mean-square displacement (RMS). FD and RMS measure relative contributions of angular rotation and uniformity of motion effects across the brain (Yan et al., 2013).
● DVARS: DVARS is a whole brain measure of the temporal derivative (D) of image intensity computed by obtaining the root mean square variance across voxels (VARS; Goto et al., 2016.) As such it reflects time-varying signals and large values are often attributable to artifacts such as in-scanner motion.
● fMRI-T1/T2 co-registration quality: Because of the limited spatial resolution and reduced anatomic contrast of fMRI images compared to structural images, fMRI images are co-registered to the structural image prior to normalization to template space. Poor co-registration can thus impact normalization. XCP-D calculates the Dice similarity index (Dice, 1945), overlap coefficient, and Pearson correlation between the fMRI image and the T1 image (or T2 image) to determine the quality of the registration. The Dice index equals twice the number of voxels common to both images divided by the sum of the number of voxels in each image. The overlap coefficient (Vijaymeena & Kavitha, 2016) calculates the relative number of non-zero voxels in both images. The Pearson’s correlation measures the correlations between the voxels in both images.
● fMRI-Template normalization quality: Following co-registration, the fMRI image is normalized to template space by applying the warp calculated in registration of the structural image to the template (Jahn, 2022). XCP-D calculates the dice similarity index (Dice, 1945), overlap coefficient, and Pearson correlation coefficient to quantify the alignment of the fMRI image to the template.

### Visual reports

XCP-D produces two different user-friendly, interactive .html reports. The first (DCAN-style) output is called the “Executive Summary.” The Executive Summary is an interactive web page for quick visual inspection of structural and functional registration, surface quality, physiological and non-physiological artifacts, and post-processing success (see select elements in **Figure 2**; a full example is provided in **Supplemental Figure 1**). It is particularly useful for assessing co-registration, normalization, and surface alignment. For example, it includes an interactive BrainSprite (https://github.com/brainsprite/brainsprite) viewer that overlays pial and white matter surfaces on the template image. This allows users to quickly assess the quality of the surface registration. Further information regarding co-registration and normalization quality is depicted in contour plots. The Executive Summary also includes a carpet plot for all runs, which depicts the fMRI timeseries before and after confound regression. These carpet plots are displayed alongside the FD plots and DVARS timeseries to allow users to rapidly assess denoising success. Additionally, XCP-D also provides a NiPreps style report that depicts similar information in a different layout (see **Supplemental Figure 2**). In both reports, XCP-D also produces a “methods boilerplate” that details the methods applied along with citations as relevant for users. This automatically generated description of the methods ensures fidelity of reporting and can be directly copied into publications’ methods sections.

**Figure 2:**
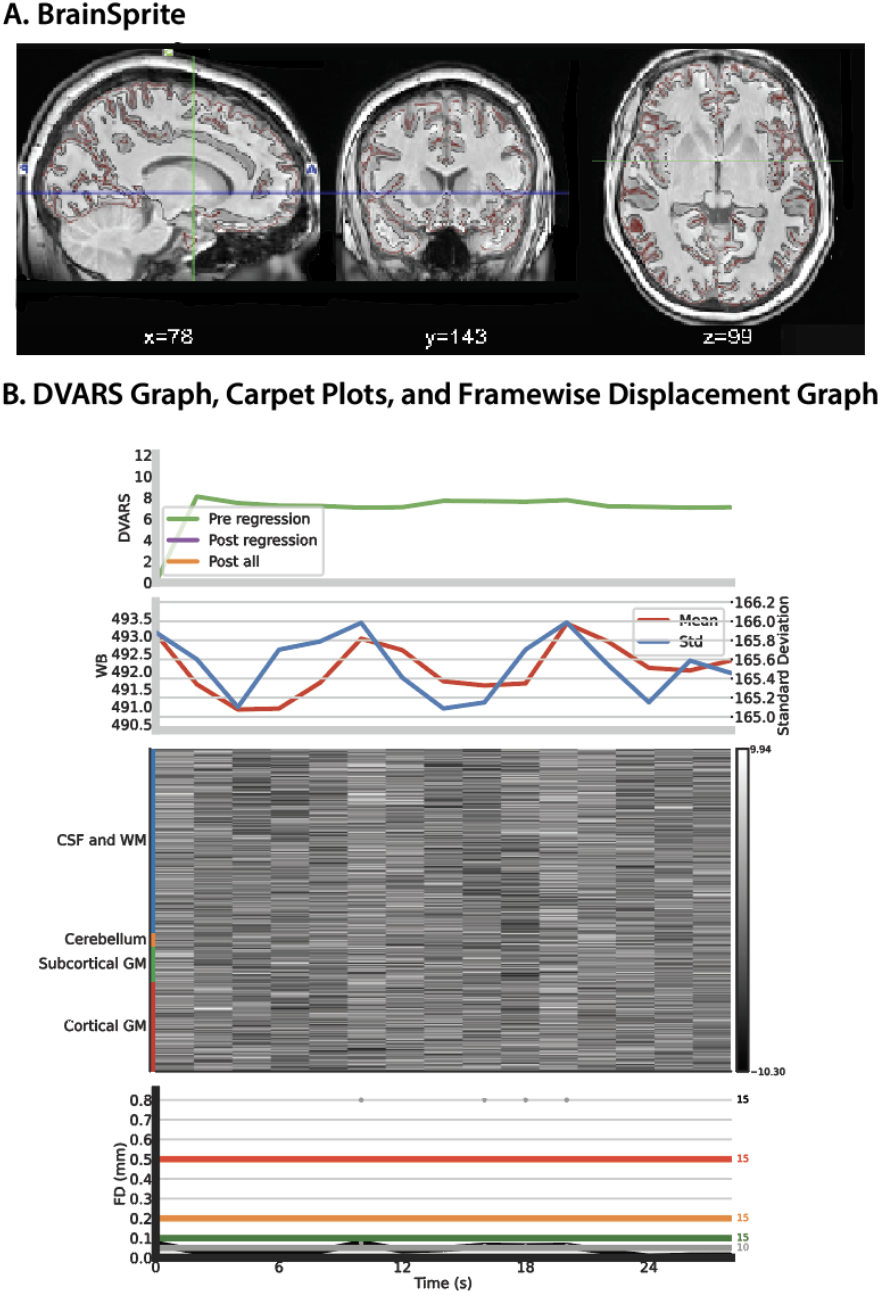
Selected elements of the XCP-D Executive Summary. Panel A depicts the BrainSprite viewer that overlays white and pial matter on the template, followed by (Panel B) a carpet plot and graphs depicting FD and DVARS. *FD: Framewise displacement; DVARS: temporal derivative (D) of image intensity computed by obtaining the root mean square variance across voxels (VARS)*.

### Anatomical Workflow

The optional anatomical workflow in XCP-D serves two main purposes. First, it is used to warp several surfaces derived from the structural images from fsnative to fsLR space, which is useful as part of the visual reports for assessing normalization to the fsLR template. To this end, the workflow generates surf.gii files in fsLR space for the gray matter / pial matter border and the white matter/gray matter border. It also generates HCP-style inflated surfaces for visualization purposes. The workflow can be enabled via the `–-warp-surfaces-native2std` flag.

Second, XCP-D will parcellate morphometric surface files – including cortical thickness, depth, and curvature – generated in pre-processing by sMRIPrep (Esteban et al., 2020) or HCP pipelines. XCP-D parcellates these morphometric files using the same atlases that are used for creating functional connectivity matrices as well as other surface features like ALFF and ReHo. This functionality facilitates analyses of both fMRI and structural imaging features when data is processed using XCP-D.

### Concatenation

XCP-D also offers users the option of concatenating fully denoised timeseries across fMRI runs based on the `run` entity specified (notably, different tasks are not concatenated); this also yields QC metrics that are concatenated. Notably, this option should be used with some caution as it will double the size of output data in the derivatives folder. Users can concatenate runs by specifying the `–-combineruns` flag.

## RESULTS

Below, we demonstrate the utility of XCP-D in two ways. First, we provide a detailed walkthrough with bundled example data. Second, we apply it to data from three large-scale datasets.

### WALKTHROUGH

#### The XCP-D workflow for processing an fMRIPrep dataset (example subjects)

The following walkthrough details the workflow for post-processing a dataset using XCP-D on a HPC – specifically, a RedHat Enterprise Linux-based system, using Singularity. To do so, we use an example dataset that is bundled with the software within the container. This container contains three example subjects from a study on executive function, which is available on OpenNeuro at https://openneuro.org/datasets/ds004450. These subjects are organized in a BIDS-compatible manner with T1s, two resting-state runs, and corresponding field maps for the three subjects. Both .nii.gz and .json files are available for each of these scans, along with a dataset_description.json, and riper derivatives. For the purposes of this walkthrough, commands for a minimal XCP-D run will be demonstrated.

All commands are run in a directory named `XCPD_test`. The XCP-D walkthrough container with the bundled subjects can be downloaded via Singularity, by running the following bash script:

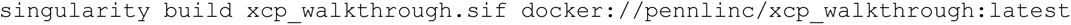

XCP-D can then be run on example subjects via Singularity, by running the following bash script:

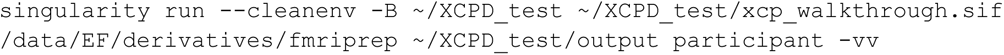

This script runs XCP-D using all the default options. The `--cleanenv` flags ensures that environment variables from local machines are ignored so that appropriate packages from within the container are used, and `-B` mounts the input files on local devices to the image. The three arguments here correspond to the mandatory arguments of: fmriprep directory (`/data/EF/derivatives/fmriprep`), output directory (∼/`XCPD_test/output)`, and analysis level (`participant`).

This will produce XCP-D derivatives under the folder `XCPD_test/output.` The outputs will include a dataset description, logs, citation information, processed anatomical and functional derivatives, as well as .svg figures. See **Supplemental Figure 3** for the expected directory structure of output from one example subject.

#### Application of XCP-D to three example datasets

To illustrate the utility of XCP-D to diverse data, we processed a total of 600 subjects from three datasets. Specifically, we processed n=200 participants each from the Philadelphia Neurodevelopmental Cohort (PNC; Satterthwaite et al., 2014; Satterthwaite et al., 2016), the Healthy Connectome Project - Young Adults (HCP-YA; Glasser et al., 2013) sample, and the Adolescent Brain Cognitive Development (ABCD; Volkow et al., 2017) study^®^. Note that the ABCD data repository grows and changes over time. The ABCD data used in this report came from https://doi.org/10.17605/OSF.IO/PSV5M.

Notably, prior to post-processing with XCP-D, each of these datasets were pre-processed using different tools. The PNC was processed using fMRIPrep (Esteban et al., 2018), ABCD was processed using ABCD-BIDS (Feczko et al., 2021), and the HCP-YA sample was processed via the HCP minimal processing pipelines (Glasser et al., 2013). All testing data had high quality structural images and greater than 5 minutes of high-quality resting-state fMRI data.

The following command was used to process the data (via the CIFTI surface-based workflow, with the anatomical workflow enabled):

### PNC

> singularity run –cleanenv -B ${PWD} –env FS_LICENSE=${PWD}/code/license.txt pennlinc-containers/.datalad/environments/xcp/image ${PWD}/inputs/data/fmriprep xcp participant –combineruns –nthreads 1 –omp-nthreads 1 –mem_gb 10 –smoothing 2 –min_coverage 0.5 – min_time 100 –dummy-scans auto –random-seed 0 –bpf-order 2 –lower-bpf 0.01 –upper-bpf 0.08 –motion-filter-type lp –band-stop-min 6 –motion-filter-order 4 –head-radius auto –exact-time 300 480 600 –despike –participant_label $subid -p 36P -f 0.3 –cifti –warp-surfaces-native2std –dcan-qc -w ${PWD}/.git/tmp/wkdir -vvv –input-type fmriprep

### ABCD

> singularity run –cleanenv -B ${PWD} pennlinc-containers/.datalad/environments/xcp/image inputs/data xcp participant – combineruns –nthreads 1 –omp-nthreads 1 –mem_gb 10 –smoothing 2 –min_coverage 0.5 –min_time 100 –dummy-scans 6 –random-seed 0 –bpf-order 2 -lower-bpf 0.01 –upper-bpf 0.08 –motion-filter-type notch –band-stop-min 15 –band-stop-max 25 –motion-filter-order 4 –head-radius auto –exact-time 300 480 600 –despike – participant_label $subid -p 36P -f 0.3 –cifti –warp-surfaces-native2std – dcan-qc -w ${PWD}/.git/tmp/wkdir -v –input-type dcan

### HCP-YA

> singularity run –cleanenv -B ${PWD} pennlinc-containers/.datalad/environments/xcp/image inputs/data xcp participant – combineruns –nthreads 1 –omp-nthreads 1 –mem_gb 10 –smoothing 2 –min_coverage 0.5 –min_time 100 –dummy-scans 7 –random-seed 0 –bpf-order 2 –lower-bpf 0.01 –upper-bpf 0.08 –motion-filter-type notch –band-stop-min 12 –band-stop-max 18 –motion-filter-order 4 –head-radius auto –exact-time 300 480 600 –despike – participant_label $subid -p 36P -f 0.3 –cifti –warp-surfaces-native2std – dcan-qc -w ${PWD}/.git/tmp/wkdir -v –input-type hcp

XCP-D completed successfully for all participants in all datasets. Among other outputs, XCP-D generated functional connectivity matrices (**Figure 3**) and parcellated cortical thickness information for each participant (**Figure 4**). Two small parcels in the medial temporal lobe cortex lacked coverage in the PNC. Notably, the correlation between the mean connectivity matrices is 0.93 for ABCD and PNC, 0.90 for ABCD and HCP, and 0.92 for PNC. The correlation between cortical thickness measures is 0.90 for ABCD and PNC, 0.95 for ABCD and HCP, and 0.85 for PNC and HCP.

**Figure 3:**
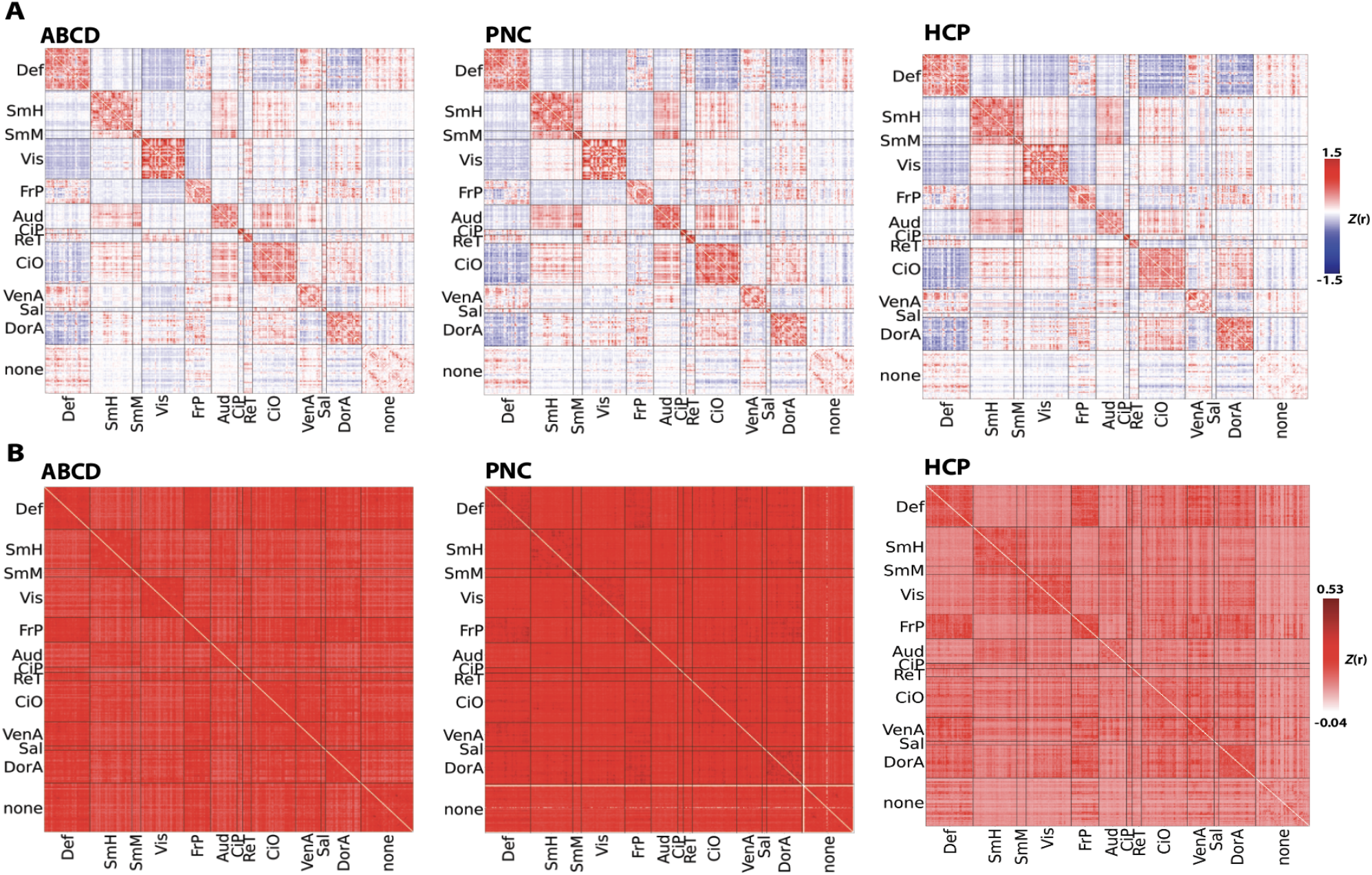
Mean (Panel A) and standard deviation (Panel B) functional connectivity generated by XCP-D for each dataset in our large-scale application, displayed after Fisher’s *Z* transformation. Data are displayed using the Gordon atlas (Gordon et al., 2016). *Def: default mode network; SmH: somatomotor hands network; SmM: somatomotor mouth network; Vis: visual network; FrP: Frontoparietal network; Aud: auditory network; CiP: cinguloparietal network; CiO: cingulo-opercular network; VenA: ventral attention network; Sal: salience network; DorA: dorsal attention network*

**Figure 4:**
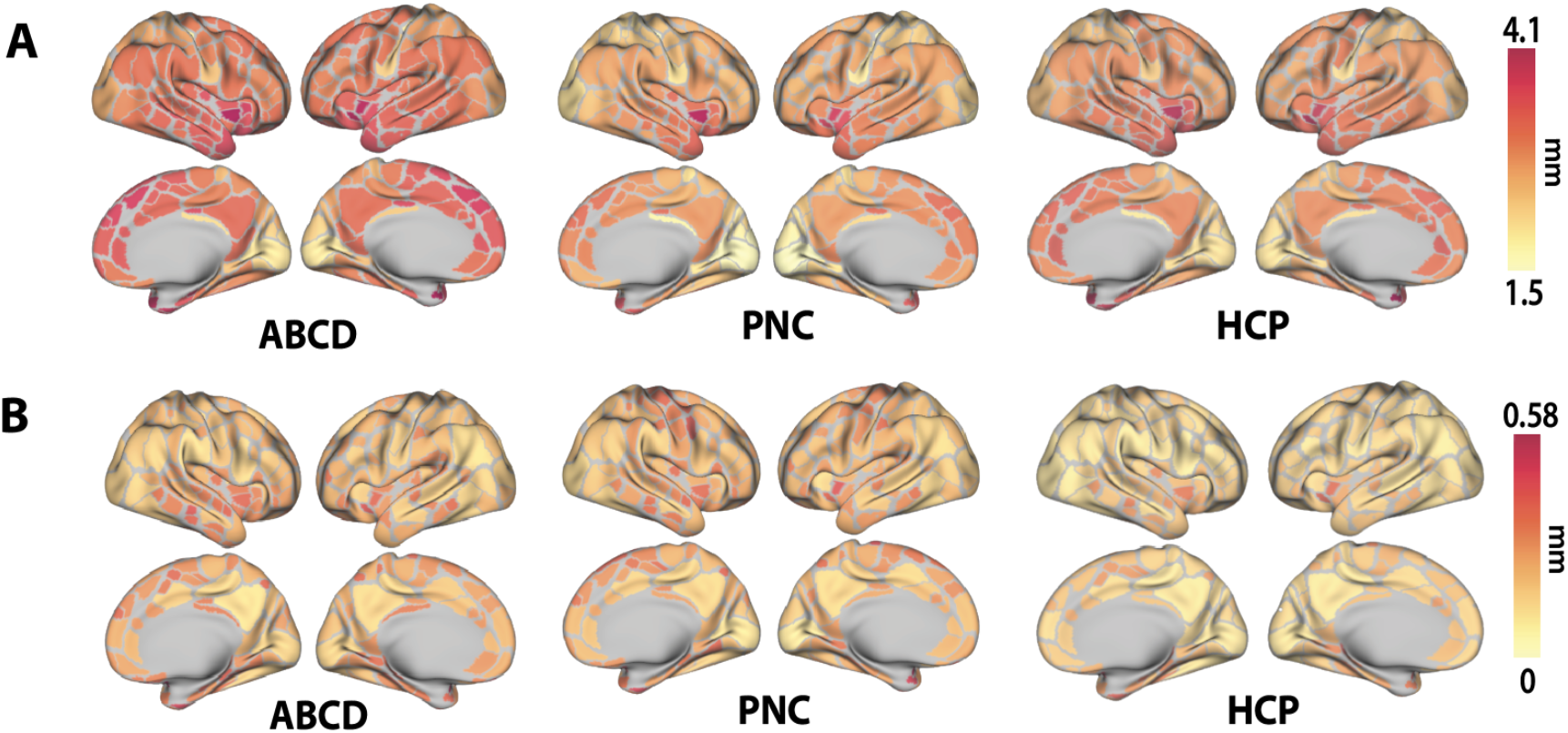
The mean (Panel A) and standard deviation (Panel B) of cortical thickness calculated by XCP-D of each dataset in our large-scale application. Data are displayed using the Gordon atlas (Gordon et al., 2016).

## DISCUSSION

Functional imaging is an essential tool for human neuroscience research. In contrast to pre-processing, where the field has gravitated towards use of standardized pipelines such as fMRIPrep, there has been a relative lack of standardization in fMRI post-processing. While several options for post-processing exist, they are often incompatible with common pre-processing methods, lack standardized output, and may not include software engineering best practices such as CI testing. While the steps used to generate the minimally pre-processed data are often quite similar, post-processing strategies used and derived measures often diverge substantially across data resources. XCP-D seeks to fill this gap and provide a post-processing workflow that is compatible with data pre-processed with several widely used strategies. XCP-D’s open and modular codebase in Nipype includes extensive CI testing, produces many measures of quality control, and yields analysis-ready derived measures that are named according to the BIDS standard. Together, XCP-D provides rigorous, accessible, and generalizable fMRI post-processing.

The derived measures generated by XCP-D include many of the most broadly used features of brain function and structure. Functional measures include connectivity matrices from multiple atlases as well as voxel- and vertex-wise maps of fluctuation amplitude and (ALFF) and regional homogeneity (ReHo). While XCP-D does not include extensive structural image processing or image registration, it does consume the structural features generated by pre-processing pipelines, rename them according to current BIDS standards, and apply the same parcellations used for the functional images. Summarizing functional and structural features in the many contemporary atlases included in XCP – including multi-scale atlases like the Schaefer parcellation (Schaefer et al., 2018) – facilitates multi-modal data integration and analysis. Multi-modal analyses are further accelerated by the recent integration of this same atlas bundle into our existing pipelines for diffusion MRI processing (QSIPrep; Cieslak et al., 2021) and processing arterial spin-labeled MRI (ASLPrep; Adebimpe et al., 2022) for calculation of cerebral blood flow.

Beyond such analysis-ready derived features, XCP-D produces an extensive set of quality control measures. These measures include indices of both image registration (e.g., Dice coefficient, overlap index) and denoising performance (e.g., the correlation of DVARS and motion before and after denoising). Together, such measures facilitate scalable quality assurance for large datasets and allow users to identify problematic datasets that can be further evaluated using the detailed reports generated for each participant. As part of our “glass-box” design philosophy, these single-participant reports allow users to examine key intermediate steps in the processing workflow. One particularly useful feature is the interactive BrainSprite that depicts the fully processed structural images along with overlays of the functional images. This visualization allows users to rapidly assess the success of image co-registration and atlas normalization. Additionally, the report includes tailored carpet plots that display the functional timeseries before and after post-processing, facilitating rapid visualization of artifacts related to in-scanner motion. Each participant’s report closes with an automatically generated boilerplate summary of the methods used by XCP-D for the configuration specified, along with relevant citations and references. This text enables users to determine if the desired processing occurred as expected and ensures accurate methods reporting.

In the design of XCP-D, we integrated multiple software engineering features to ensure stability and rigor. First, all XCP-D development is open, version-controlled, and clearly documented via detailed pull requests on GitHub. XCP-D implements branch protection rules that require reviews from at least one XCP-D developer before pull requests can be merged or changes can be released. We have benefitted from substantial community input and strive to quickly respond to bug reports from users. Second, XCP-D has a highly modular design in Nipype to reduce code duplication, enforce standardized workflows, facilitate integration testing, and allow for extensibility over time. Third, XCP-D is a BIDS-App, and we have made every effort to adhere to the standards described by BIDS, including the BIDS extension proposals (BEPs) related to derived data and functional networks. Fourth, XCP-D modules are subjected to extensive CI testing using CircleCI. These tests do not simply check that a file was produced but draw upon diverse example data and knowledge of each module’s operation to ensure that processing was executed correctly (for example, checking that a spike in the data is no longer present after despiking). These tests make the software more sustainable over time and mitigate risk of updates introducing occult errors. We track CI coverage using CodeCov; at present, 81% of the XCP-D codebase is covered by CI tests. Fifth and finally, XCP-D is containerized and distributed via Docker and Singularity, which wraps all dependencies to allow the software to be easily deployed in most computing environments.

There are many tools that denoise fMRI data, produce resting-state derivatives, and/or produce structural derivatives, including C-PAC (Configurable Pipeline for the Analysis of Connectomes; Craddock et al., 2013), CONN (Whitfield-Gabrieli & Nieto-Castanon, 2012), connectomemapper3 (Tourbier et al., 2022), CCS (Connectome Computation System, Xu et al., 2015), and DPARSF (Data Processing Assistant for Resting-State fMRI; Gan & Feng et al, 2010). One major difference between these tools and XCP-D is our dedicated focus on consuming data pre-processed by other widely used tools such as fMRIPrep. As such, XCP-D fills an important niche in the neuroimaging software ecosystem. Much of the post-processing that XCP-D provides can be performed using tools included in Nilearn, FSL, AFNI, and other software libraries. However, this would require users to assemble a pipeline themselves from component tools, and as such necessitate a higher degree of methodological proficiency. Furthermore, such user-assembled custom pipelines inevitably result in greater heterogeneity of methods used and usually reduce generalizability across efforts.

XCP-D has several limitations. Although XCP-D currently offers multiple denoising options, the range of denoising options described in the literature is vast and many are not currently supported. For example, XCP-D does not provide dedicated support for physiological confounds such as respiration or heart rate measures (Frederick et al., 2012), although these signals can be modeled as a “custom confound” supplied by the user. Similarly, we do not currently support denoising methods such as phase regression, which suppresses signal from large veins by removing the linear fit between magnitude and phase timeseries from the magnitude timeseries (Knudsen et al., 2023). Also, XCP-D cannot be used to analyze task data. Such functionality is provided by FitLins (Markiewicz, 2022), NiBetaSeries (Kent et al., 2020), and other packages. XCP-D also does not support group-level analyses.

These limitations notwithstanding, XCP-D provides generalizable, accessible, and robust post-processing for fMRI data. XCP-D’s ability to post-process data from ABCD-BIDS, HCP-YA and fMRIPrep allows the same denoising, confound regression, and generation of derivates for large-scale data resources that provide minimally pre-processed data; this could be invaluable for combining data across lifespan data resources. Moving forward, we plan to integrate additional advanced denoising methods, provide dedicated methods for handling physiological data, and extend the pre-processing data types supported to include infant data pre-processed using NiBabies. As an open-source, collaborative software package, we welcome bug reports, feature suggestions, pull requests, and contributions from the community.

## DISCLOSURES

Dr. Damien A. Fair is a co-founder, director, and equity holder of Turing Medical. Dr. Max Bertolero is an employee of Turing Medical.

## ACKNOWLEDGMENTS

This study was supported by grants from the National Institutes of Health: R37MH125829 (T.D.S. & DAF), R01MH112847 (T.D.S.), R01MH113550 (T.D.S. & D.S.B.), R01MH120482 (T.D.S.), R01MH112847 (T.D.S. & R.T.S), R01EB022573 (T.D.S.), DA041148 (D.A.F.), DA04112 (D.A.F.), MH115357 (D.A.F.), MH096773 (D.A.F.), MH122066 (D.A.F.), MH121276 (D.A.F.), MH124567 (D.A.F.), NS129521 (D.A.F.) and National Institute of Mental Health: 5T32MH019112-32 (A.S.K.). Additional support was provided by the AE Foundation and the Penn/CHOP Lifespan Brain Institute. Data used in the preparation of this article were obtained from the Adolescent Brain Cognitive Development (ABCD) Study (https://abcdstudy.org), held in the NIMH Data Archive (NDA). This is a multisite, longitudinal study designed to recruit more than 10,000 children aged 9-10 and follow them over 10 years into early adulthood. The ABCD Study is supported by the National Institutes of Health and additional federal partners under award numbers U01DA041048, U01DA050989, U01DA051016, U01DA041022, U01DA051018, U01DA051037, U01DA050987, U01DA041174, U01DA041106, U01DA041117, U01DA041028, U01DA041134, U01DA050988, U01DA051039, U01DA041156, U01DA041025, U01DA041120, U01DA051038, U01DA041148, U01DA041093, U01DA041089, U24DA041123, U24DA041147. A full list of supporters is available at https://abcdstudy.org/federal-partners.html. A listing of participating sites and a complete listing of the study investigators can be found at https://abcdstudy.org/consortium_members/. ABCD consortium investigators designed and implemented the study and/or provided data but did not necessarily participate in the analysis or writing of this report. This manuscript reflects the views of the authors and may not reflect the opinions or views of the NIH or ABCD consortium investigators.

