## Supplemental Figure 2 for "XCP-D: A Robust Pipeline for the post-processing of fMRI data"

### Summary

- Subject ID: 113922
- BOLD series: 18

### Processing Summary

#### Reports for: task EMOTION, direction LR.

##### Summary

- BOLD volume space: fsLR
- Repetition Time (TR): 0.72
- Mean Framewise Displacement: 0.1371
- Mean Relative RMS Motion: 0.1067
- Max Relative RMS Motion: 0.1848
- DVARS Before and After Processing : 480.9099, 23.3339
- Correlation between DVARS and FD Before and After Processing : 0.1961, 0.0396
- Number of Volumes Censored : 0

##### Alignment of functional and anatomical MRI data (surface driven)

bbregister was used to coregister functional and anatomical MRI data.

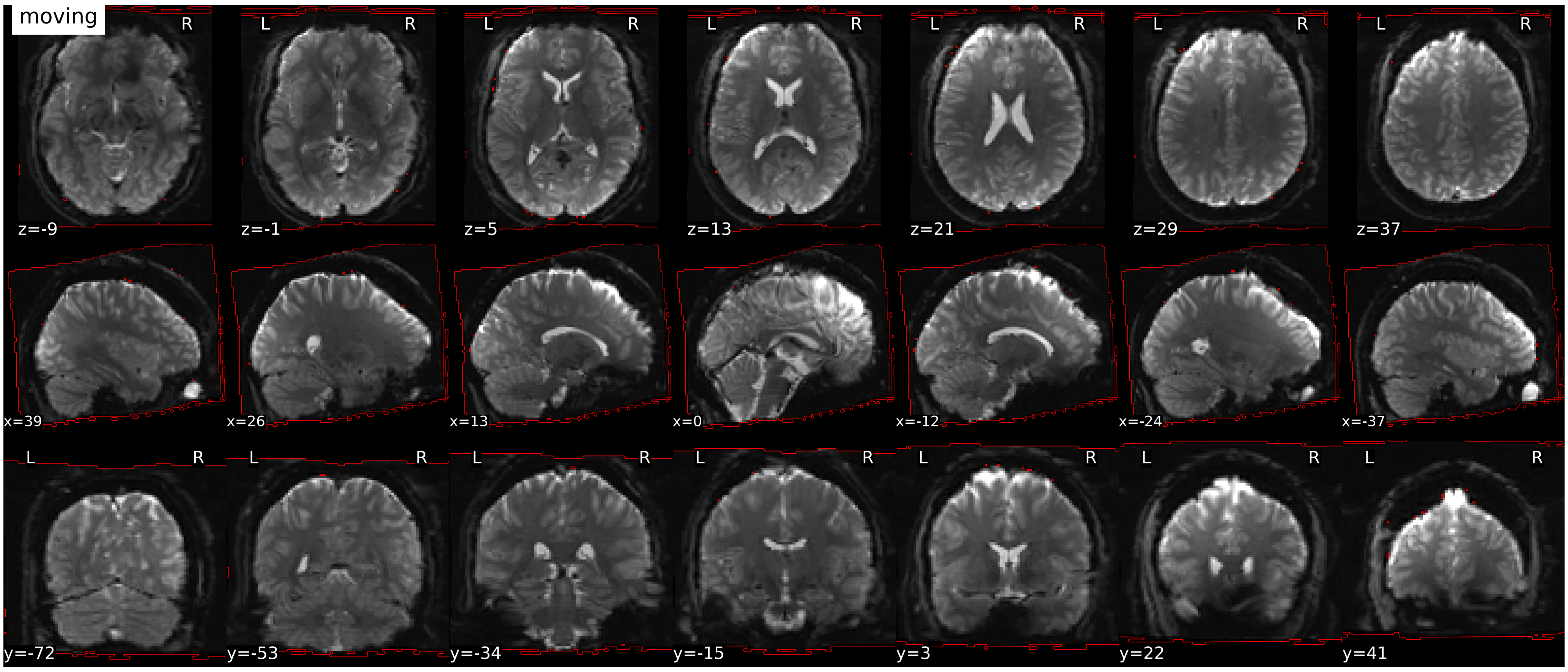


Get figure file: sub-113922/figures/sub-113922_task-EMOTION_dir-LR_space-MNI152NLin6Asym_desc-bbregister_bold.svg

##### Carpet Plot Before Postprocessing

FD and DVARS are two measures of in-scanner motion. This plot shows standardized FD, DVARS, and then a carpet plot for the time series of each voxel/vertex's time series of activity.

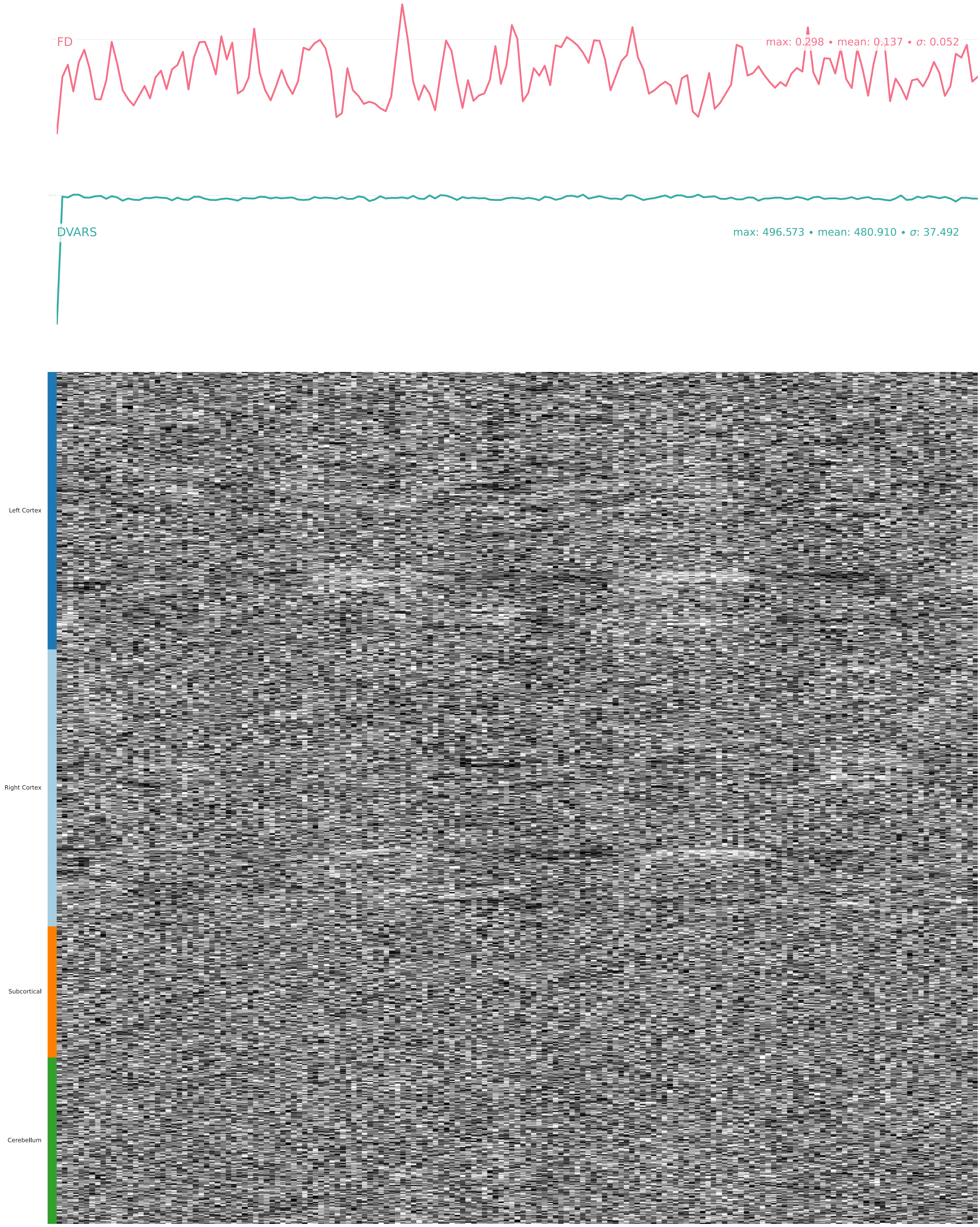


Get figure file: [sub-113922/figures/sub-113922_task-EMOTION_dir-LR_space-fsLR_desc-preprocessing_bold.svg](sub-113922/figures/sub-113922_task-EMOTION_dir-LR_space-fsLR_desc-preprocessing_bold.svg)

#### Framewise Displacement and Censored Volumes

Framewise displacement (FD) is used to flag high-motion volumes, which are then censored as part of the denoising procedure. If motion filtering is requested, then the six translation and rotation motion parameters are filtered to remove respiratory effects before FD is calculated and outlier volumes are identified.

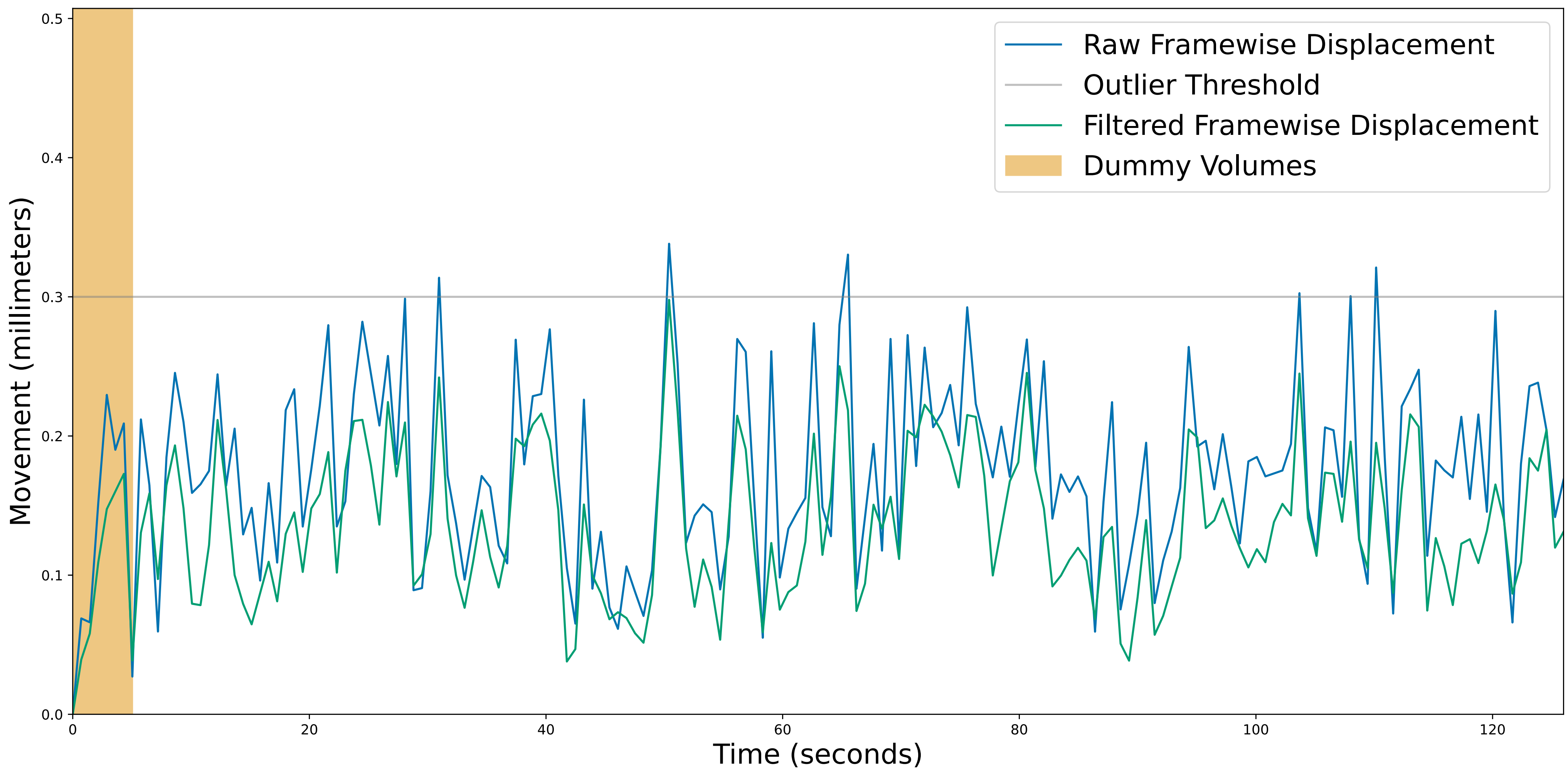


Get figure file: sub-113922/figures/sub-113922_task-EMOTION_dir-LR_space-fsLR_desc-censoring_motion.svg

#### Design Matrix for Confound Regression

The "design matrix" represents the confounds that are used to denoise the BOLD data.

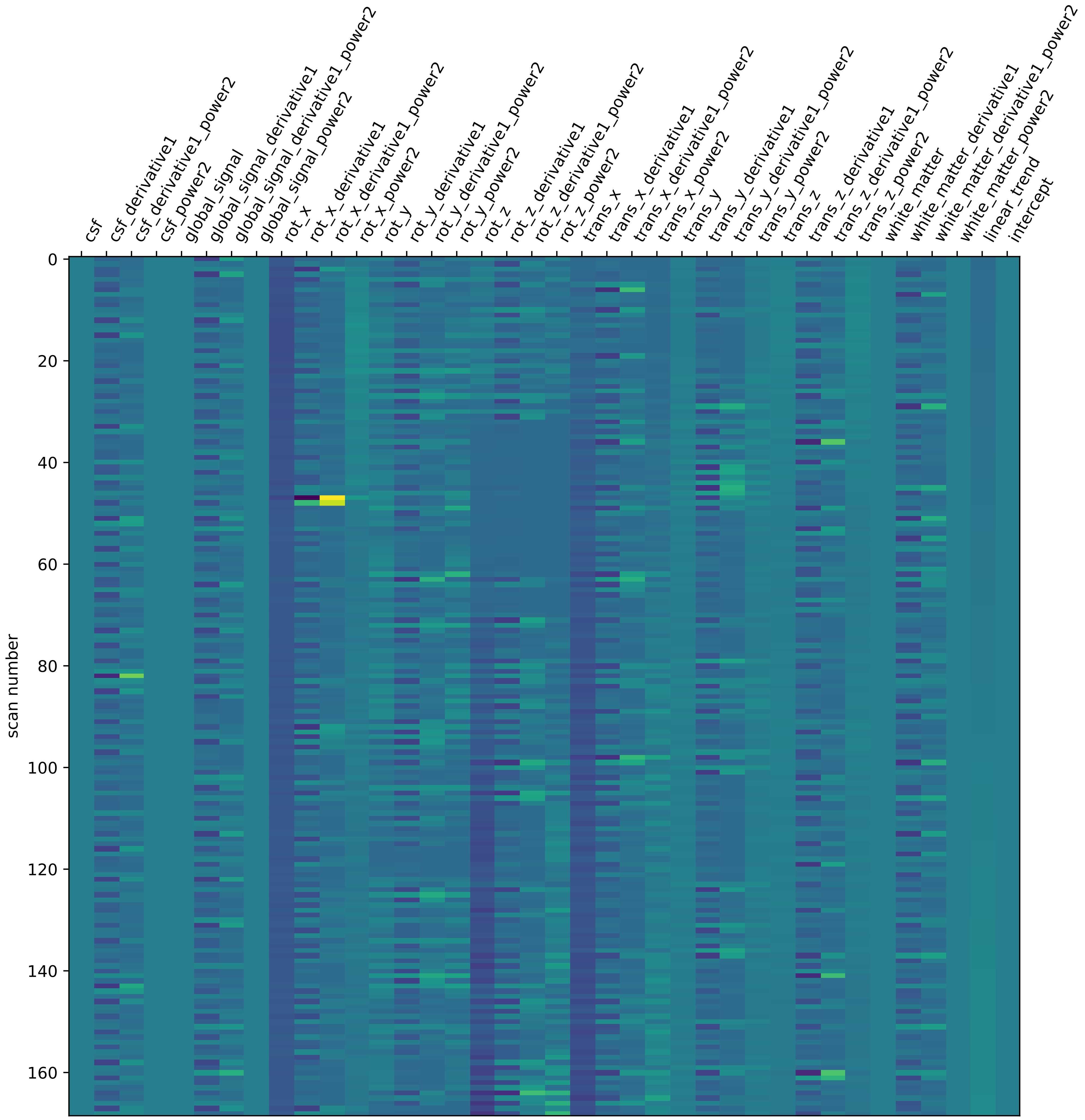


Get figure file: [sub-113922/figures/sub-113922_task-EMOTION_dir-LR_design.svg](sub-113922/figures/sub-113922_task-EMOTION_dir-LR_design.svg)

#### Carpet Plot After Postprocessing

FD and DVARS are two measures of in-scanner motion. This plot shows standardized FD, DVARS, and then a carpet plot for the time series of each voxel/vertex's time series of activity.

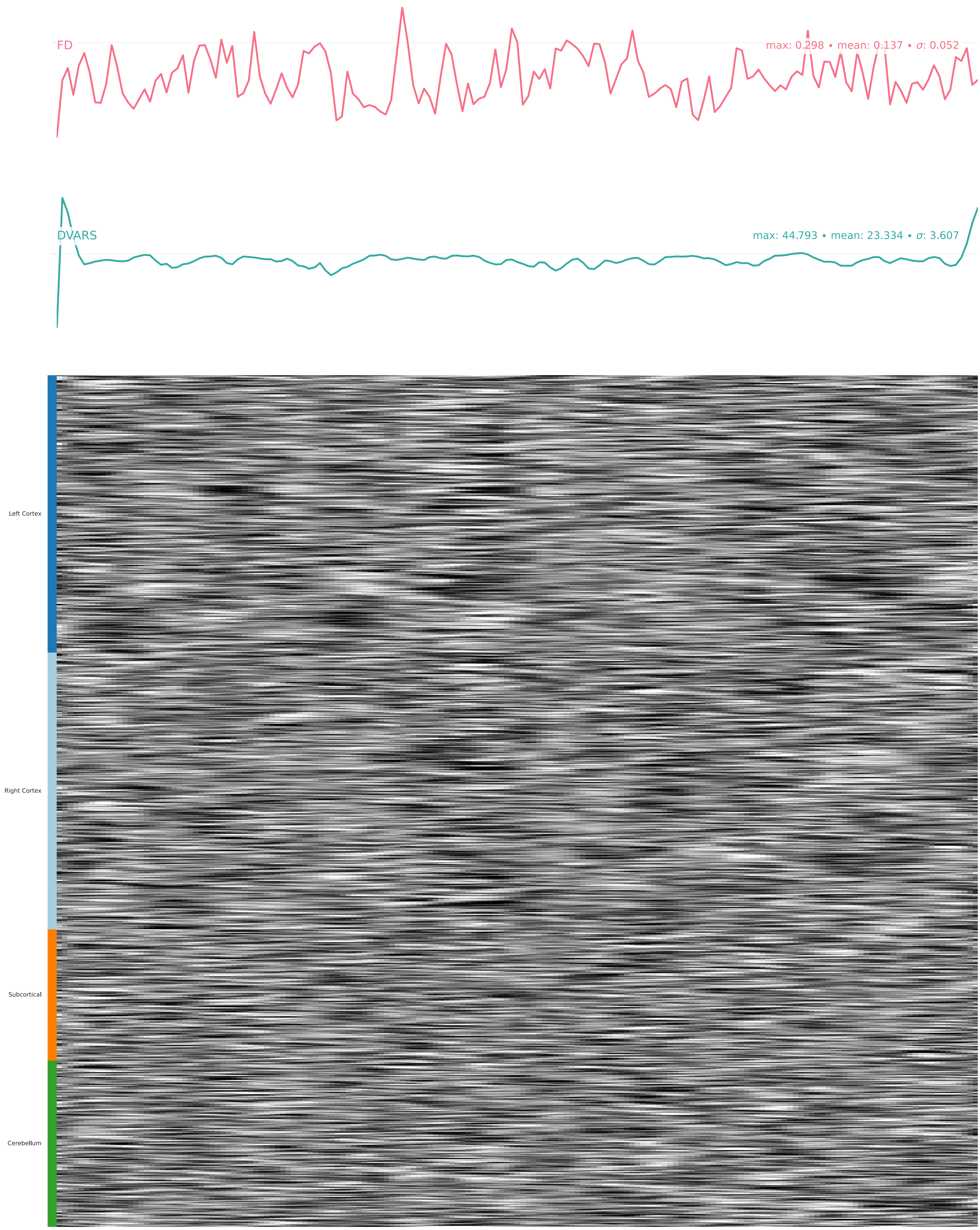


Get figure file: sub-113922/figures/sub-113922_task-EMOTION_dir-LR_space-fsLR_desc-postprocessing_bold.svg

#### Correlation Heatmaps from Four Atlases

This plot shows heatmaps from ROI-to-ROI correlations from four atlases.

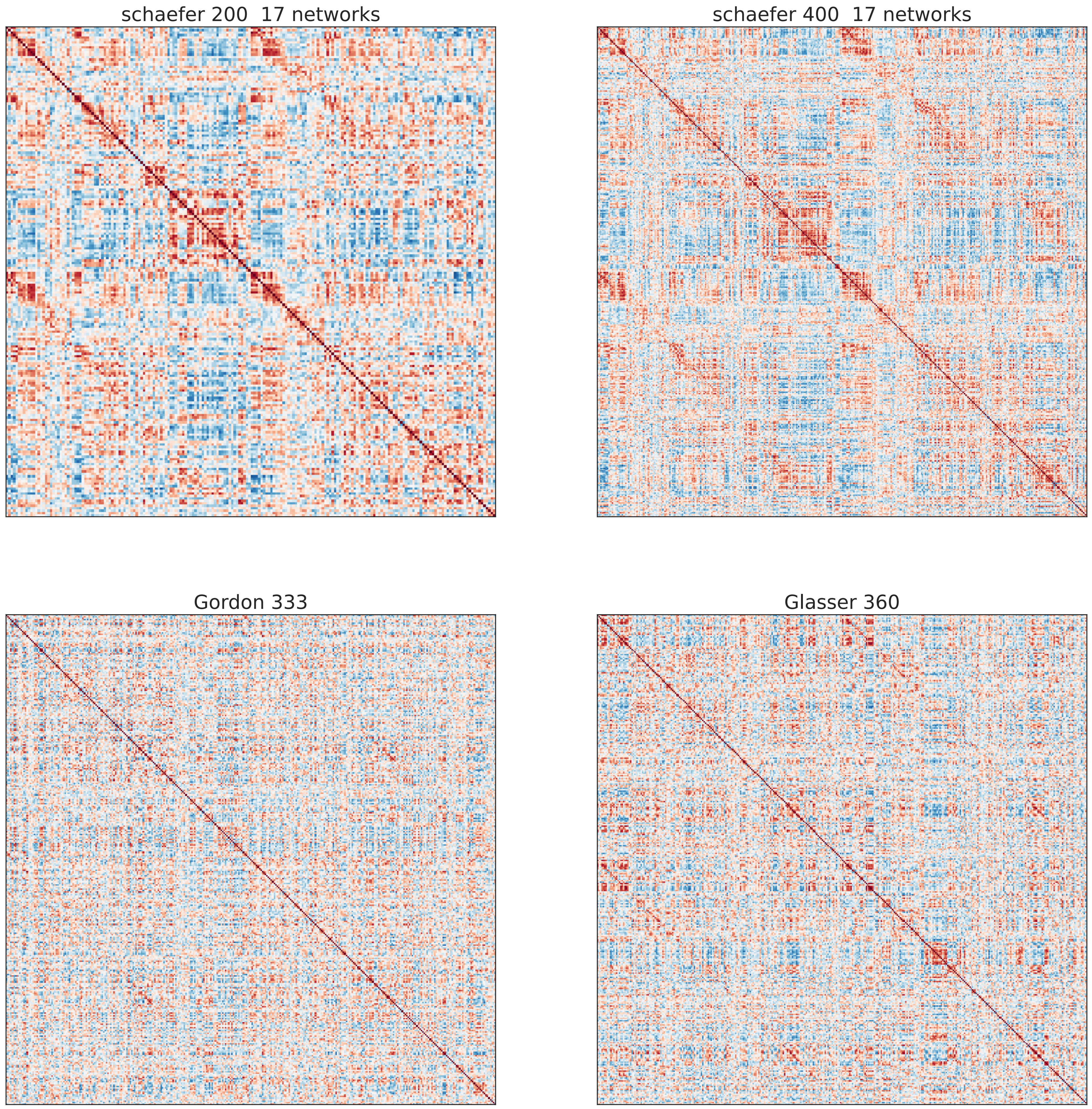


Get figure file: sub-113922/figures/sub-113922_task-EMOTION_dir-LR_space-fsLR_desc-connectivityplot_bold.svg

### Reports for: task EMOTION, direction RL.

#### Summary

- BOLD volume space: fsLR
- Repetition Time (TR): 0.72
- Mean Framewise Displacement: 0.0893
- Mean Relative RMS Motion: 0.2495
- Max Relative RMS Motion: 0.3074
- DVARS Before and After Processing : 478.2646, 23.1072
- Correlation between DVARS and FD Before and After Processing : 0.1763, 0.1443
- Number of Volumes Censored : 0

#### Alignment of functional and anatomical MRI data (surface driven)

bbregister was used to coregister functional and anatomical MRI data.

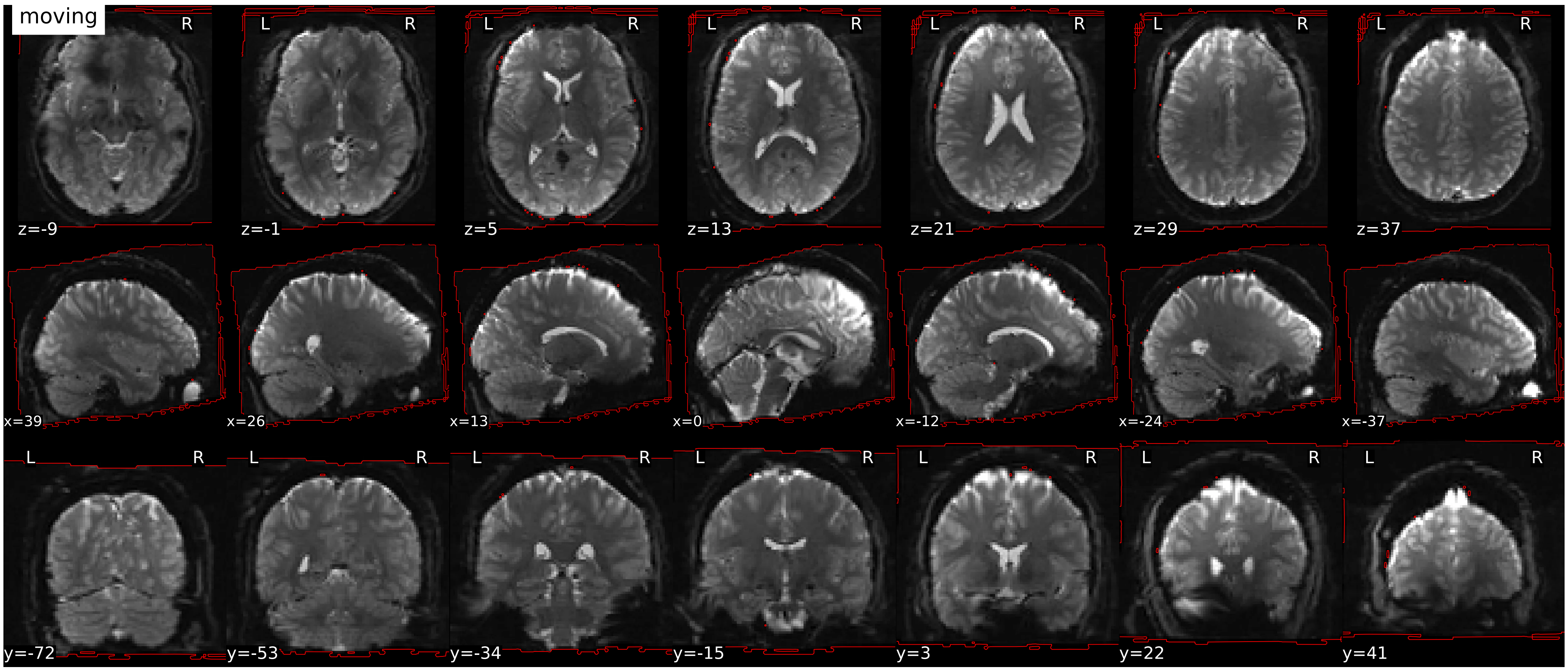


Get figure file: [sub-113922/figures/sub-113922_task-EMOTION_dir-RL_space-MNI152NLin6Asym_desc-bbregister_bold.svg](sub-113922/figures/sub-113922_task-EMOTION_dir-RL_space-MNI152NLin6Asym_desc-bbregister_bold.svg)

#### Carpet Plot Before Postprocessing

FD and DVARS are two measures of in-scanner motion. This plot shows standardized FD, DVARS, and then a carpet plot for the time series of each voxel/vertex's time series of activity.

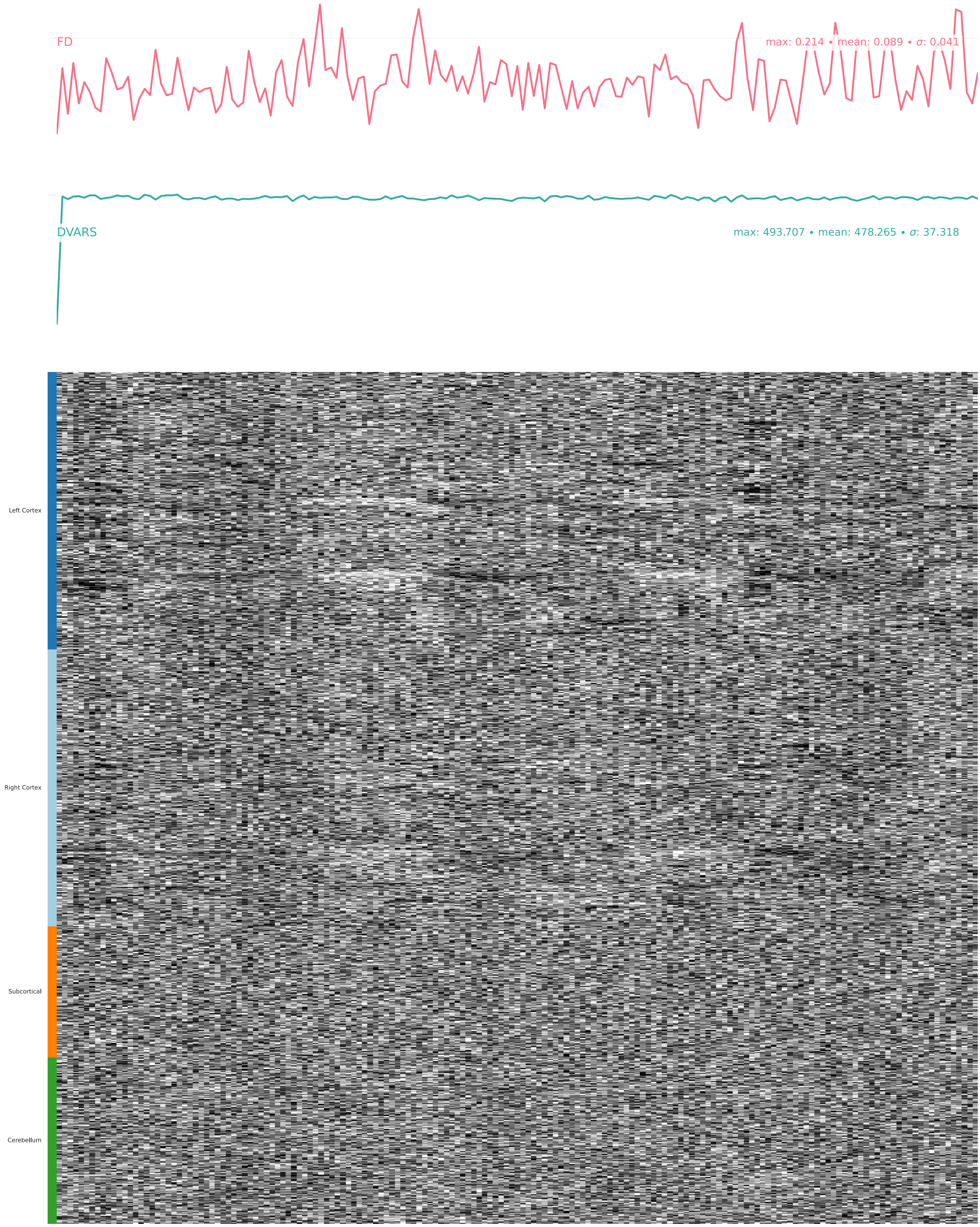


Get figure file: sub-113922/figures/sub-113922_task-EMOTION_dir-RL_space-fsLR_desc-preprocessing_bold.svg

#### Framewise Displacement and Censored Volumes

Framewise displacement (FD) is used to flag high-motion volumes, which are then censored as part of the denoising procedure. If motion filtering is requested, then the six translation and rotation motion parameters are filtered to remove respiratory effects before FD is calculated and outlier volumes are identified.

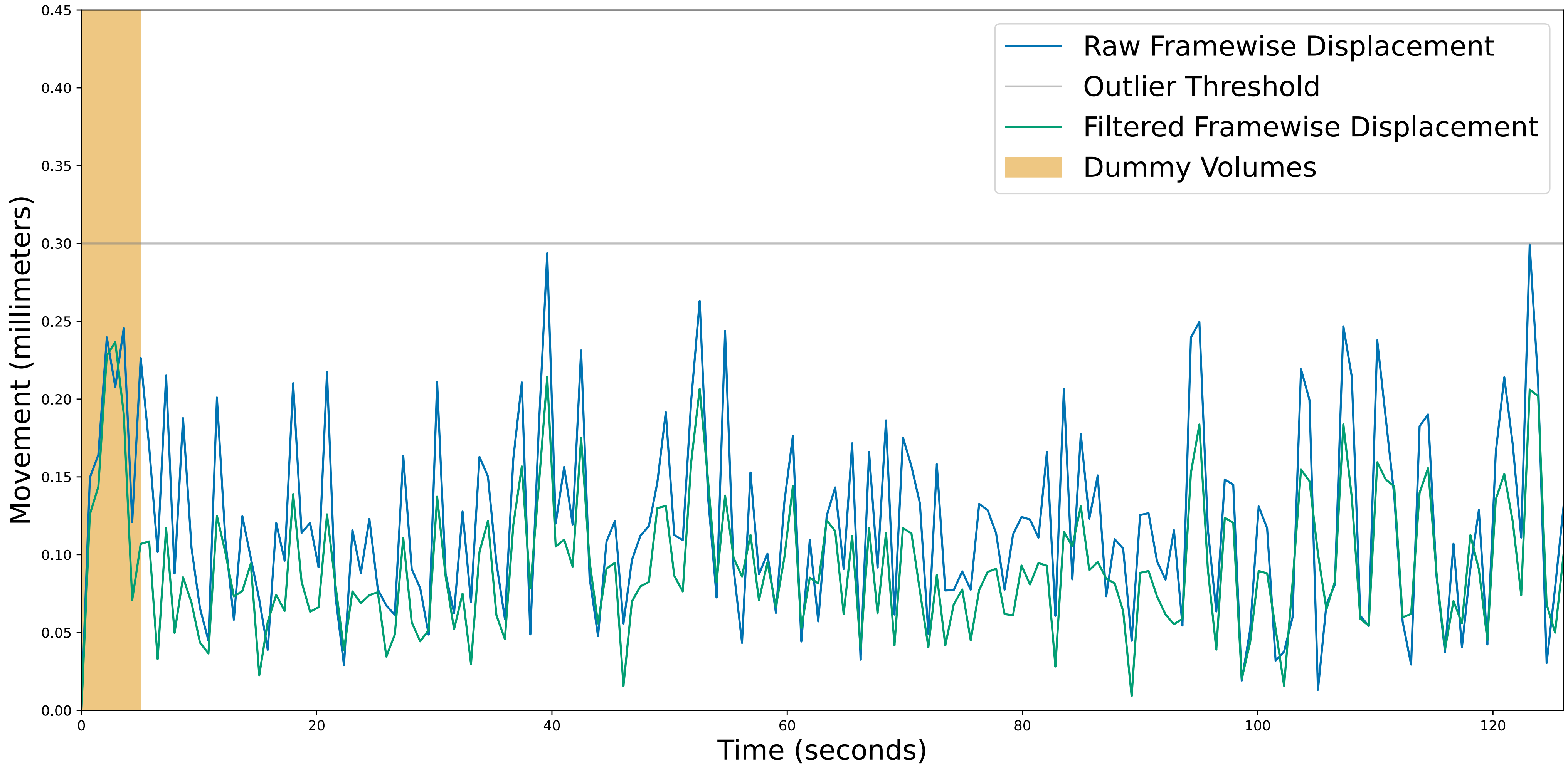


Get figure file: sub-113922/figures/sub-113922_task-EMOTION_dir-RL_space-fsLR_desc-censoring_motion.svg

#### Design Matrix for Confound Regression

The "design matrix" represents the confounds that are used to denoise the BOLD data.

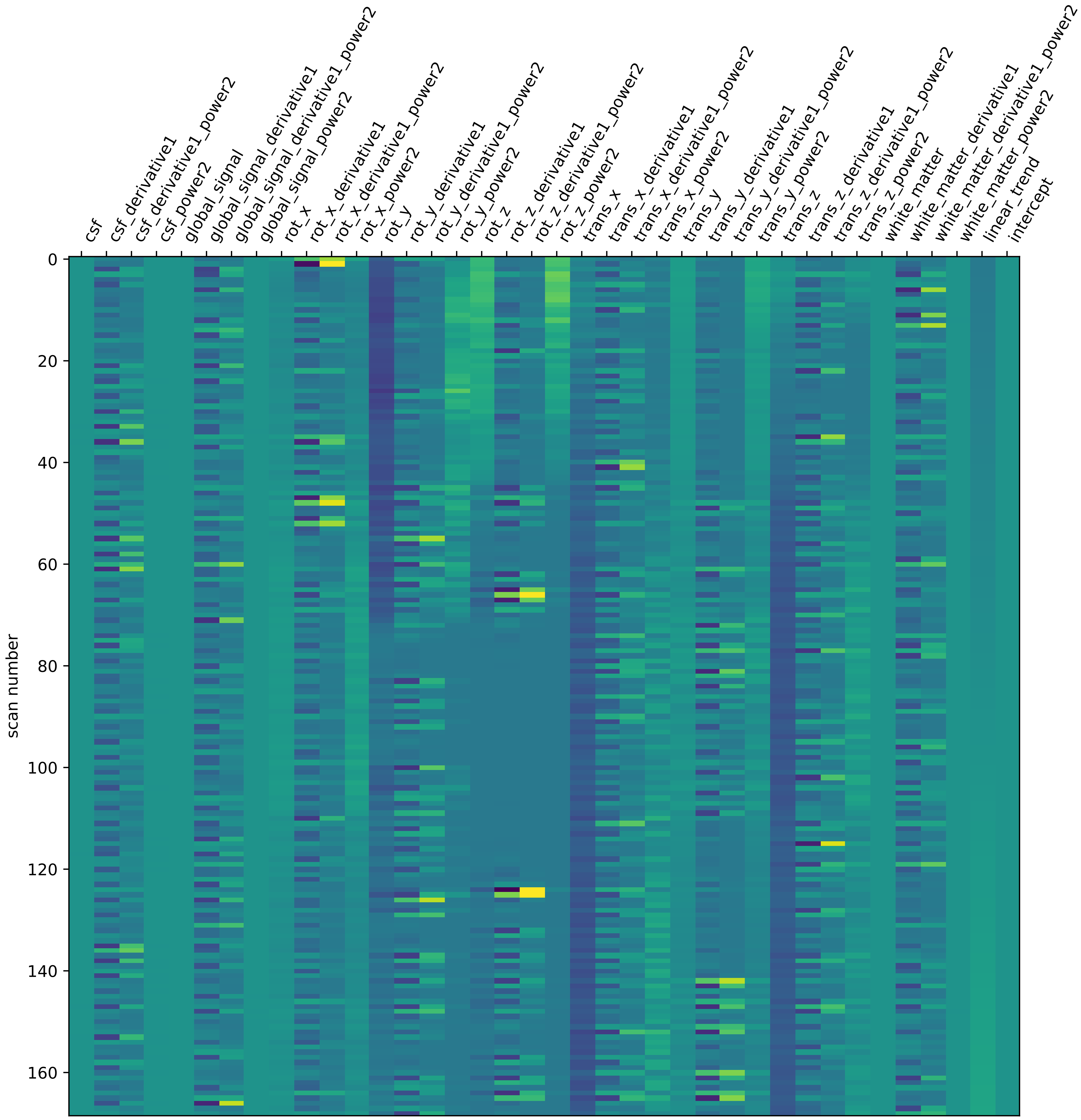


Get figure file: sub-113922/figures/sub-113922_task-EMOTION_dir-RL_design.svg

#### Carpet Plot After Postprocessing

FD and DVARS are two measures of in-scanner motion. This plot shows standardized FD, DVARS, and then a carpet plot for the time series of each voxel/vertex's time series of activity.

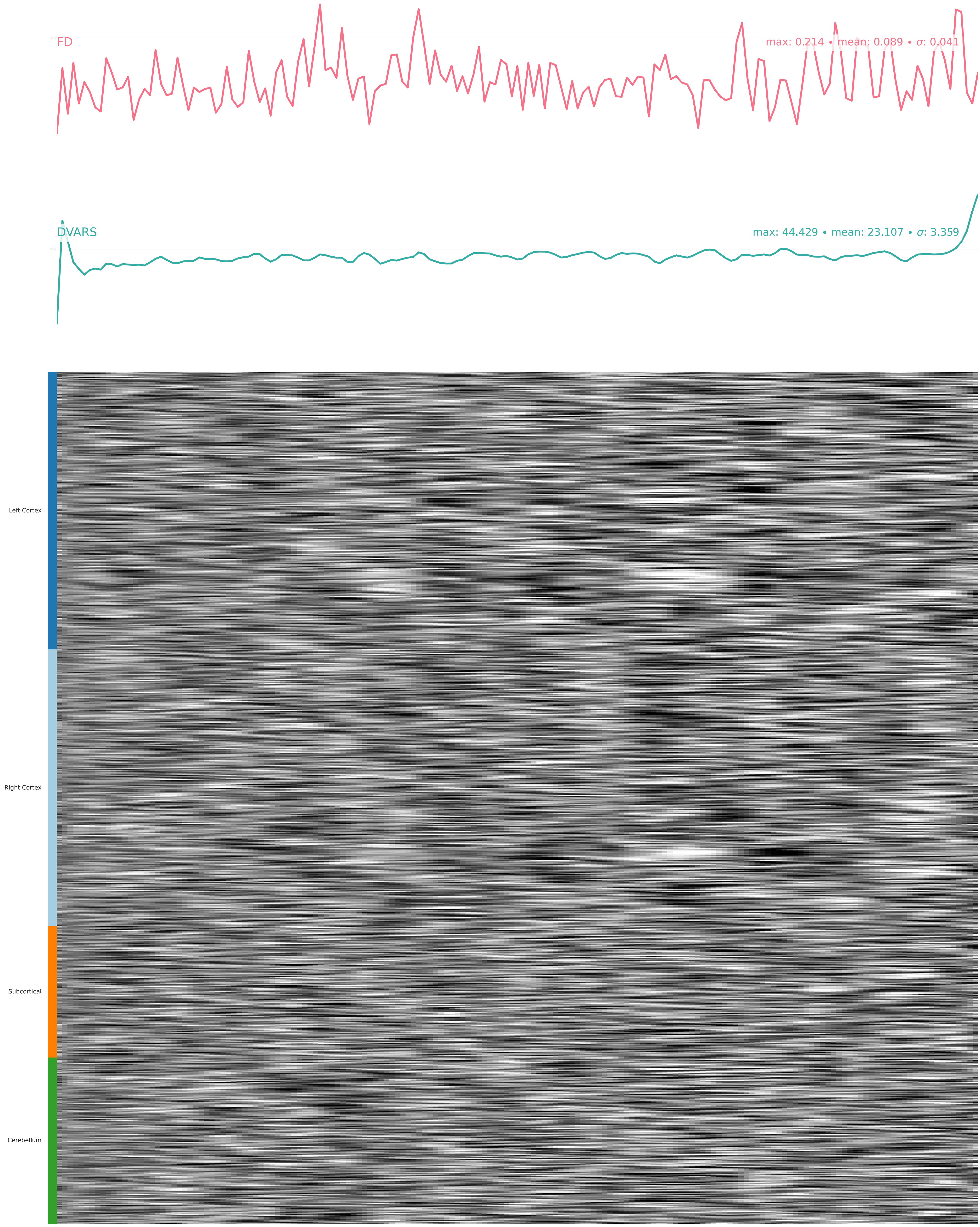


Get figure file: [sub-113922/figures/sub-113922_task-EMOTION_dir-RL_space-fsLR_desc-postprocessing_bold.svg](sub-113922/figures/sub-113922_task-EMOTION_dir-RL_space-fsLR_desc-postprocessing_bold.svg)

#### Correlation Heatmaps from Four Atlases

This plot shows heatmaps from ROI-to-ROI correlations from four atlases.

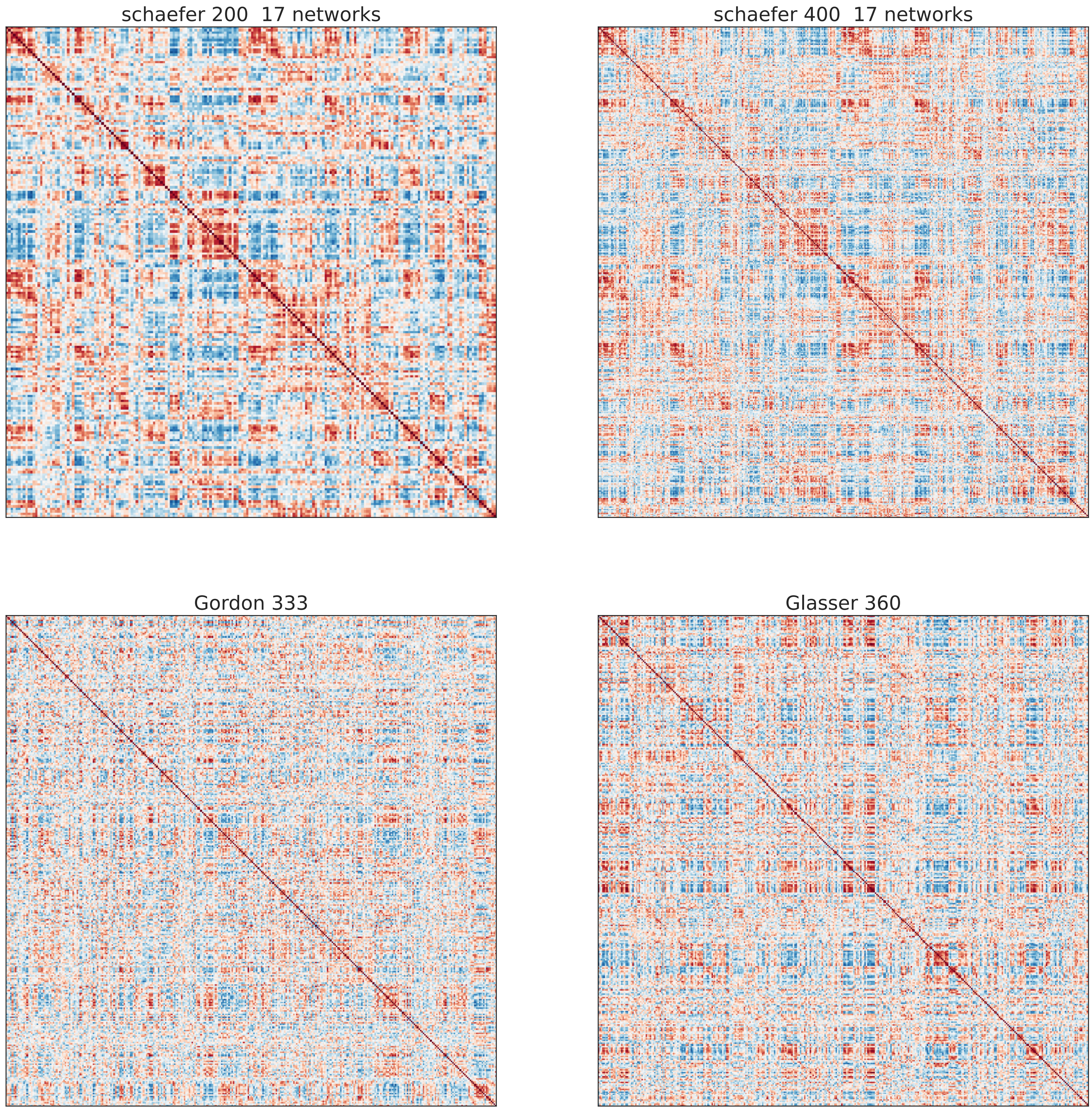


Get figure file: sub-113922/figures/sub-113922_task-EMOTION_dir-RL_space-fsLR_desc-connectivityplot_bold.svg

### Reports for: task GAMBLING, direction LR.

#### Summary

- BOLD volume space: fsLR
- Repetition Time (TR): 0.72
- Mean Framewise Displacement: 0.1329
- Mean Relative RMS Motion: 0.2238
- Max Relative RMS Motion: 0.4807
- DVARS Before and After Processing : 469.4174, 24.2365
- Correlation between DVARS and FD Before and After Processing : 0.166, -0.0095
- Number of Volumes Censored : 0

#### Alignment of functional and anatomical MRI data (surface driven)

bbregister was used to coregister functional and anatomical MRI data.

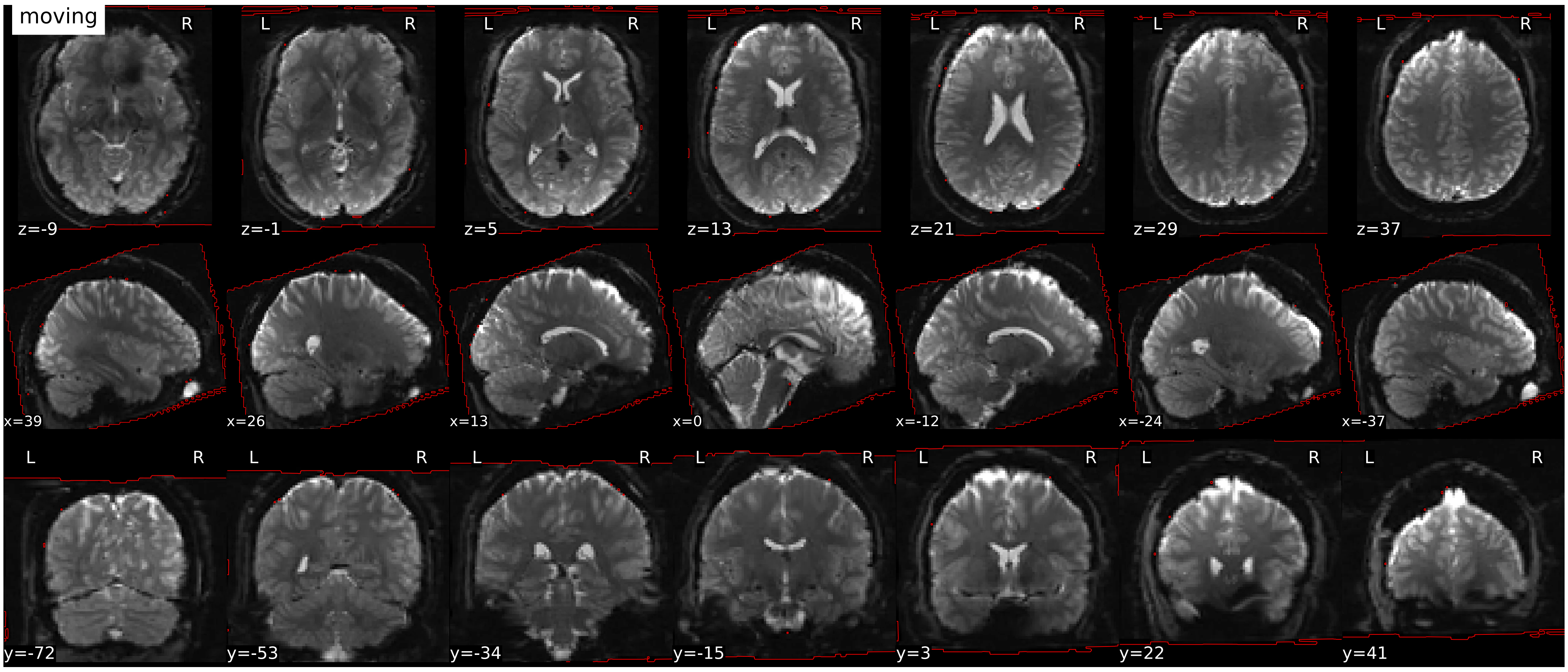


Get figure file: sub-113922/figures/sub-113922_task-GAMBLING_dir-LR_space-MNI152NLin6Asym_desc-bbregister_bold.svg

### Carpet Plot Before Postprocessing

FD and DVARS are two measures of in-scanner motion. This plot shows standardized FD, DVARS, and then a carpet plot for the time series of each voxel/vertex's time series of activity.

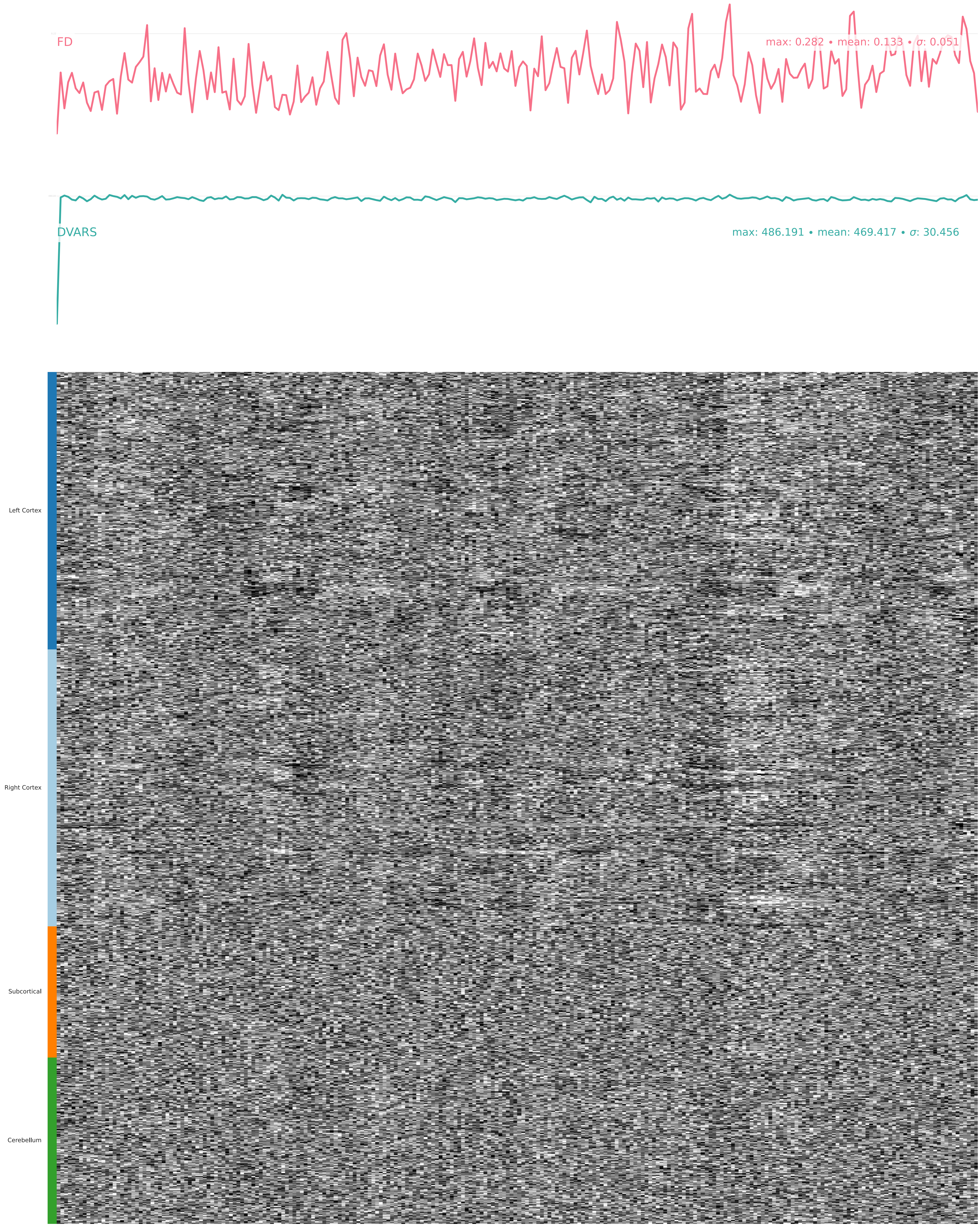


Get figure file: sub-113922/figures/sub-113922_task-GAMBLING_dir-LR_space-fsLR_desc-preprocessing_bold.svg

#### Framewise Displacement and Censored Volumes

Framewise displacement (FD) is used to flag high-motion volumes, which are then censored as part of the denoising procedure. If motion filtering is requested, then the six translation and rotation motion parameters are filtered to remove respiratory effects before FD is calculated and outlier volumes are identified.

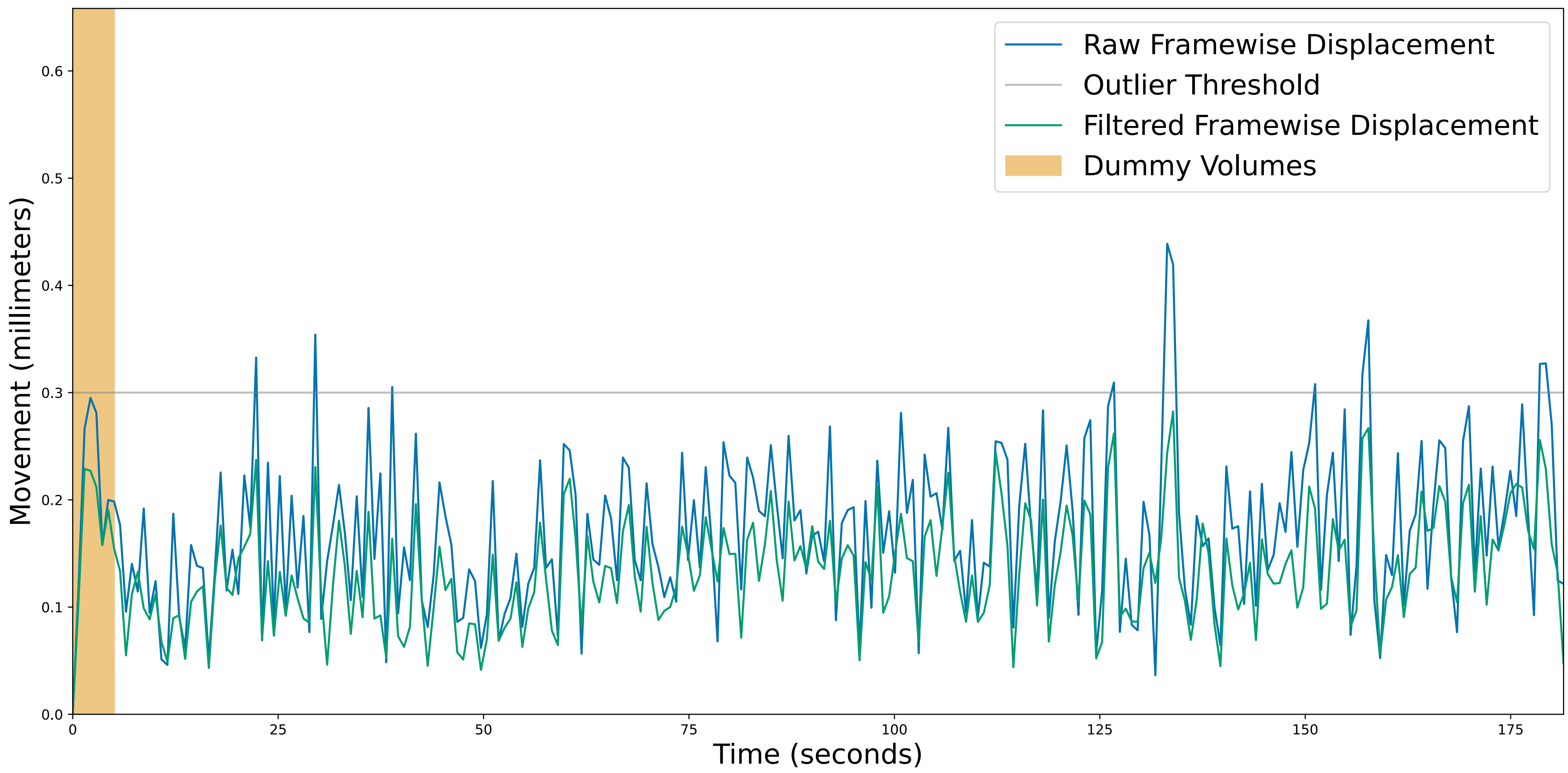


Get figure file: [sub-113922/figures/sub-113922_task-GAMBLING_dir-LR_space-fsLR_desc-censoring_motion.svg](sub-113922/figures/sub-113922_task-GAMBLING_dir-LR_space-fsLR_desc-censoring_motion.svg)

#### Design Matrix for Confound Regression

The "design matrix" represents the confounds that are used to denoise the BOLD data.

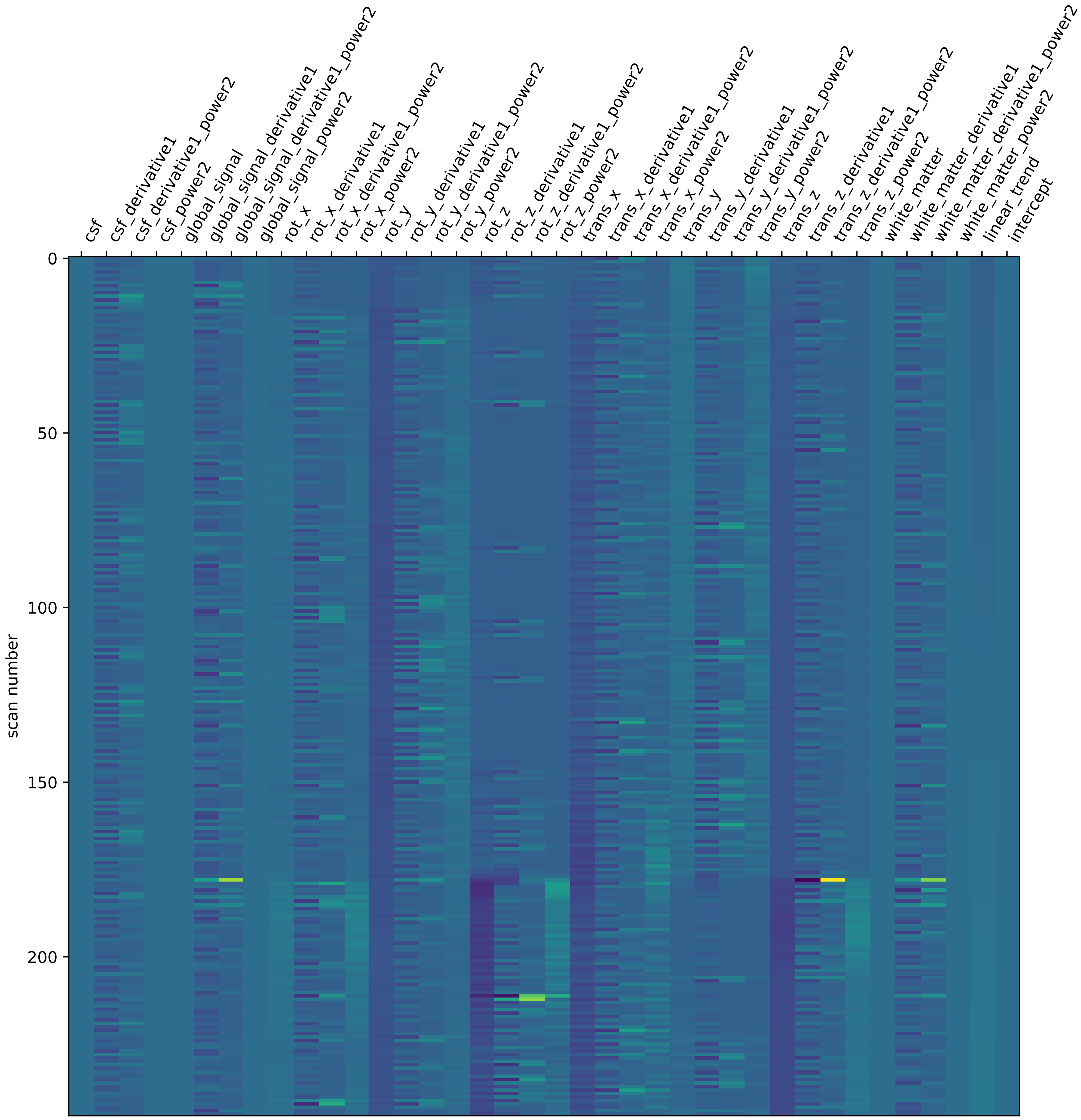


Get figure file: [sub-113922/figures/sub-113922_task-GAMBLING_dir-LR_design.svg](sub-113922/figures/sub-113922_task-GAMBLING_dir-LR_design.svg)

#### Carpet Plot After Postprocessing

FD and DVARS are two measures of in-scanner motion. This plot shows standardized FD, DVARS, and then a carpet plot for the time series of each voxel/vertex's time series of activity.

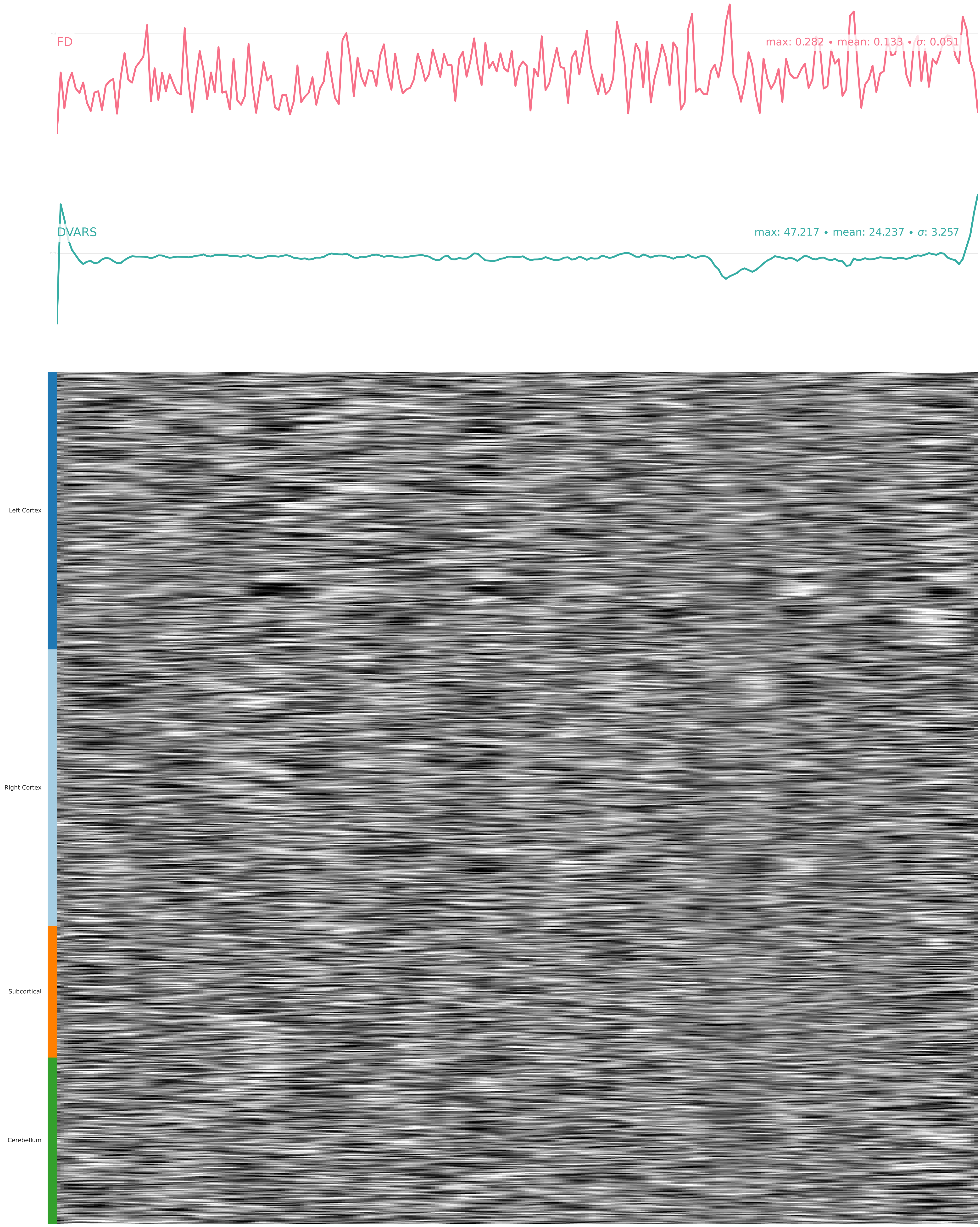


Get figure file: sub-113922/figures/sub-113922_task-GAMBLING_dir-LR_space-fsLR_desc-postprocessing_bold.svg

#### Correlation Heatmaps from Four Atlases

This plot shows heatmaps from ROI-to-ROI correlations from four atlases.

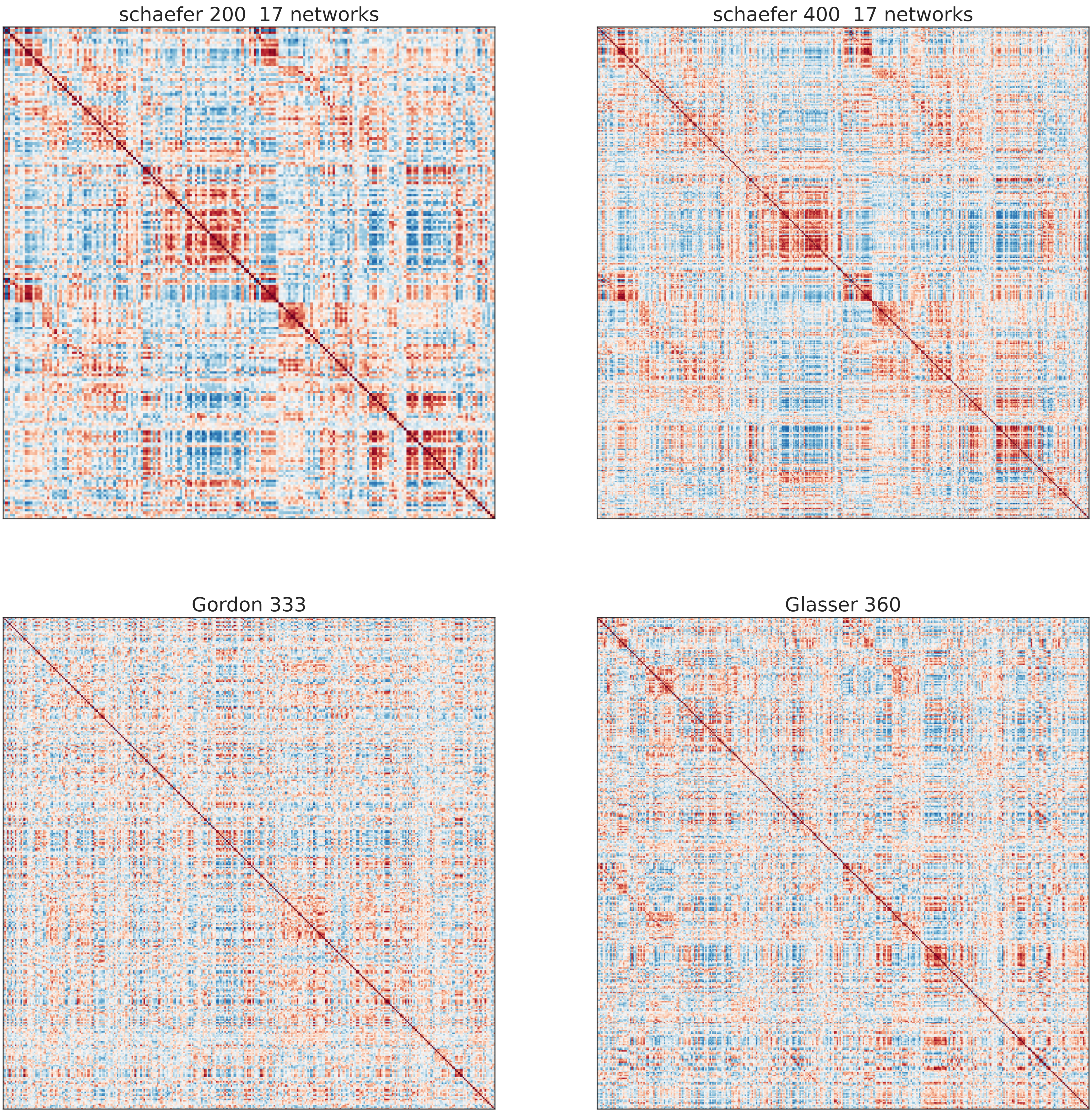


Get figure file: [sub-113922/figures/sub-113922_task-GAMBLING_dir-LR_space-fsLR_desc-connectivityplot_bold.svg]

### Reports for: task GAMBLING, direction RL.

#### Summary

- BOLD volume space: fsLR
- Repetition Time (TR): 0.72
- Mean Framewise Displacement: 0.128
- Mean Relative RMS Motion: 0.3525
- Max Relative RMS Motion: 0.5246
- DVARS Before and After Processing : 473.3058, 25.3794
- Correlation between DVARS and FD Before and After Processing : 0.3746, 0.2324
- Number of Volumes Censored : 6

#### Alignment of functional and anatomical MRI data (surface driven)

bbregister was used to coregister functional and anatomical MRI data.

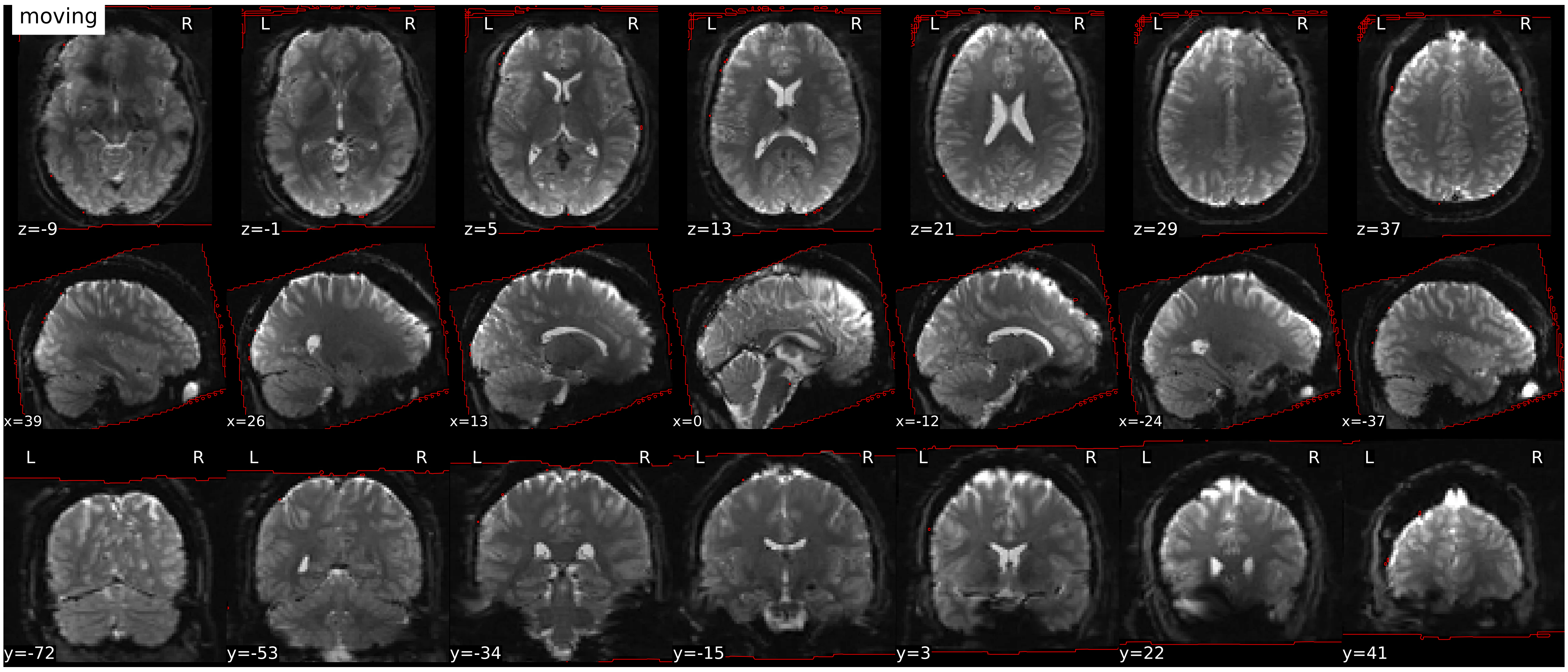


Get figure file: sub-113922/figures/sub-113922_task-GAMBLING_dir-RL_space-MNI152NLin6Asym_desc-bbregister_bold.svg

#### Carpet Plot Before Postprocessing

FD and DVARS are two measures of in-scanner motion. This plot shows standardized FD, DVARS, and then a carpet plot for the time series of each voxel/vertex's time series of activity.

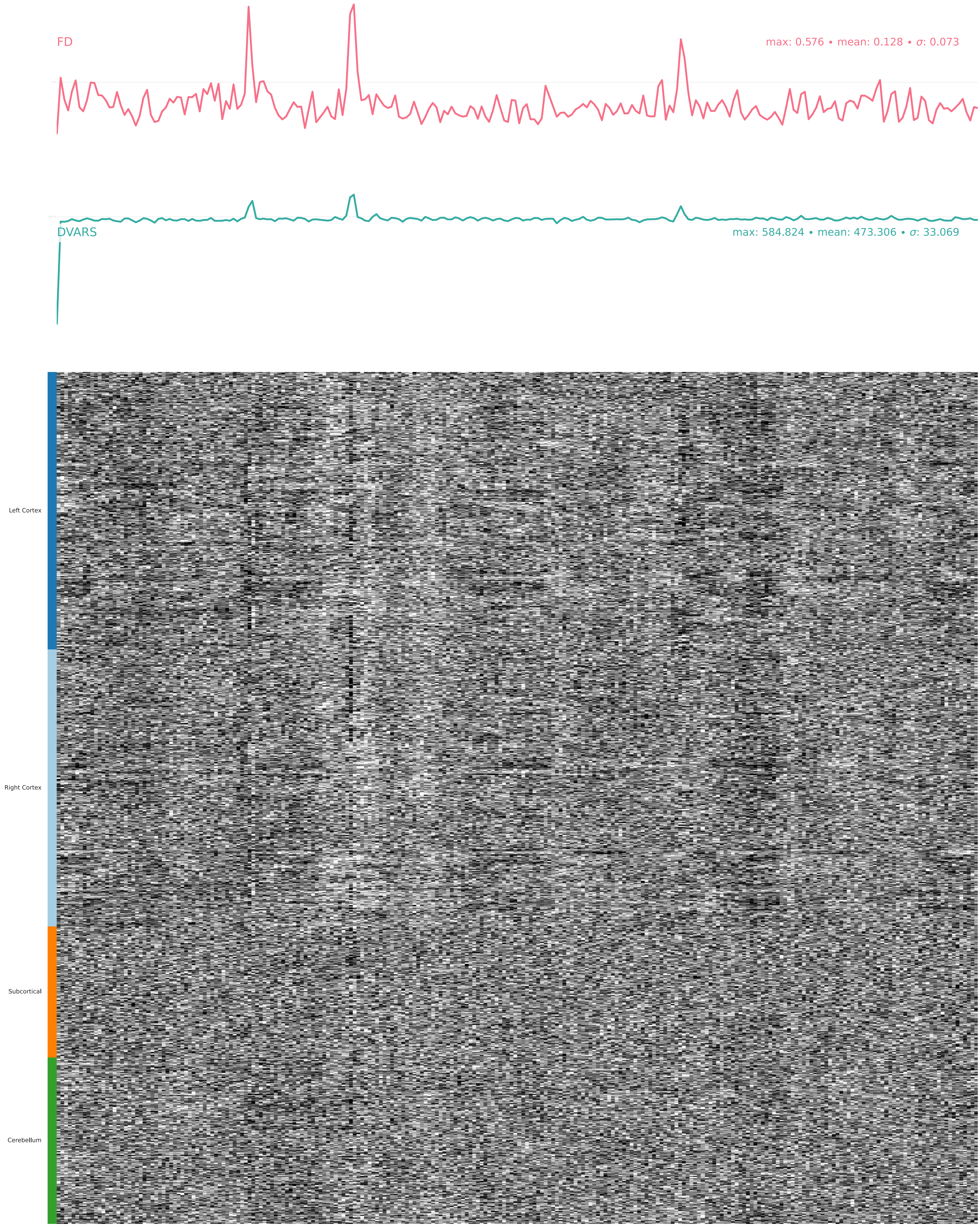


Get figure file: sub-113922/figures/sub-113922_task-GAMBLING_dir-RL_space-fsLR_desc-preprocessing_bold.svg

#### Framewise Displacement and Censored Volumes

Framewise displacement (FD) is used to flag high-motion volumes, which are then censored as part of the denoising procedure. If motion filtering is requested, then the six translation and rotation motion parameters are filtered to remove respiratory effects before FD is calculated and outlier volumes are identified.

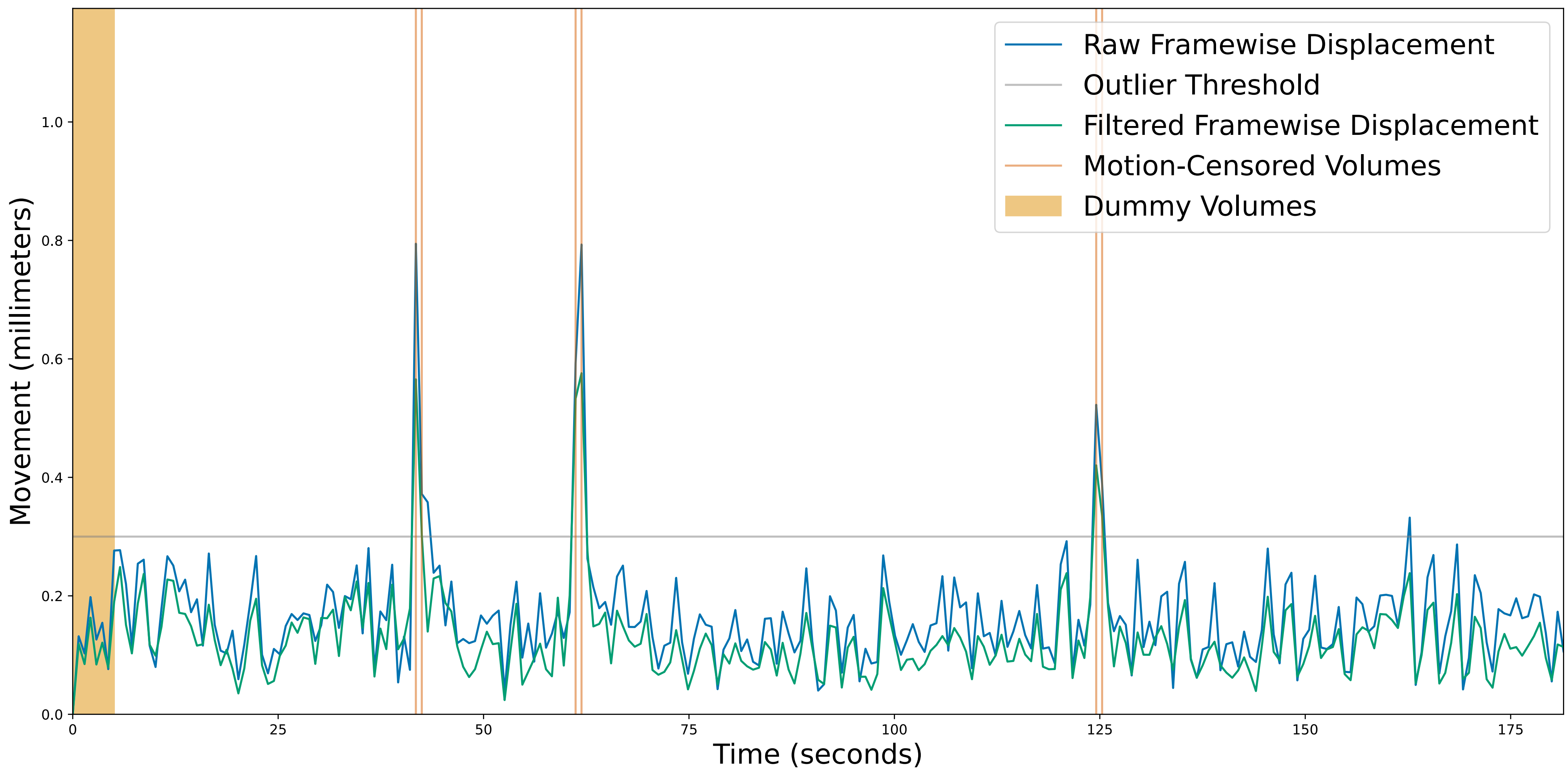


Get figure file: sub-113922/figures/sub-113922_task-GAMBLING_dir-RL_space-fsLR_desc-censoring_motion.svg

#### Design Matrix for Confound Regression

The "design matrix" represents the confounds that are used to denoise the BOLD data.

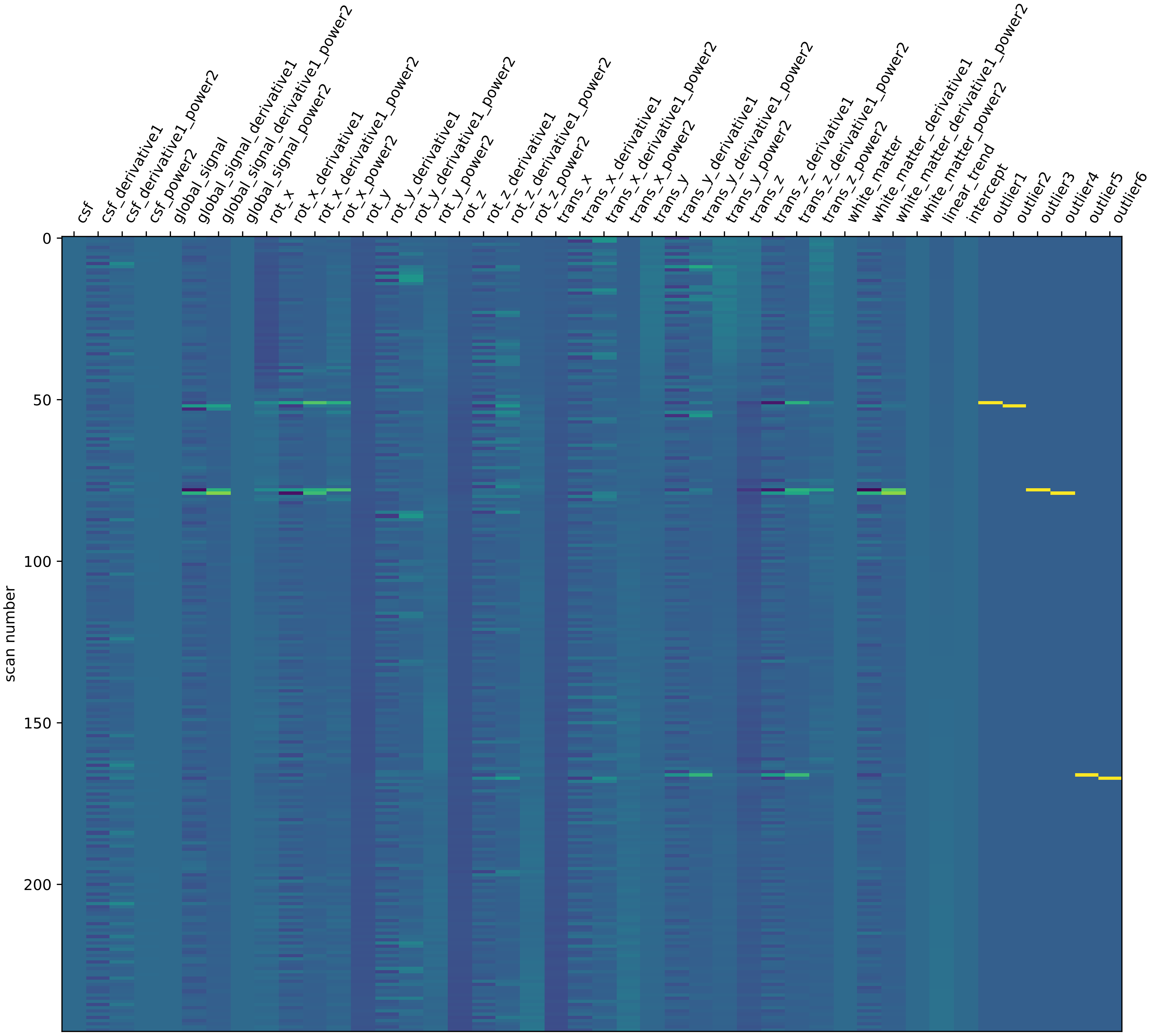


Get figure file: sub-113922/figures/sub-113922_task-GAMBLING_dir-RL_design.svg

#### Carpet Plot After Postprocessing

FD and DVARS are two measures of in-scanner motion. This plot shows standardized FD, DVARS, and then a carpet plot for the time series of each voxel/vertex's time series of activity.

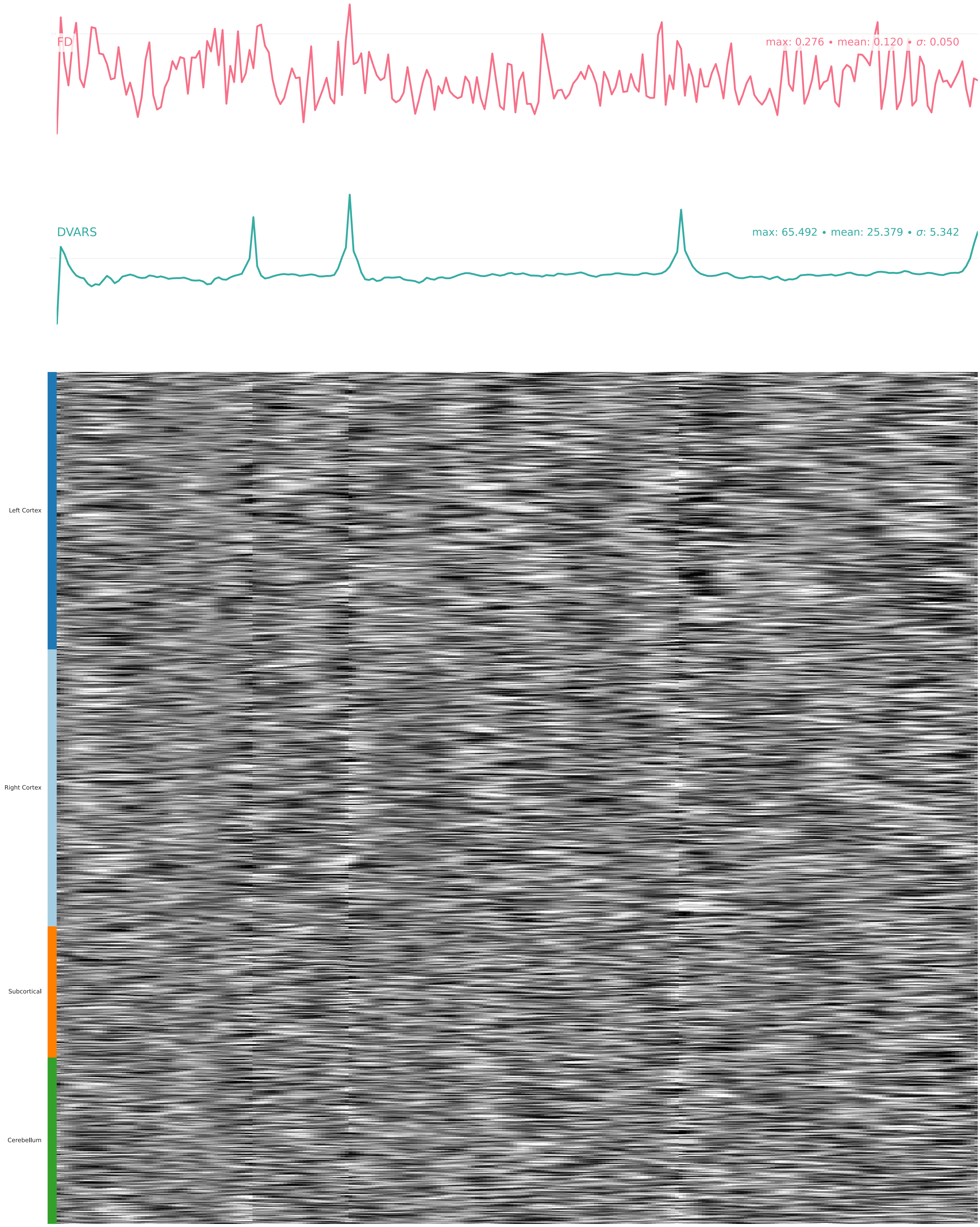


Get figure file: sub-113922/figures/sub-113922_task-GAMBLING_dir-RL_space-fsLR_desc-postprocessing_bold.svg

#### Correlation Heatmaps from Four Atlases

This plot shows heatmaps from ROI-to-ROI correlations from four atlases.

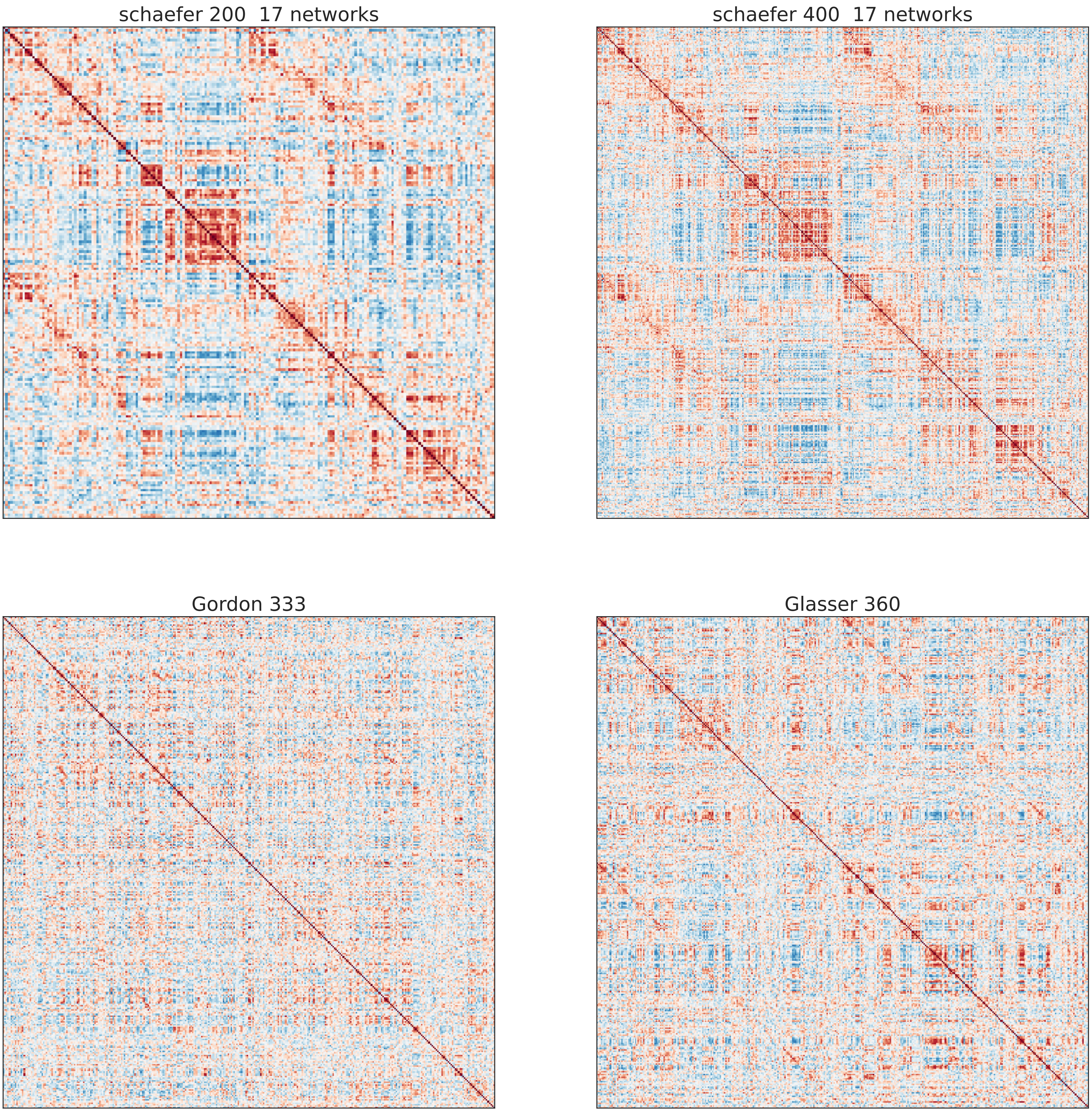


Get figure file: sub-113922/figures/sub-113922_task-GAMBLING_dir-RL_space-fsLR_desc-connectivityplot_bold.svg

### Reports for: task LANGUAGE, direction LR.

#### Summary

- BOLD volume space: fsLR
- Repetition Time (TR): 0.72
- Mean Framewise Displacement: 0.098
- Mean Relative RMS Motion: 0.1956
- Max Relative RMS Motion: 0.312
- DVARS Before and After Processing : 478.9544, 25.1346
- Correlation between DVARS and FD Before and After Processing : 0.172, -0.0018
- Number of Volumes Censored : 1

#### Alignment of functional and anatomical MRI data (surface driven)

bbregister was used to coregister functional and anatomical MRI data.

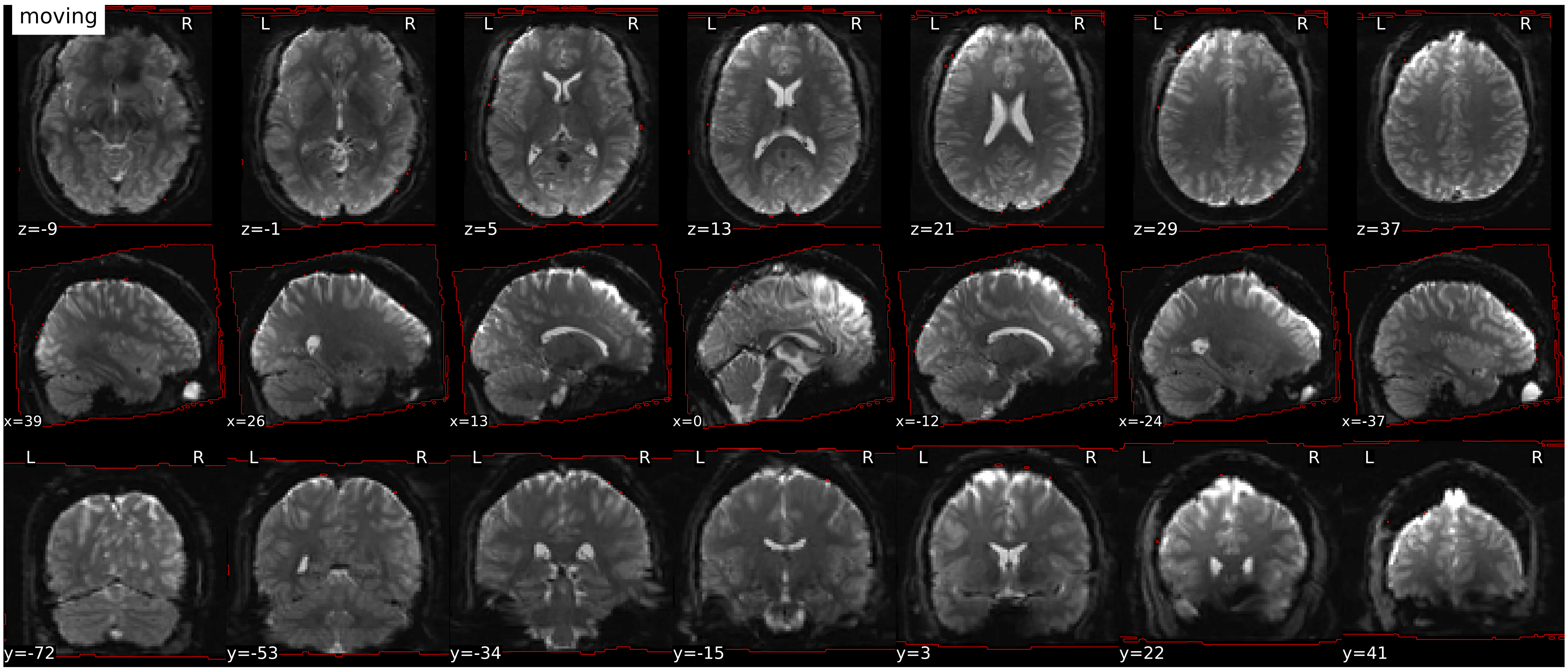


Get figure file: sub-113922/figures/sub-113922_task-LANGUAGE_dir-LR_space-MNI152NLin6Asym_desc-bbregister_bold.svg

#### Carpet Plot Before Postprocessing

FD and DVARS are two measures of in-scanner motion. This plot shows standardized FD, DVARS, and then a carpet plot for the time series of each voxel/vertex's time series of activity.

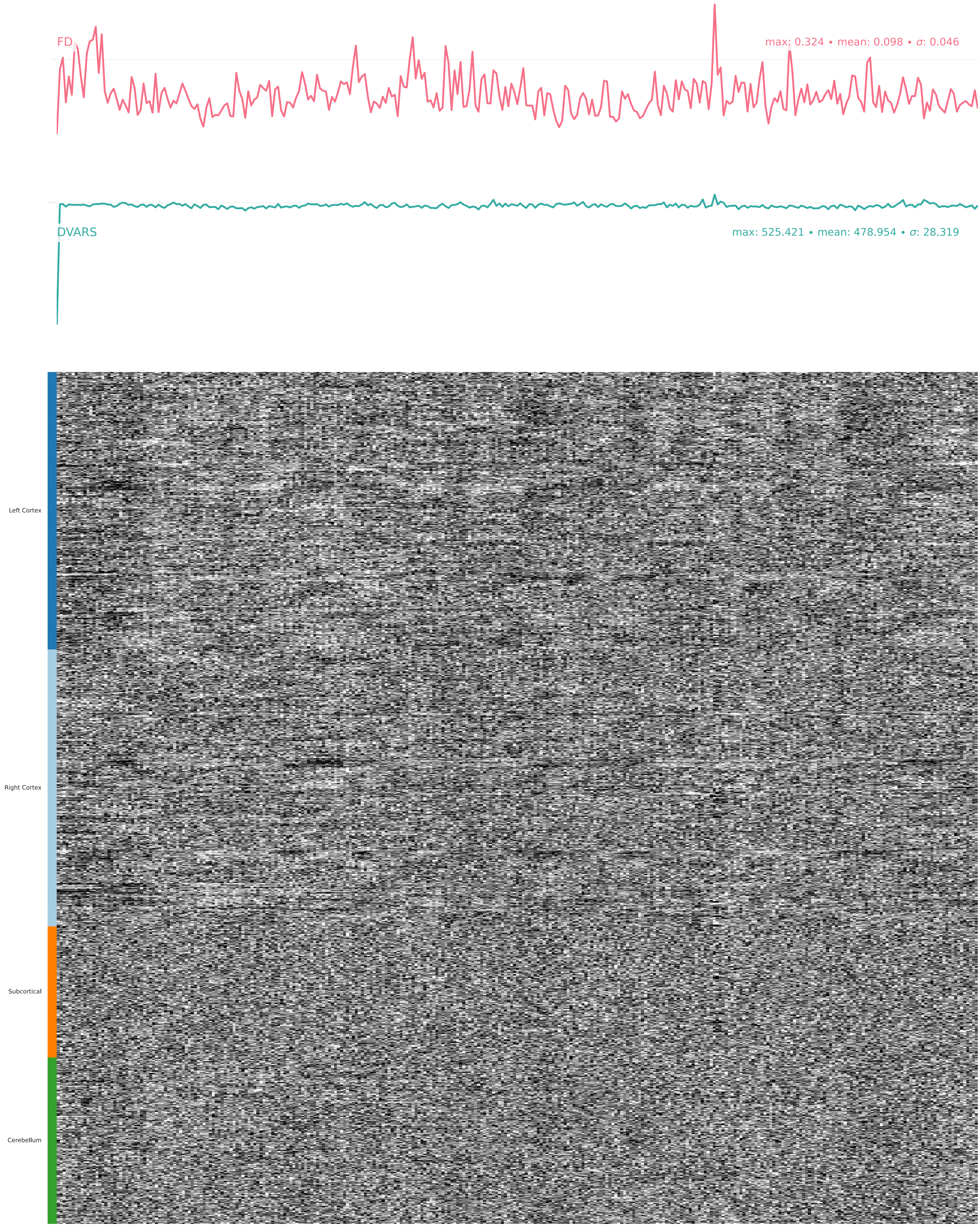


Get figure file: sub-113922/figures/sub-113922_task-LANGUAGE_dir-LR_space-fsLR_desc-preprocessing_bold.svg

#### Framewise Displacement and Censored Volumes

Framewise displacement (FD) is used to flag high-motion volumes, which are then censored as part of the denoising procedure. If motion filtering is requested, then the six translation and rotation motion parameters are filtered to remove respiratory effects before FD is calculated and outlier volumes are identified.

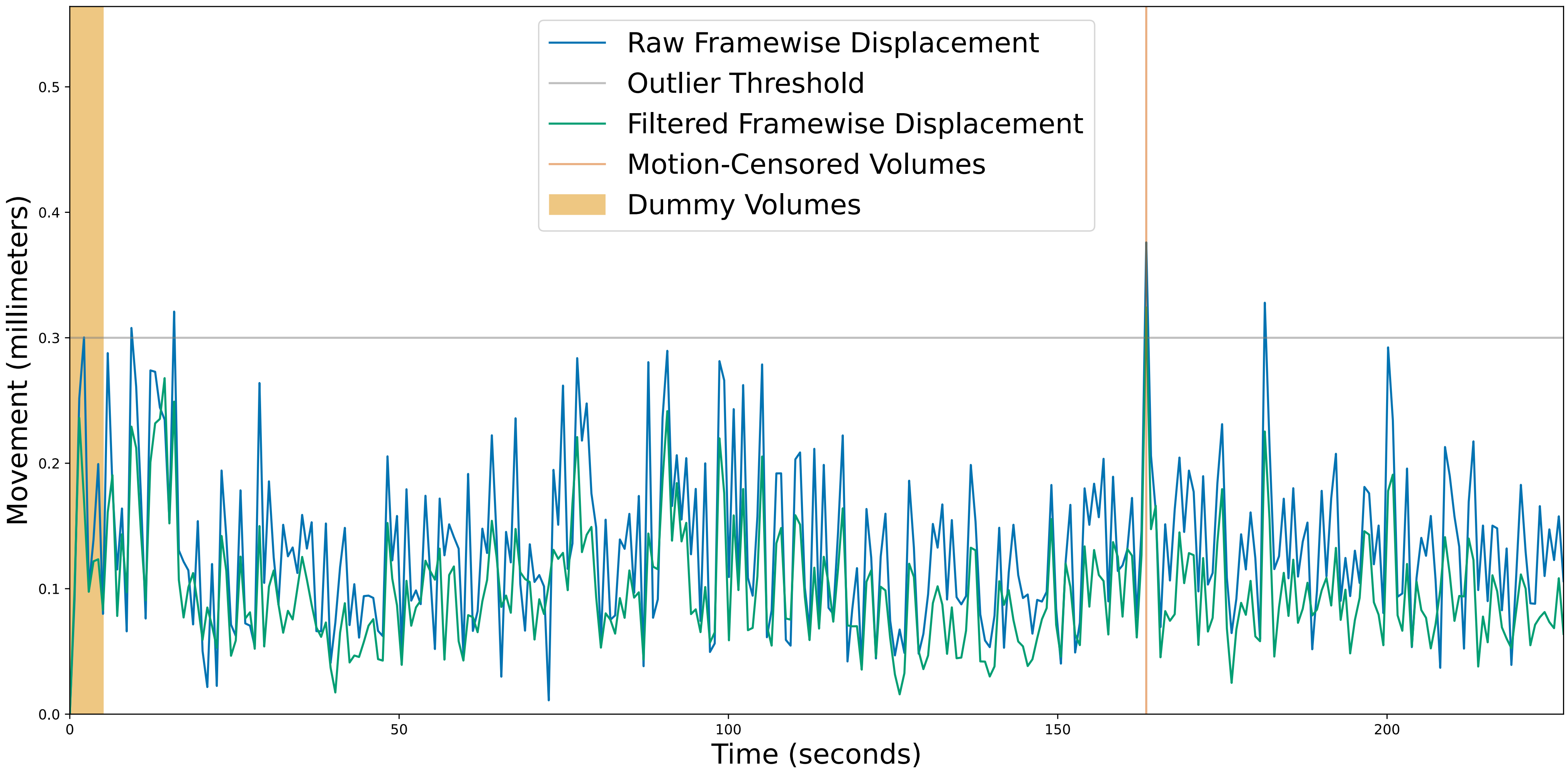


Get figure file: sub-113922/figures/sub-113922_task-LANGUAGE_dir-LR_space-fsLR_desc-censoring_motion.svg

#### Design Matrix for Confound Regression

The "design matrix" represents the confounds that are used to denoise the BOLD data.

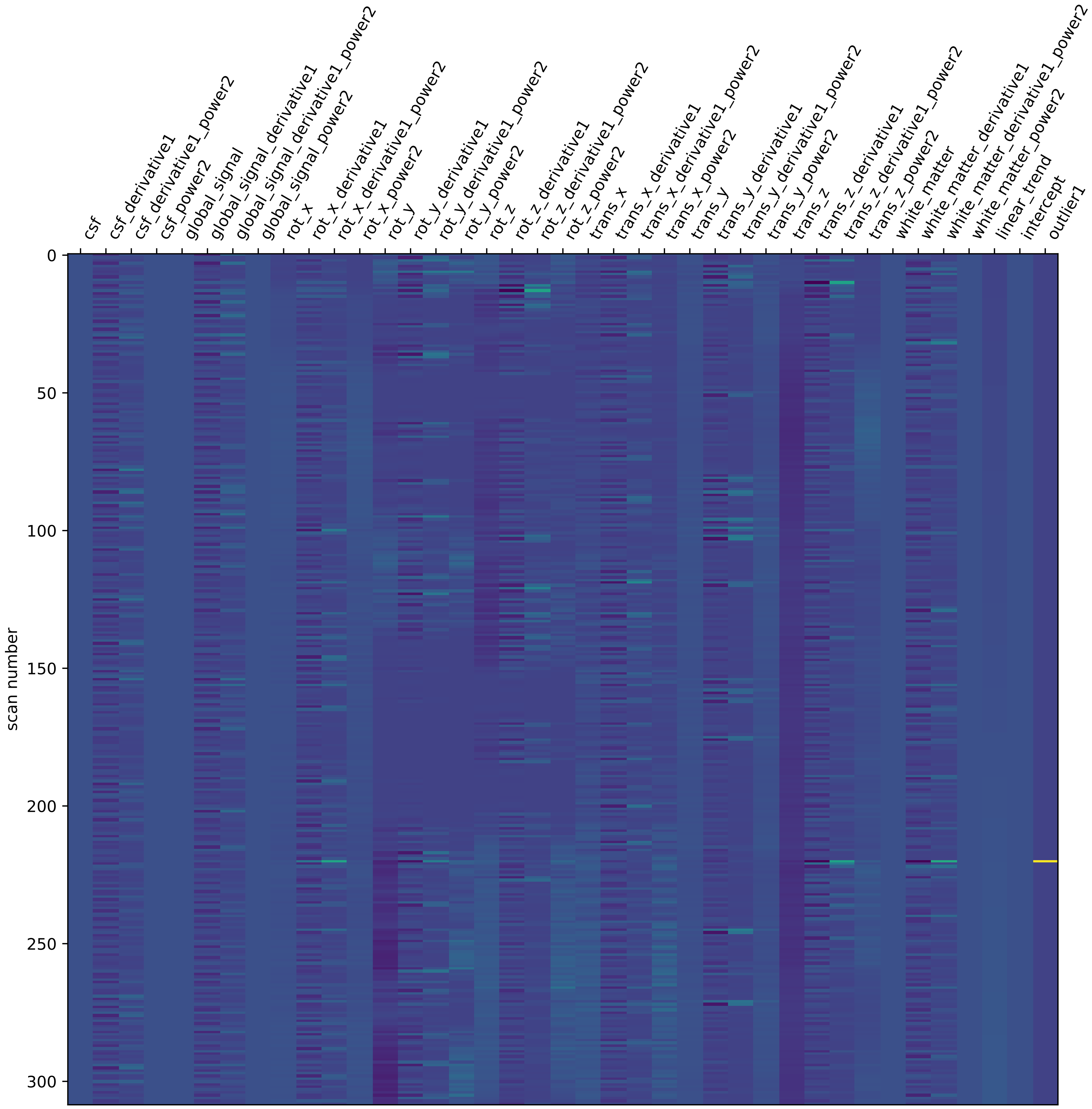


Get figure file: sub-113922/figures/sub-113922_task-LANGUAGE_dir-LR_design.svg

#### Carpet Plot After Postprocessing

FD and DVARS are two measures of in-scanner motion. This plot shows standardized FD, DVARS, and then a carpet plot for the time series of each voxel/vertex's time series of activity.

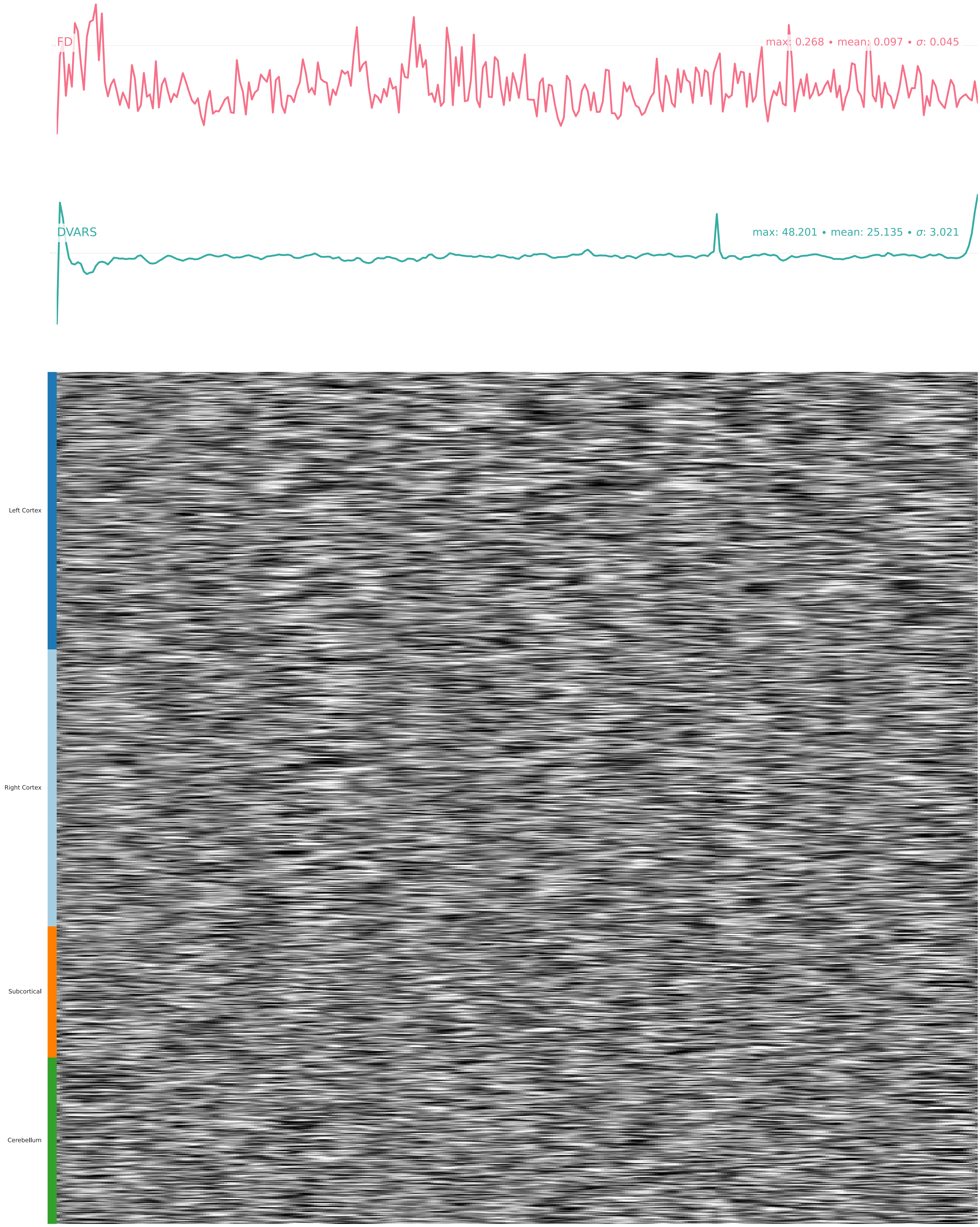


Get figure file: sub-113922/figures/sub-113922_task-LANGUAGE_dir-LR_space-fsLR_desc-postprocessing_bold.svg

#### Correlation Heatmaps from Four Atlases

This plot shows heatmaps from ROI-to-ROI correlations from four atlases.

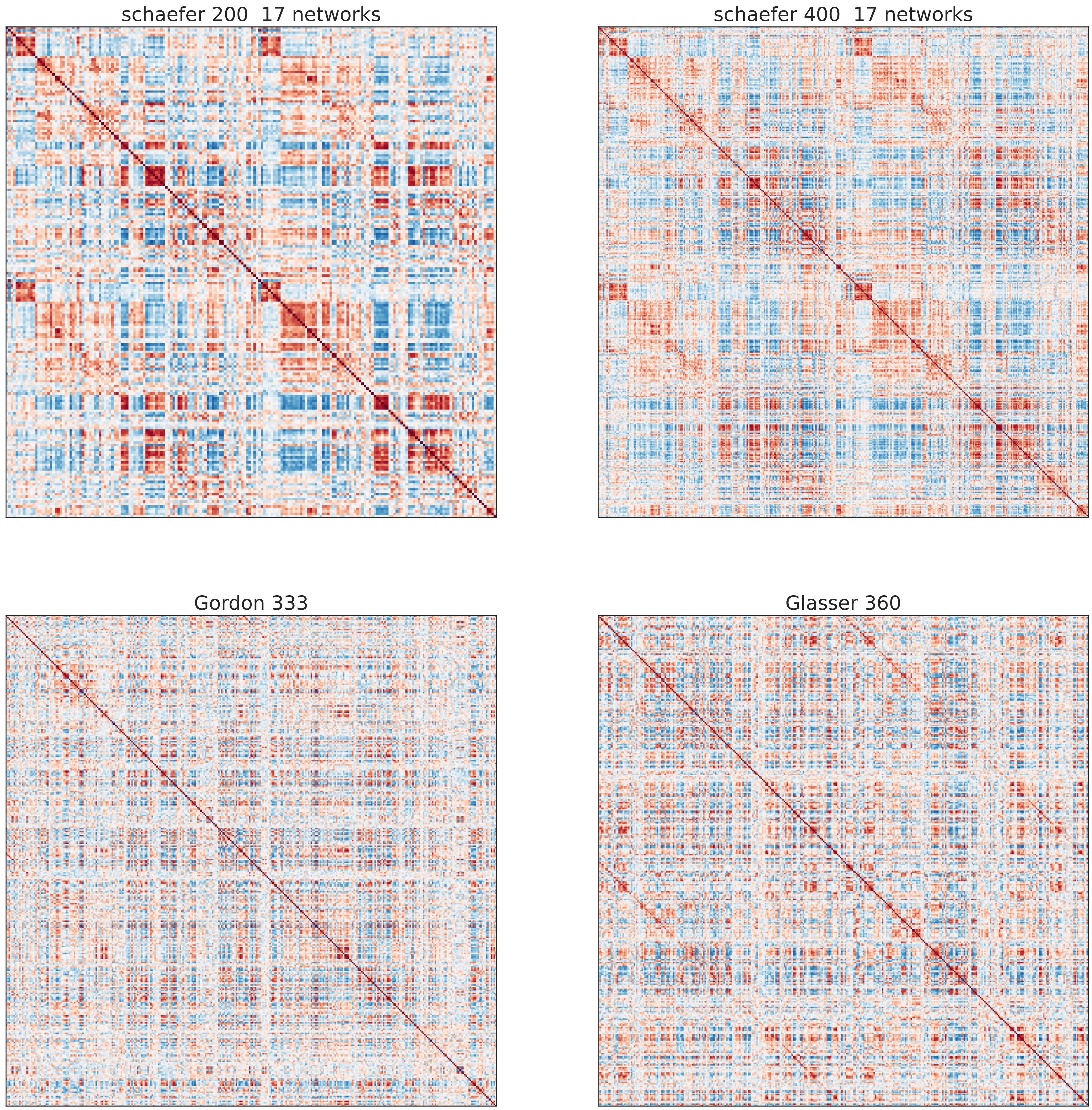


Get figure file: sub-113922/figures/sub-113922_task-LANGUAGE_dir-LR_space-fsLR_desc-connectivityplot_bold.svg

### Reports for: task LANGUAGE, direction RL.

#### Summary

- BOLD volume space: fsLR
- Repetition Time (TR): 0.72
- Mean Framewise Displacement: 0.1248
- Mean Relative RMS Motion: 0.19
- Max Relative RMS Motion: 0.3258
- DVARS Before and After Processing : 482.0385, 25.1094
- Correlation between DVARS and FD Before and After Processing : 0.1733, -0.0017
- Number of Volumes Censored : 1

#### Alignment of functional and anatomical MRI data (surface driven)

bbregister was used to coregister functional and anatomical MRI data.

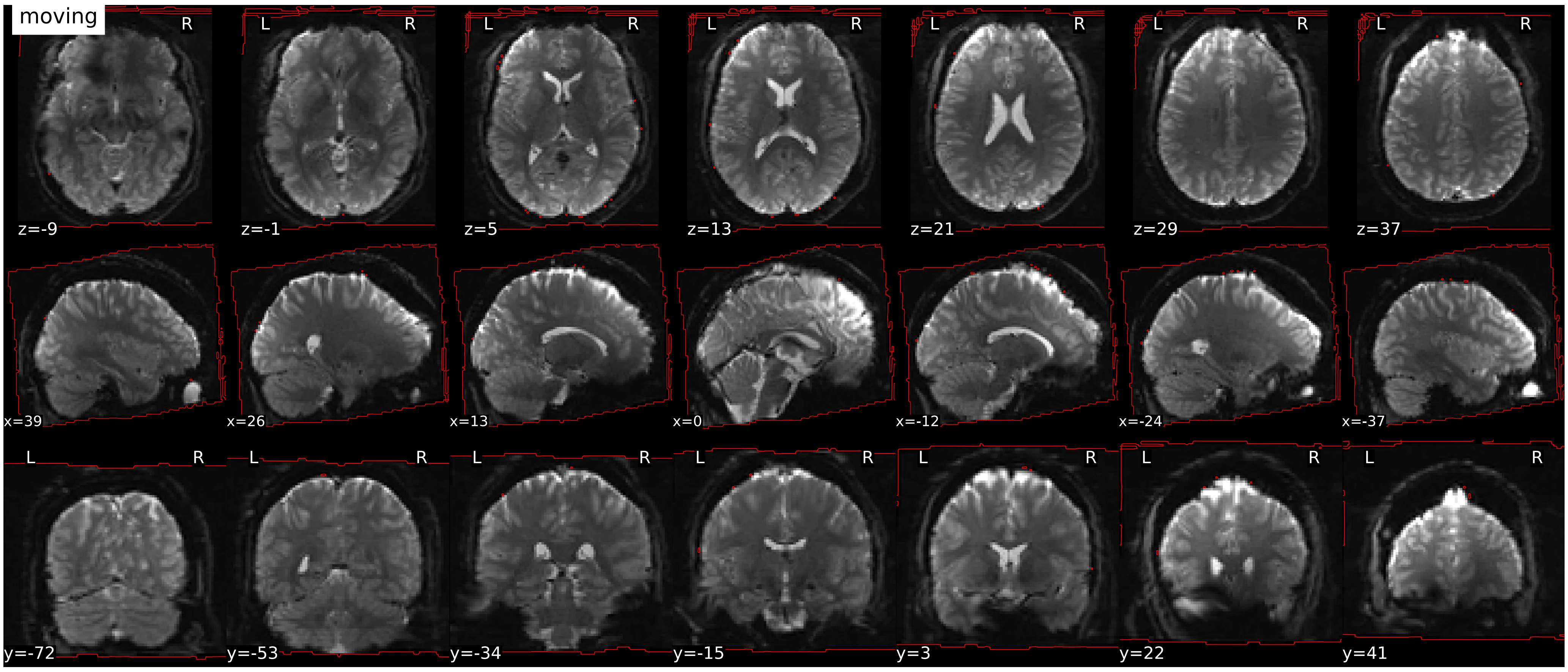


Get figure file: sub-113922/figures/sub-113922_task-LANGUAGE_dir-RL_space-MNI152NLin6Asym_desc-bbregister_bold.svg

#### Carpet Plot Before Postprocessing

FD and DVARS are two measures of in-scanner motion. This plot shows standardized FD, DVARS, and then a carpet plot for the time series of each voxel/vertex's time series of activity.

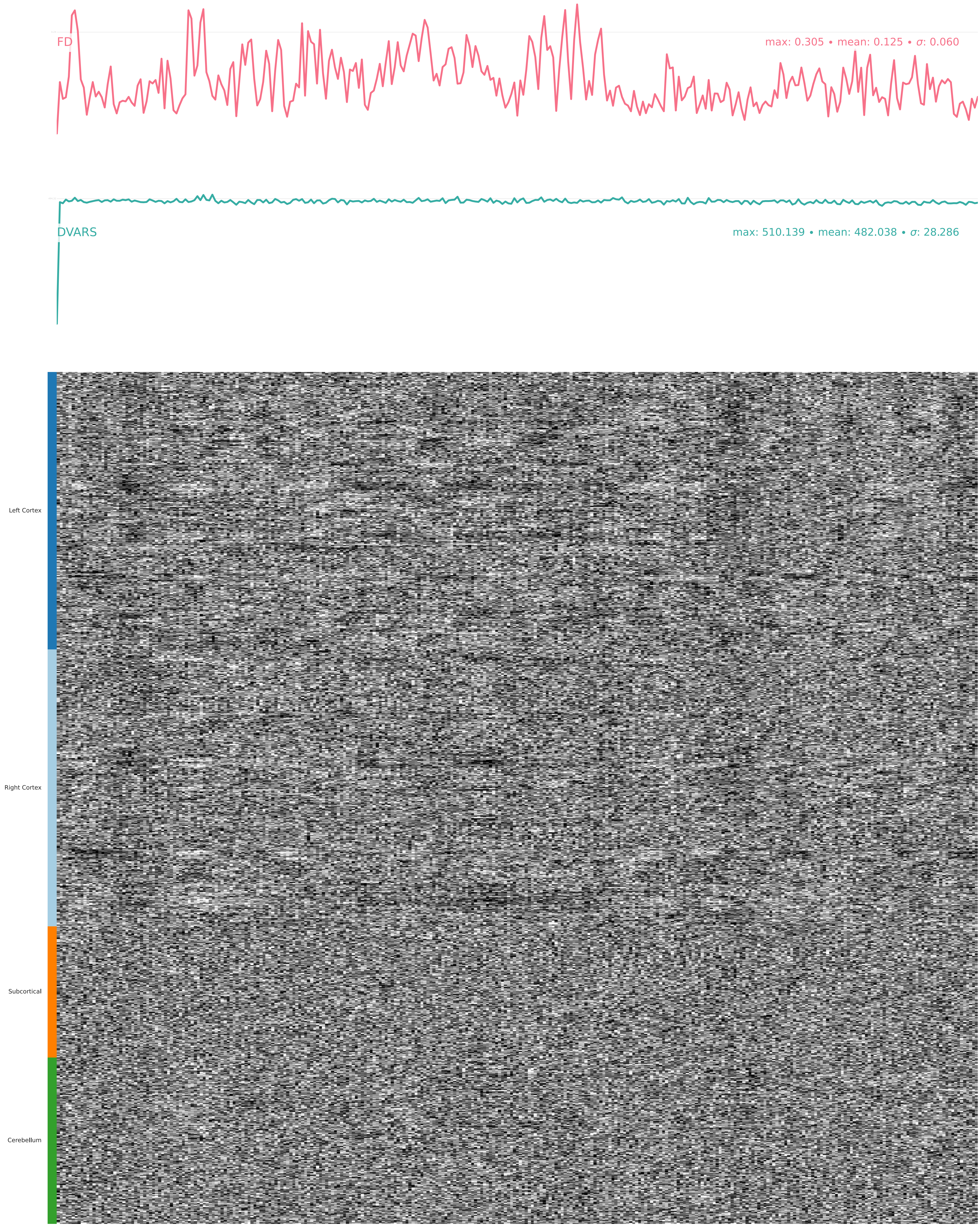


Get figure file: [sub-113922/figures/sub-113922_task-LANGUAGE_dir-RL_space-fsLR_desc-preprocessing_bold.svg](sub-113922/figures/sub-113922_task-LANGUAGE_dir-RL_space-fsLR_desc-preprocessing_bold.svg)

#### Framewise Displacement and Censored Volumes

Framewise displacement (FD) is used to flag high-motion volumes, which are then censored as part of the denoising procedure. If motion filtering is requested, then the six translation and rotation motion parameters are filtered to remove respiratory effects before FD is calculated and outlier volumes are identified.

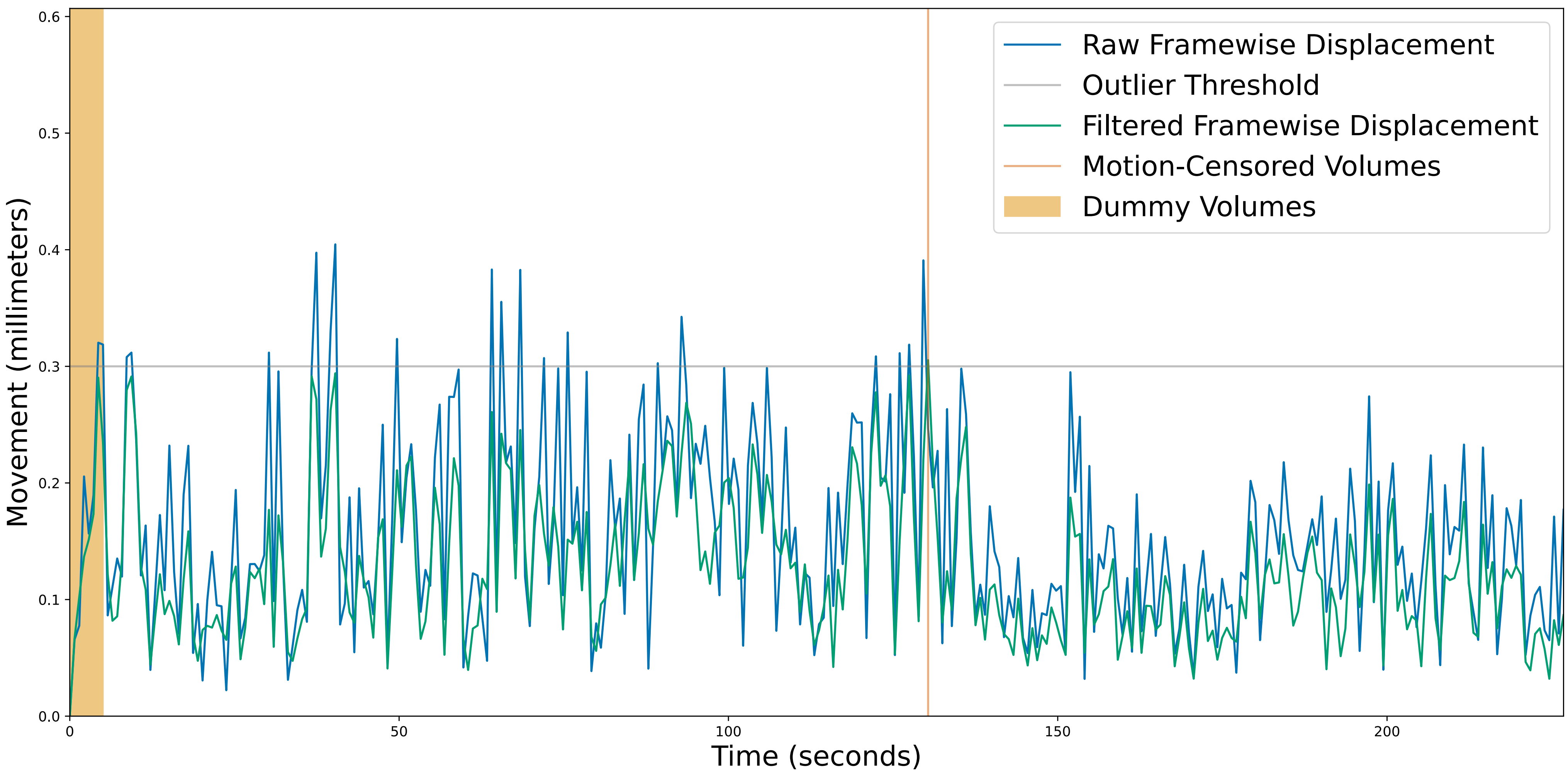


Get figure file: [sub-113922/figures/sub-113922_task-LANGUAGE_dir-RL_space-fsLR_desc-censoring_motion.svg](sub-113922/figures/sub-113922_task-LANGUAGE_dir-RL_space-fsLR_desc-censoring_motion.svg)

#### Design Matrix for Confound Regression

The "design matrix" represents the confounds that are used to denoise the BOLD data.

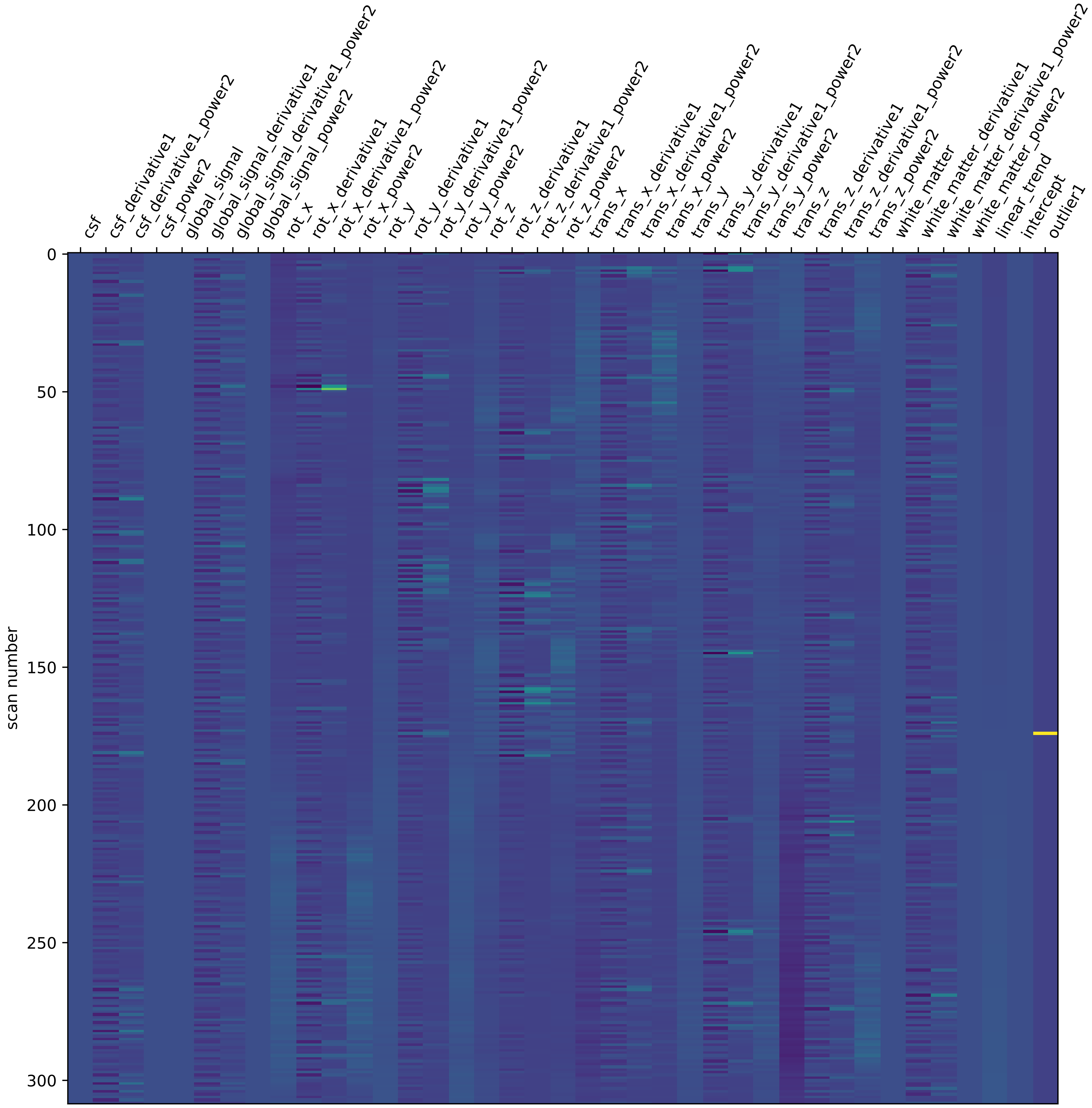


Get figure file: [sub-113922/figures/sub-113922_task-LANGUAGE_dir-RL_design.svg](sub-113922/figures/sub-113922_task-LANGUAGE_dir-RL_design.svg)

#### Carpet Plot After Postprocessing

FD and DVARS are two measures of in-scanner motion. This plot shows standardized FD, DVARS, and then a carpet plot for the time series of each voxel/vertex's time series of activity.

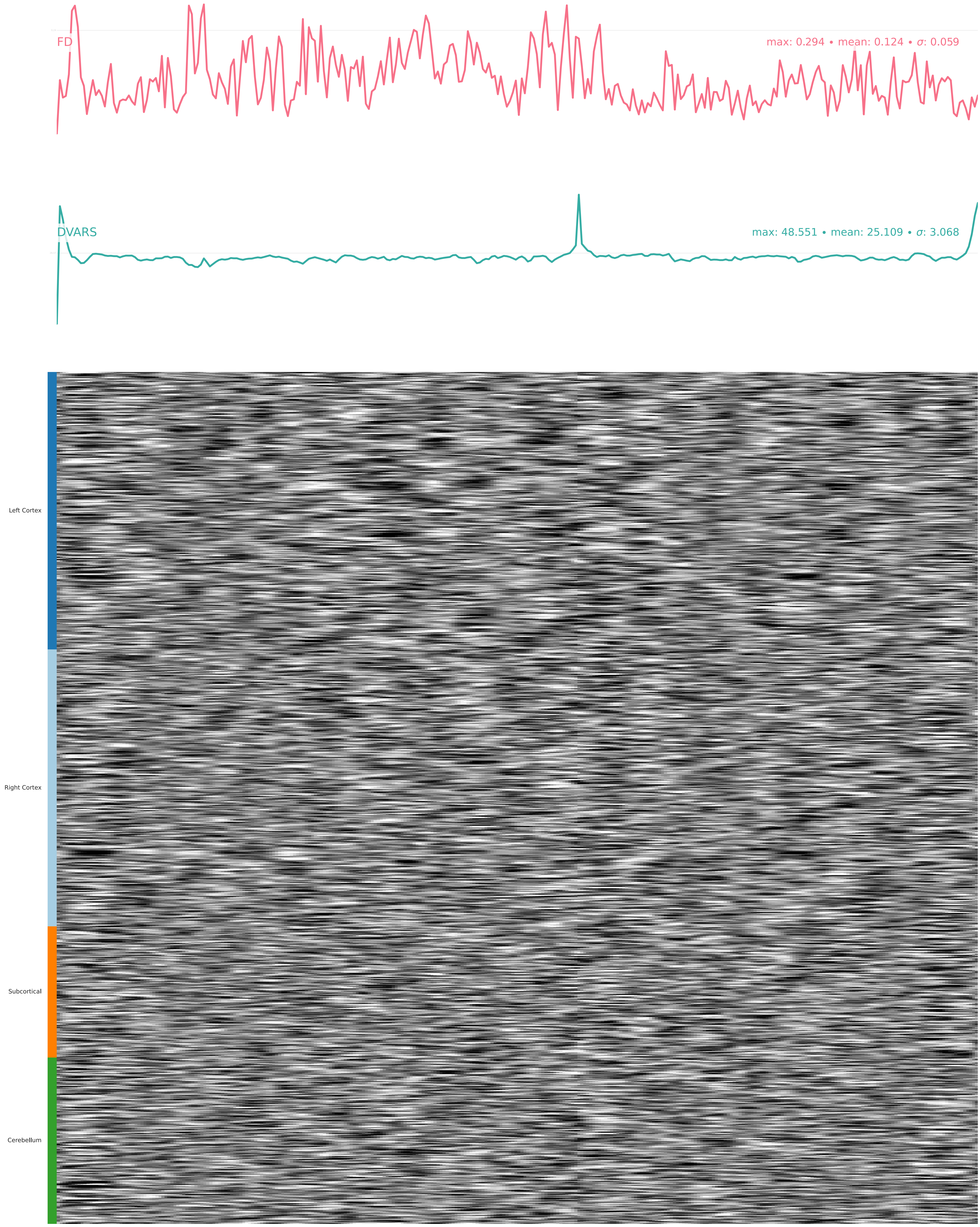


Get figure file: sub-113922/figures/sub-113922_task-LANGUAGE_dir-RL_space-fsLR_desc-postprocessing_bold.svg

#### Correlation Heatmaps from Four Atlases

This plot shows heatmaps from ROI-to-ROI correlations from four atlases.

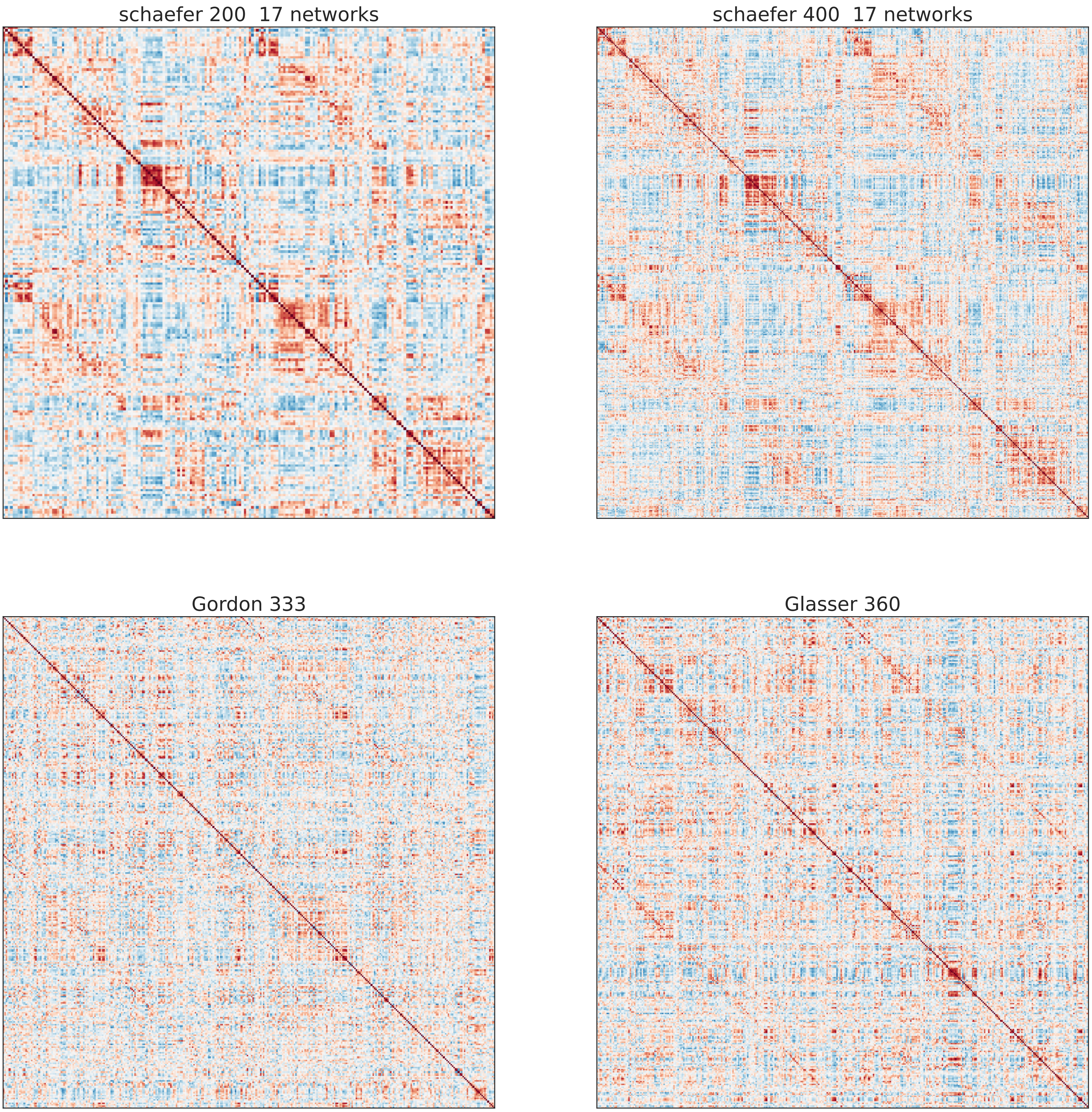


Get figure file: sub-113922/figures/sub-113922_task-LANGUAGE_dir-RL_space-fsLR_desc-connectivityplot_bold.svg

### Reports for: task MOTOR, direction LR.

#### Summary

- BOLD volume space: fsLR
- Repetition Time (TR): 0.72
- Mean Framewise Displacement: 0.1293
- Mean Relative RMS Motion: 0.2346
- Max Relative RMS Motion: 0.4636
- DVARS Before and After Processing : 472.9467, 162.9215
- Correlation between DVARS and FD Before and After Processing : 0.4314, 0.0208
- Number of Volumes Censored : 10

#### Alignment of functional and anatomical MRI data (surface driven)

bbregister was used to coregister functional and anatomical MRI data.

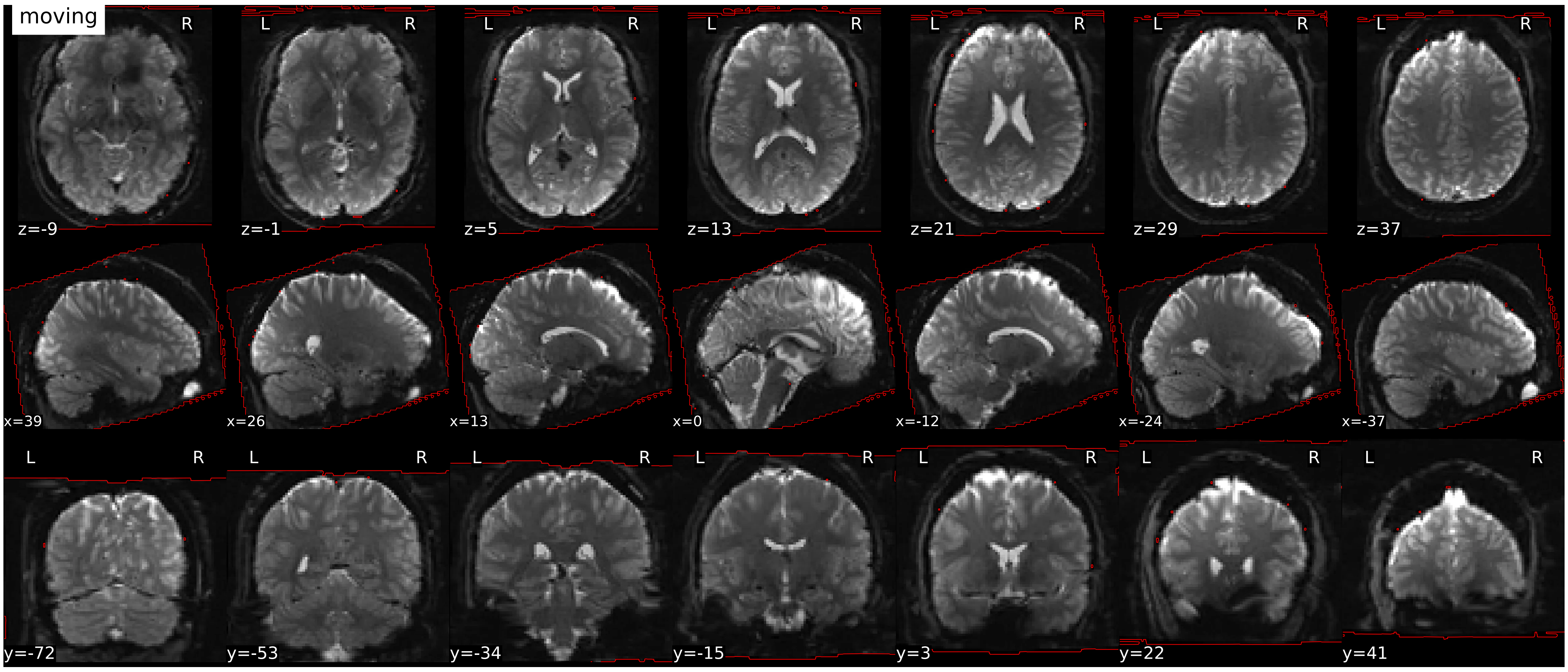


Get figure file: sub-113922/figures/sub-113922_task-MOTOR_dir-LR_space-MNI152NLin6Asym_desc-bbregister_bold.svg

#### Carpet Plot Before Postprocessing

FD and DVARS are two measures of in-scanner motion. This plot shows standardized FD, DVARS, and then a carpet plot for the time series of each voxel/vertex's time series of activity.

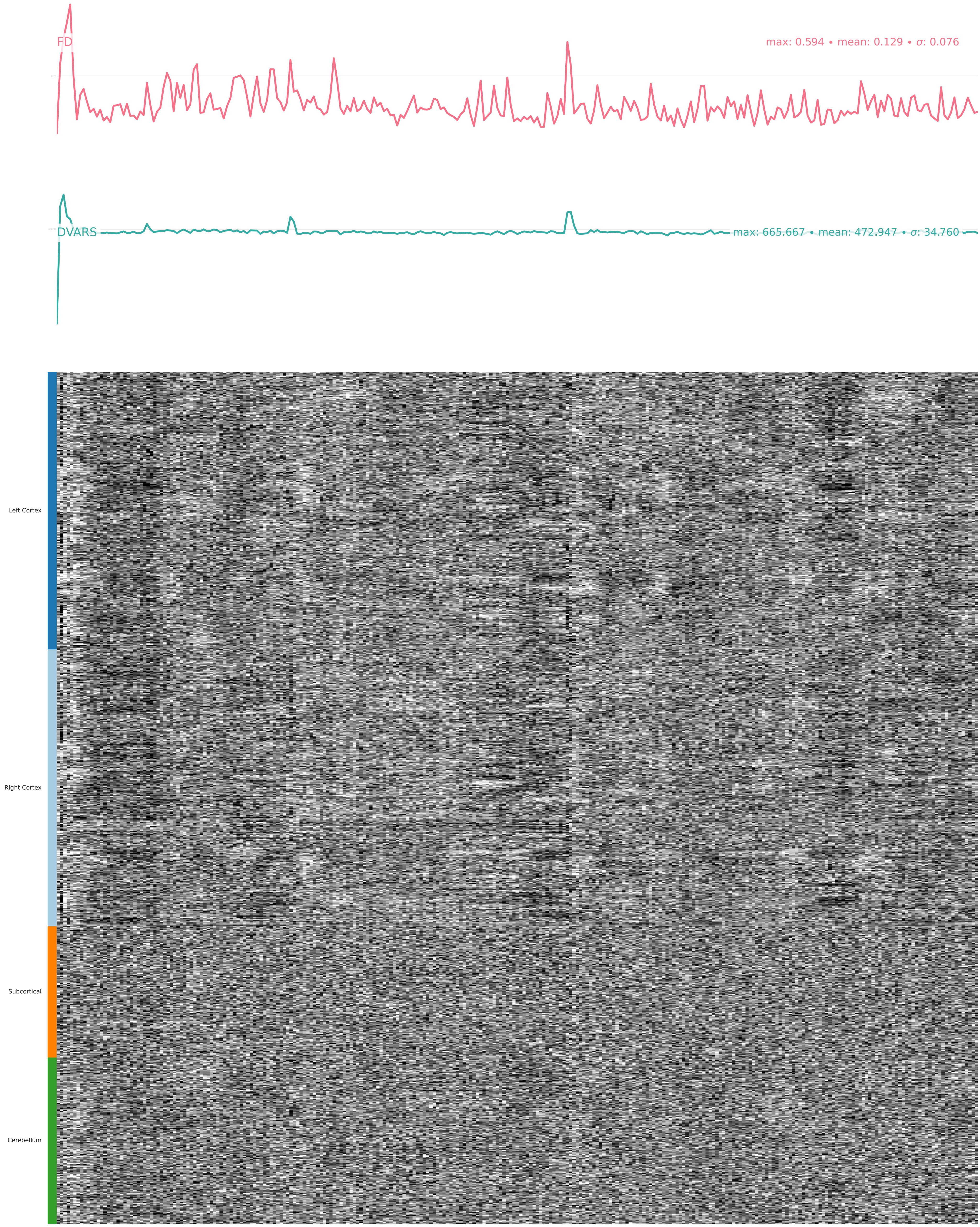


Get figure file: [sub-113922/figures/sub-113922_task-MOTOR_dir-LR_space-fsLR_desc-preprocessing_bold.svg](sub-113922/figures/sub-113922_task-MOTOR_dir-LR_space-fsLR_desc-preprocessing_bold.svg)

#### Framewise Displacement and Censored Volumes

Framewise displacement (FD) is used to flag high-motion volumes, which are then censored as part of the denoising procedure. If motion filtering is requested, then the six translation and rotation motion parameters are filtered to remove respiratory effects before FD is calculated and outlier volumes are identified.

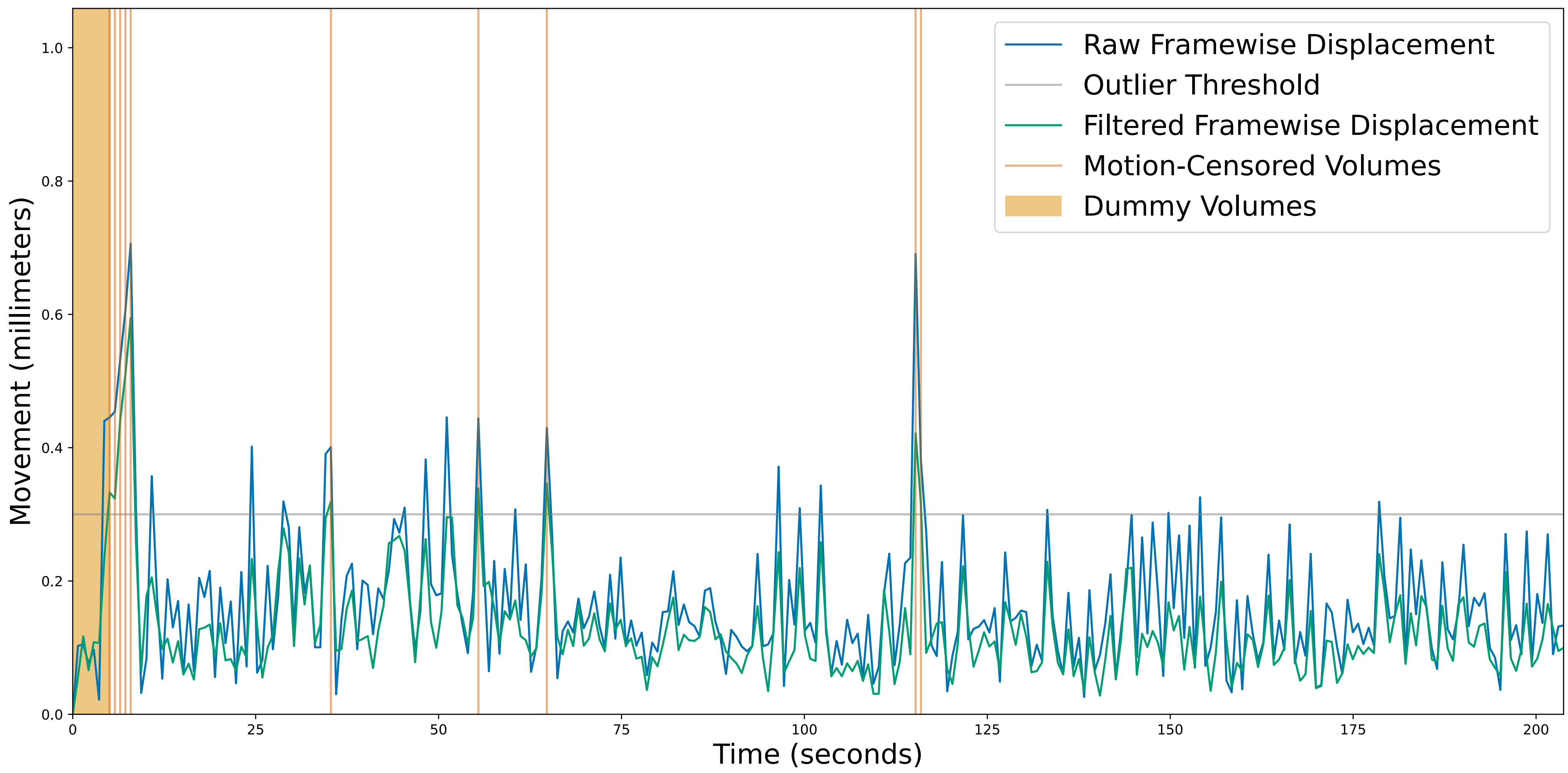


Get figure file: sub-113922/figures/sub-113922_task-MOTOR_dir-LR_space-fsLR_desc-censoring_motion.svg

#### Design Matrix for Confound Regression

The "design matrix" represents the confounds that are used to denoise the BOLD data.

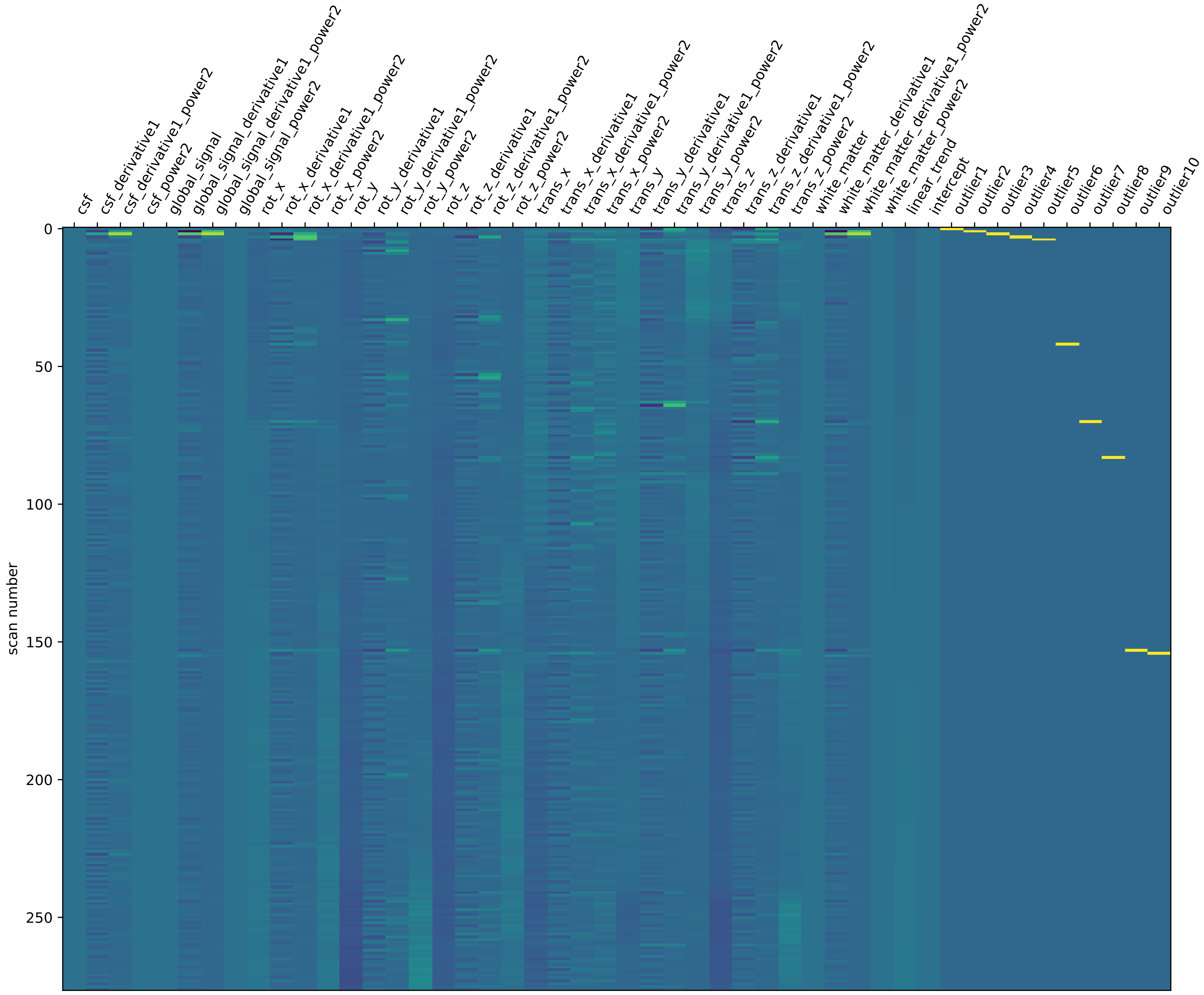


Get figure file: sub-113922/figures/sub-113922_task-MOTOR_dir-LR_design.svg

### Carpet Plot After Postprocessing

FD and DVARS are two measures of in-scanner motion. This plot shows standardized FD, DVARS, and then a carpet plot for the time series of each voxel/vertex's time series of activity.

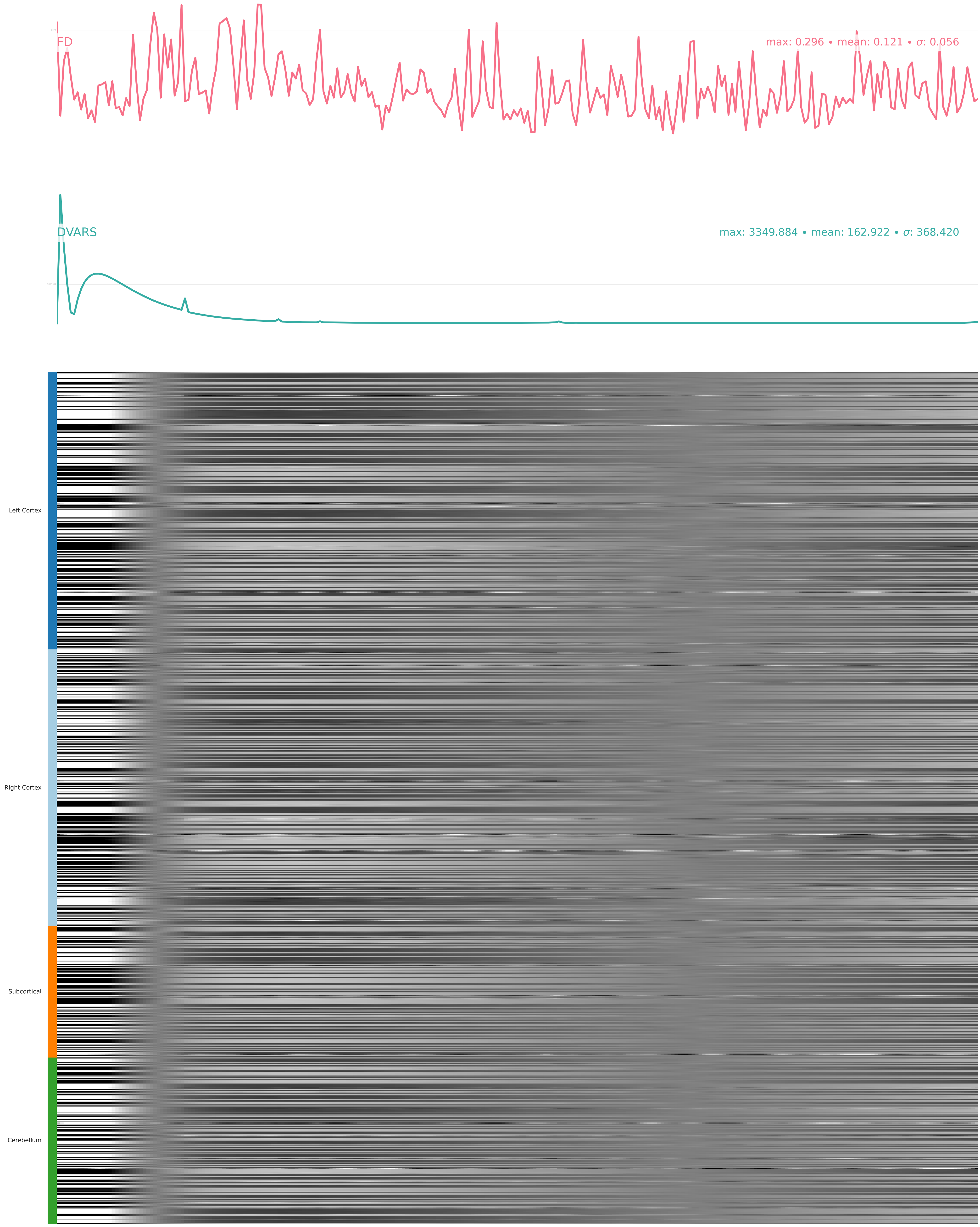


Get figure file: sub-113922/figures/sub-113922_task-MOTOR_dir-LR_space-fsLR_desc-postprocessing_bold.svg

#### Correlation Heatmaps from Four Atlases

This plot shows heatmaps from ROI-to-ROI correlations from four atlases.

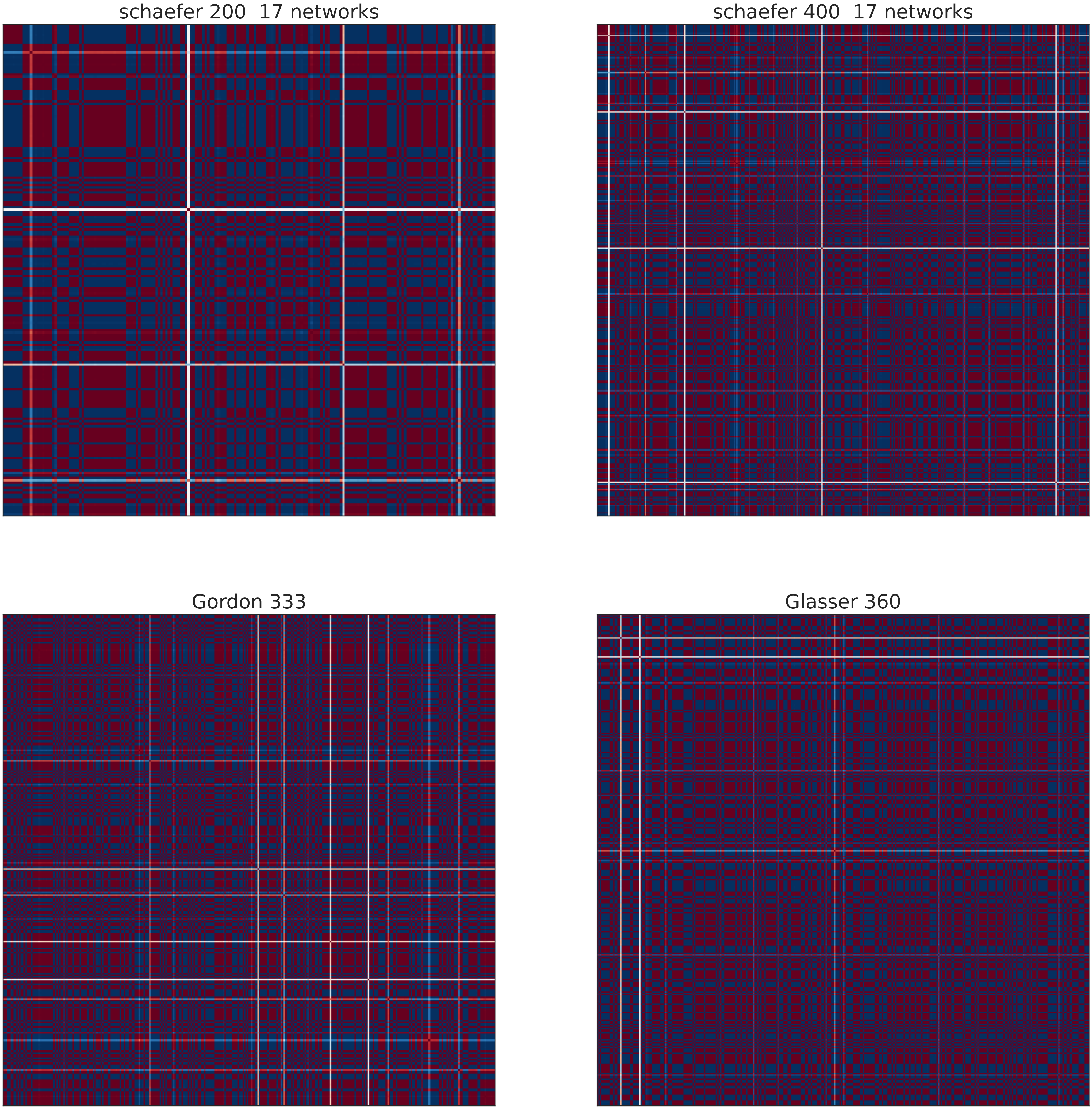


Get figure file: [sub-113922/figures/sub-113922_task-MOTOR_dir-LR_space-fsLR_desc-connectivityplot_bold.svg](sub-113922/figures/sub-113922_task-MOTOR_dir-LR_space-fsLR_desc-connectivityplot_bold.svg)

### Reports for: task MOTOR, direction RL.

#### Summary

- BOLD volume space: fsLR
- Repetition Time (TR): 0.72
- Mean Framewise Displacement: 0.1569
- Mean Relative RMS Motion: 0.3683
- Max Relative RMS Motion: 0.8812
- DVARS Before and After Processing : 479.2293, 30.2661
- Correlation between DVARS and FD Before and After Processing : 0.7269, 0.2294
- Number of Volumes Censored : 20

#### Alignment of functional and anatomical MRI data (surface driven)

bbregister was used to coregister functional and anatomical MRI data.

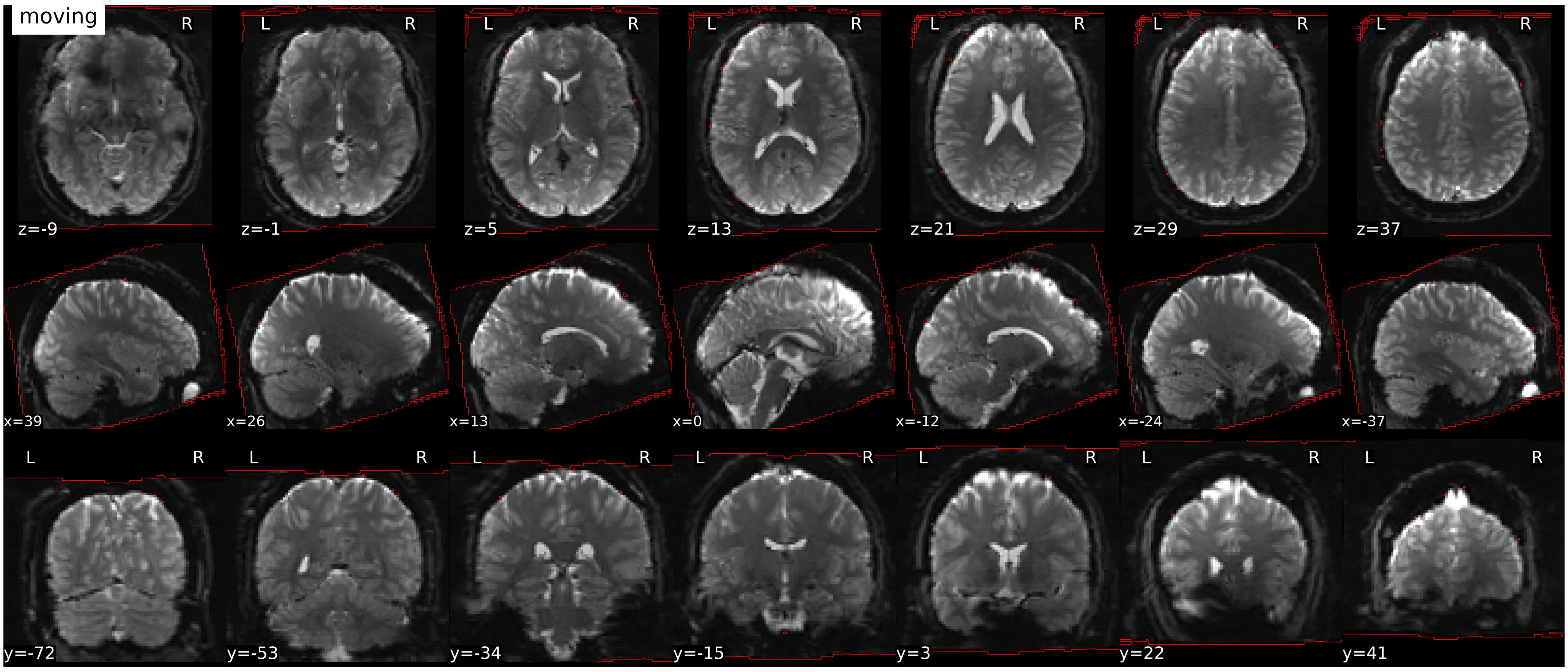


Get figure file: sub-113922/figures/sub-113922_task-MOTOR_dir-RL_space-MNI152NLin6Asym_desc-bbregister_bold.svg

#### Carpet Plot Before Postprocessing

FD and DVARS are two measures of in-scanner motion. This plot shows standardized FD, DVARS, and then a carpet plot for the time series of each voxel/vertex's time series of activity.

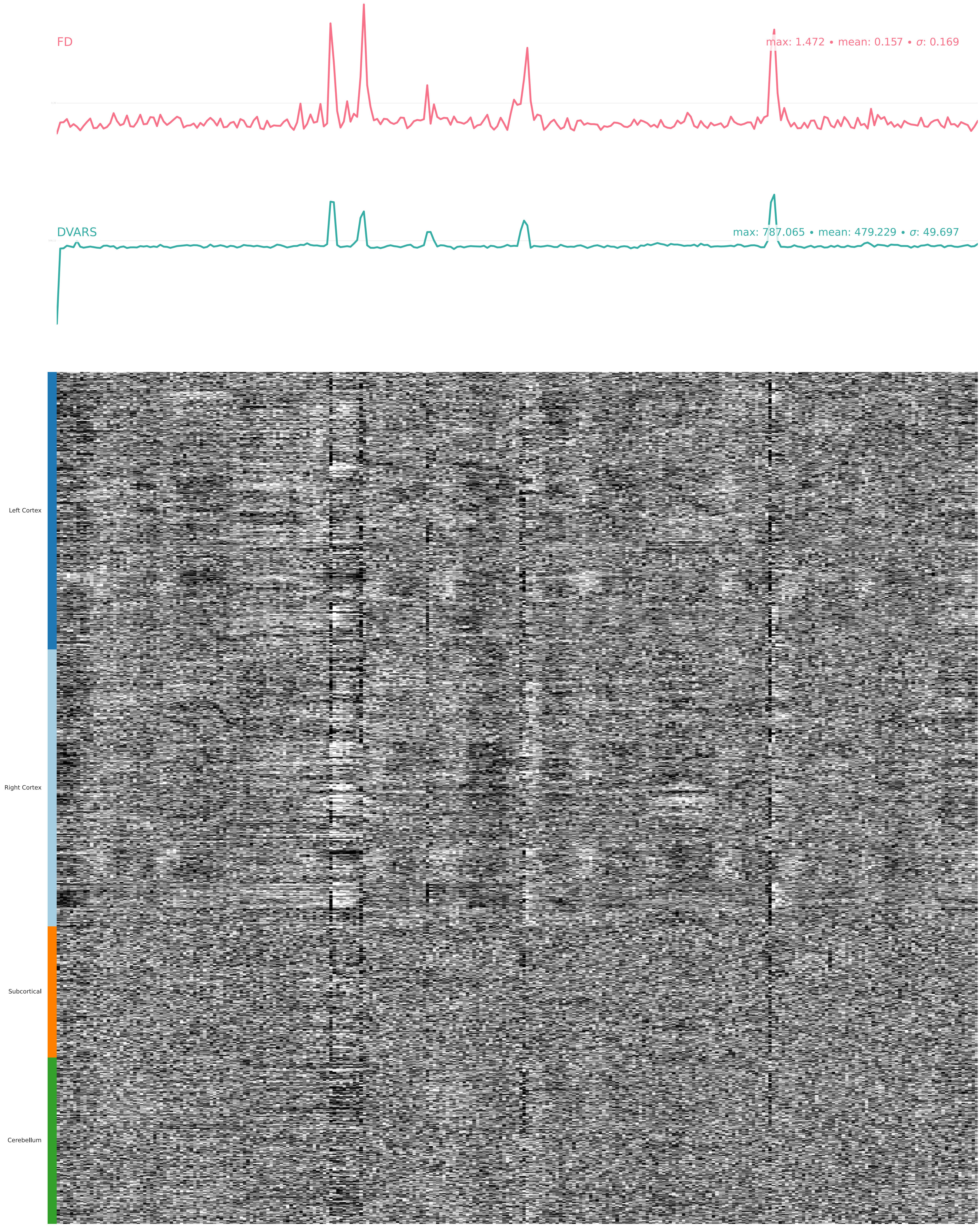


Get figure file: sub-113922/figures/sub-113922_task-MOTOR_dir-RL_space-fsLR_desc-preprocessing_bold.svg

#### Framewise Displacement and Censored Volumes

Framewise displacement (FD) is used to flag high-motion volumes, which are then censored as part of the denoising procedure. If motion filtering is requested, then the six translation and rotation motion parameters are filtered to remove respiratory effects before FD is calculated and outlier volumes are identified.

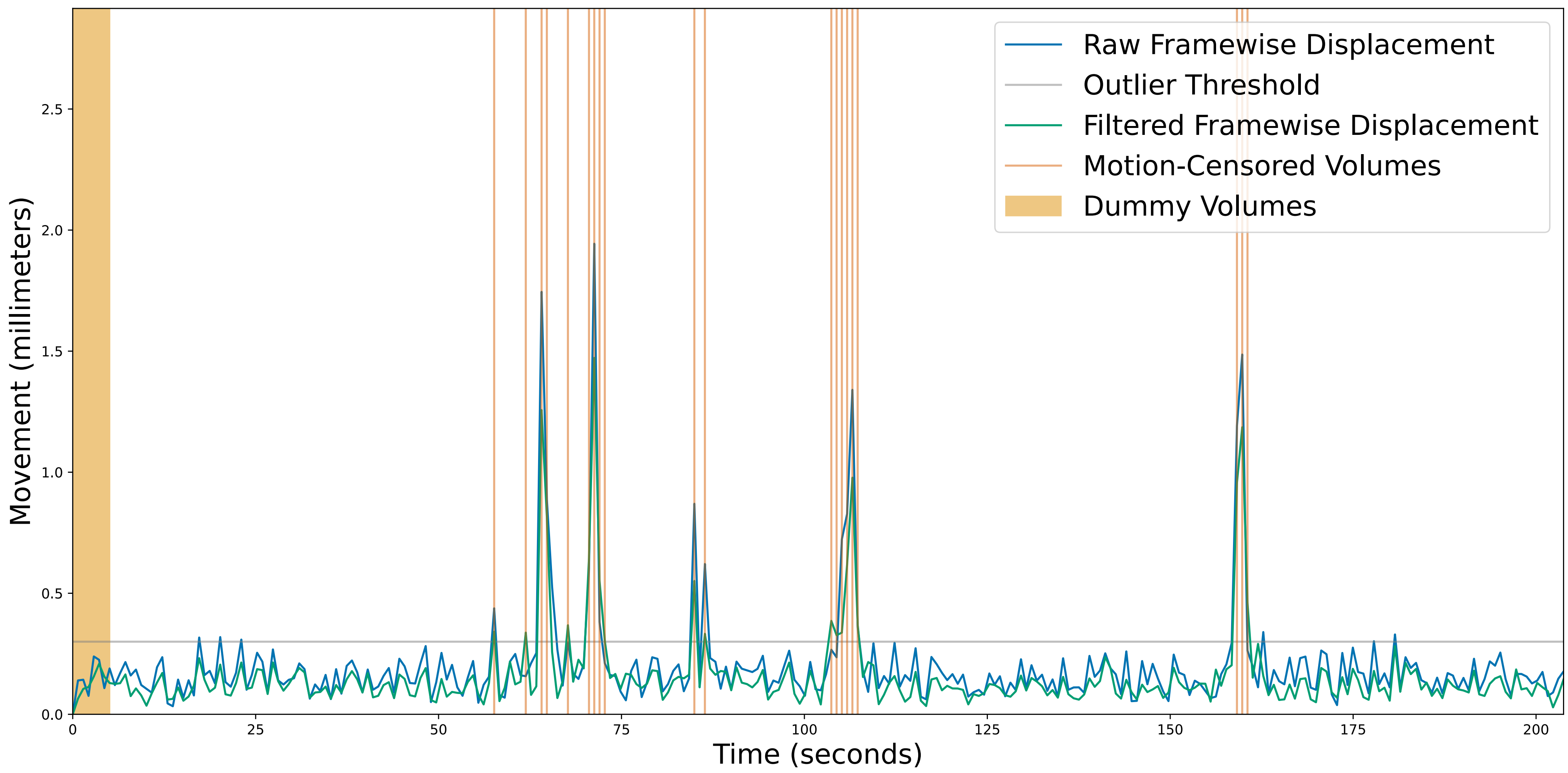


Get figure file: [sub-113922/figures/sub-113922_task-MOTOR_dir-RL_space-fsLR_desc-censoring_motion.svg](sub-113922/figures/sub-113922_task-MOTOR_dir-RL_space-fsLR_desc-censoring_motion.svg)

#### Design Matrix for Confound Regression

The "design matrix" represents the confounds that are used to denoise the BOLD data.

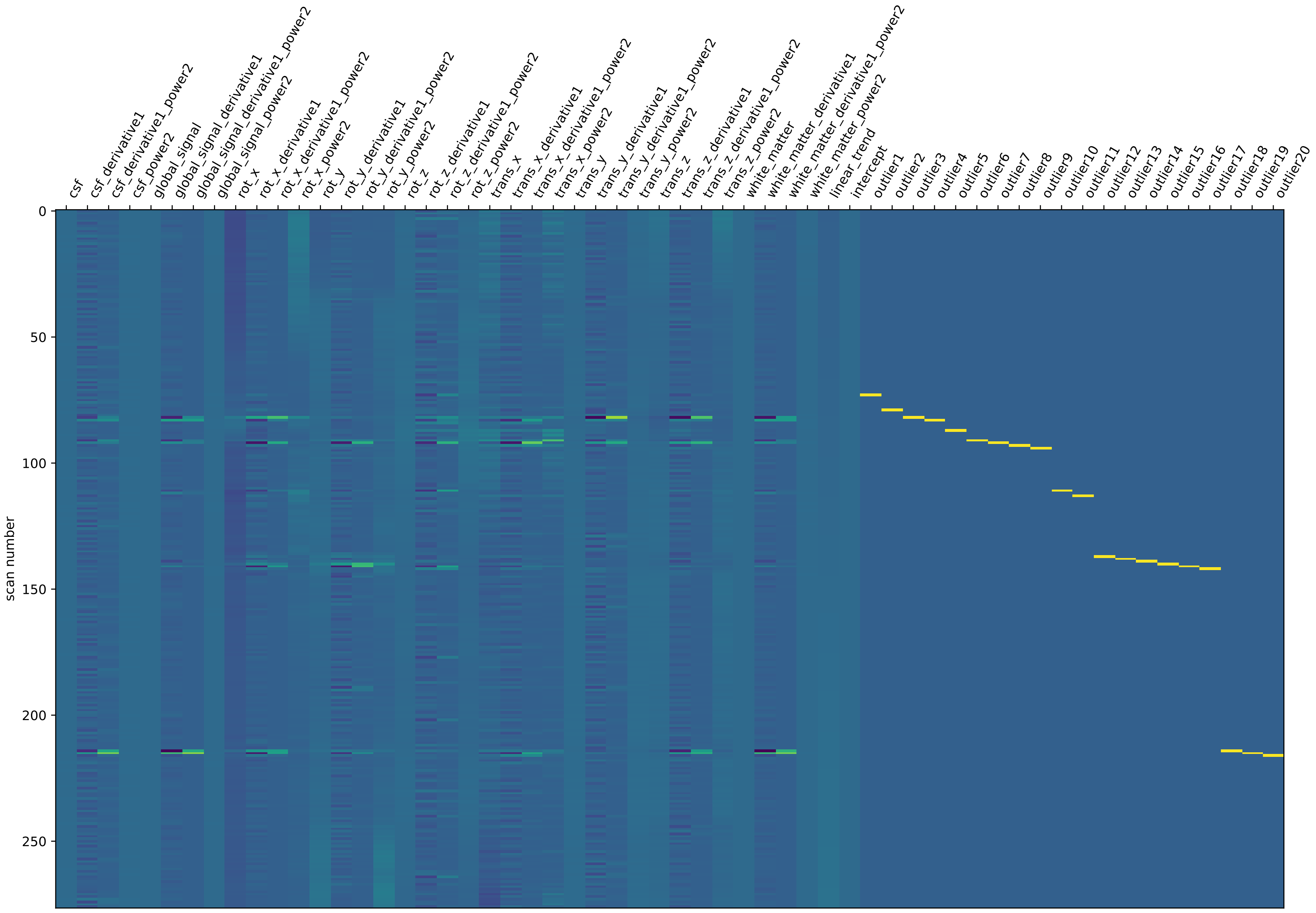


Get figure file: [sub-113922/figures/sub-113922_task-MOTOR_dir-RL_design.svg](sub-113922/figures/sub-113922_task-MOTOR_dir-RL_design.svg)

#### Carpet Plot After Postprocessing

FD and DVARS are two measures of in-scanner motion. This plot shows standardized FD, DVARS, and then a carpet plot for the time series of each voxel/vertex's time series of activity.

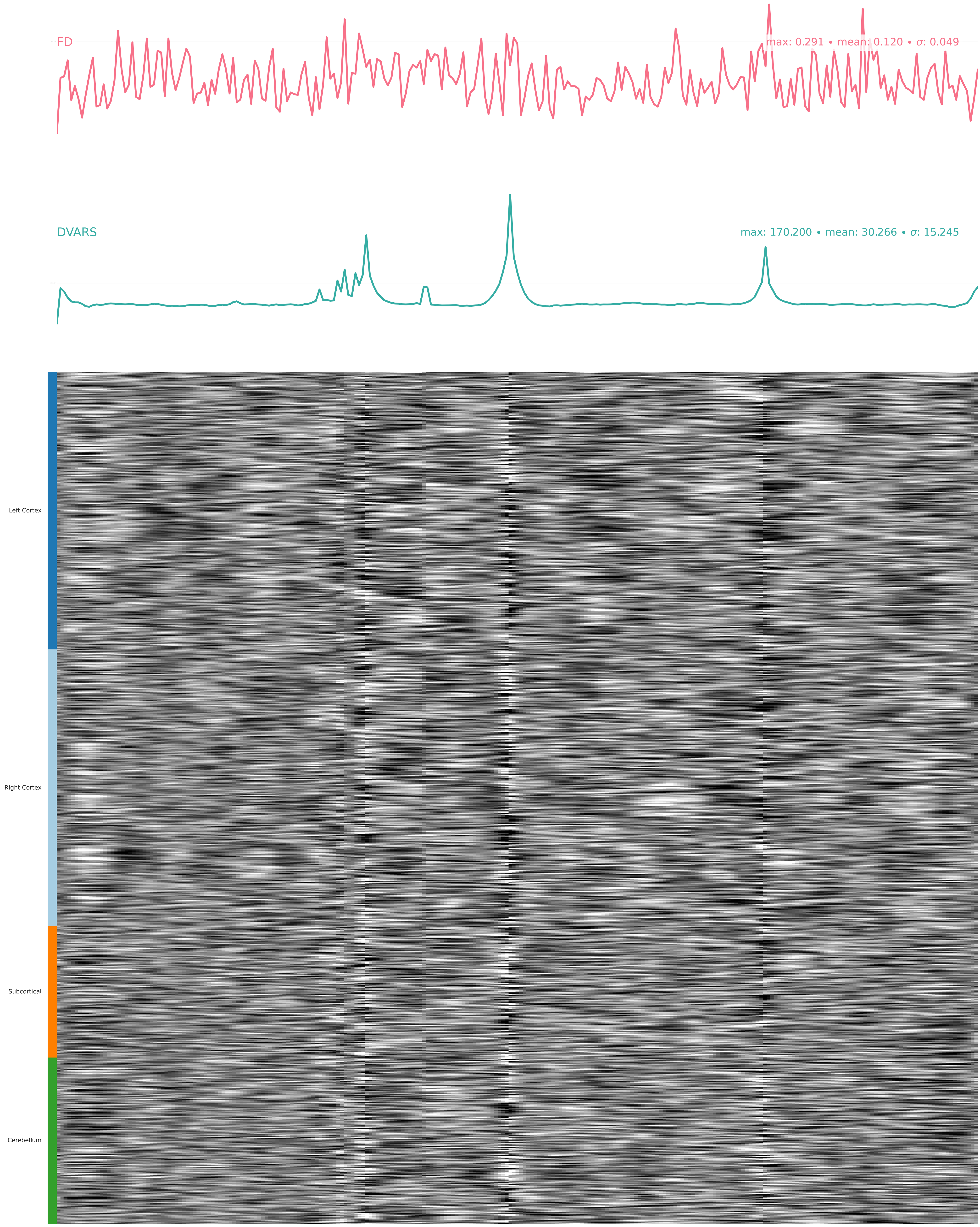


Get figure file: sub-113922/figures/sub-113922_task-MOTOR_dir-RL_space-fsLR_desc-postprocessing_bold.svg

#### Correlation Heatmaps from Four Atlases

This plot shows heatmaps from ROI-to-ROI correlations from four atlases.

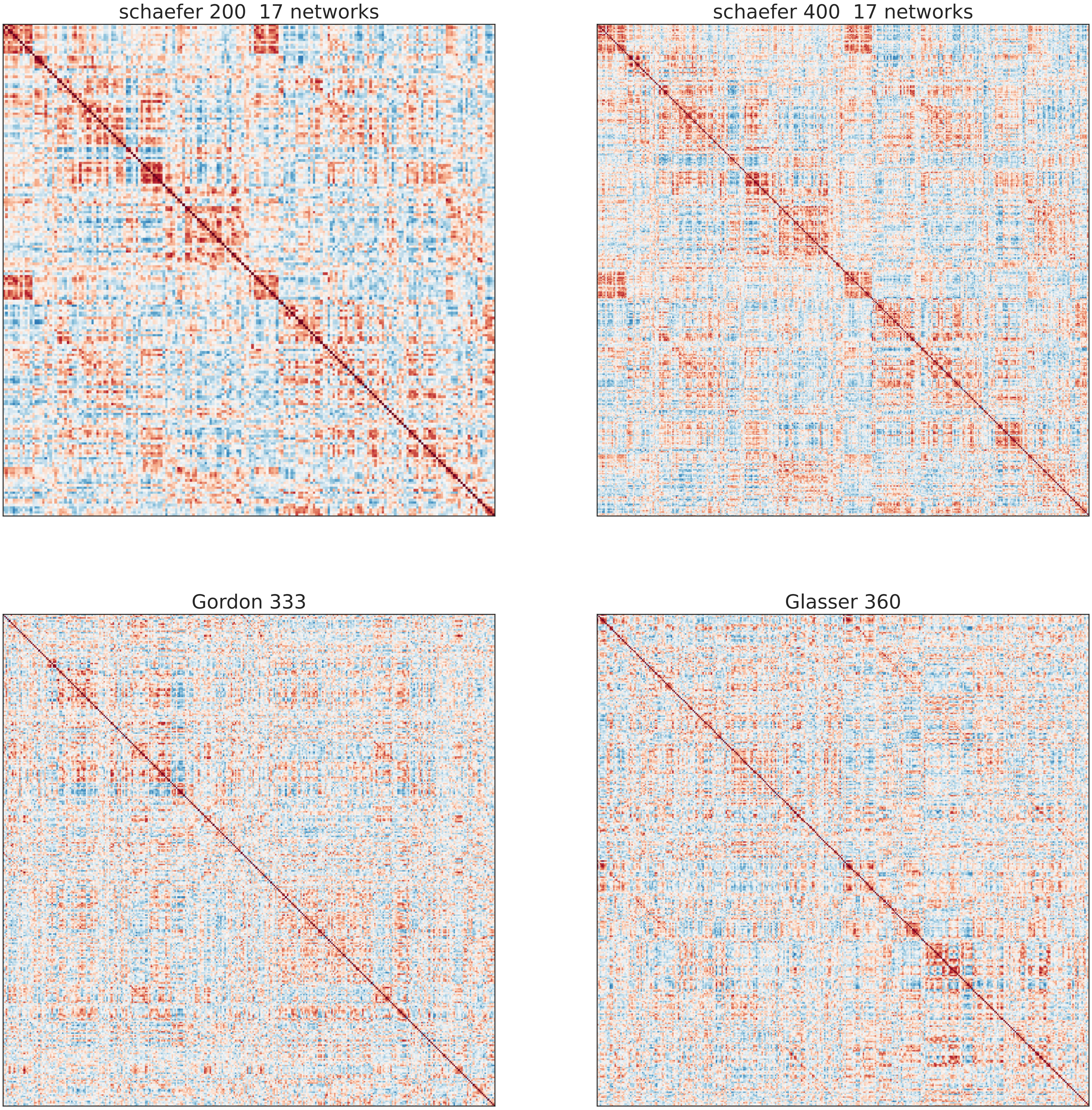


Get figure file: sub-113922/figures/sub-113922_task-MOTOR_dir-RL_space-fsLR_desc-connectivityplot_bold.svg

### Reports for: task RELATIONAL, direction LR.

#### Summary

- BOLD volume space: fsLR
- Repetition Time (TR): 0.72
- Mean Framewise Displacement: 0.1318
- Mean Relative RMS Motion: 0.3973
- Max Relative RMS Motion: 0.5749
- DVARS Before and After Processing : 482.9591, 25.2759
- Correlation between DVARS and FD Before and After Processing : 0.6727, 0.2149
- Number of Volumes Censored : 10

#### Alignment of functional and anatomical MRI data (surface driven)

bbregister was used to coregister functional and anatomical MRI data.

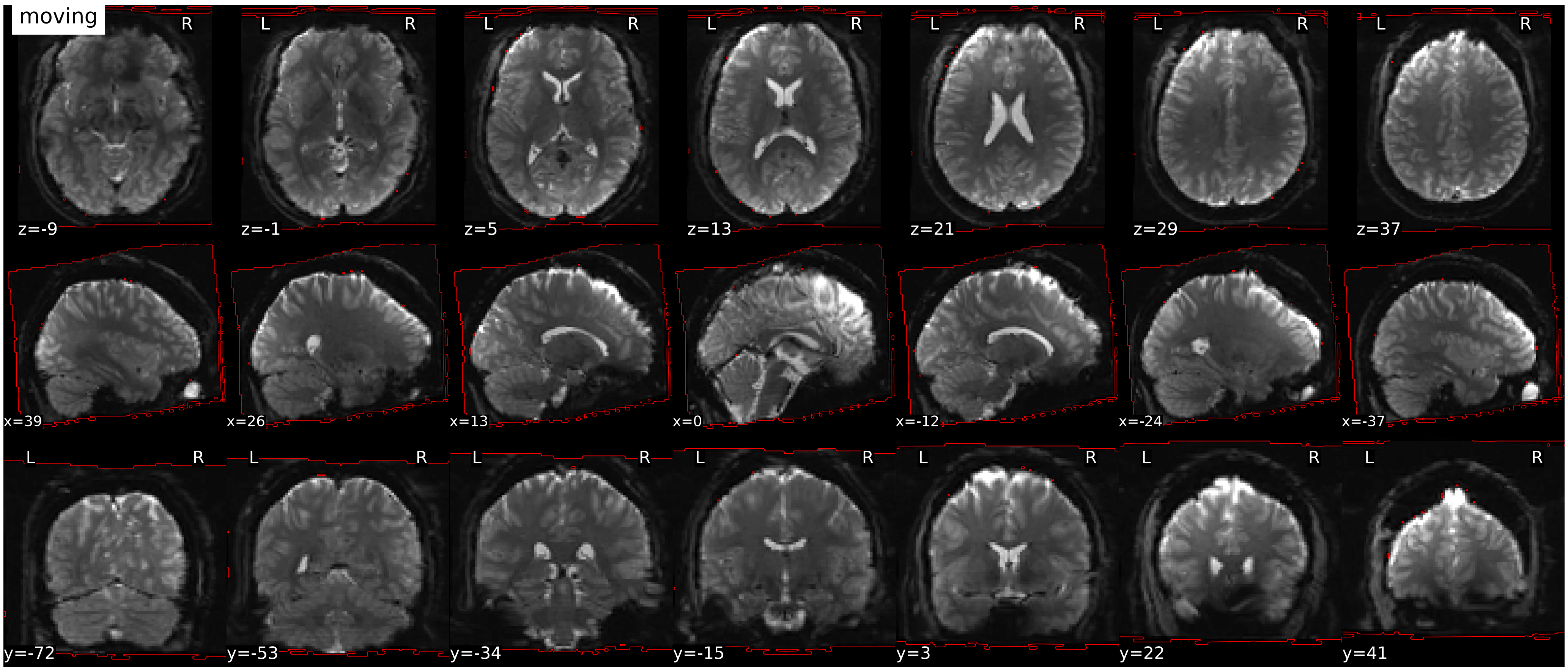


Get figure file: sub-113922/figures/sub-113922_task-RELATIONAL_dir-LR_space-MNI152NLin6Asym_desc-bbregister_bold.svg

#### Carpet Plot Before Postprocessing

FD and DVARS are two measures of in-scanner motion. This plot shows standardized FD, DVARS, and then a carpet plot for the time series of each voxel/vertex's time series of activity.

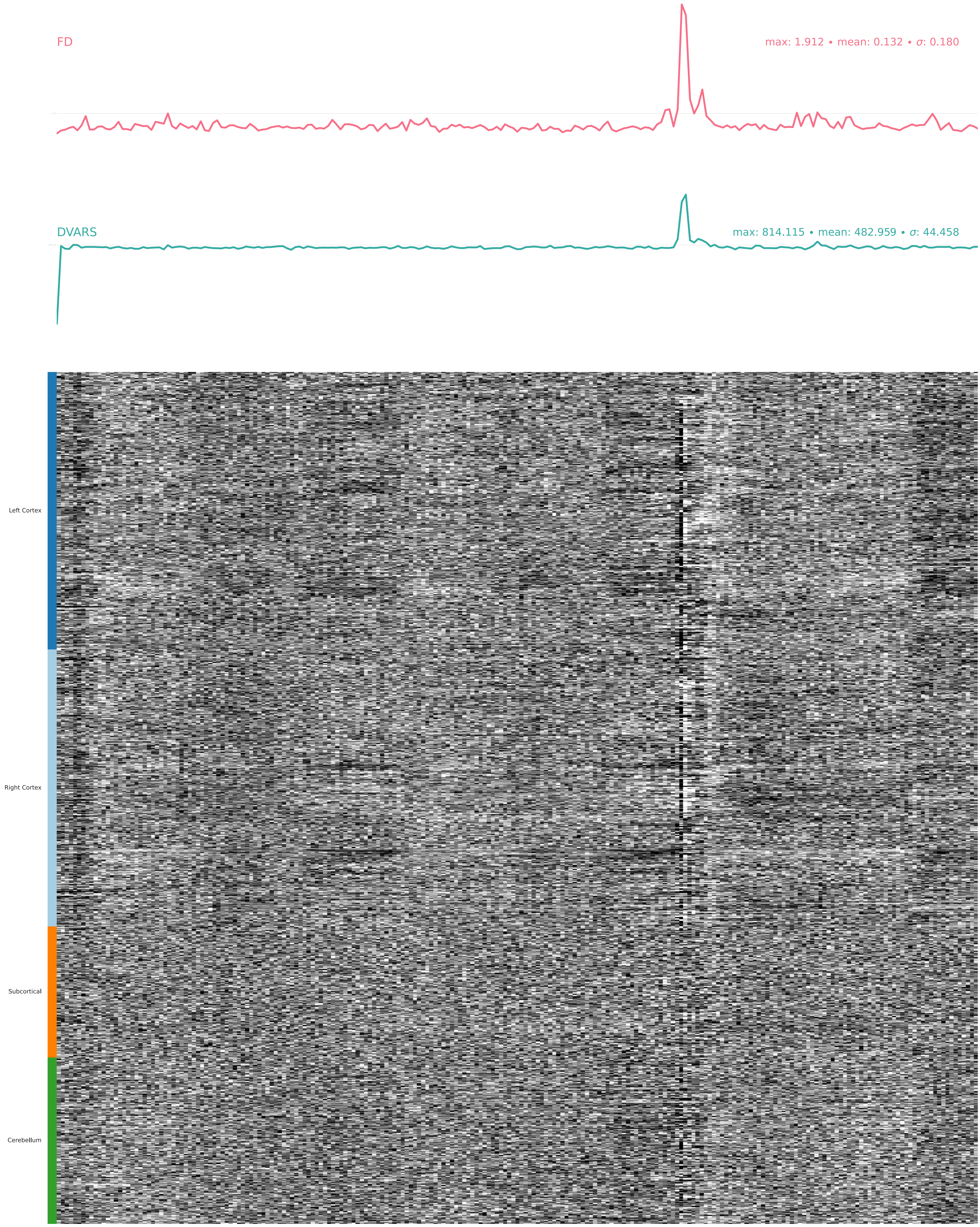


Get figure file: sub-113922/figures/sub-113922_task-RELATIONAL_dir-LR_space-fsLR_desc-preprocessing_bold.svg

#### Framewise Displacement and Censored Volumes

Framewise displacement (FD) is used to flag high-motion volumes, which are then censored as part of the denoising procedure. If motion filtering is requested, then the six translation and rotation motion parameters are filtered to remove respiratory effects before FD is calculated and outlier volumes are identified.

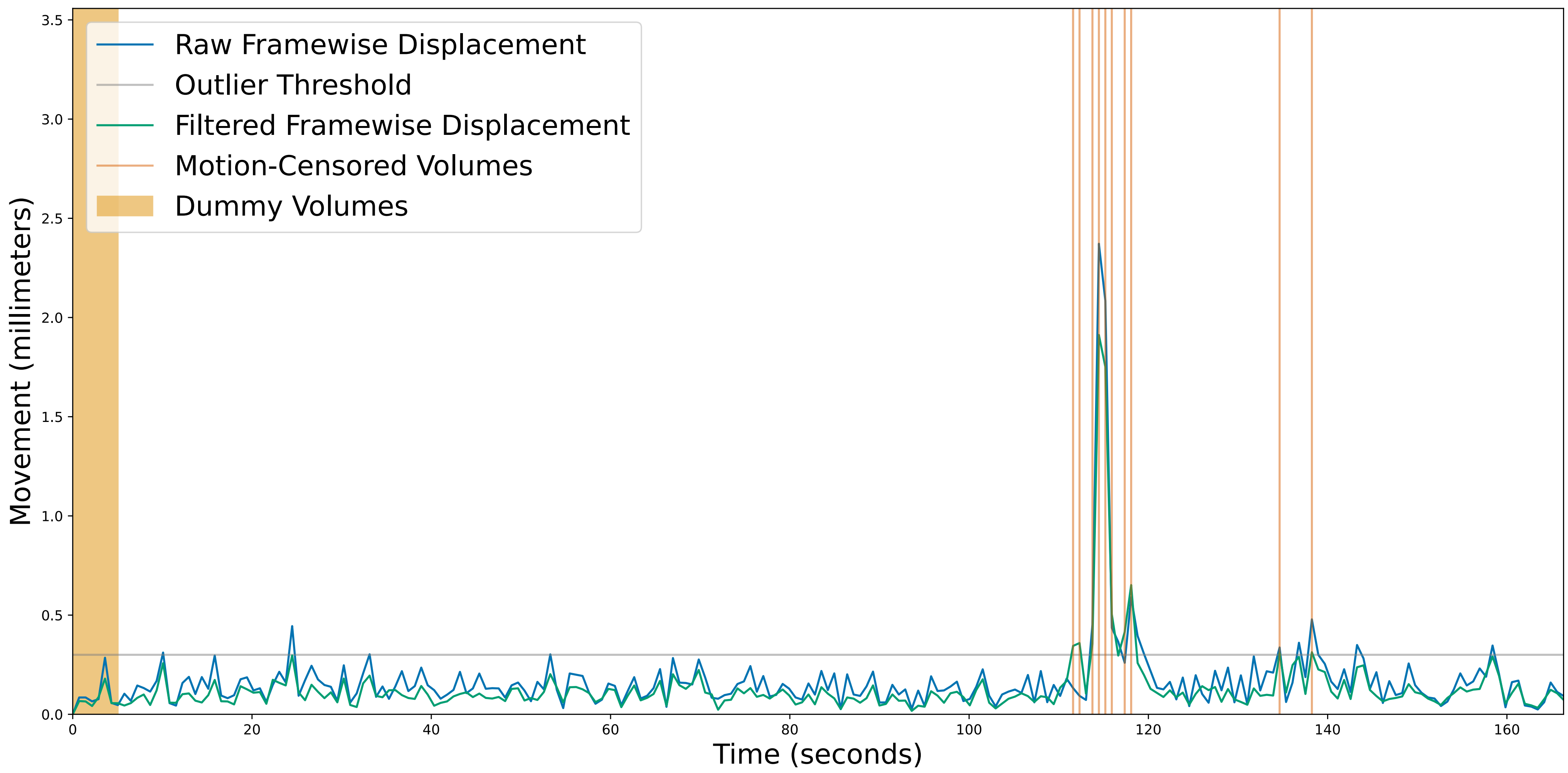


Get figure file: [sub-113922/figures/sub-113922_task-RELATIONAL_dir-LR_space-fsLR_desc-censoring_motion.svg](sub-113922/figures/sub-113922_task-RELATIONAL_dir-LR_space-fsLR_desc-censoring_motion.svg)

#### Design Matrix for Confound Regression

The "design matrix" represents the confounds that are used to denoise the BOLD data.

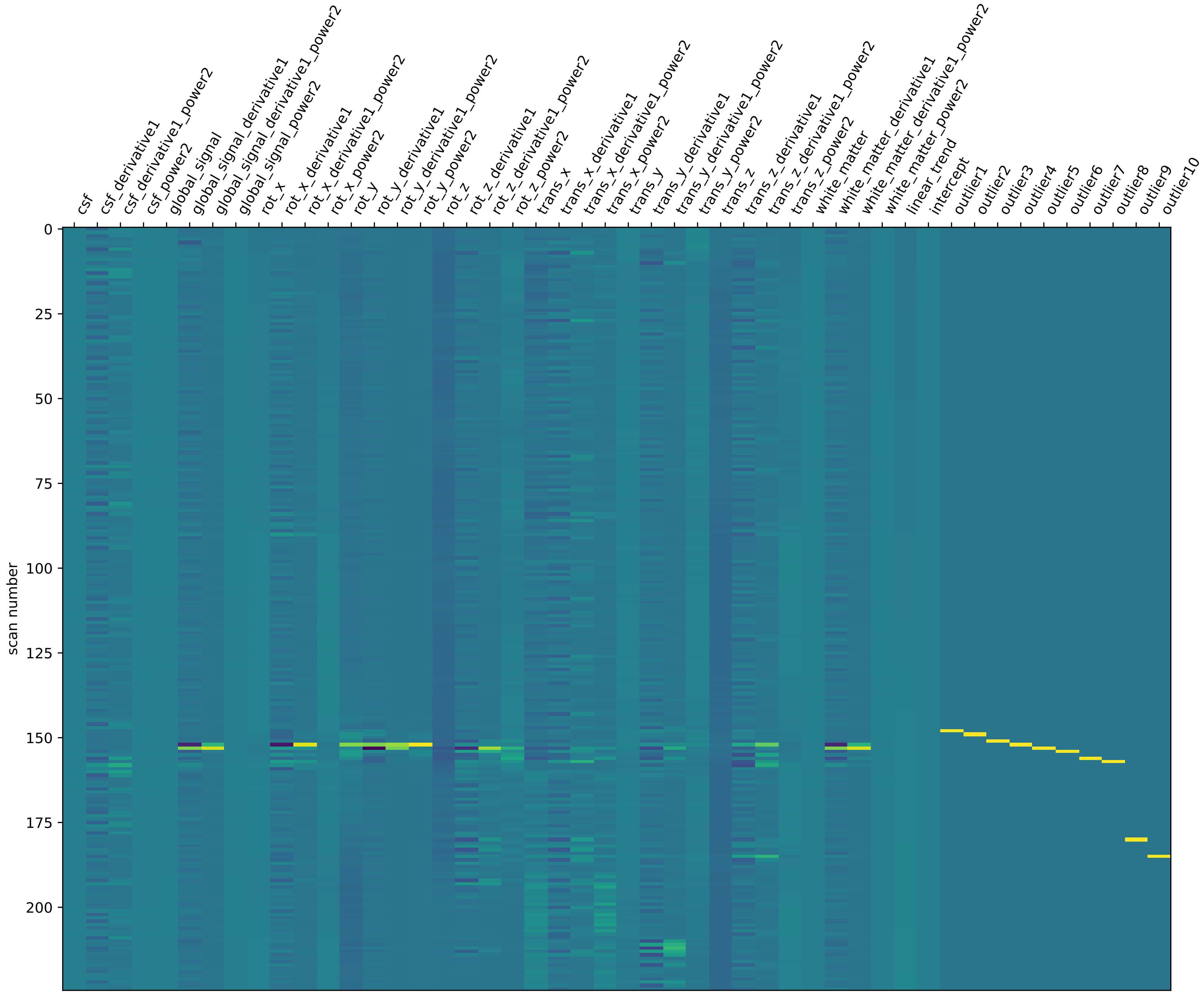


Get figure file: [sub-113922/figures/sub-113922_task-RELATIONAL_dir-LR_design.svg](sub-113922/figures/sub-113922_task-RELATIONAL_dir-LR_design.svg)

### Carpet Plot After Postprocessing

FD and DVARS are two measures of in-scanner motion. This plot shows standardized FD, DVARS, and then a carpet plot for the time series of each voxel/vertex's time series of activity.

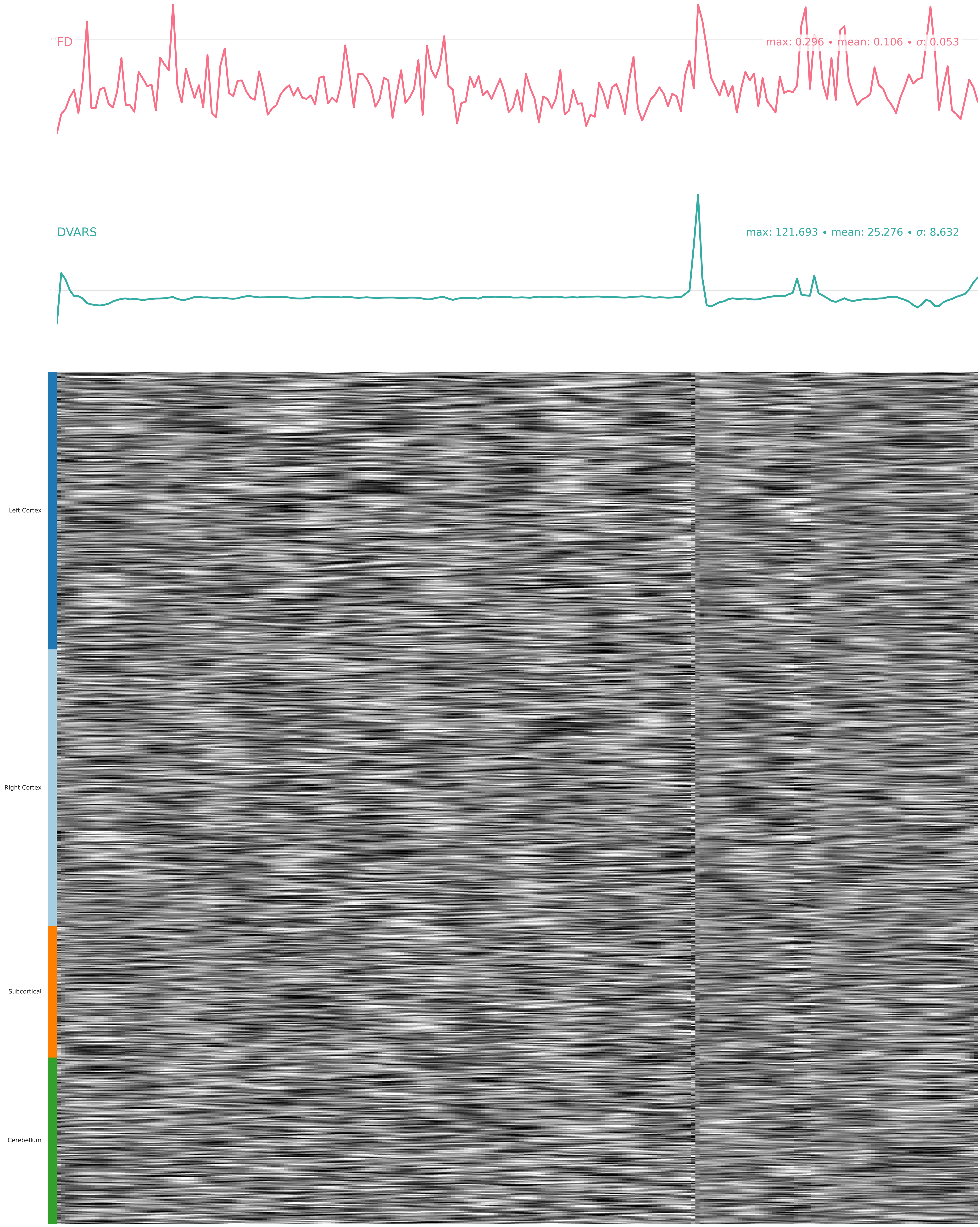


Get figure file: sub-113922/figures/sub-113922_task-RELATIONAL_dir-LR_space-fsLR_desc-postprocessing_bold.svg

#### Correlation Heatmaps from Four Atlases

This plot shows heatmaps from ROI-to-ROI correlations from four atlases.

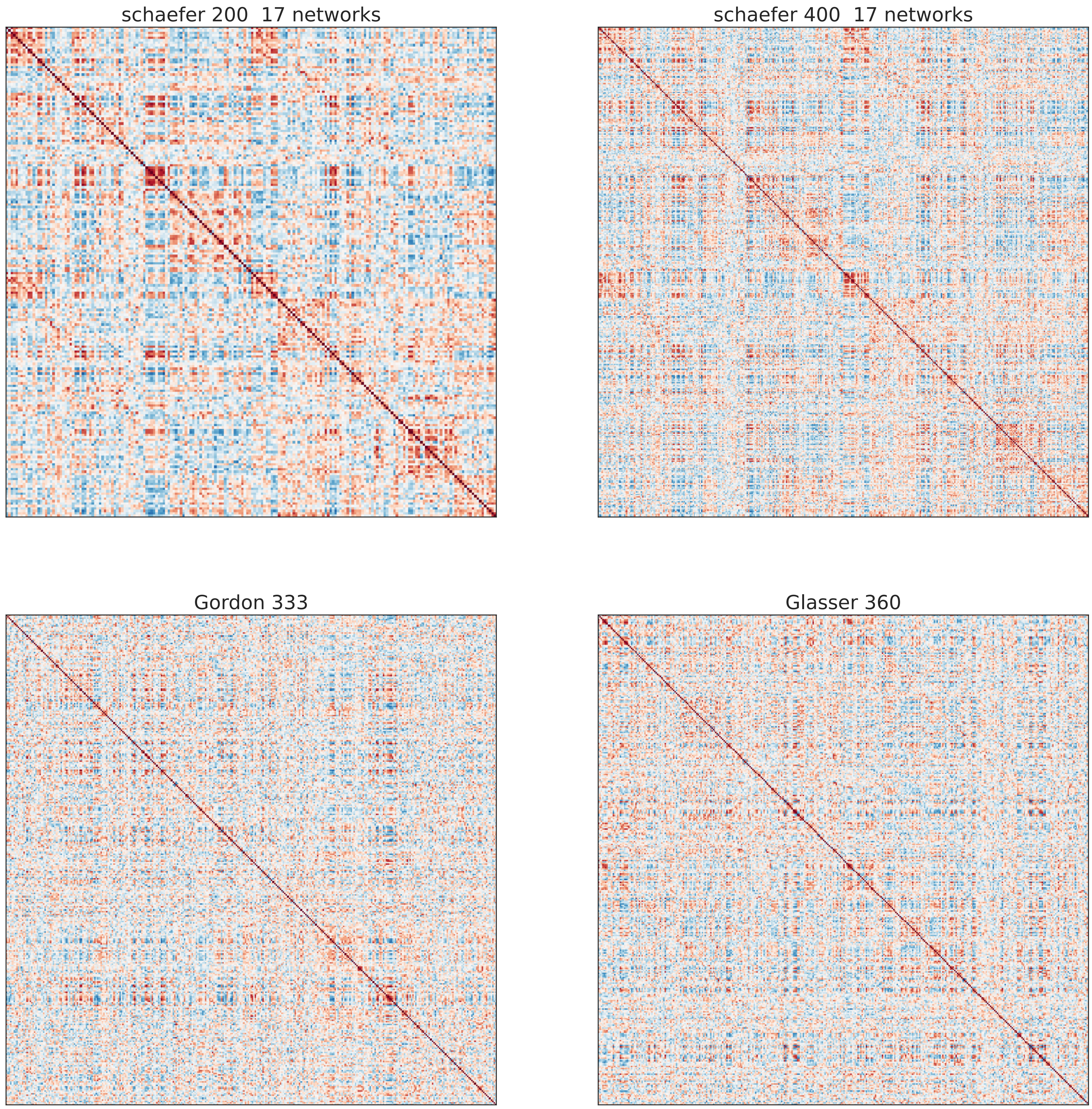


Get figure file: [sub-113922/figures/sub-113922_task-RELATIONAL_dir-LR_space-fsLR_desc-connectivityplot_bold.svg]

### Reports for: task RELATIONAL, direction RL.

#### Summary

- BOLD volume space: fsLR
- Repetition Time (TR): 0.72
- Mean Framewise Displacement: 0.114
- Mean Relative RMS Motion: 0.6506
- Max Relative RMS Motion: 0.9029
- DVARS Before and After Processing : 476.7603, 24.7755
- Correlation between DVARS and FD Before and After Processing : 0.2801, 0.0977
- Number of Volumes Censored : 4

### Alignment of functional and anatomical MRI data (surface driven)

bbregister was used to coregister functional and anatomical MRI data.

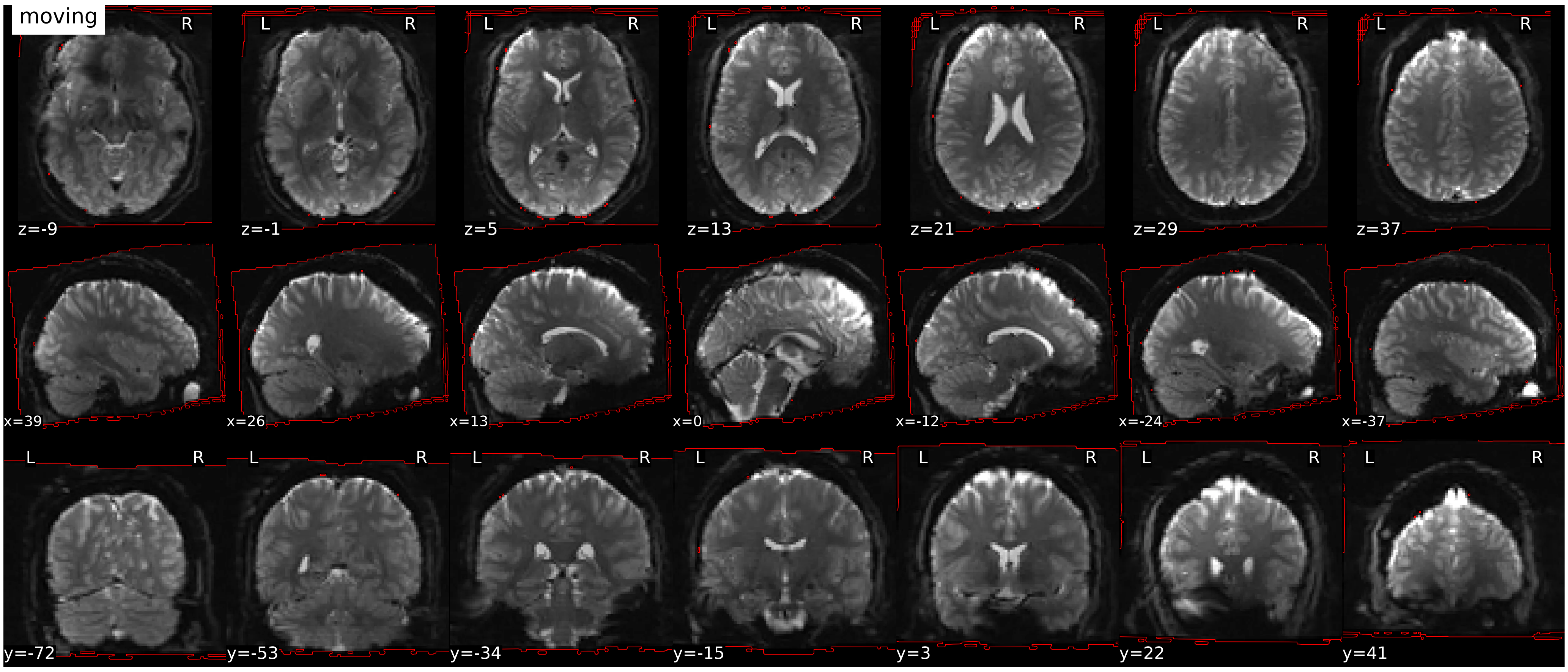


Get figure file: sub-113922/figures/sub-113922_task-RELATIONAL_dir-RL_space-MNI152NLin6Asym_desc-bbregister_bold.svg

### Carpet Plot Before Postprocessing

FD and DVARS are two measures of in-scanner motion. This plot shows standardized FD, DVARS, and then a carpet plot for the time series of each voxel/vertex's time series of activity.

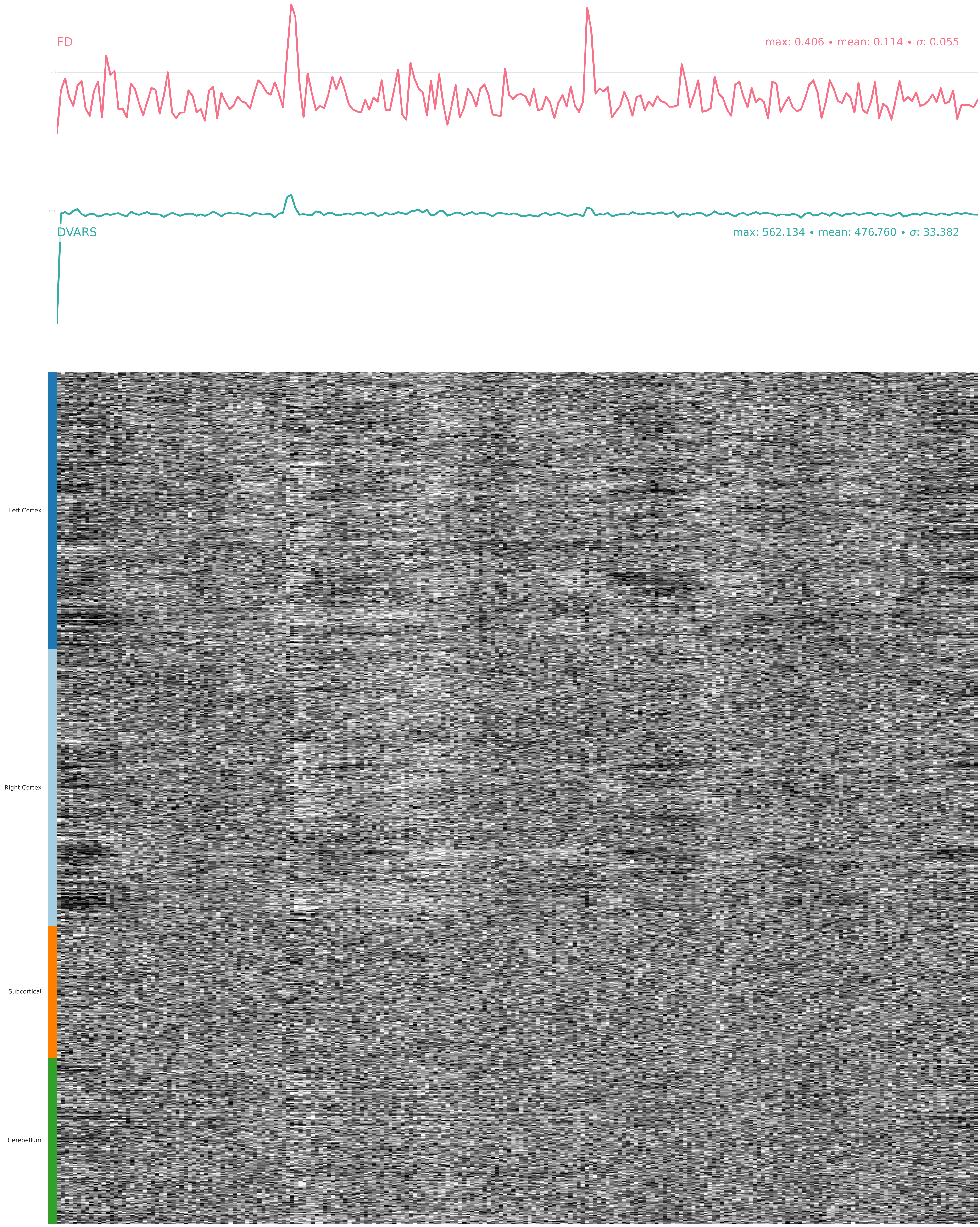


Get figure file: sub-113922/figures/sub-113922_task-RELATIONAL_dir-RL_space-fsLR_desc-preprocessing_bold.svg

#### Framewise Displacement and Censored Volumes

Framewise displacement (FD) is used to flag high-motion volumes, which are then censored as part of the denoising procedure. If motion filtering is requested, then the six translation and rotation motion parameters are filtered to remove respiratory effects before FD is calculated and outlier volumes are identified.

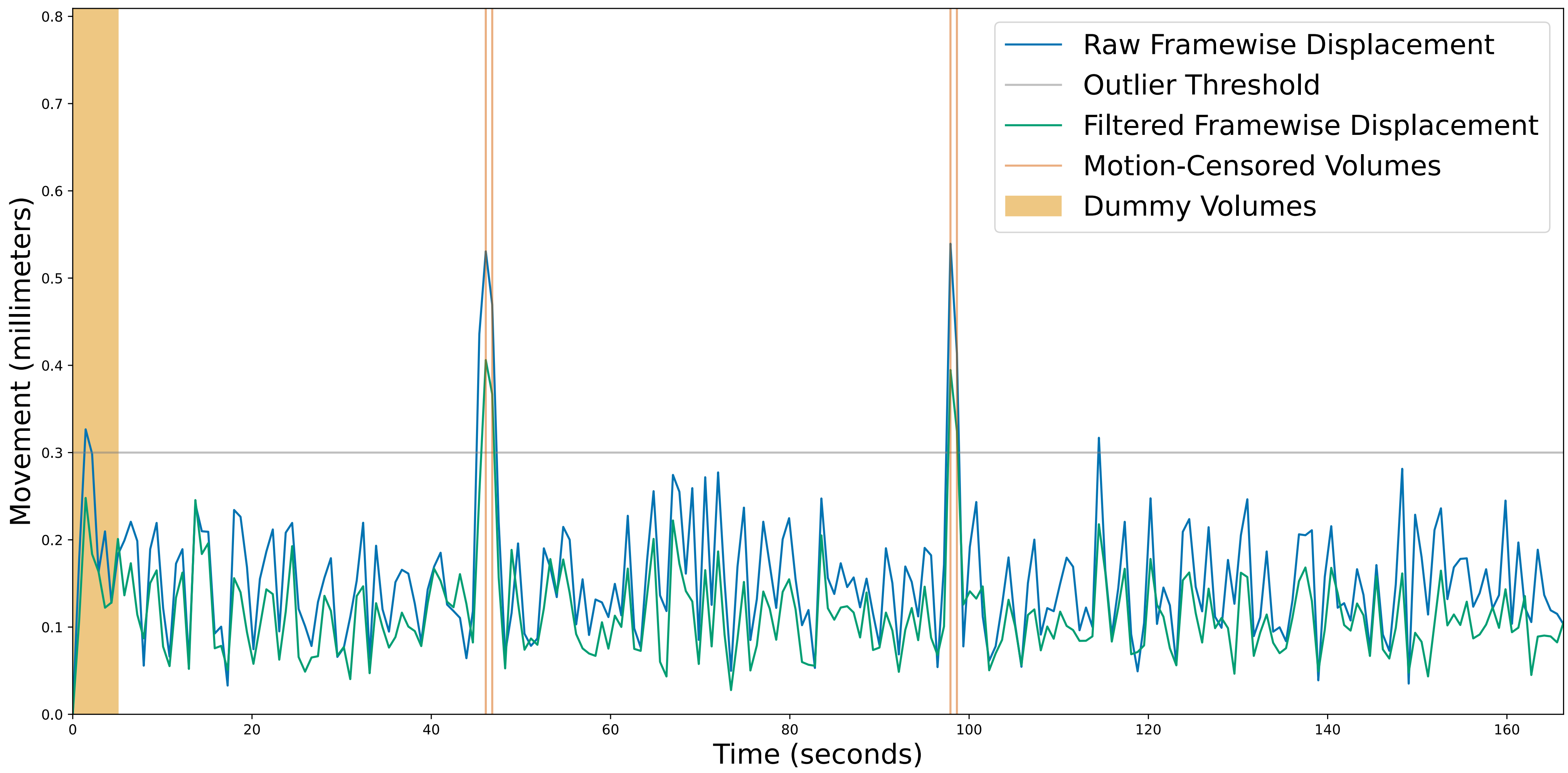


Get figure file: sub-113922/figures/sub-113922_task-RELATIONAL_dir-RL_space-fsLR_desc-censoring_motion.svg

#### Design Matrix for Confound Regression

The "design matrix" represents the confounds that are used to denoise the BOLD data.

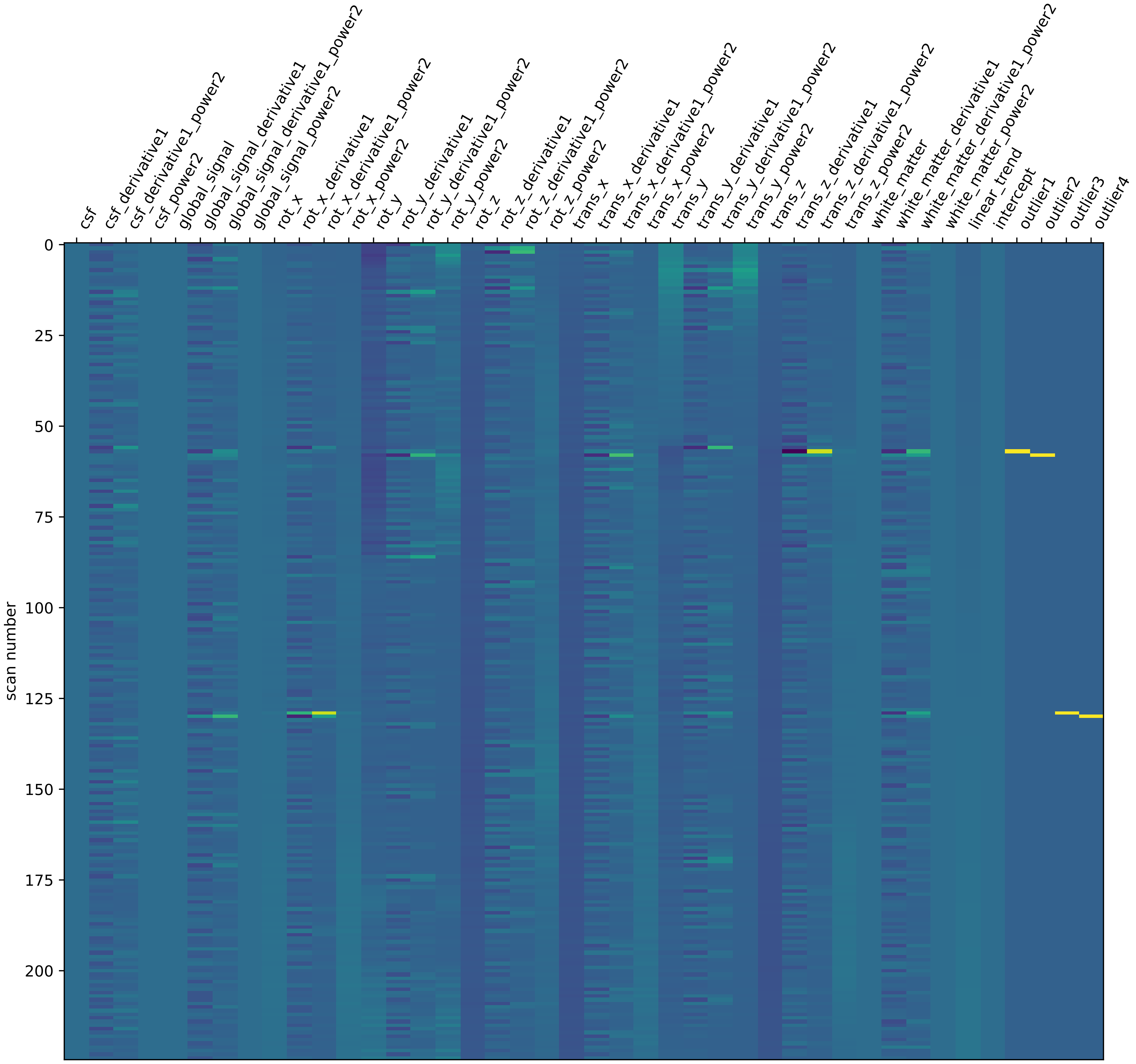


Get figure file: sub-113922/figures/sub-113922_task-RELATIONAL_dir-RL_design.svg

#### Carpet Plot After Postprocessing

FD and DVARS are two measures of in-scanner motion. This plot shows standardized FD, DVARS, and then a carpet plot for the time series of each voxel/vertex's time series of activity.

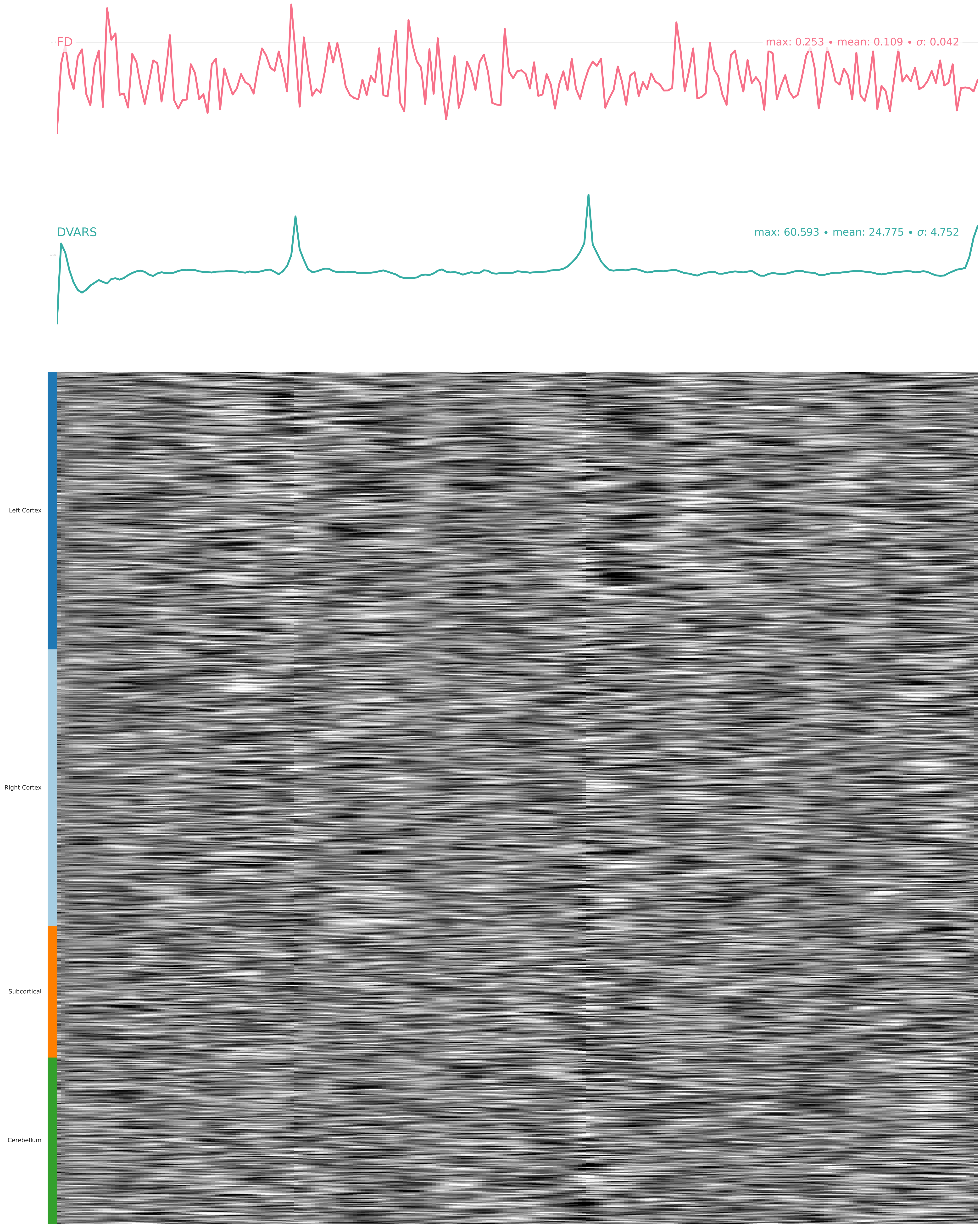


Get figure file: [sub-113922/figures/sub-113922_task-RELATIONAL_dir-RL_space-fsLR_desc-postprocessing_bold.svg](sub-113922/figures/sub-113922_task-RELATIONAL_dir-RL_space-fsLR_desc-postprocessing_bold.svg)

#### Correlation Heatmaps from Four Atlases

This plot shows heatmaps from ROI-to-ROI correlations from four atlases.

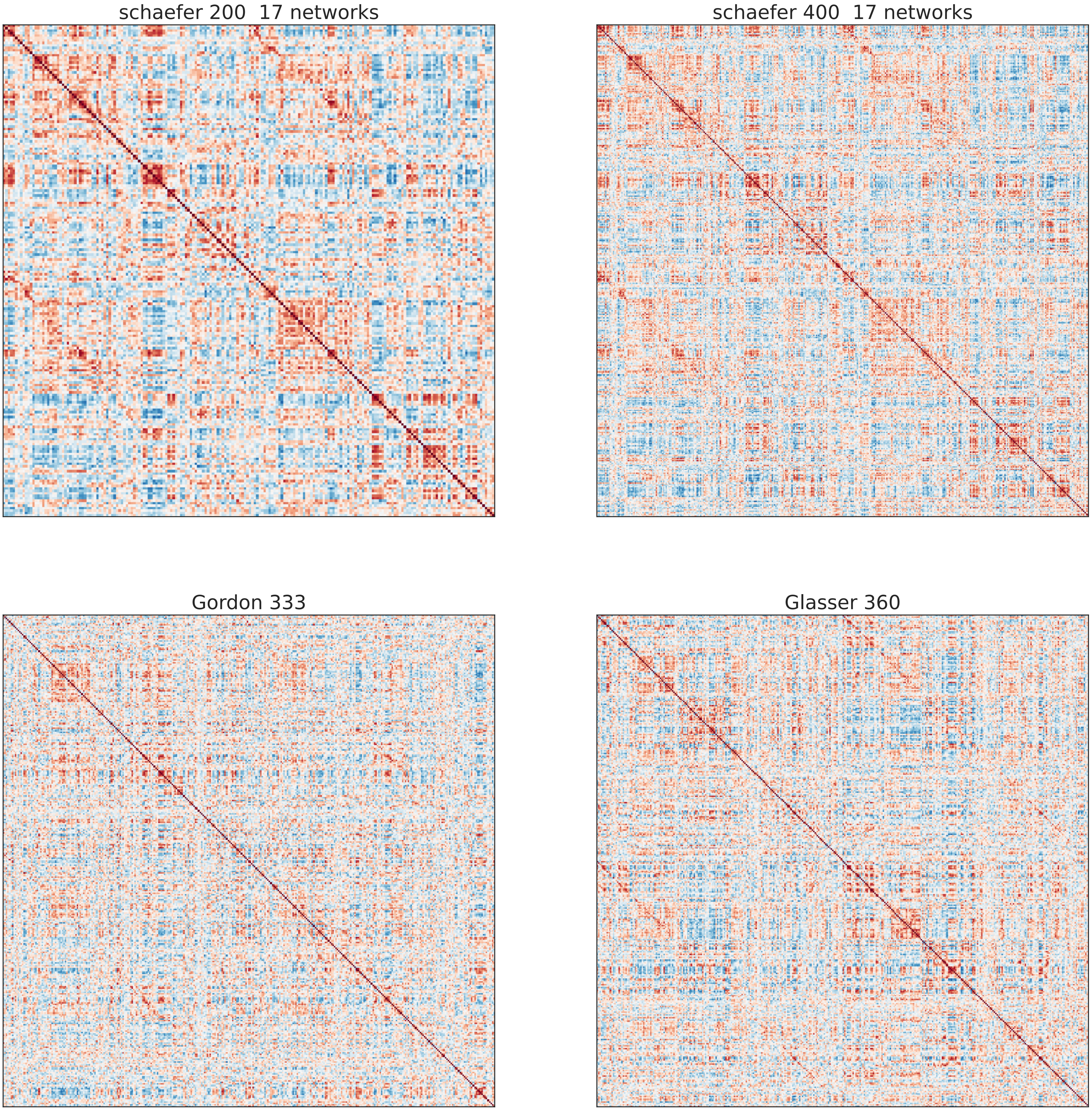


Get figure file: sub-113922/figures/sub-113922_task-RELATIONAL_dir-RL_space-fsLR_desc-connectivityplot_bold.svg

### Reports for: task REST1, direction LR.

#### Summary

- BOLD volume space: fsLR
- Repetition Time (TR): 0.72
- Mean Framewise Displacement: 0.1645
- Mean Relative RMS Motion: 1.8504
- Max Relative RMS Motion: 3.0288
- DVARS Before and After Processing : 482.219, 34.2559
- Correlation between DVARS and FD Before and After Processing : 0.3745, 0.3174
- Number of Volumes Censored : 133

#### Alignment of functional and anatomical MRI data (surface driven)

bbregister was used to coregister functional and anatomical MRI data.

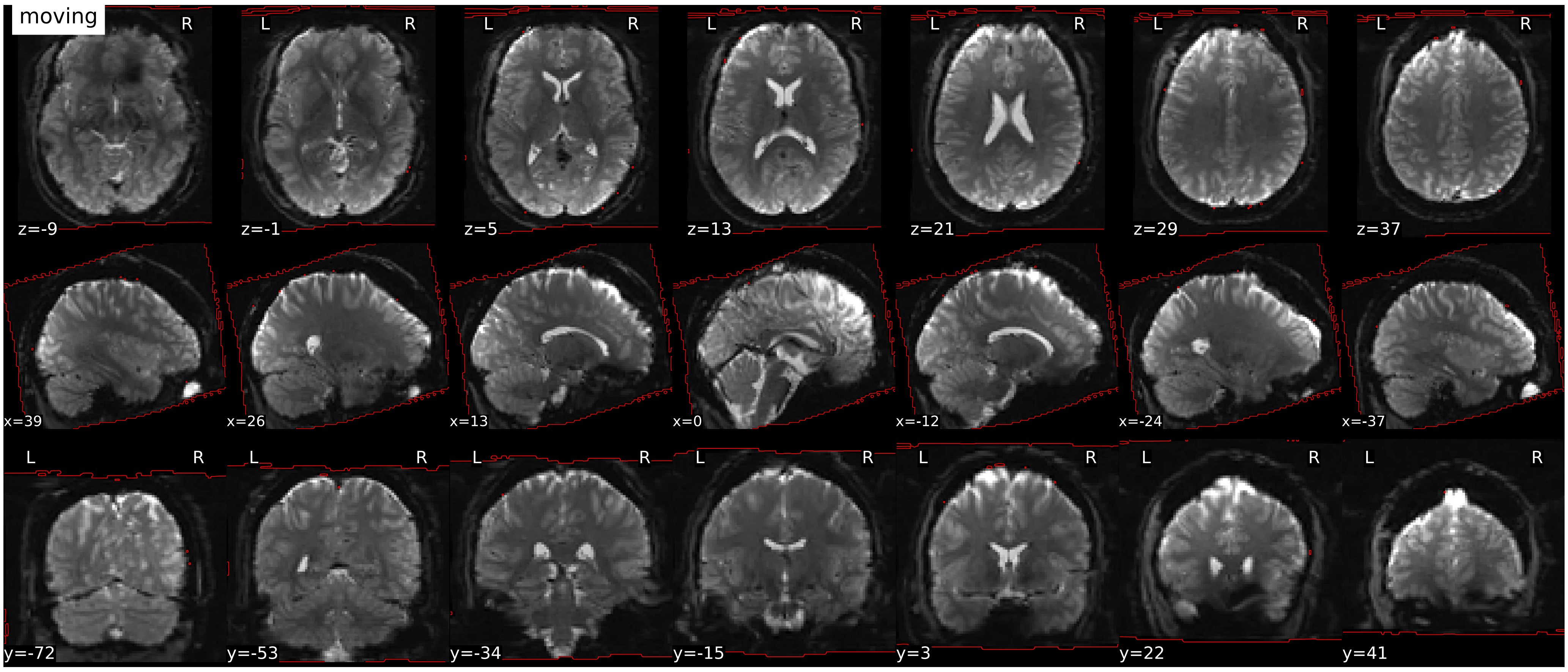


Get figure file: sub-113922/figures/sub-113922_task-REST1_dir-LR_space-MNI152NLin6Asym_desc-bbregister_bold.svg

#### Carpet Plot Before Postprocessing

FD and DVARS are two measures of in-scanner motion. This plot shows standardized FD, DVARS, and then a carpet plot for the time series of each voxel/vertex's time series of activity.

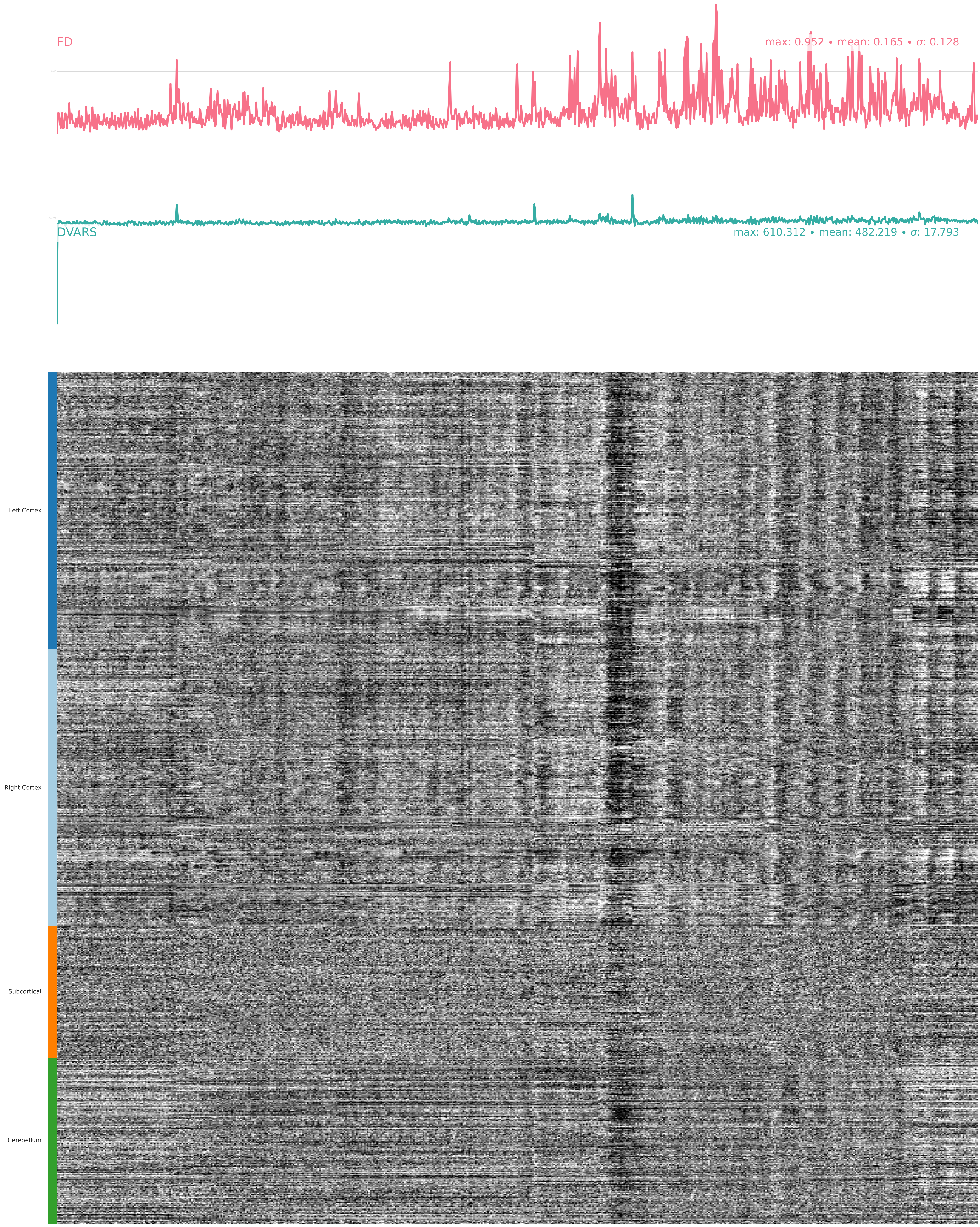


Get figure file: sub-113922/figures/sub-113922_task-REST1_dir-LR_space-fsLR_desc-preprocessing_bold.svg

#### Framewise Displacement and Censored Volumes

Framewise displacement (FD) is used to flag high-motion volumes, which are then censored as part of the denoising procedure. If motion filtering is requested, then the six translation and rotation motion parameters are filtered to remove respiratory effects before FD is calculated and outlier volumes are identified.

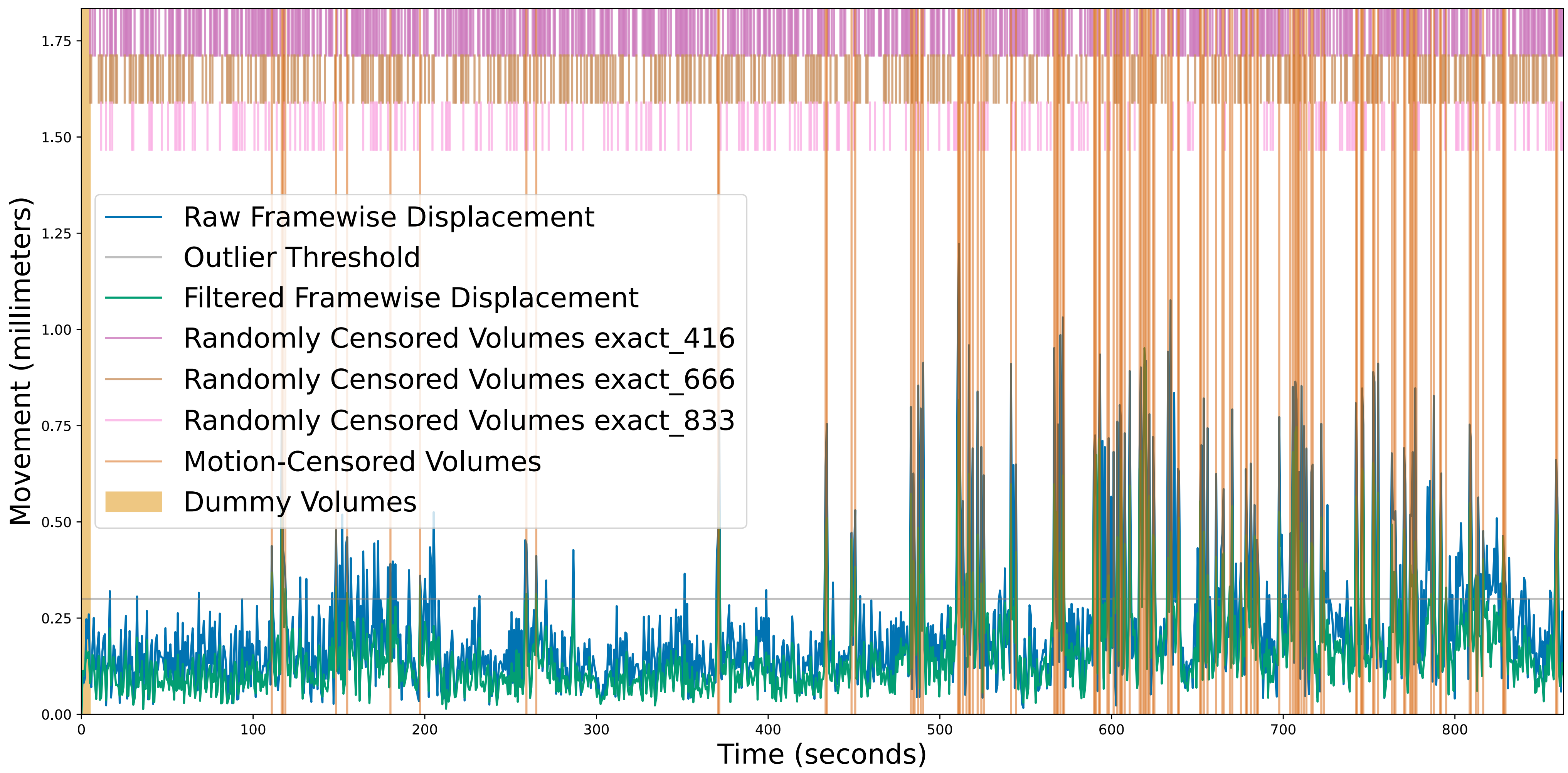


Get figure file: [sub-113922/figures/sub-113922_task-REST1_dir-LR_space-fsLR_desc-censoring_motion.svg](sub-113922/figures/sub-113922_task-REST1_dir-LR_space-fsLR_desc-censoring_motion.svg)

#### Design Matrix for Confound Regression

The "design matrix" represents the confounds that are used to denoise the BOLD data.

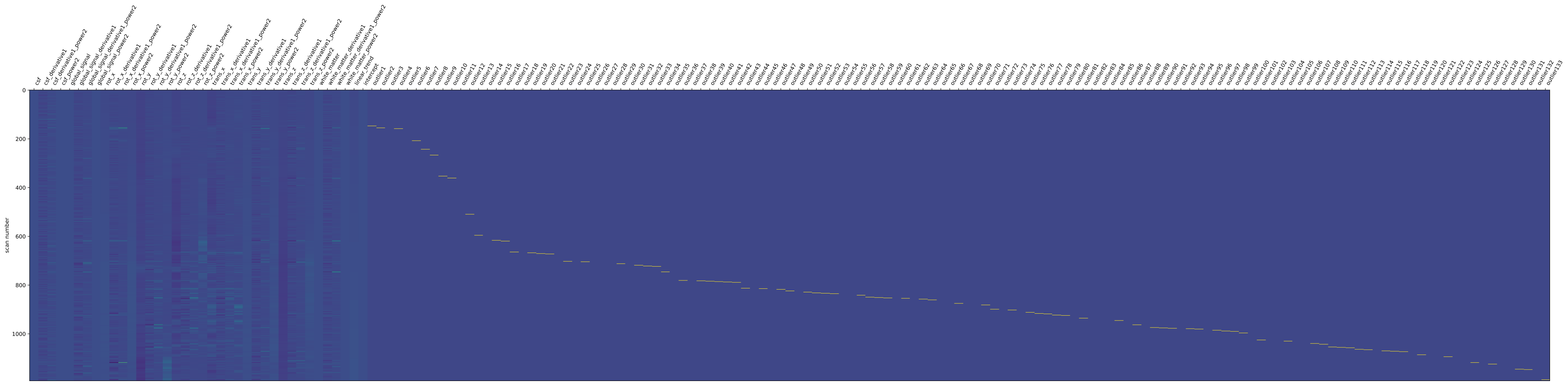

Get figure file: [sub-113922/figures/sub-113922_task-REST1_dir-LR_design.svg](sub-113922/figures/sub-113922_task-REST1_dir-LR_design.svg)

#### Carpet Plot After Postprocessing

FD and DVARS are two measures of in-scanner motion. This plot shows standardized FD, DVARS, and then a carpet plot for the time series of each voxel/vertex's time series of activity.

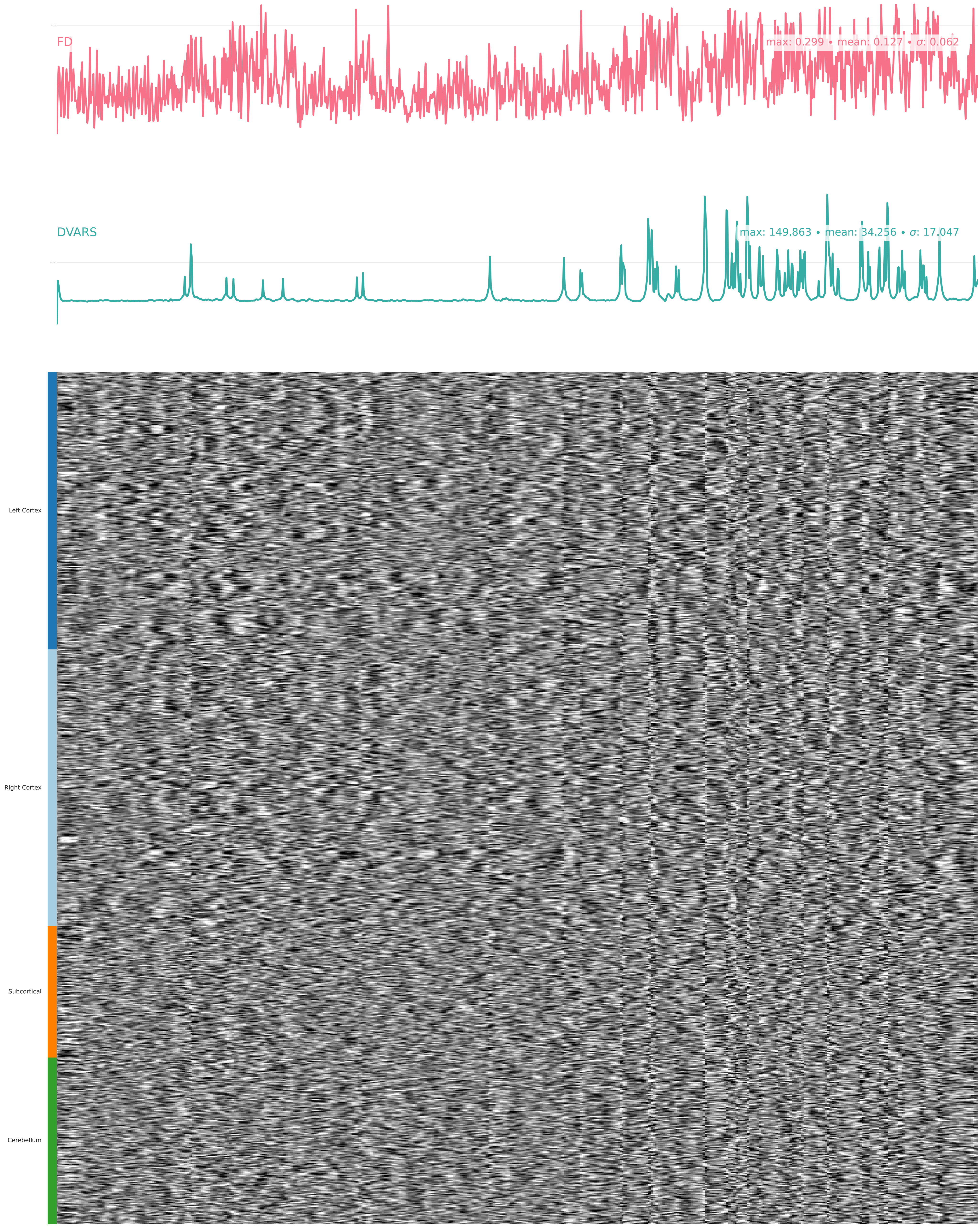


Get figure file: sub-113922/figures/sub-113922_task-REST1_dir-LR_space-fsLR_desc-postprocessing_bold.svg

#### Correlation Heatmaps from Four Atlases

This plot shows heatmaps from ROI-to-ROI correlations from four atlases.

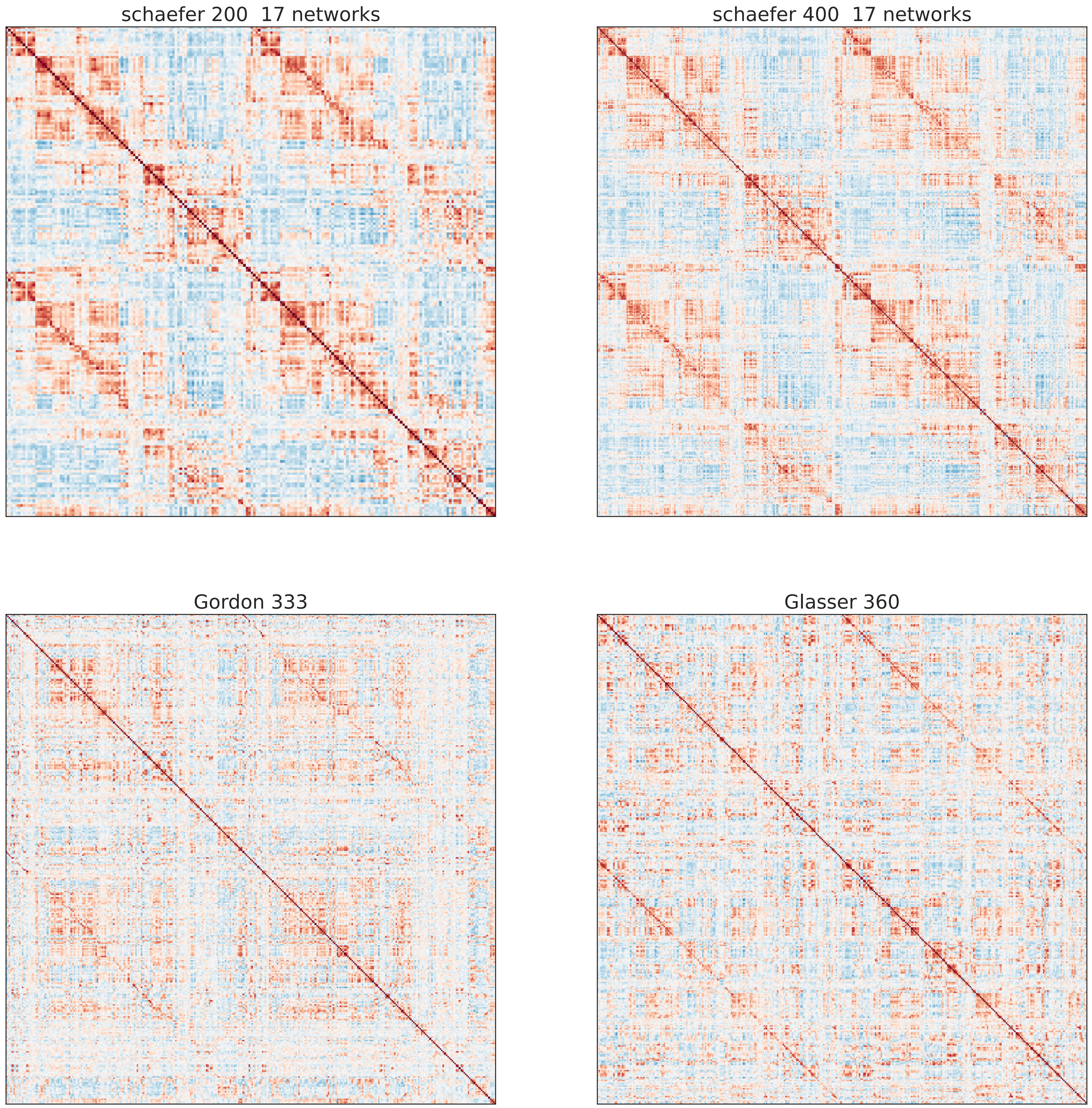


Get figure file: sub-113922/figures/sub-113922_task-REST1_dir-LR_space-fsLR_desc-connectivityplot_bold.svg

### Reports for: task REST1, direction RL.

#### Summary

- BOLD volume space: fsLR
- Repetition Time (TR): 0.72
- Mean Framewise Displacement: 0.1082
- Mean Relative RMS Motion: 0.9401
- Max Relative RMS Motion: 2.0634
- DVARS Before and After Processing : 481.2201, 27.8924
- Correlation between DVARS and FD Before and After Processing : 0.1555, 0.2317
- Number of Volumes Censored : 16

#### Alignment of functional and anatomical MRI data (surface driven)

bbregister was used to coregister functional and anatomical MRI data.

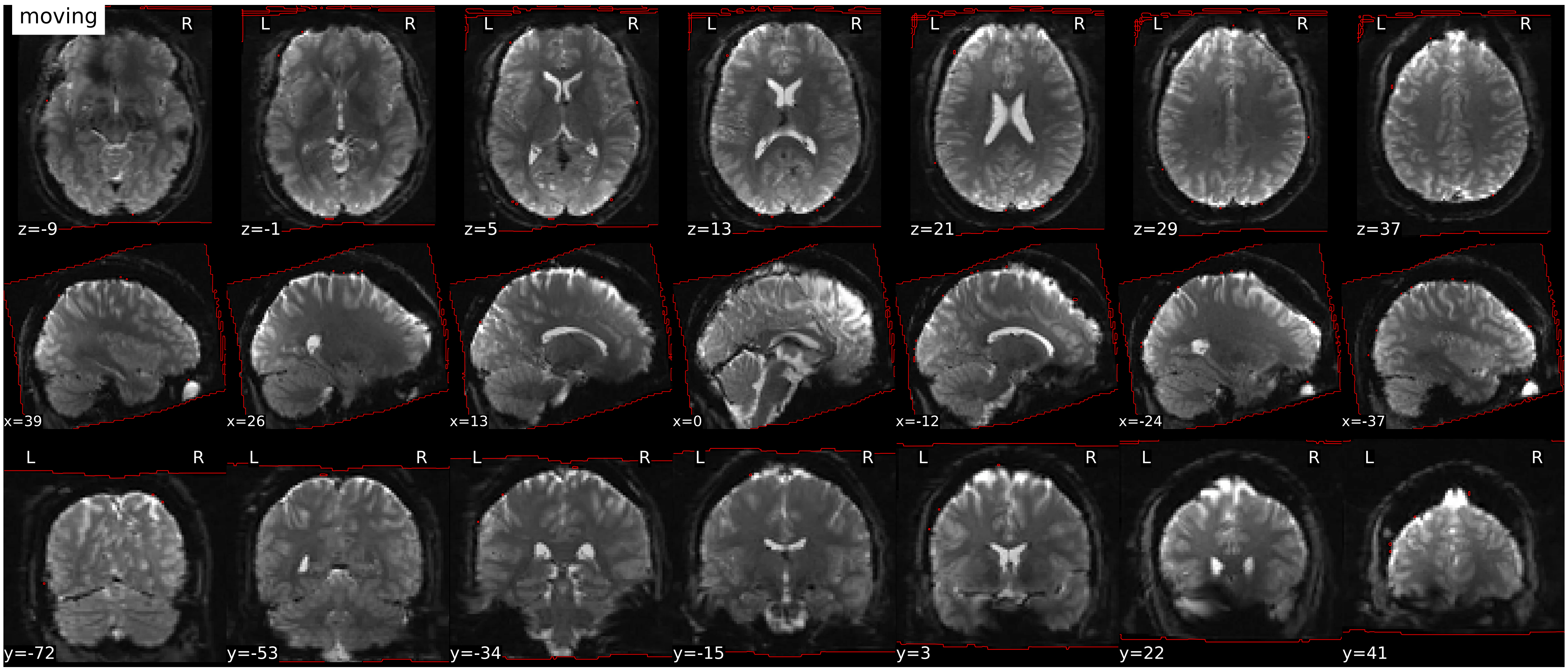


Get figure file: sub-113922/figures/sub-113922_task-REST1_dir-RL_space-MNI152NLin6Asym_desc-bbregister_bold.svg

#### Carpet Plot Before Postprocessing

FD and DVARS are two measures of in-scanner motion. This plot shows standardized FD, DVARS, and then a carpet plot for the time series of each voxel/vertex's time series of activity.

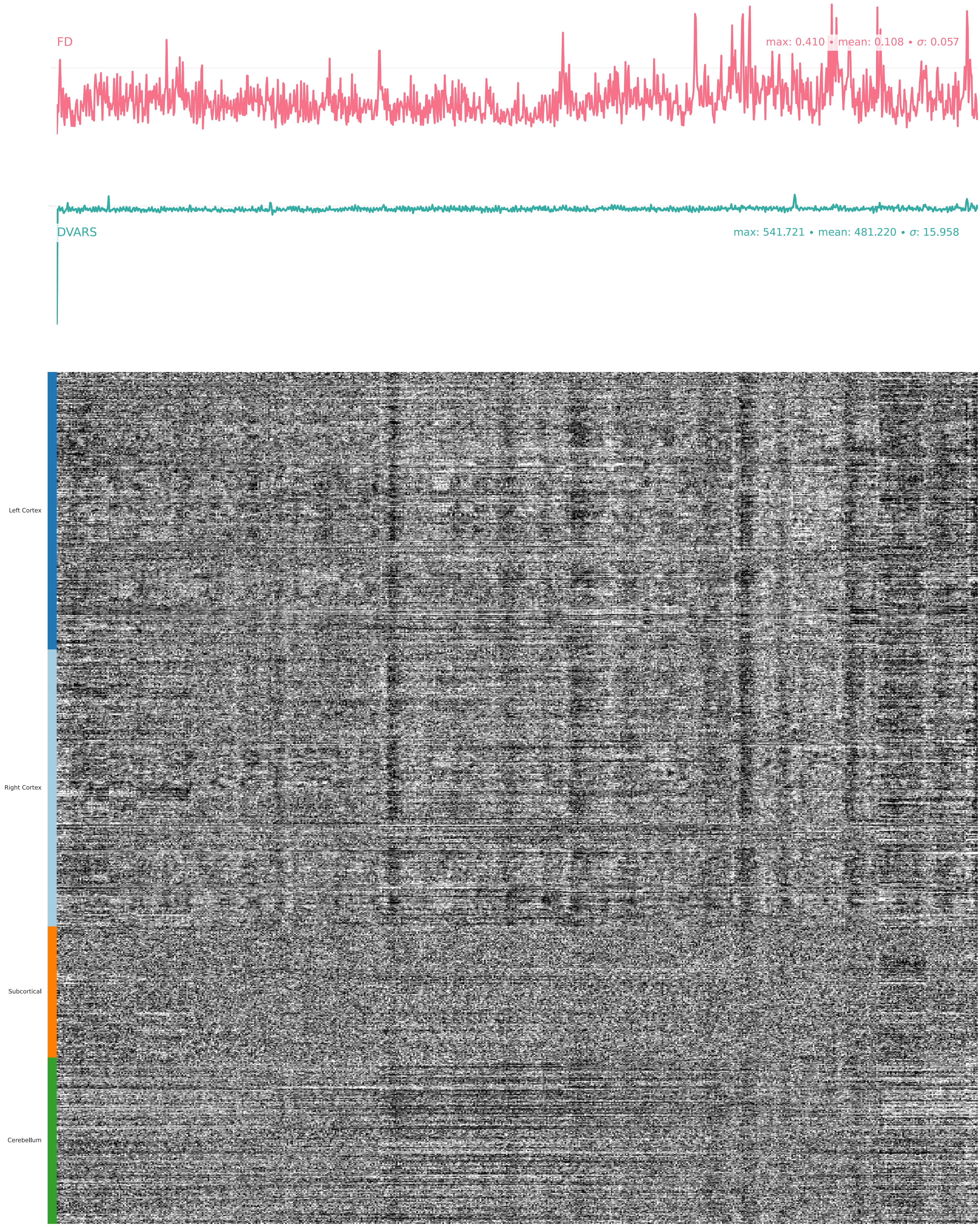


Get figure file: sub-113922/figures/sub-113922_task-REST1_dir-RL_space-fsLR_desc-preprocessing_bold.svg

#### Framewise Displacement and Censored Volumes

Framewise displacement (FD) is used to flag high-motion volumes, which are then censored as part of the denoising procedure. If motion filtering is requested, then the six translation and rotation motion parameters are filtered to remove respiratory effects before FD is calculated and outlier volumes are identified.

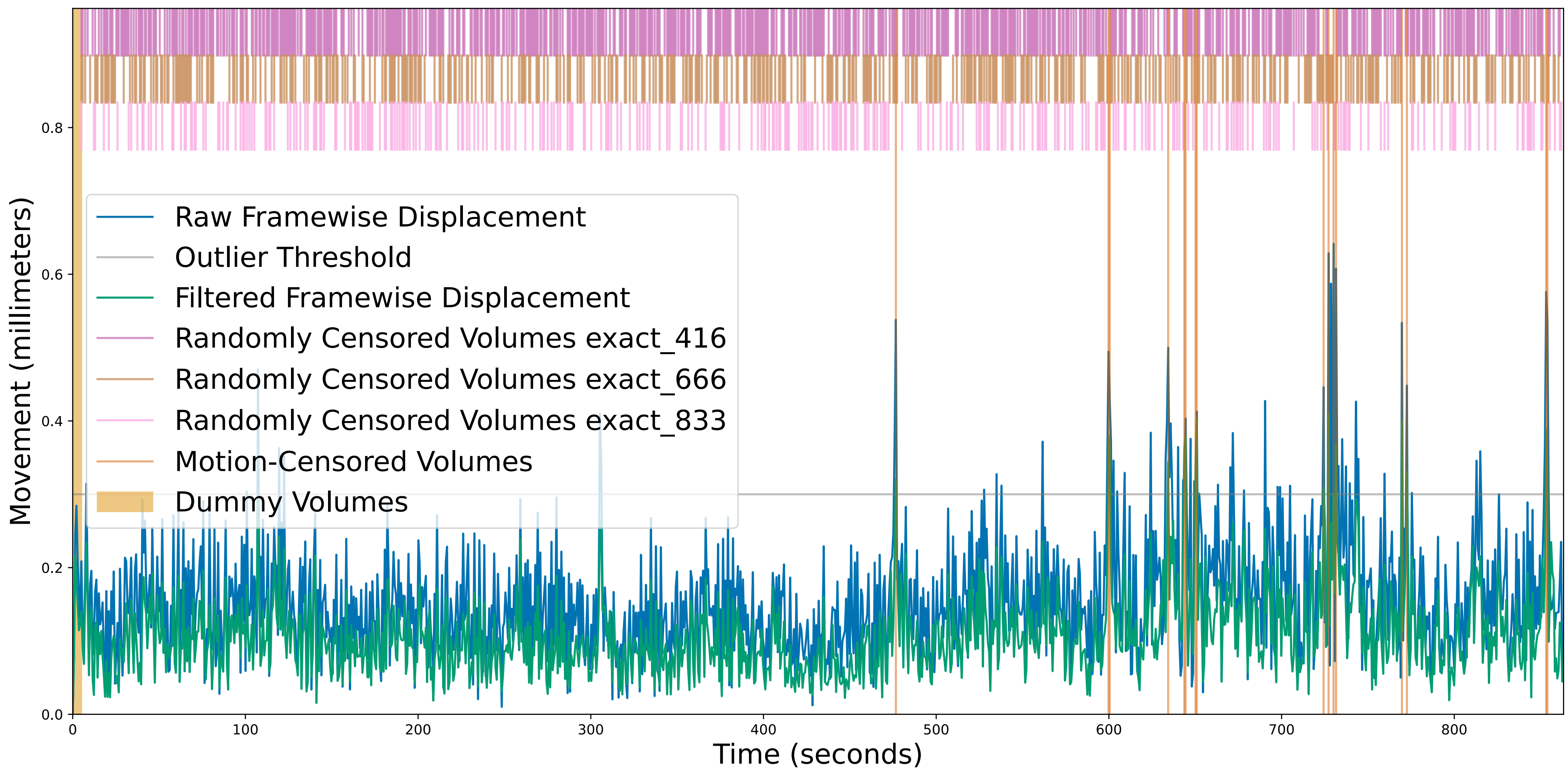


Get figure file: [sub-113922/figures/sub-113922_task-REST1_dir-RL_space-fsLR_desc-censoring_motion.svg](sub-113922/figures/sub-113922_task-REST1_dir-RL_space-fsLR_desc-censoring_motion.svg)

#### Design Matrix for Confound Regression

The "design matrix" represents the confounds that are used to denoise the BOLD data.

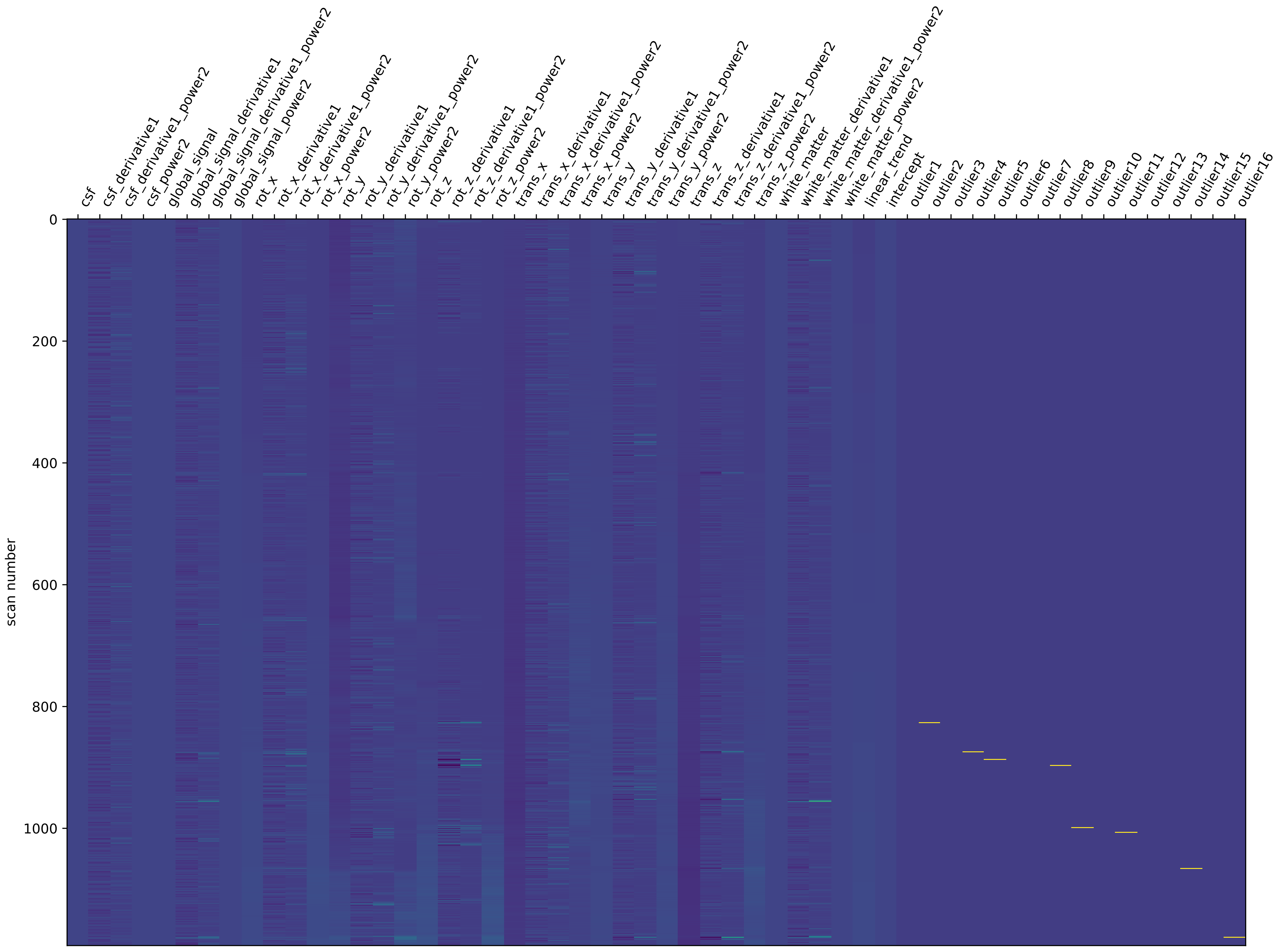


Get figure file: [sub-113922/figures/sub-113922_task-REST1_dir-RL_design.svg](sub-113922/figures/sub-113922_task-REST1_dir-RL_design.svg)

### Carpet Plot After Postprocessing

FD and DVARS are two measures of in-scanner motion. This plot shows standardized FD, DVARS, and then a carpet plot for the time series of each voxel/vertex's time series of activity.

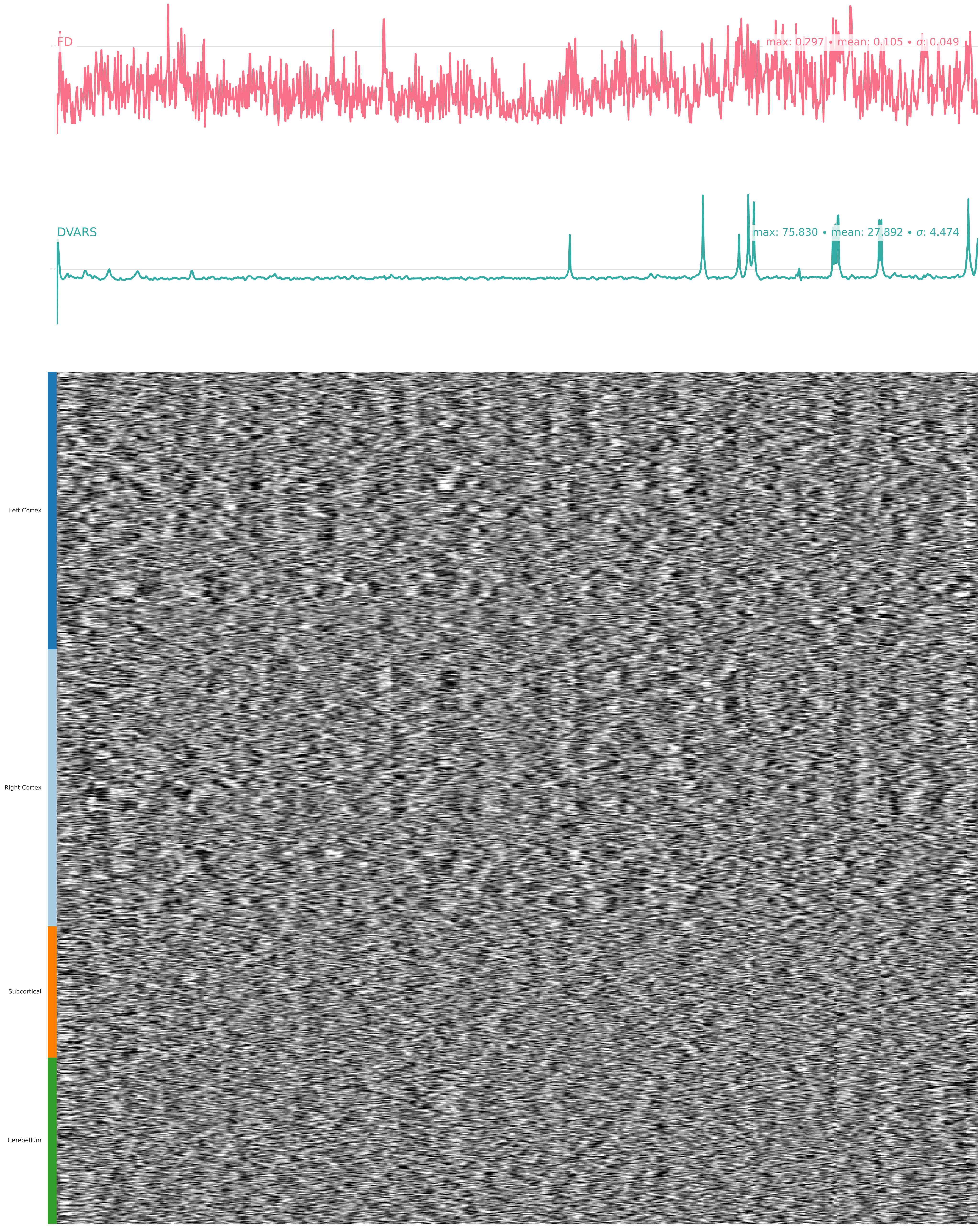


Get figure file: sub-113922/figures/sub-113922_task-REST1_dir-RL_space-fsLR_desc-postprocessing_bold.svg

#### Correlation Heatmaps from Four Atlases

This plot shows heatmaps from ROI-to-ROI correlations from four atlases.

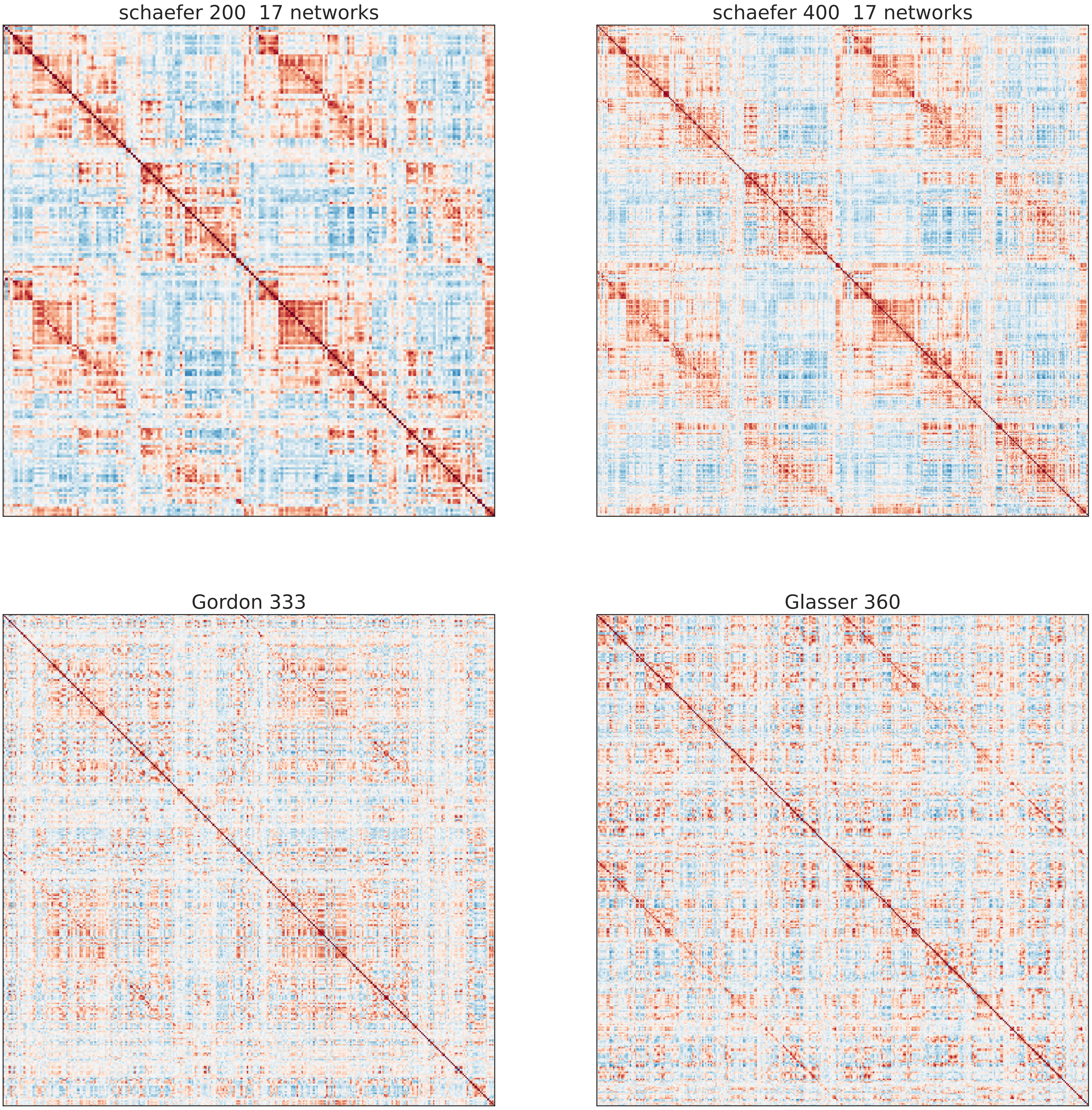


Get figure file: sub-113922/figures/sub-113922_task-REST1_dir-RL_space-fsLR_desc-connectivityplot_bold.svg

### Reports for: task REST2, direction LR.

#### Summary

- BOLD volume space: fsLR
- Repetition Time (TR): 0.72
- Mean Framewise Displacement: 0.1252
- Mean Relative RMS Motion: 0.3947
- Max Relative RMS Motion: 0.6154
- DVARS Before and After Processing : 483.8401, 26.8457
- Correlation between DVARS and FD Before and After Processing : 0.1854, 0.1333
- Number of Volumes Censored : 8

#### Alignment of functional and anatomical MRI data (surface driven)

bbregister was used to coregister functional and anatomical MRI data.

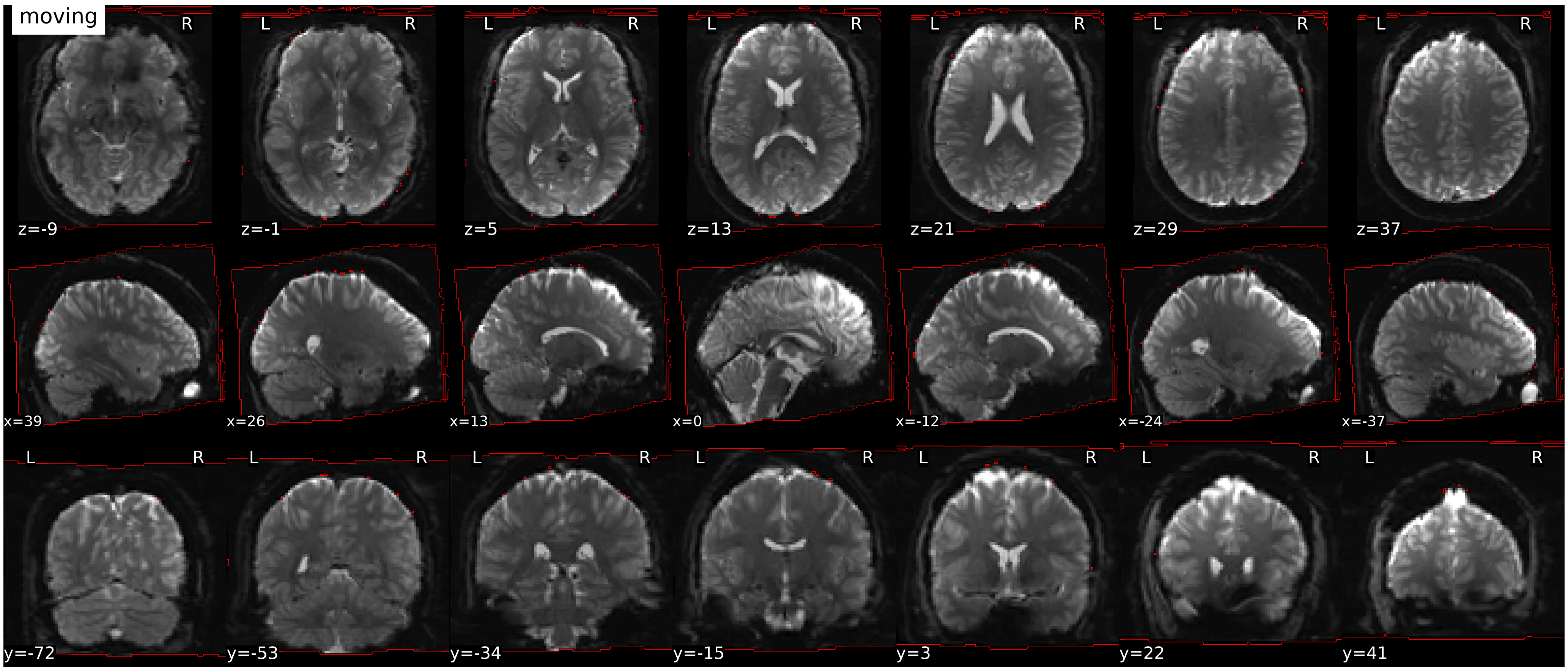


Get figure file: sub-113922/figures/sub-113922_task-REST2_dir-LR_space-MNI152NLin6Asym_desc-bbregister_bold.svg

#### Carpet Plot Before Postprocessing

FD and DVARS are two measures of in-scanner motion. This plot shows standardized FD, DVARS, and then a carpet plot for the time series of each voxel/vertex's time series of activity.

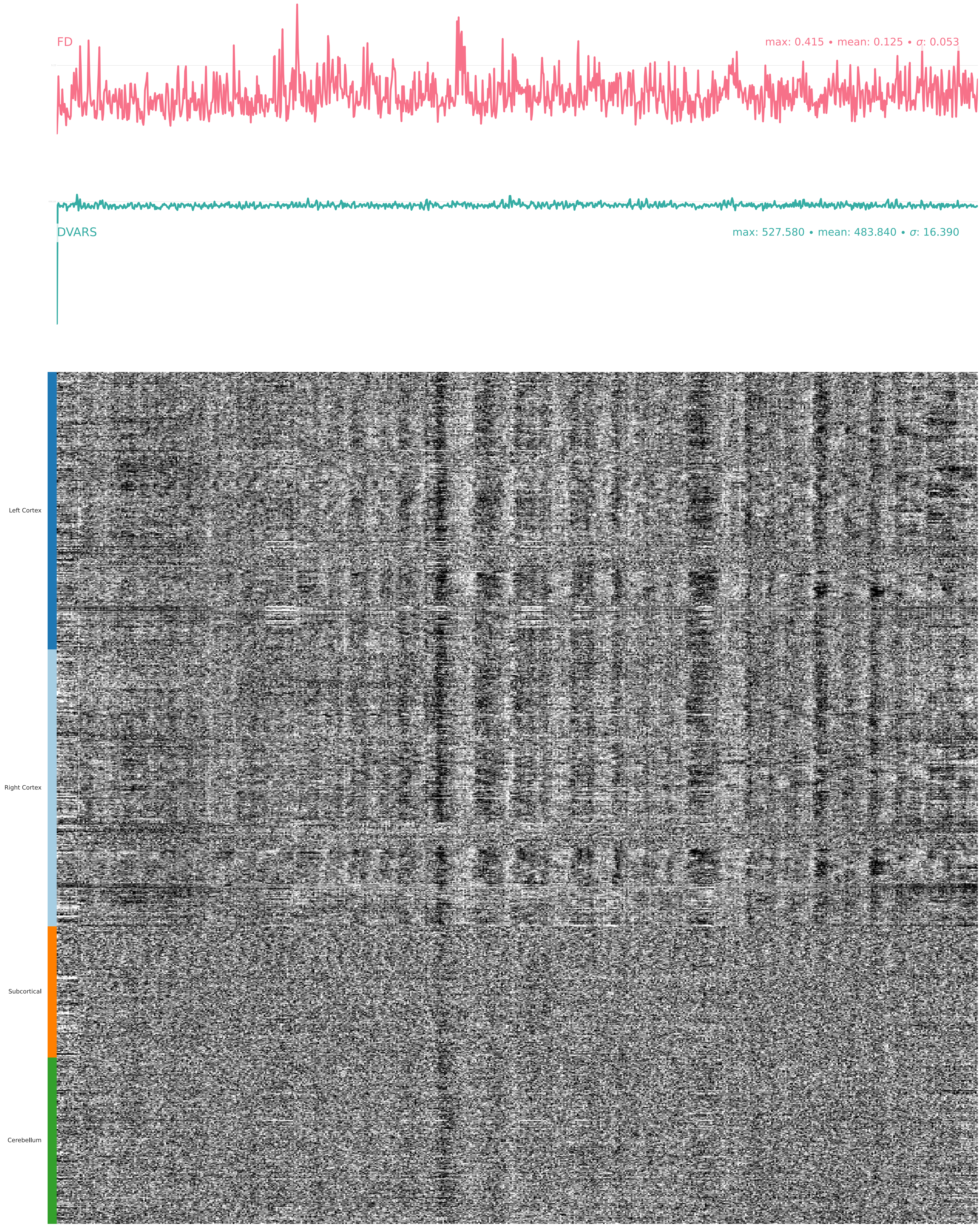


Get figure file: sub-113922/figures/sub-113922_task-REST2_dir-LR_space-fsLR_desc-preprocessing_bold.svg

#### Framewise Displacement and Censored Volumes

Framewise displacement (FD) is used to flag high-motion volumes, which are then censored as part of the denoising procedure. If motion filtering is requested, then the six translation and rotation motion parameters are filtered to remove respiratory effects before FD is calculated and outlier volumes are identified.

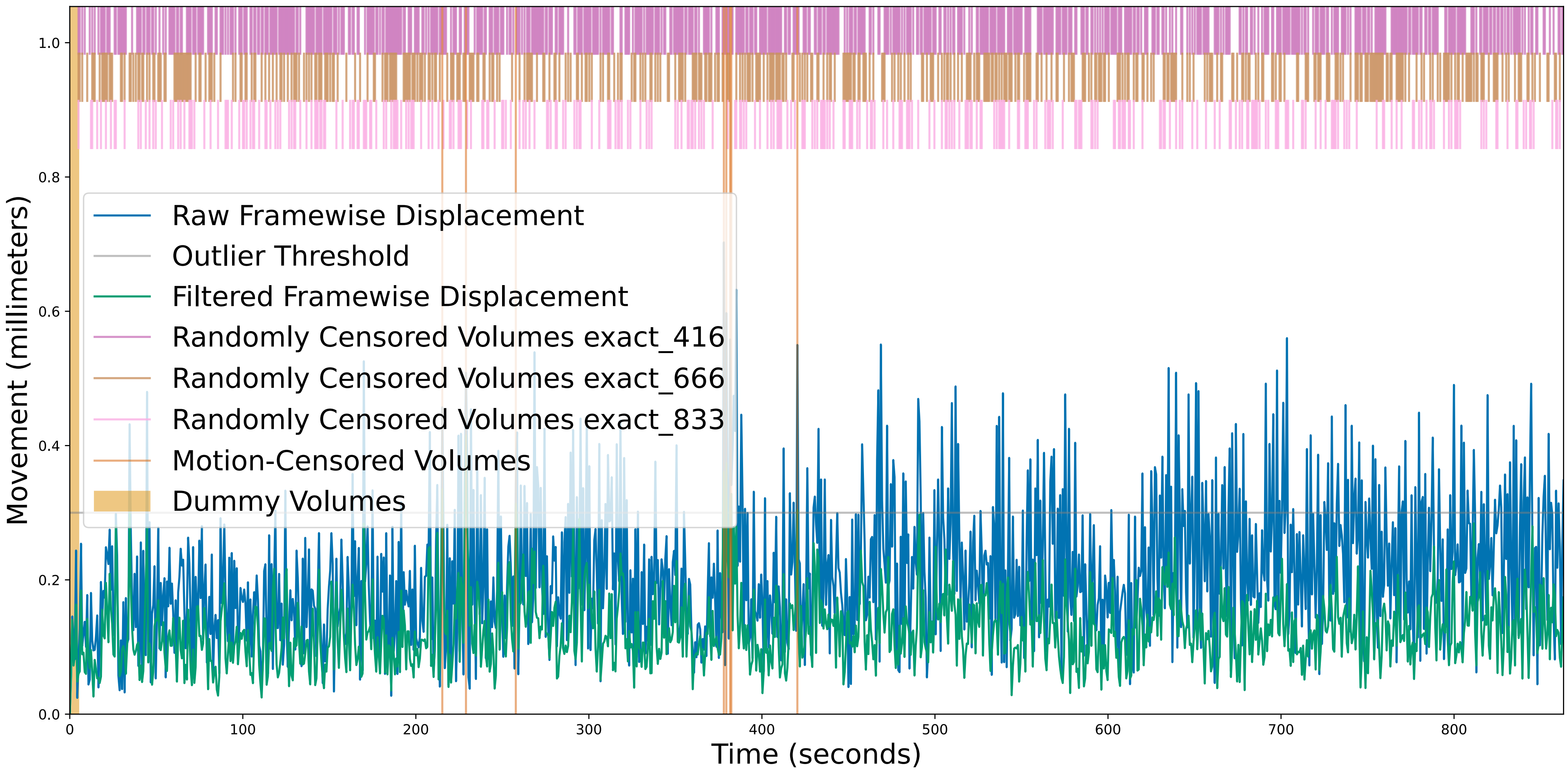


Get figure file: sub-113922/figures/sub-113922_task-REST2_dir-LR_space-fsLR_desc-censoring_motion.svg

#### Design Matrix for Confound Regression

The "design matrix" represents the confounds that are used to denoise the BOLD data.

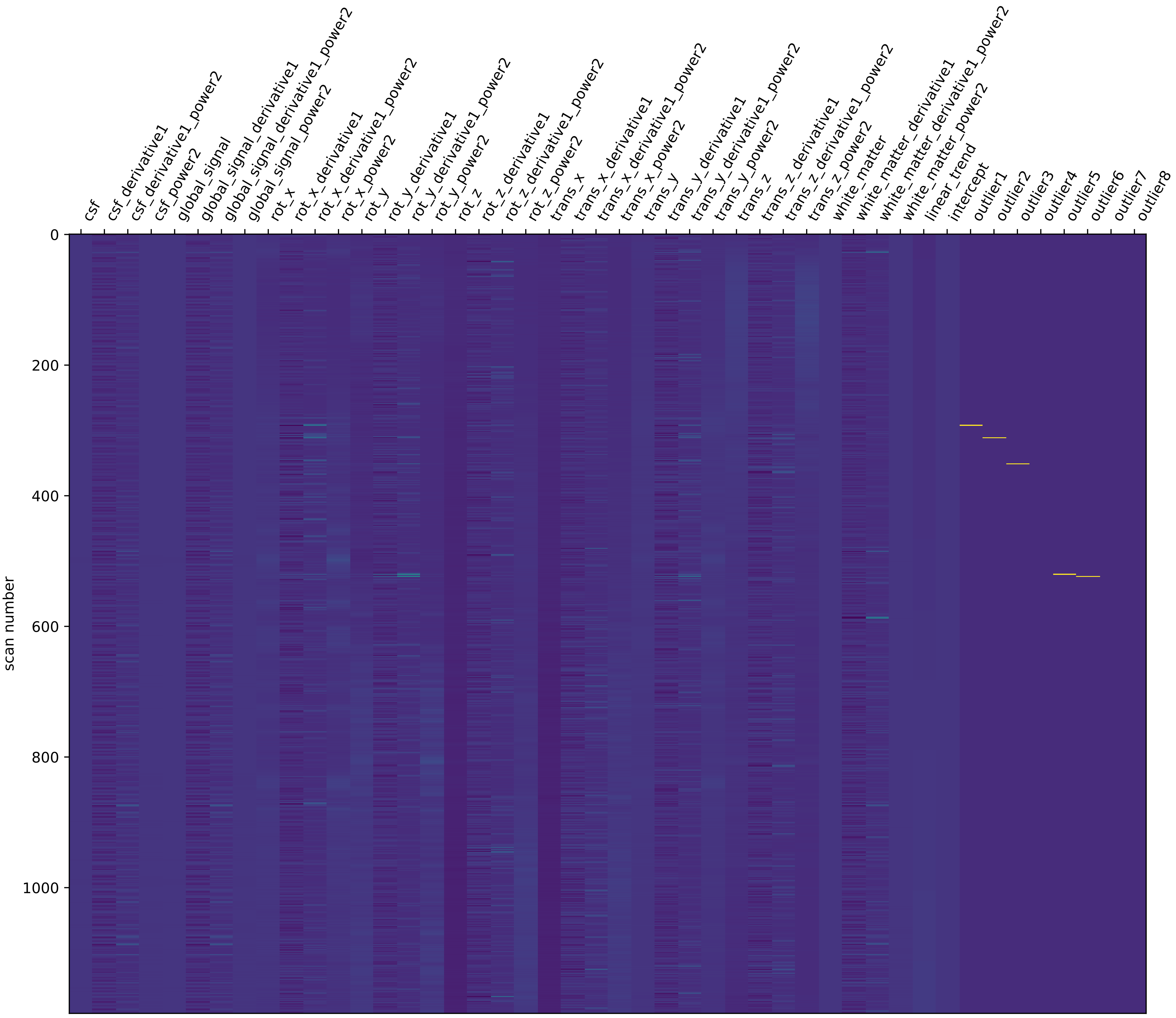


Get figure file: [sub-113922/figures/sub-113922_task-REST2_dir-LR_design.svg](sub-113922/figures/sub-113922_task-REST2_dir-LR_design.svg)

#### Carpet Plot After Postprocessing

FD and DVARS are two measures of in-scanner motion. This plot shows standardized FD, DVARS, and then a carpet plot for the time series of each voxel/vertex's time series of activity.

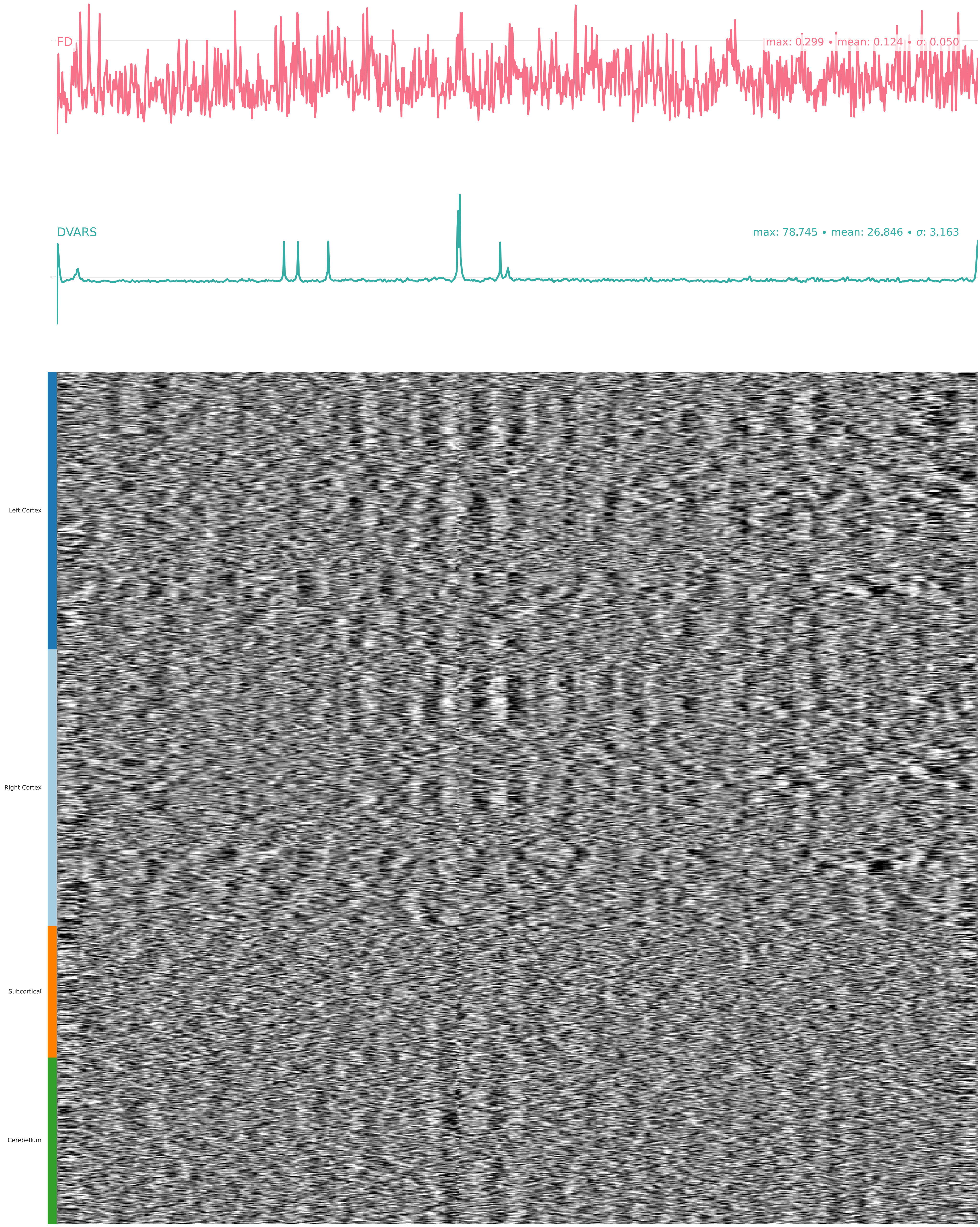


Get figure file: [sub-113922/figures/sub-113922_task-REST2_dir-LR_space-fsLR_desc-postprocessing_bold.svg]

#### Correlation Heatmaps from Four Atlases

This plot shows heatmaps from ROI-to-ROI correlations from four atlases.

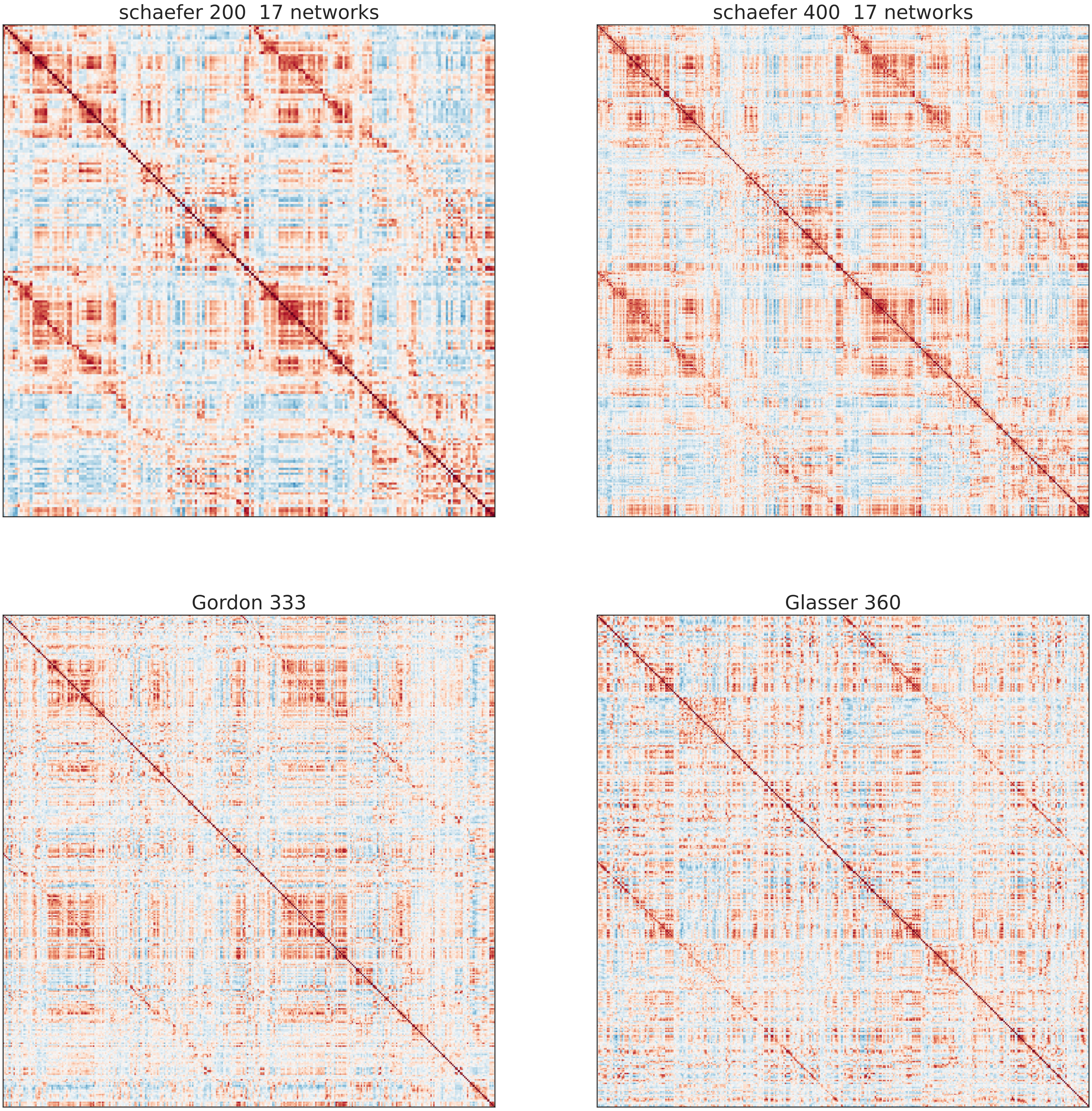


Get figure file: sub-113922/figures/sub-113922_task-REST2_dir-LR_space-fsLR_desc-connectivityplot_bold.svg

### Reports for: task REST2, direction RL.

#### Summary

- BOLD volume space: fsLR
- Repetition Time (TR): 0.72
- Mean Framewise Displacement: 0.1047
- Mean Relative RMS Motion: 0.5133
- Max Relative RMS Motion: 0.6878
- DVARS Before and After Processing : 486.2628, 27.0057
- Correlation between DVARS and FD Before and After Processing : 0.1505, 0.0738
- Number of Volumes Censored : 4

#### Alignment of functional and anatomical MRI data (surface driven)

bbregister was used to coregister functional and anatomical MRI data.

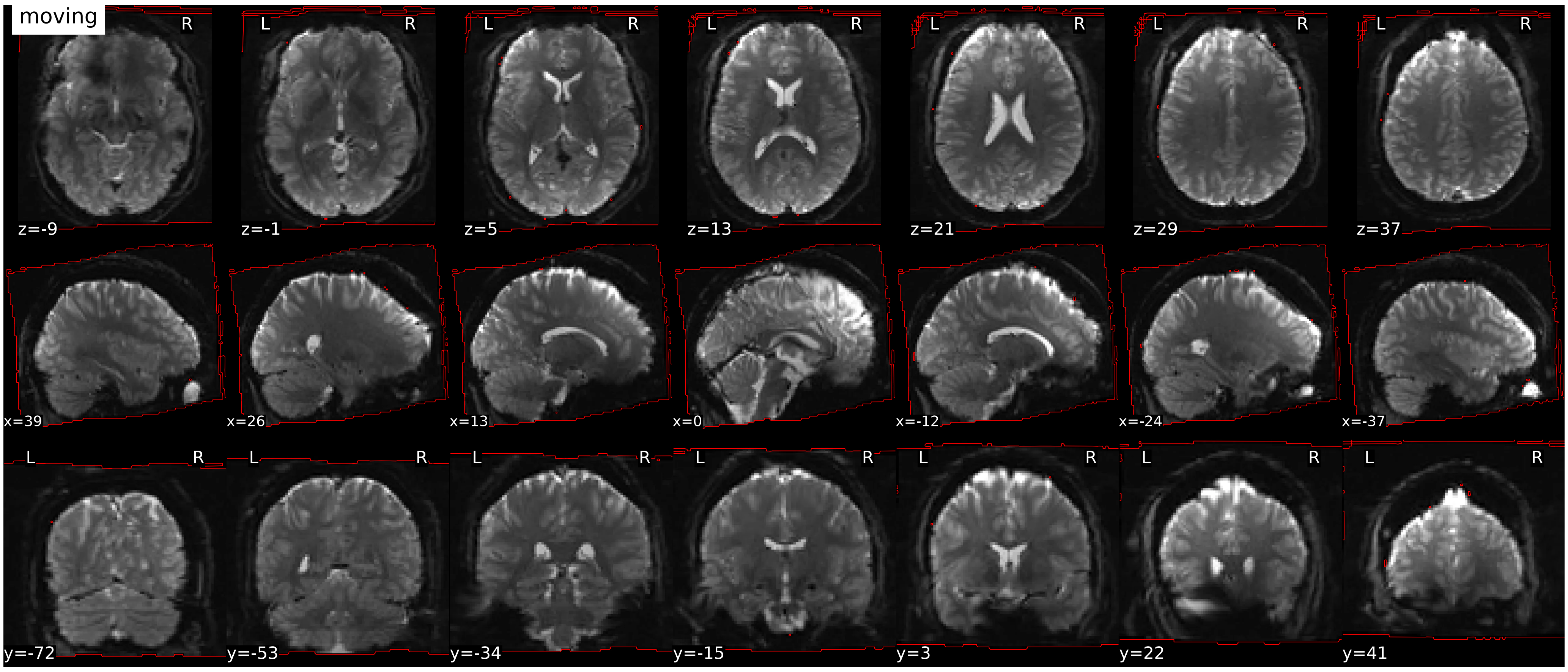


Get figure file: sub-113922/figures/sub-113922_task-REST2_dir-RL_space-MNI152NLin6Asym_desc-bbregister_bold.svg

#### Carpet Plot Before Postprocessing

FD and DVARS are two measures of in-scanner motion. This plot shows standardized FD, DVARS, and then a carpet plot for the time series of each voxel/vertex's time series of activity.

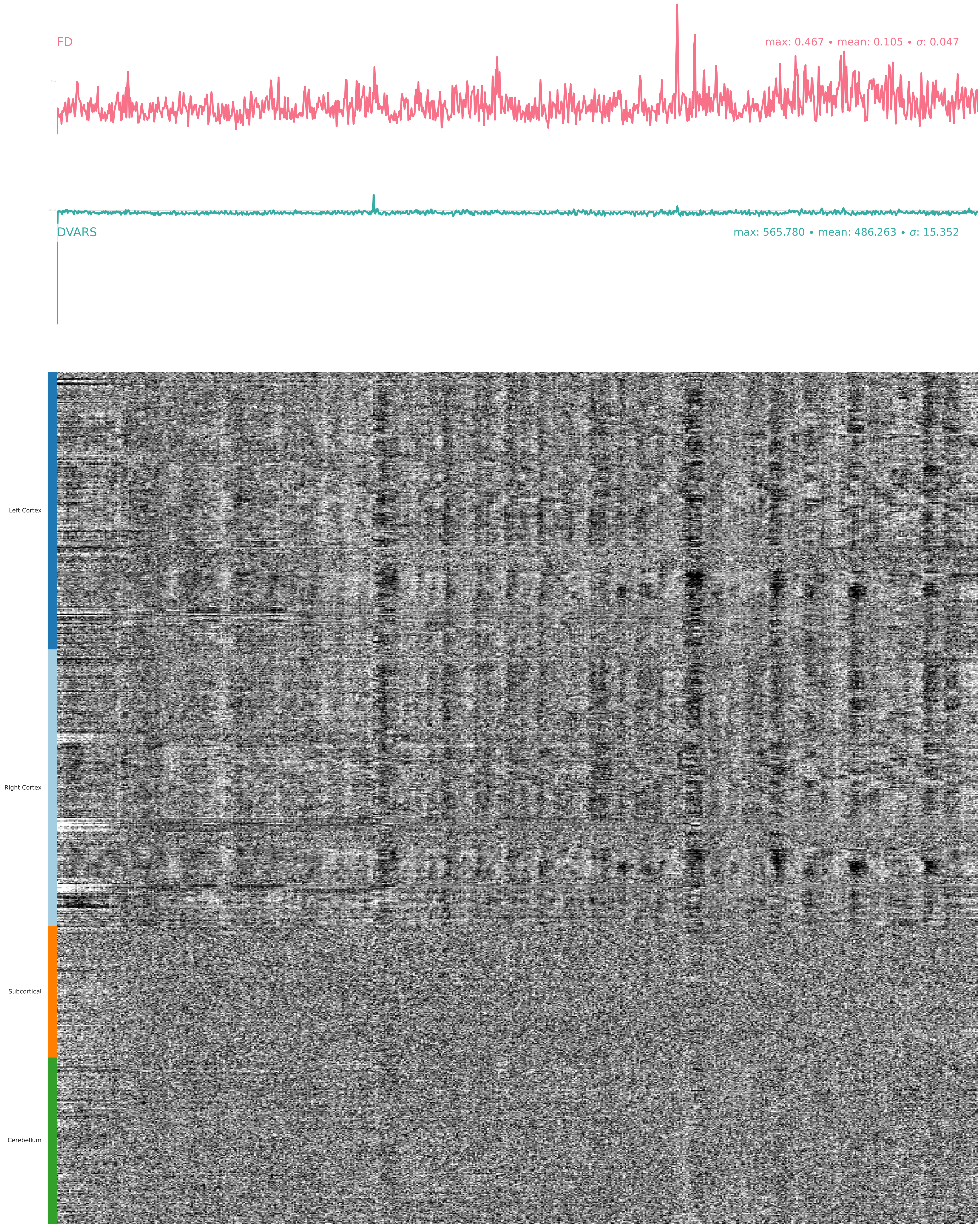


Get figure file: sub-113922/figures/sub-113922_task-REST2_dir-RL_space-fsLR_desc-preprocessing_bold.svg

#### Framewise Displacement and Censored Volumes

Framewise displacement (FD) is used to flag high-motion volumes, which are then censored as part of the denoising procedure. If motion filtering is requested, then the six translation and rotation motion parameters are filtered to remove respiratory effects before FD is calculated and outlier volumes are identified.

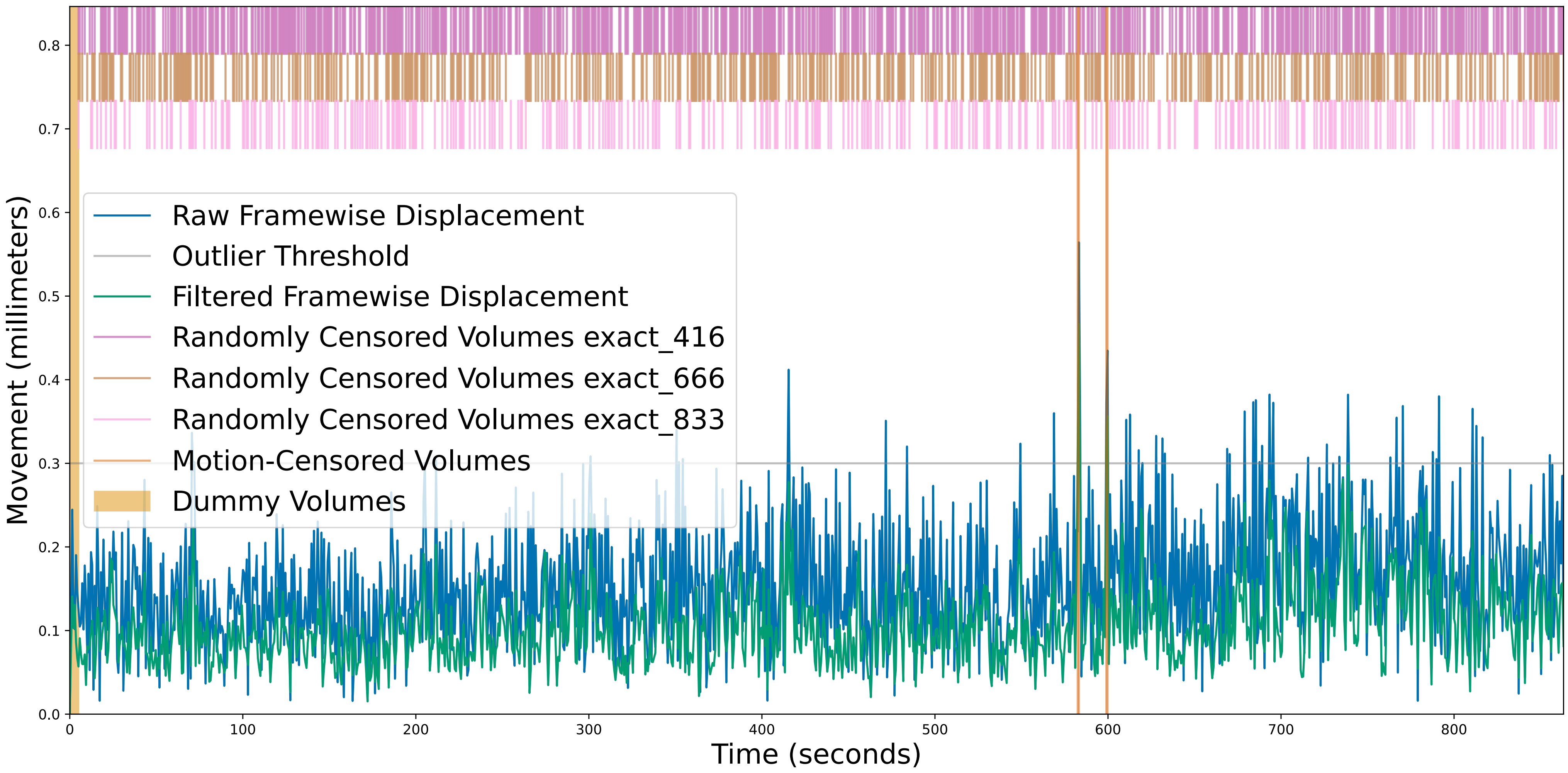


Get figure file: sub-113922/figures/sub-113922_task-REST2_dir-RL_space-fsLR_desc-censoring_motion.svg

#### Design Matrix for Confound Regression

The "design matrix" represents the confounds that are used to denoise the BOLD data.

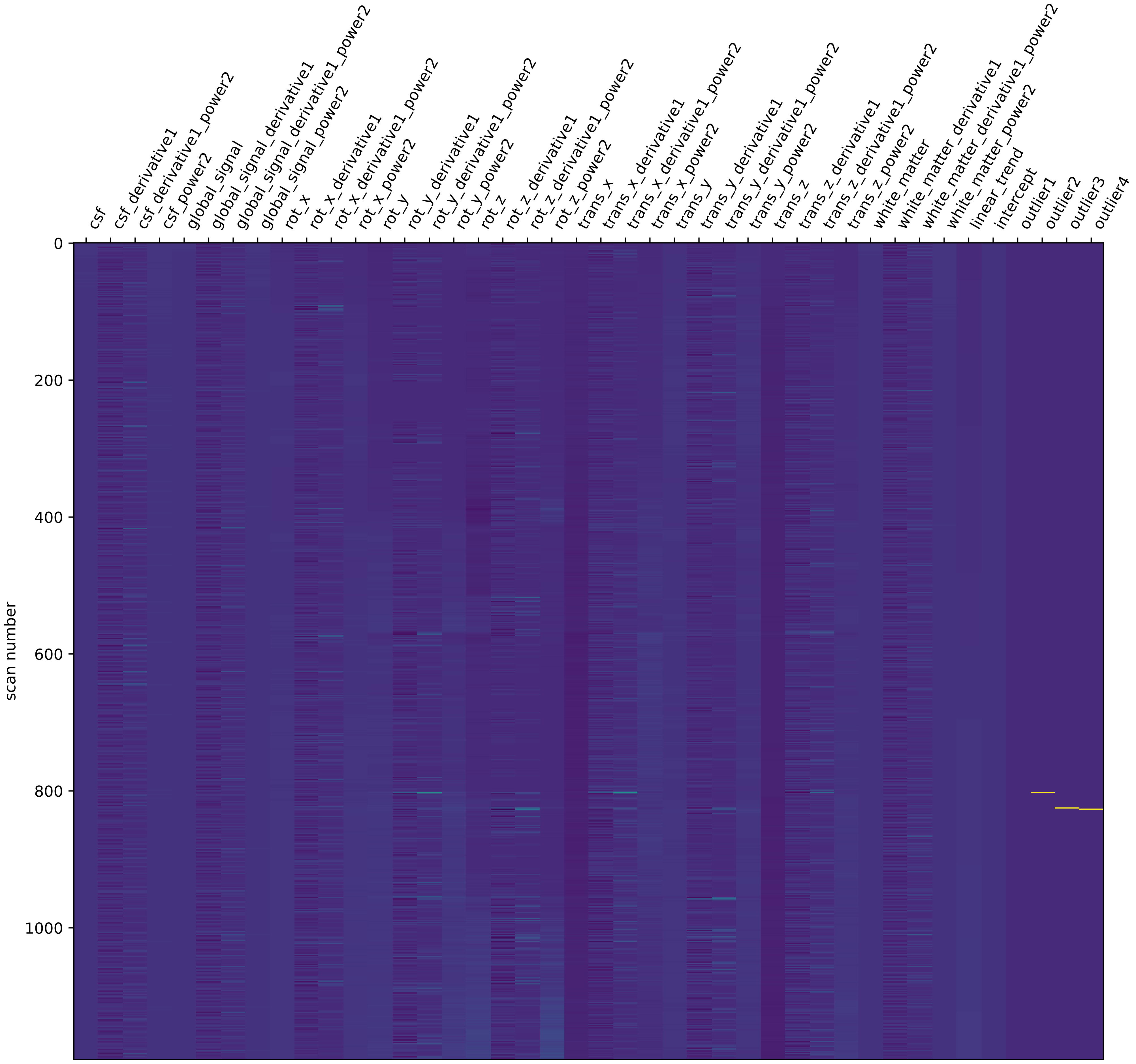


Get figure file: [sub-113922/figures/sub-113922_task-REST2_dir-RL_design.svg](sub-113922/figures/sub-113922_task-REST2_dir-RL_design.svg)

#### Carpet Plot After Postprocessing

FD and DVARS are two measures of in-scanner motion. This plot shows standardized FD, DVARS, and then a carpet plot for the time series of each voxel/vertex's time series of activity.

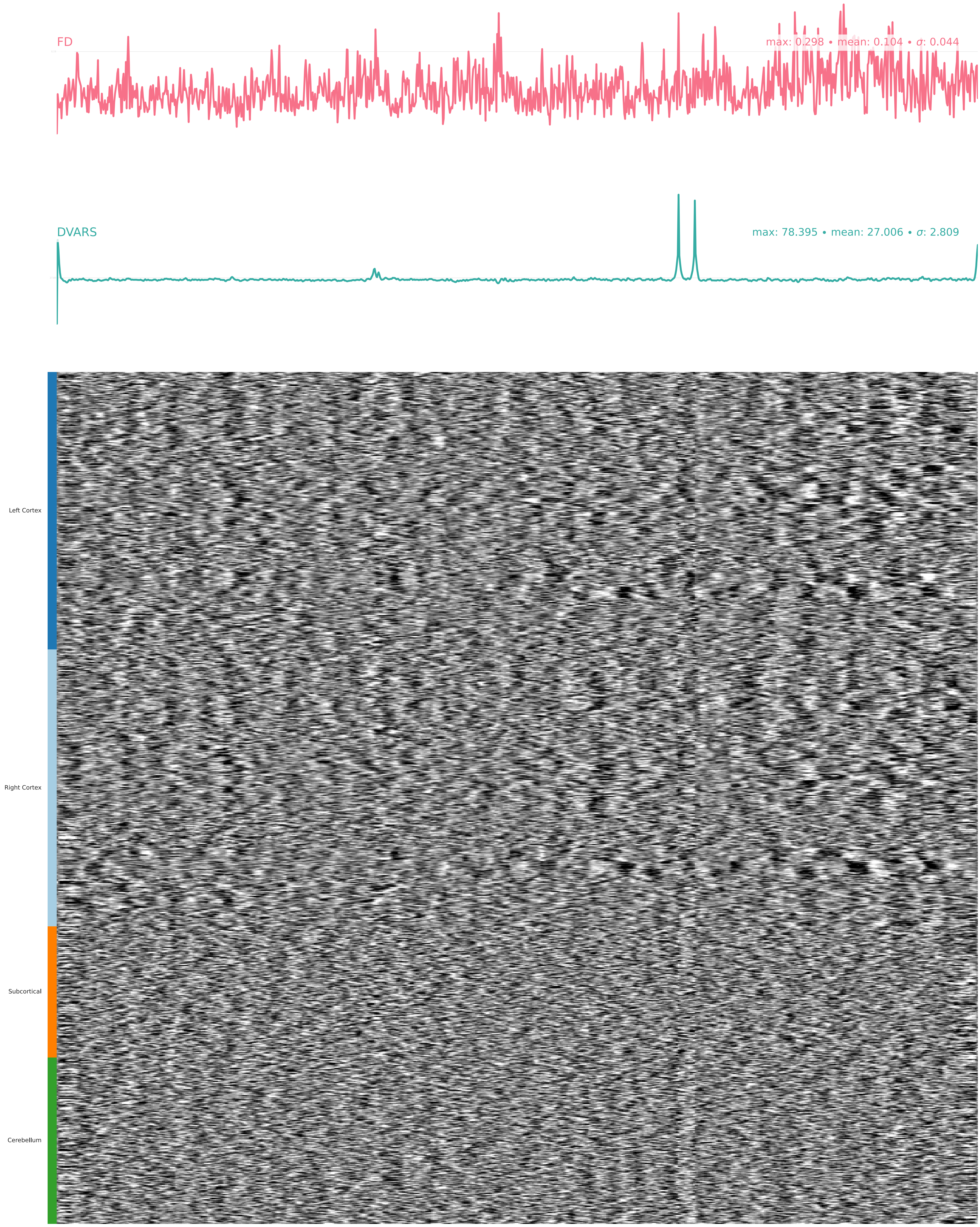


Get figure file: [sub-113922/figures/sub-113922_task-REST2_dir-RL_space-fsLR_desc-postprocessing_bold.svg](sub-113922/figures/sub-113922_task-REST2_dir-RL_space-fsLR_desc-postprocessing_bold.svg)

#### Correlation Heatmaps from Four Atlases

This plot shows heatmaps from ROI-to-ROI correlations from four atlases.

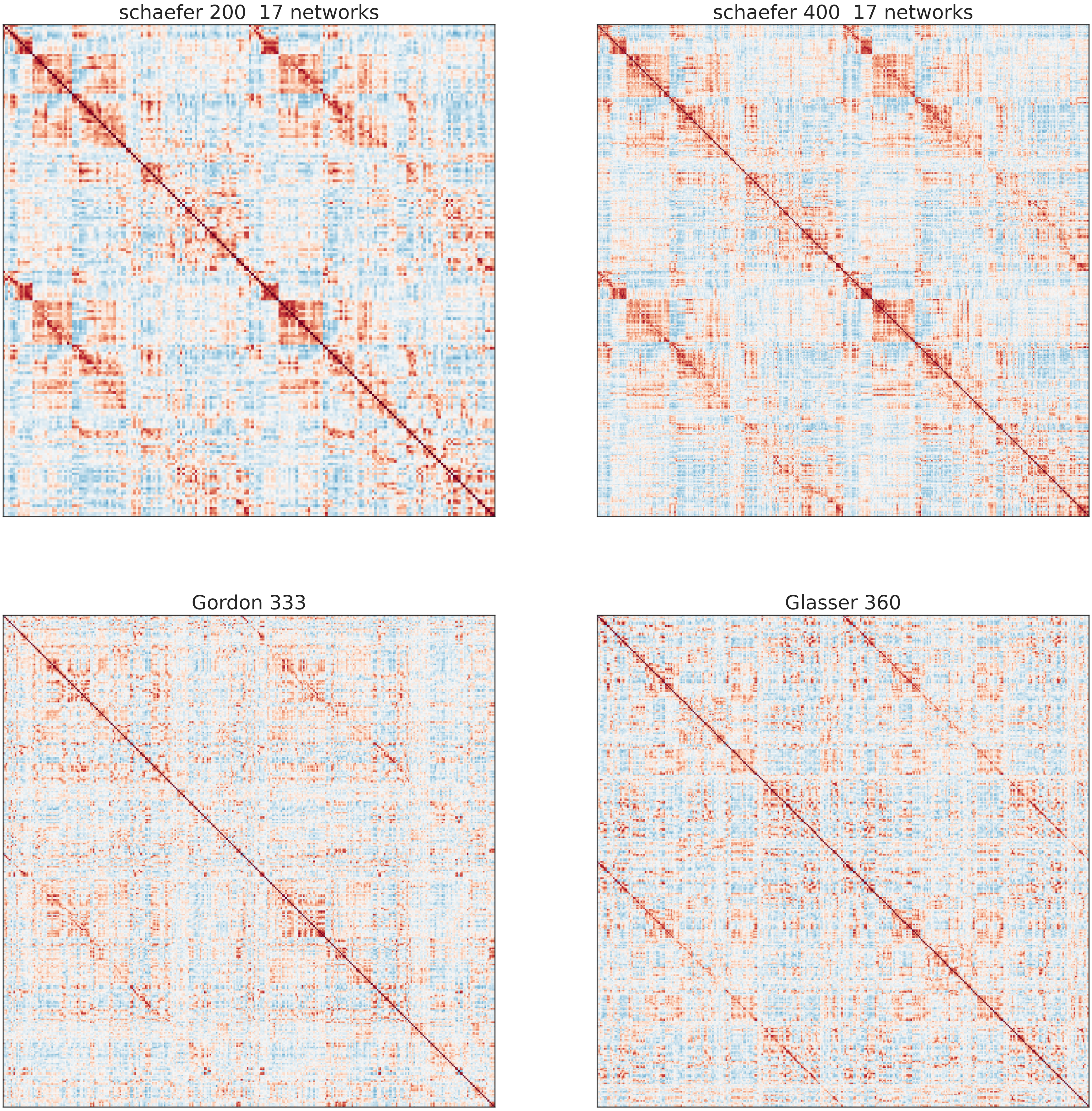


Get figure file: sub-113922/figures/sub-113922_task-REST2_dir-RL_space-fsLR_desc-connectivityplot_bold.svg

### Reports for: task SOCIAL, direction LR.

#### Summary

- BOLD volume space: fsLR
- Repetition Time (TR): 0.72
- Mean Framewise Displacement: 0.1021
- Mean Relative RMS Motion: 0.1386
- Max Relative RMS Motion: 0.2137
- DVARS Before and After Processing : 479.3113, 24.8617
- Correlation between DVARS and FD Before and After Processing : 0.113, -0.0331
- Number of Volumes Censored : 3

#### Alignment of functional and anatomical MRI data (surface driven)

bbregister was used to coregister functional and anatomical MRI data.

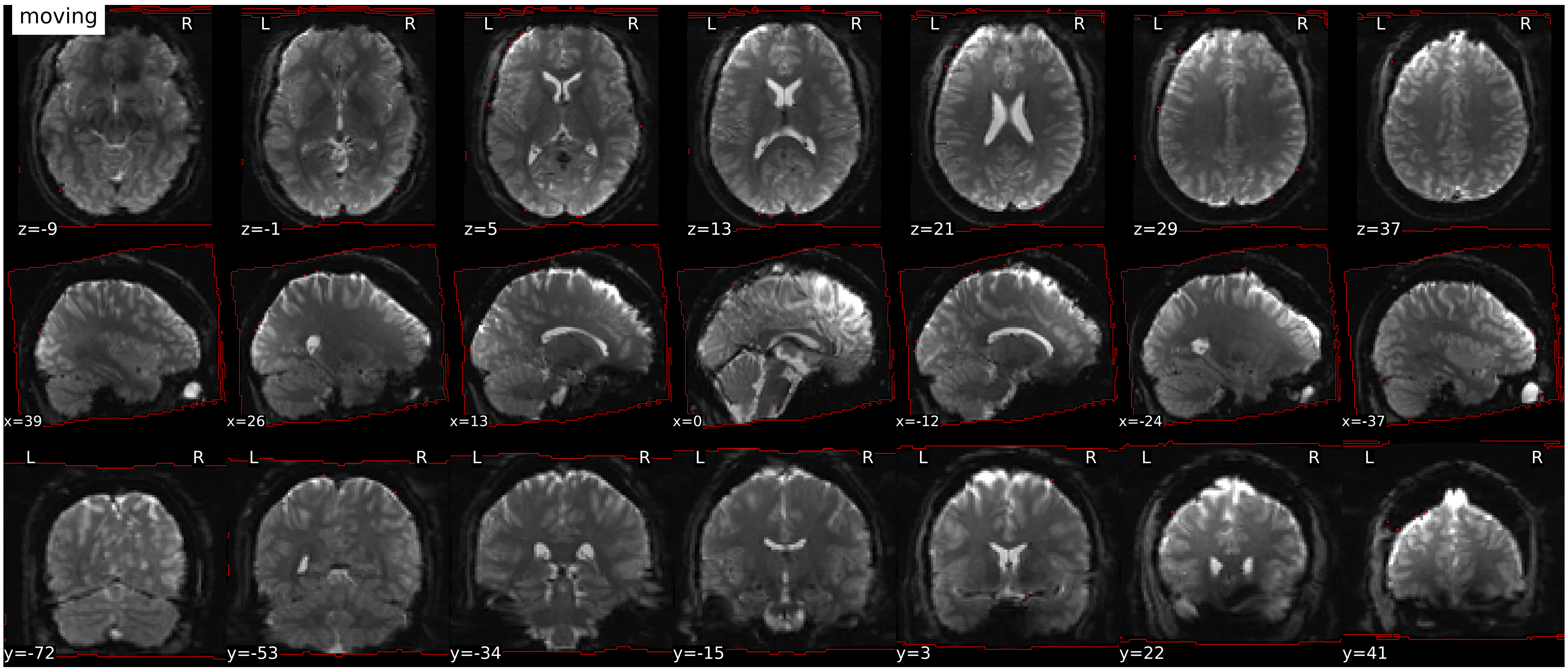


Get figure file: sub-113922/figures/sub-113922_task-SOCIAL_dir-LR_space-MNI152NLin6Asym_desc-bbregister_bold.svg

#### Carpet Plot Before Postprocessing

FD and DVARS are two measures of in-scanner motion. This plot shows standardized FD, DVARS, and then a carpet plot for the time series of each voxel/vertex's time series of activity.

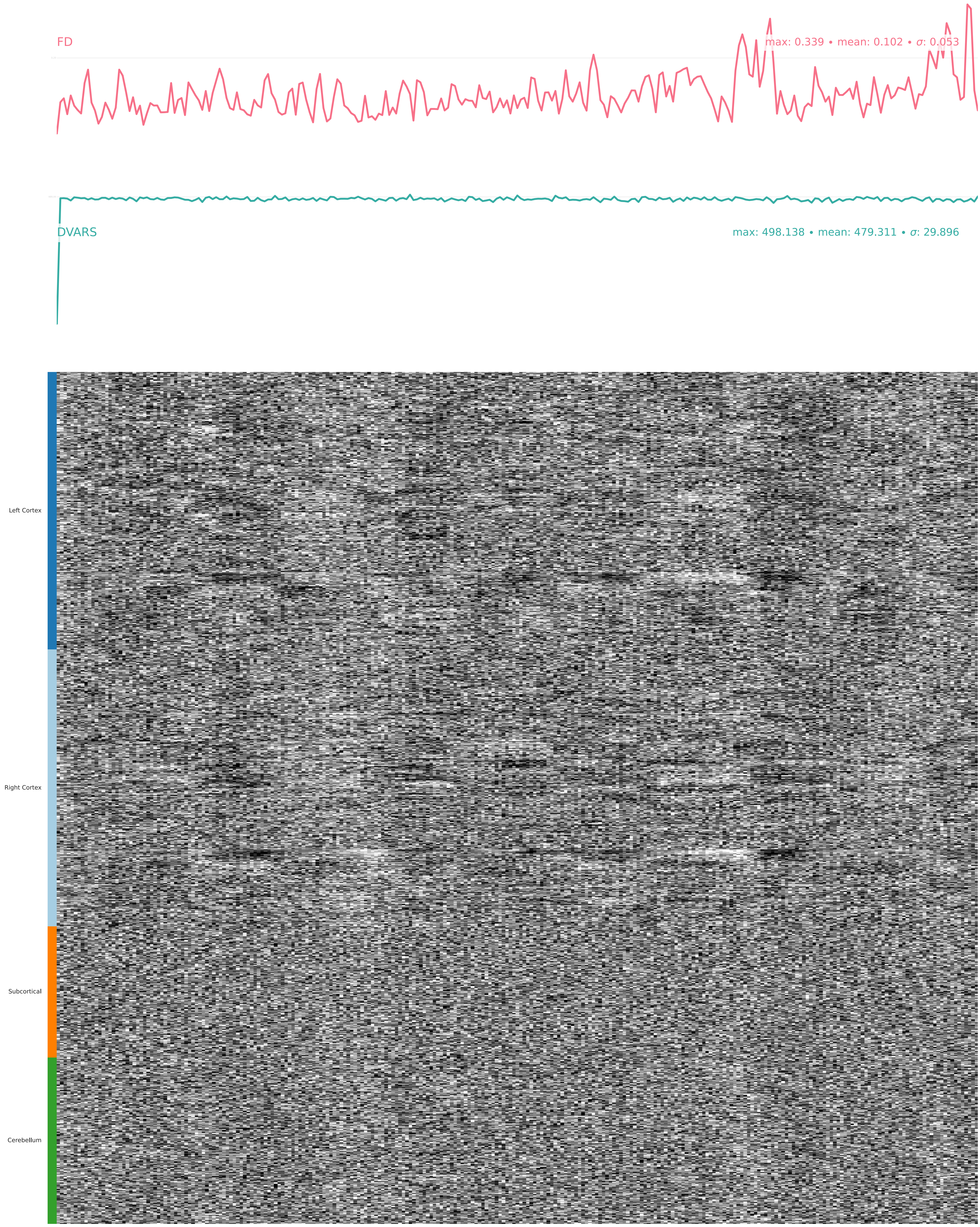


Get figure file: sub-113922/figures/sub-113922_task-SOCIAL_dir-LR_space-fsLR_desc-preprocessing_bold.svg

#### Framewise Displacement and Censored Volumes

Framewise displacement (FD) is used to flag high-motion volumes, which are then censored as part of the denoising procedure. If motion filtering is requested, then the six translation and rotation motion parameters are filtered to remove respiratory effects before FD is calculated and outlier volumes are identified.

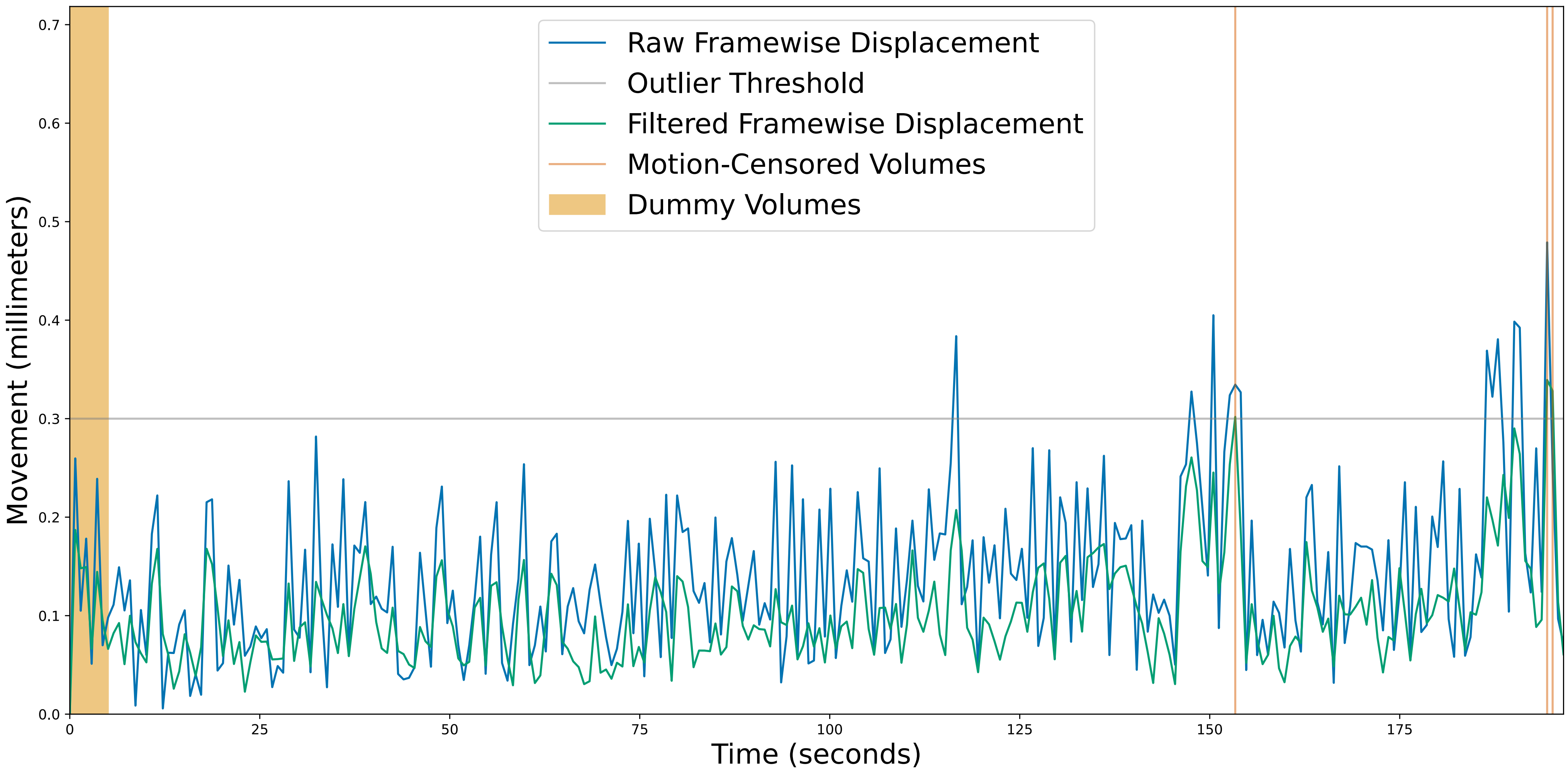


Get figure file: sub-113922/figures/sub-113922_task-SOCIAL_dir-LR_space-fsLR_desc-censoring_motion.svg

#### Design Matrix for Confound Regression

The "design matrix" represents the confounds that are used to denoise the BOLD data.

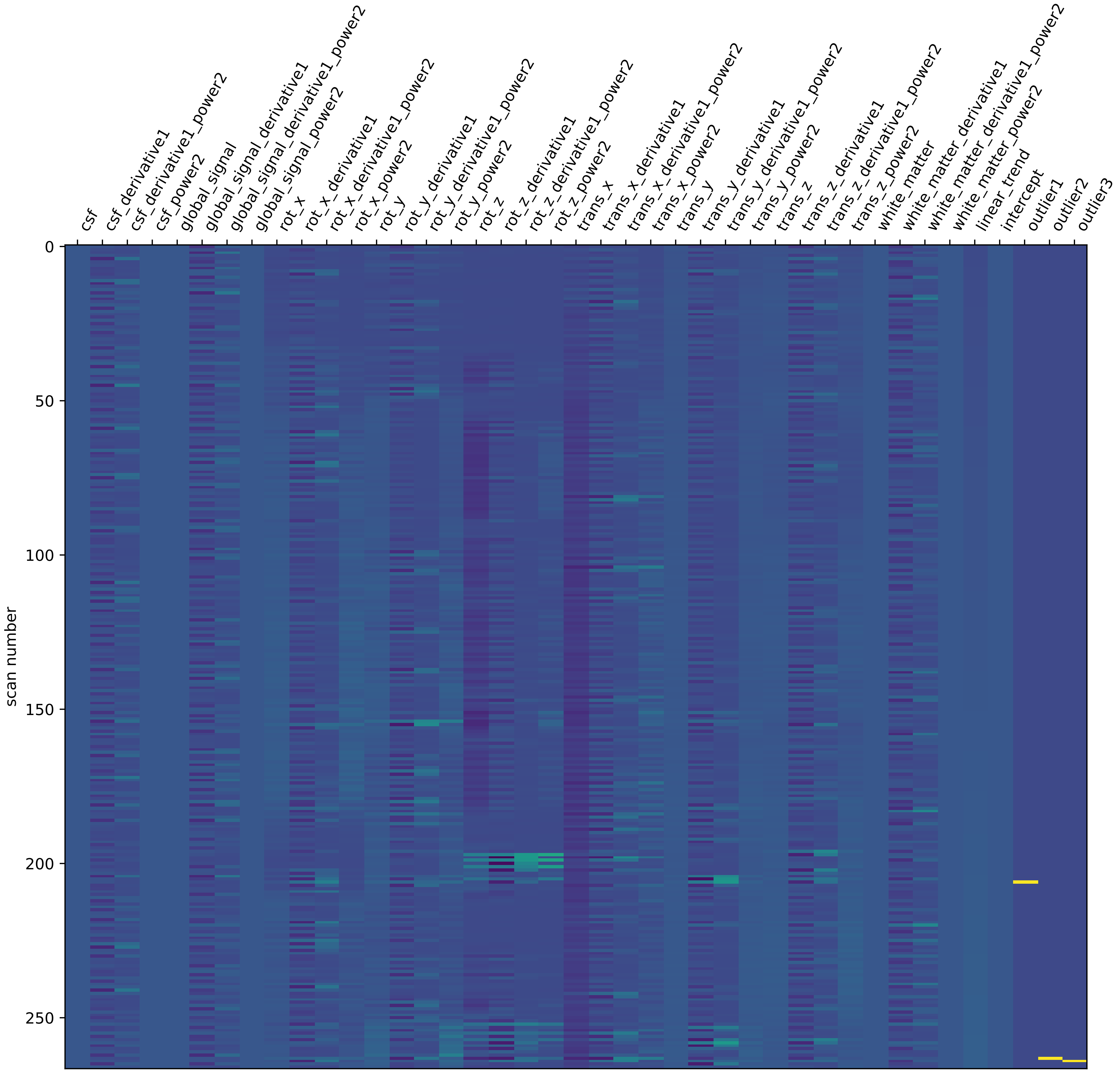


Get figure file: sub-113922/figures/sub-113922_task-SOCIAL_dir-LR_design.svg

#### Carpet Plot After Postprocessing

FD and DVARS are two measures of in-scanner motion. This plot shows standardized FD, DVARS, and then a carpet plot for the time series of each voxel/vertex's time series of activity.

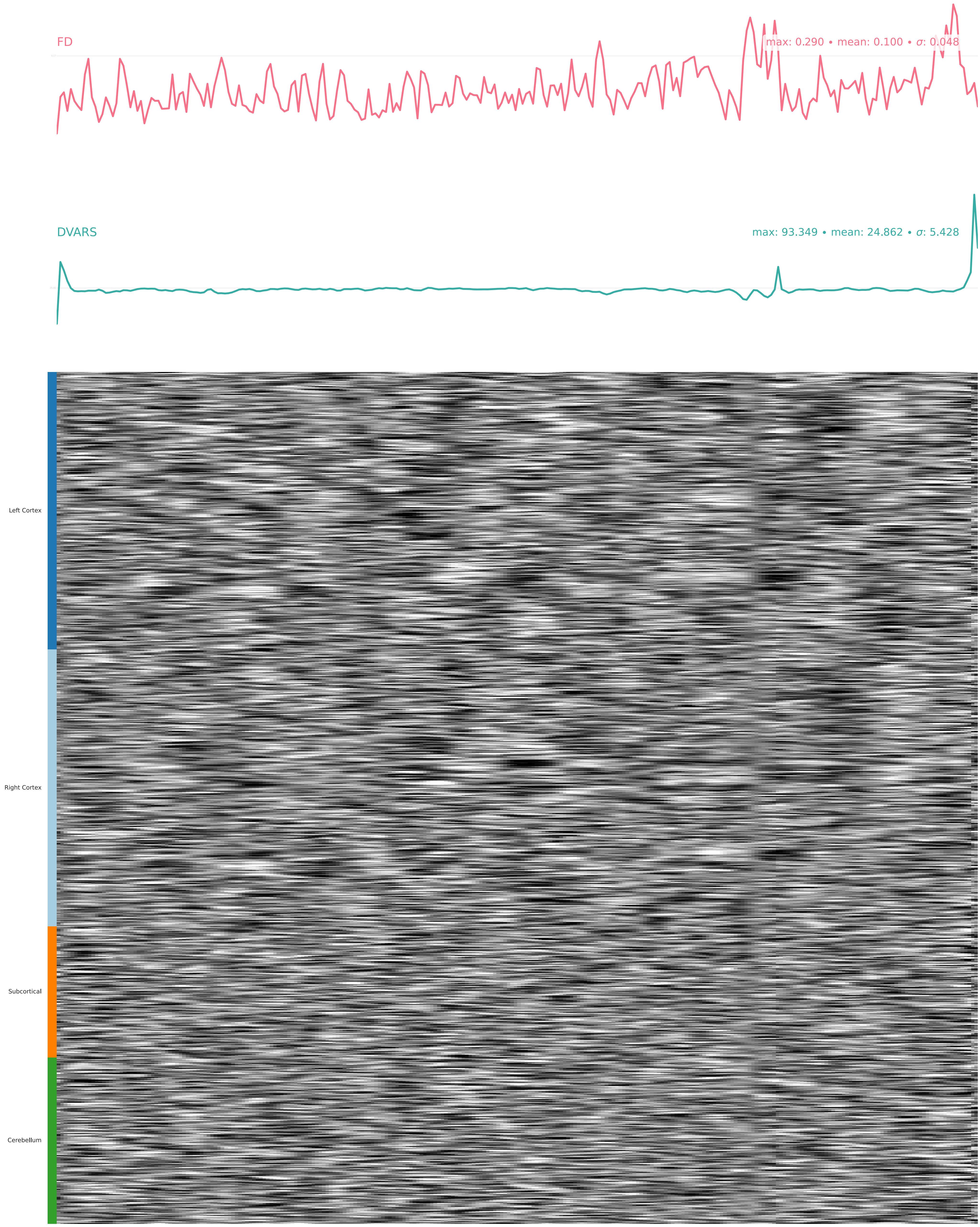


Get figure file: sub-113922/figures/sub-113922_task-SOCIAL_dir-LR_space-fsLR_desc-postprocessing_bold.svg

#### Correlation Heatmaps from Four Atlases

This plot shows heatmaps from ROI-to-ROI correlations from four atlases.

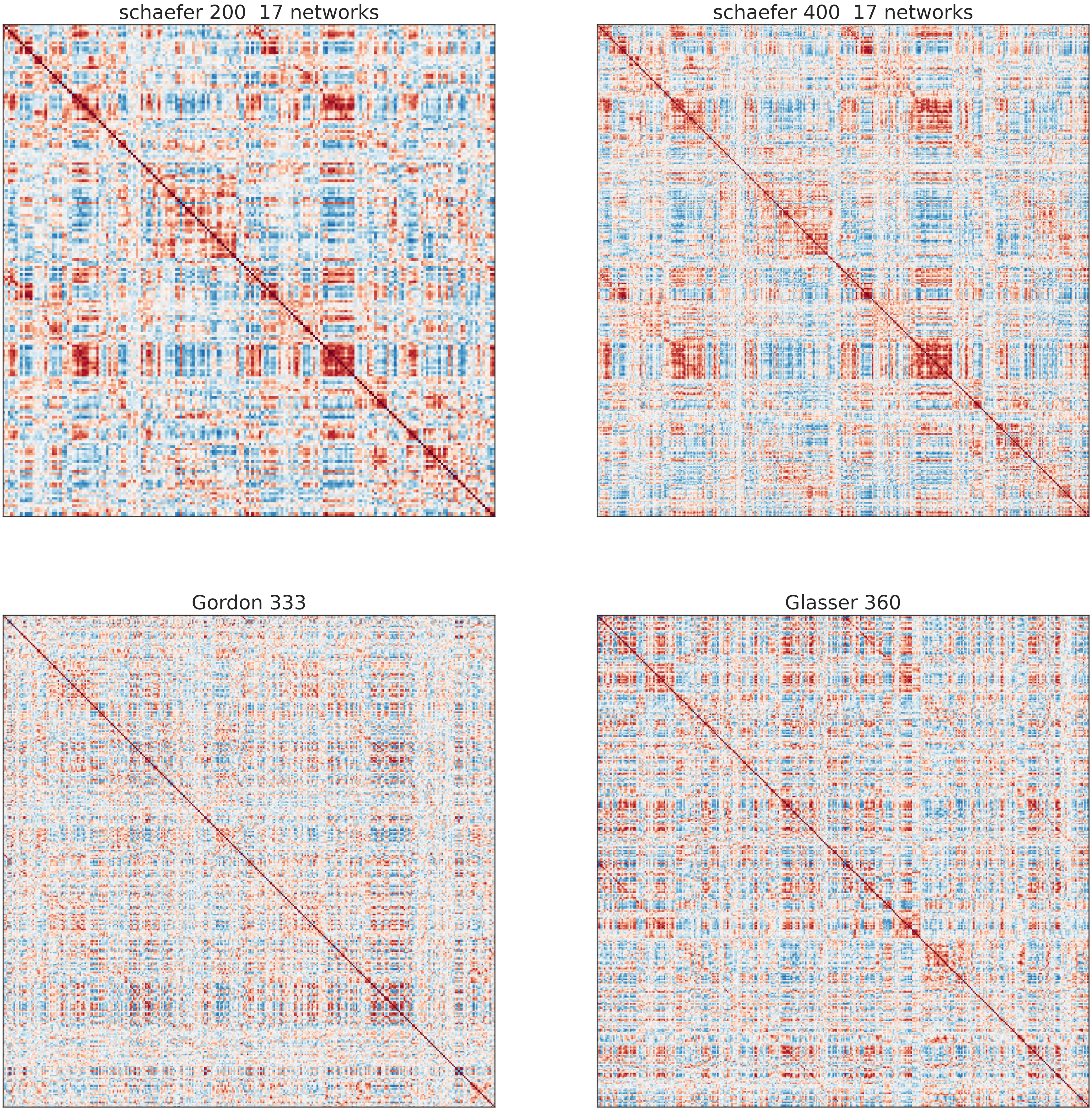


Get figure file: sub-113922/figures/sub-113922_task-SOCIAL_dir-LR_space-fsLR_desc-connectivityplot_bold.svg

### Reports for: task SOCIAL, direction RL.

#### Summary

- BOLD volume space: fsLR
- Repetition Time (TR): 0.72
- Mean Framewise Displacement: 0.1175
- Mean Relative RMS Motion: 0.2044
- Max Relative RMS Motion: 0.3919
- DVARS Before and After Processing : 485.869, 25.4153
- Correlation between DVARS and FD Before and After Processing : 0.1904, 0.108
- Number of Volumes Censored : 4

#### Alignment of functional and anatomical MRI data (surface driven)

bbregister was used to coregister functional and anatomical MRI data.

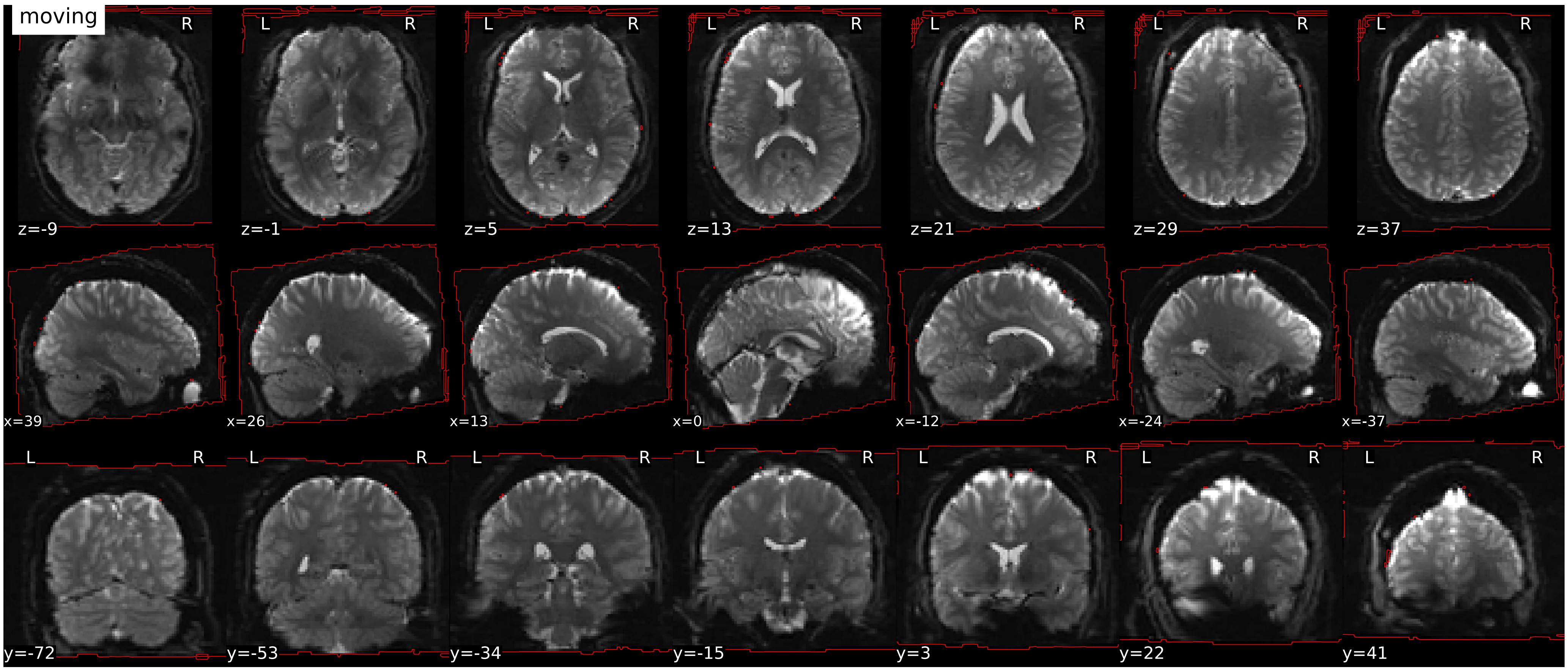


Get figure file: sub-113922/figures/sub-113922_task-SOCIAL_dir-RL_space-MNI152NLin6Asym_desc-bbregister_bold.svg

#### Carpet Plot Before Postprocessing

FD and DVARS are two measures of in-scanner motion. This plot shows standardized FD, DVARS, and then a carpet plot for the time series of each voxel/vertex's time series of activity.

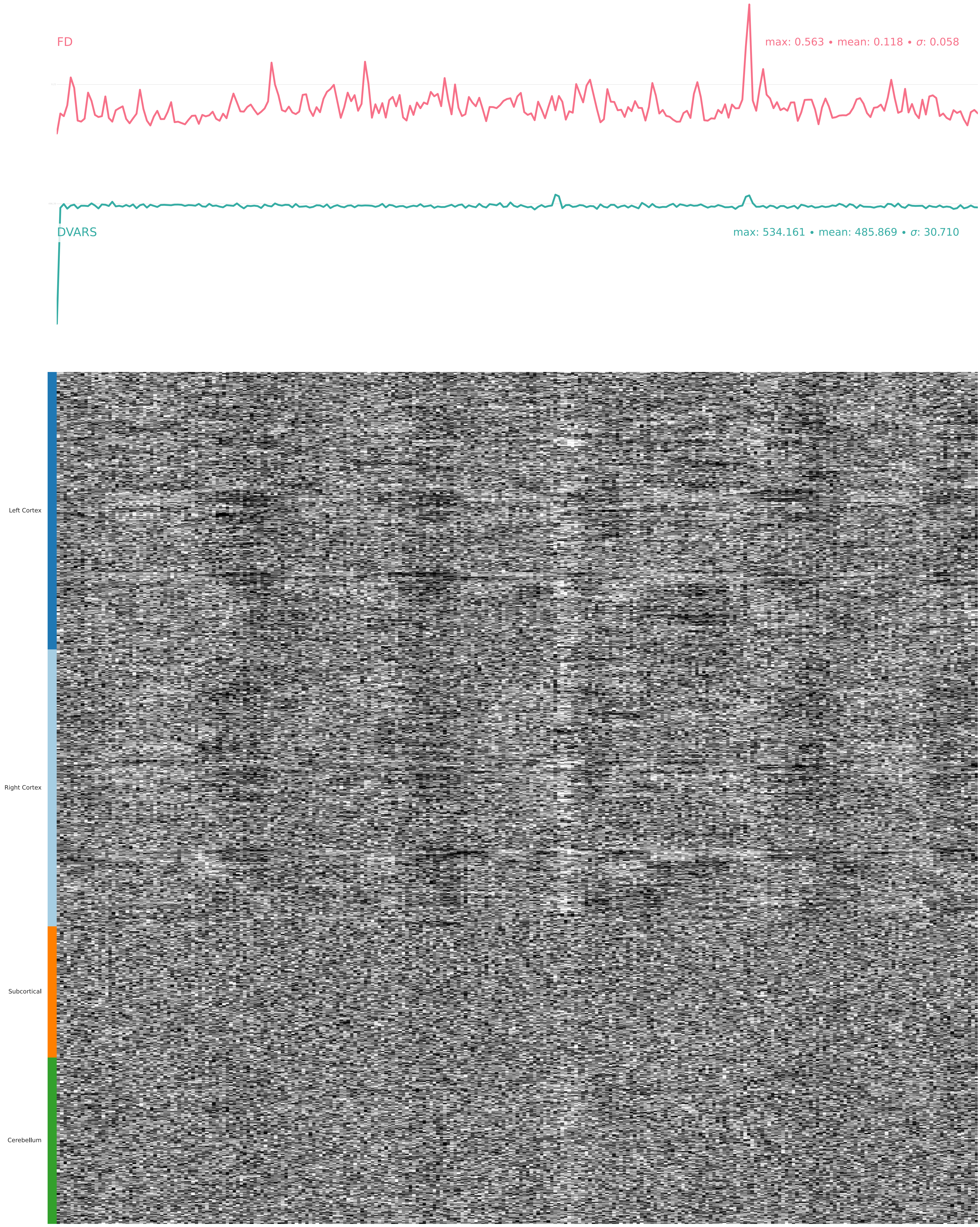


Get figure file: sub-113922/figures/sub-113922_task-SOCIAL_dir-RL_space-fsLR_desc-preprocessing_bold.svg

#### Framewise Displacement and Censored Volumes

Framewise displacement (FD) is used to flag high-motion volumes, which are then censored as part of the denoising procedure. If motion filtering is requested, then the six translation and rotation motion parameters are filtered to remove respiratory effects before FD is calculated and outlier volumes are identified.

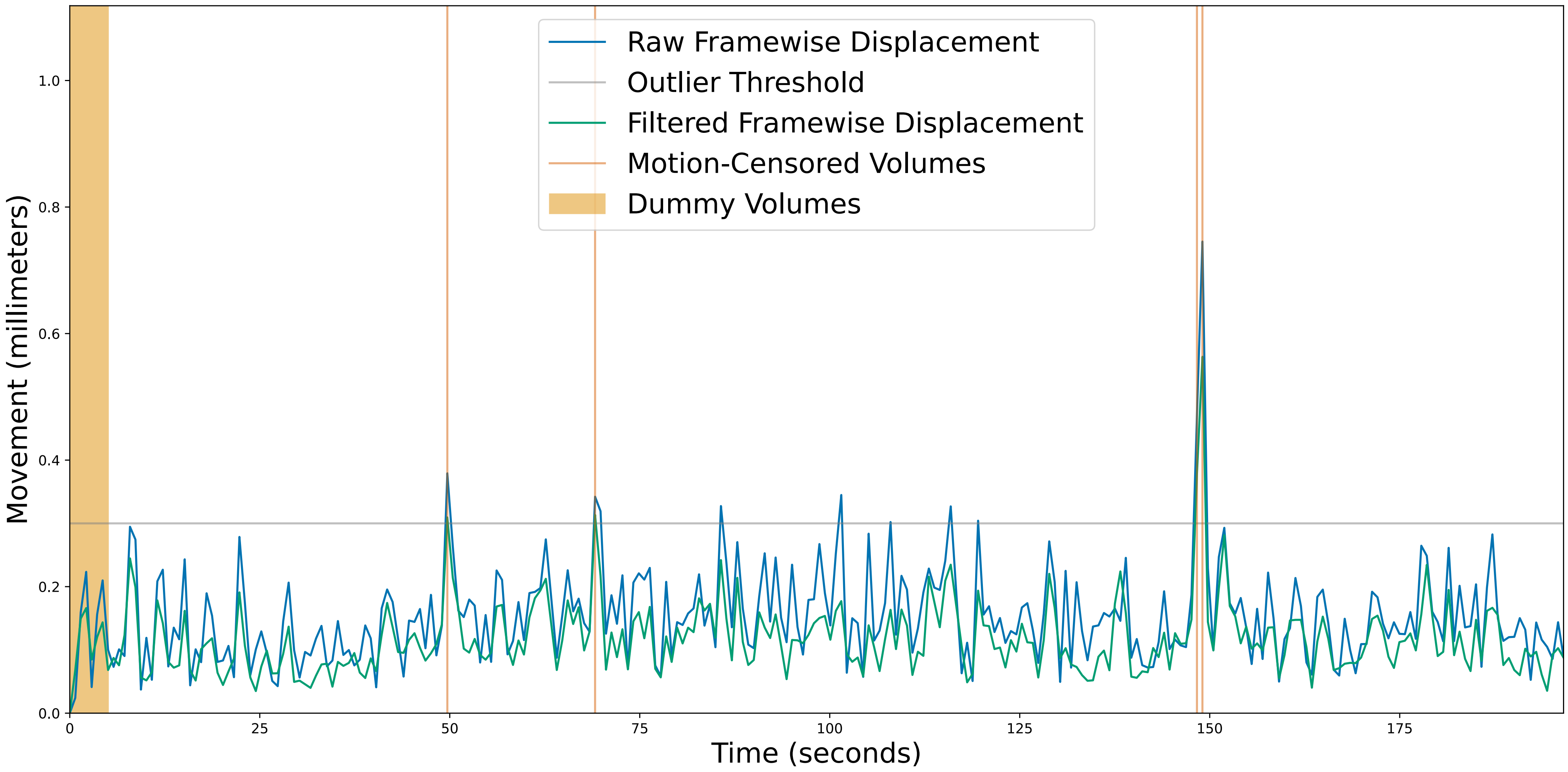


Get figure file: [sub-113922/figures/sub-113922_task-SOCIAL_dir-RL_space-fsLR_desc-censoring_motion.svg](sub-113922/figures/sub-113922_task-SOCIAL_dir-RL_space-fsLR_desc-censoring_motion.svg)

#### Design Matrix for Confound Regression

The "design matrix" represents the confounds that are used to denoise the BOLD data.

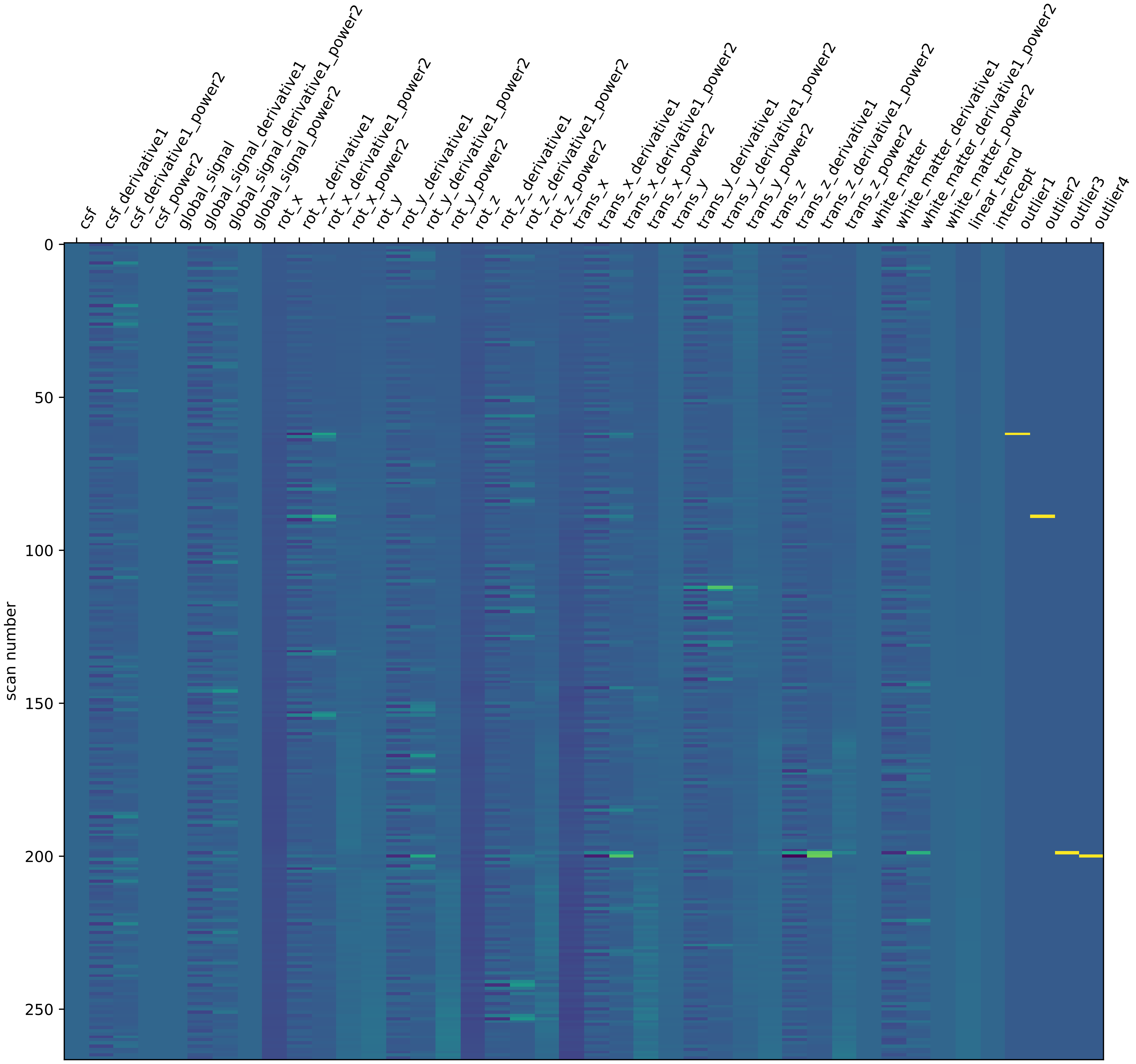


Get figure file: [sub-113922/figures/sub-113922_task-SOCIAL_dir-RL_design.svg](sub-113922/figures/sub-113922_task-SOCIAL_dir-RL_design.svg)

#### Carpet Plot After Postprocessing

FD and DVARS are two measures of in-scanner motion. This plot shows standardized FD, DVARS, and then a carpet plot for the time series of each voxel/vertex's time series of activity.

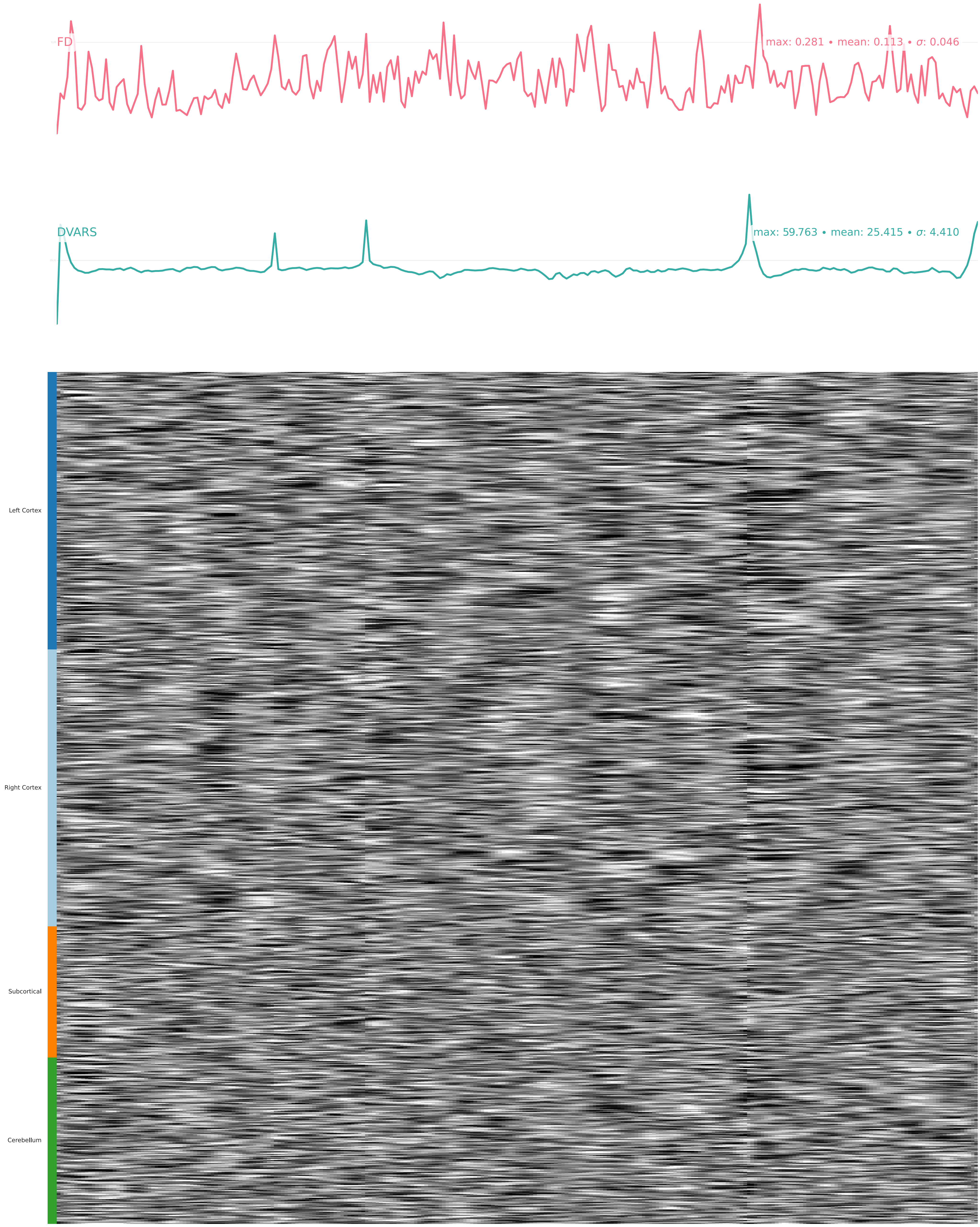


Get figure file: sub-113922/figures/sub-113922_task-SOCIAL_dir-RL_space-fsLR_desc-postprocessing_bold.svg

#### Correlation Heatmaps from Four Atlases

This plot shows heatmaps from ROI-to-ROI correlations from four atlases.

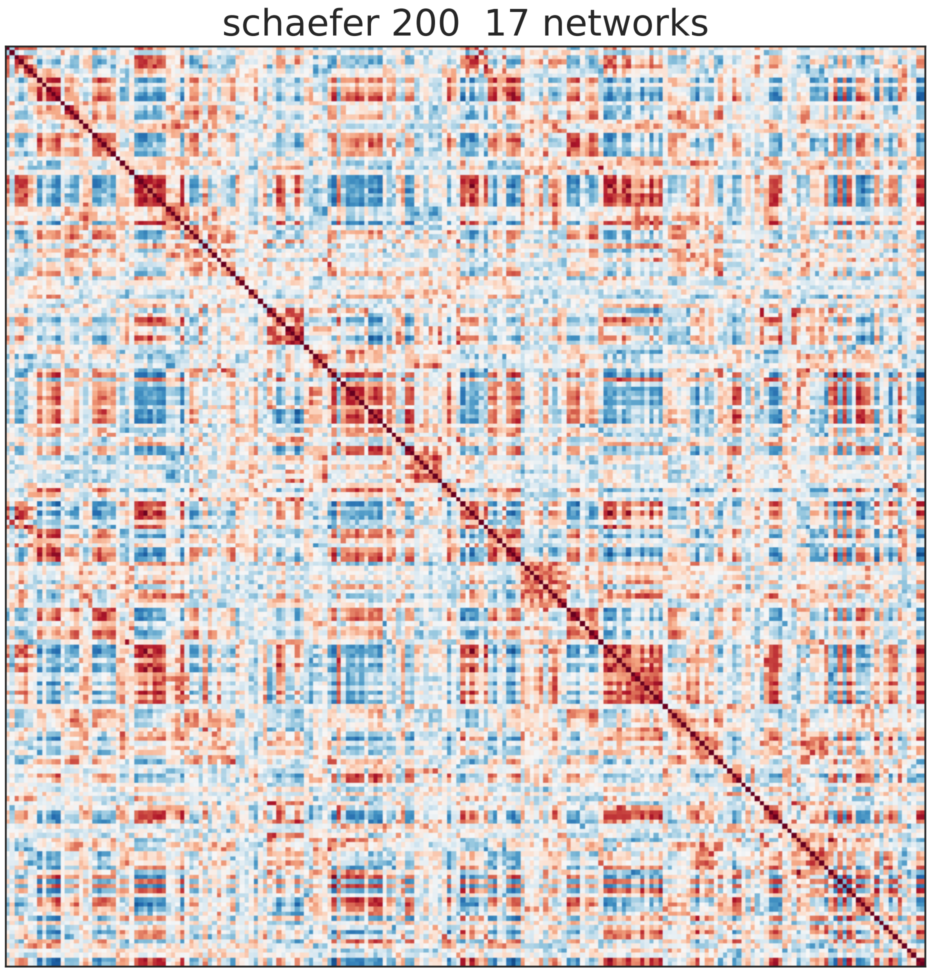

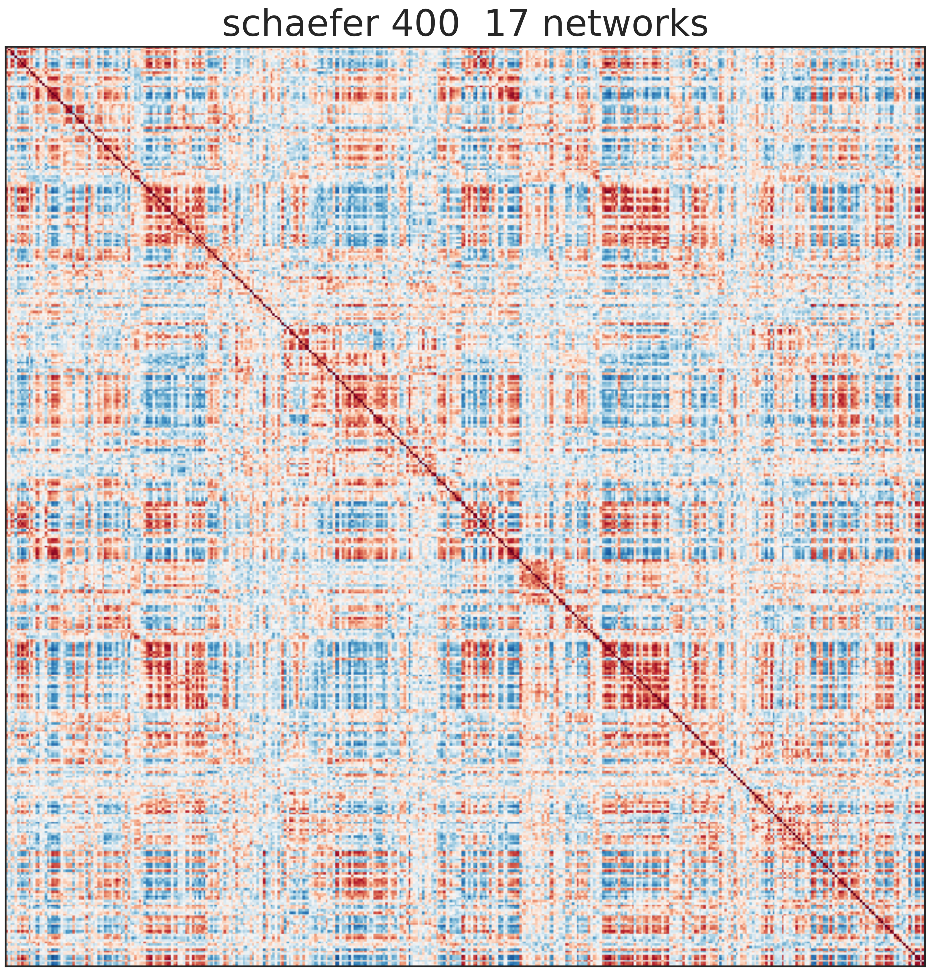

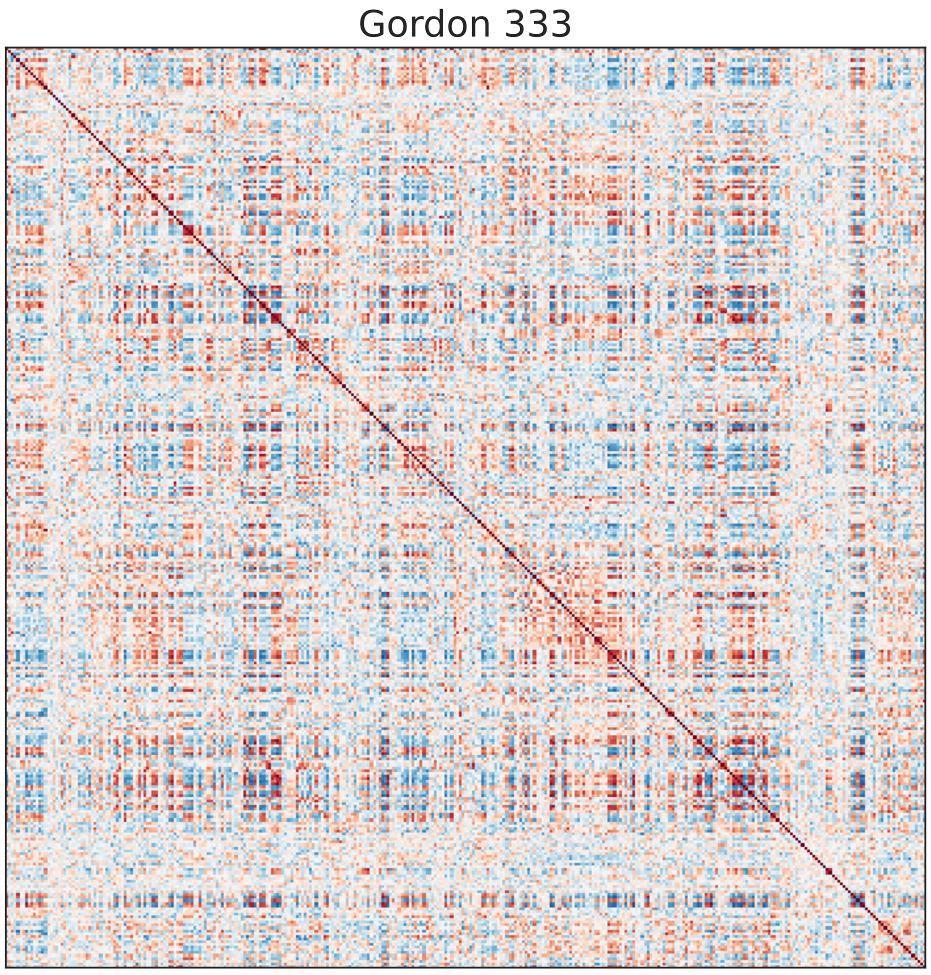

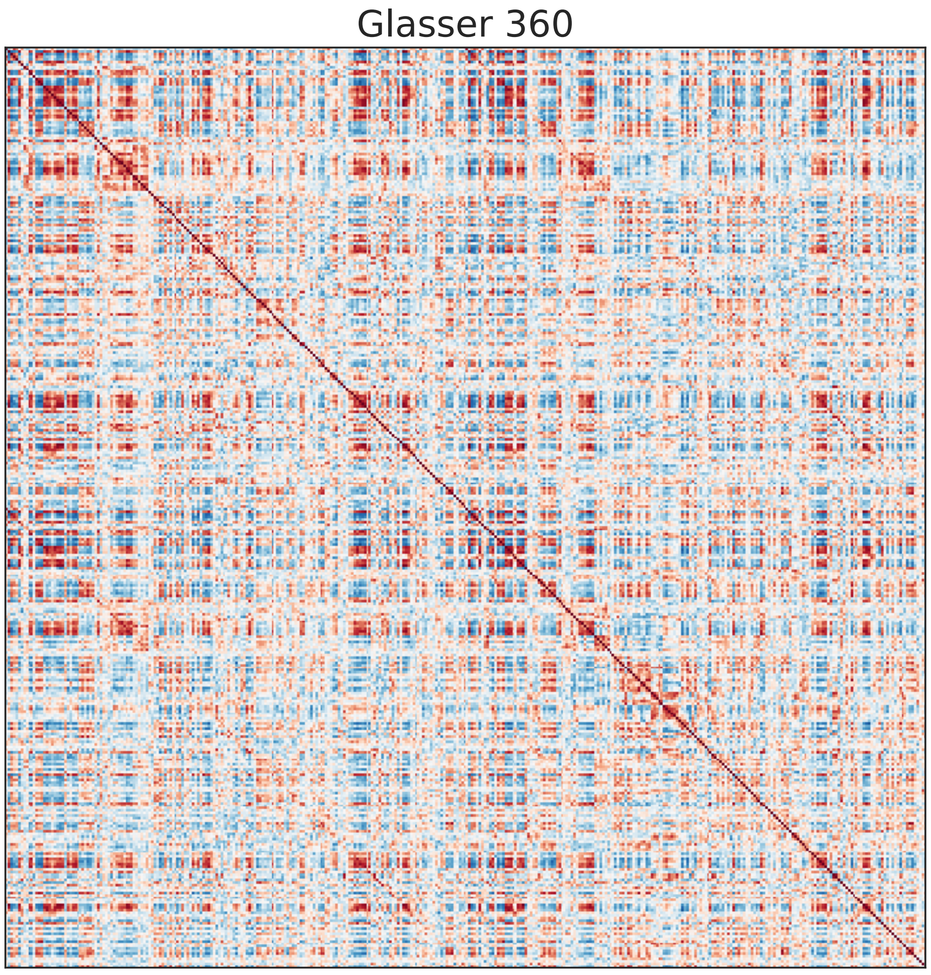


Get figure file: sub-113922/figures/sub-113922_task-SOCIAL_dir-RL_space-fsLR_desc-connectivityplot_bold.svg

### Reports for: task WM, direction LR.

#### Summary

- BOLD volume space: fsLR
- Repetition Time (TR): 0.72
- Mean Framewise Displacement: 0.1149
- Mean Relative RMS Motion: 0.5349
- Max Relative RMS Motion: 0.6891
- DVARS Before and After Processing : 471.2848, 25.3087
- Correlation between DVARS and FD Before and After Processing : 0.1509, 0.0434
- Number of Volumes Censored : 1

#### Alignment of functional and anatomical MRI data (surface driven)

bbregister was used to coregister functional and anatomical MRI data.

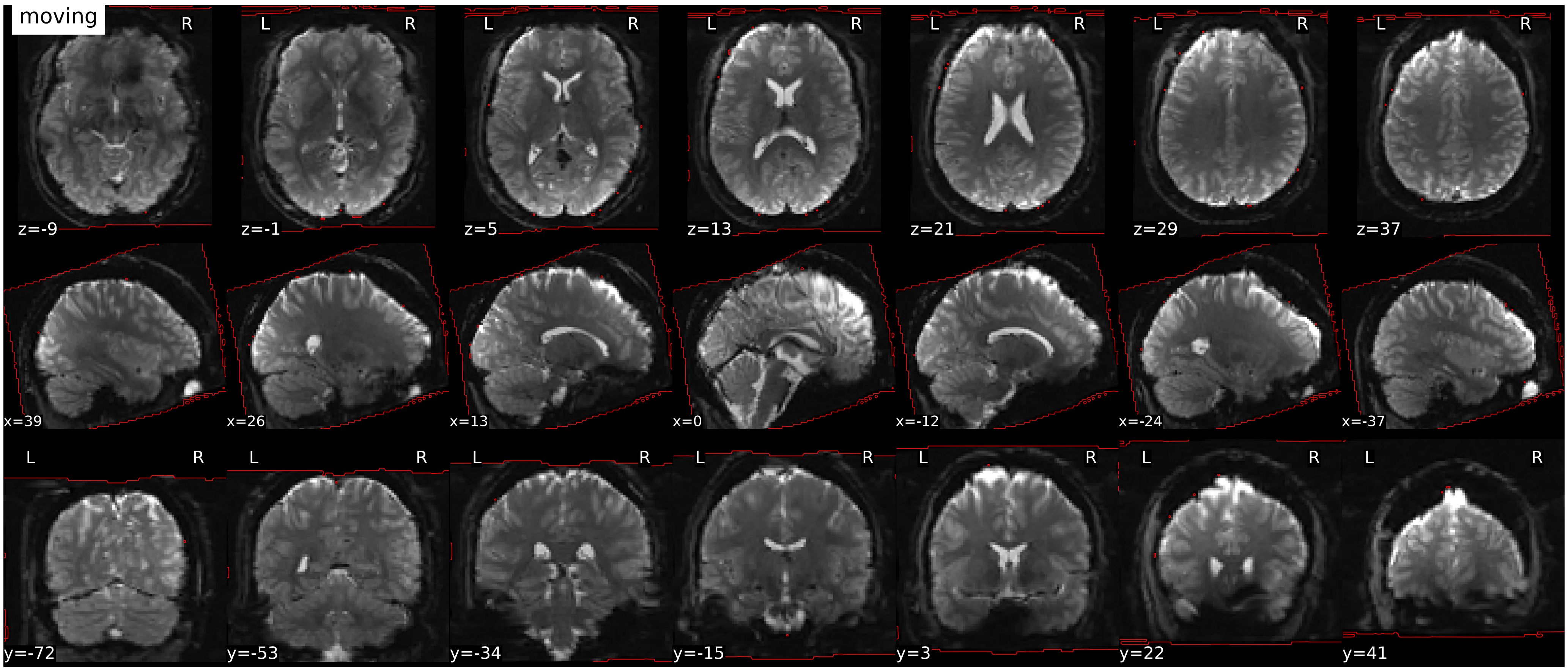


Get figure file: sub-113922/figures/sub-113922_task-WM_dir-LR_space-MNI152NLin6Asym_desc-bbregister_bold.svg

#### Carpet Plot Before Postprocessing

FD and DVARS are two measures of in-scanner motion. This plot shows standardized FD, DVARS, and then a carpet plot for the time series of each voxel/vertex's time series of activity.

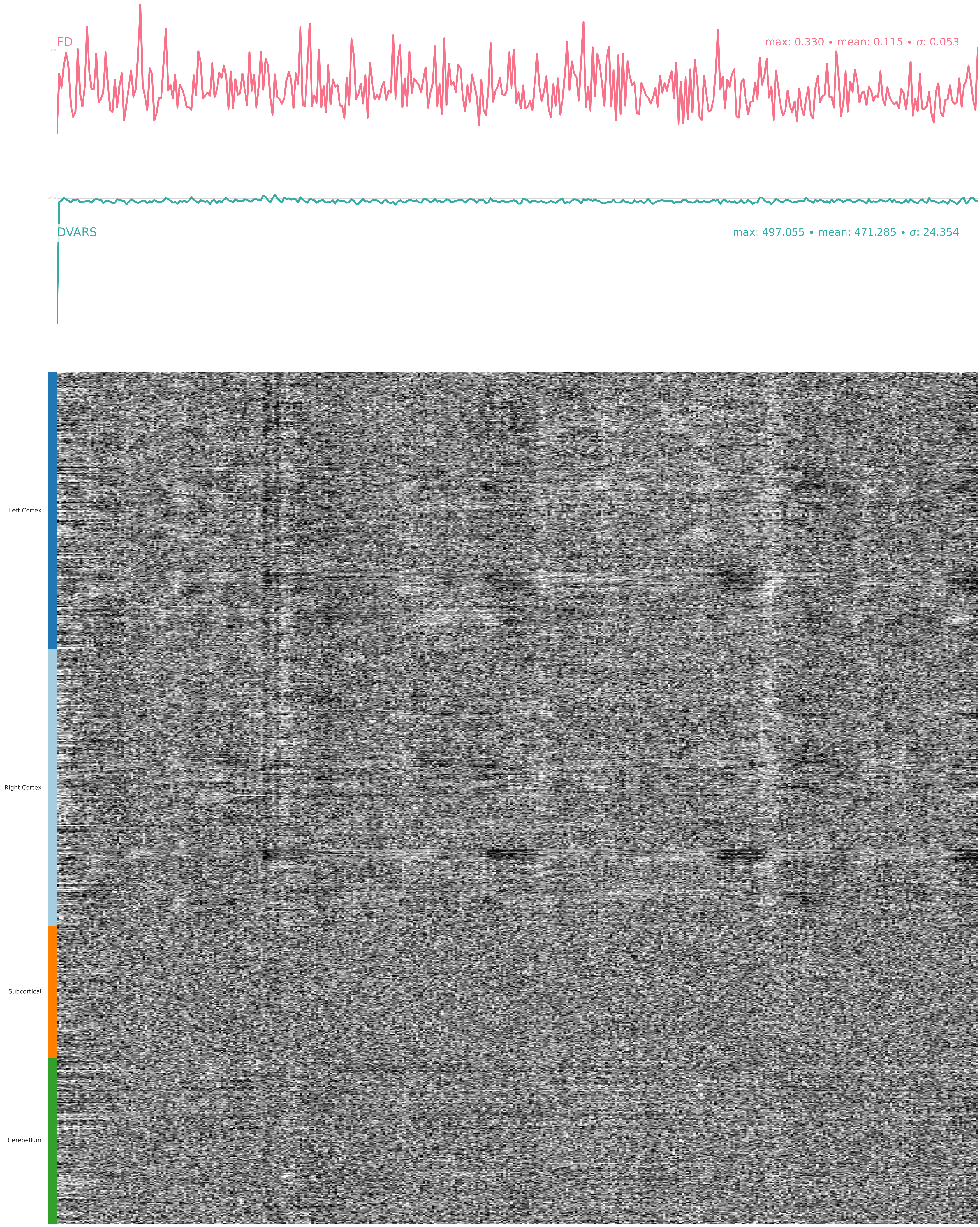


Get figure file: sub-113922/figures/sub-113922_task-WM_dir-LR_space-fsLR_desc-preprocessing_bold.svg

#### Framewise Displacement and Censored Volumes

Framewise displacement (FD) is used to flag high-motion volumes, which are then censored as part of the denoising procedure. If motion filtering is requested, then the six translation and rotation motion parameters are filtered to remove respiratory effects before FD is calculated and outlier volumes are identified.

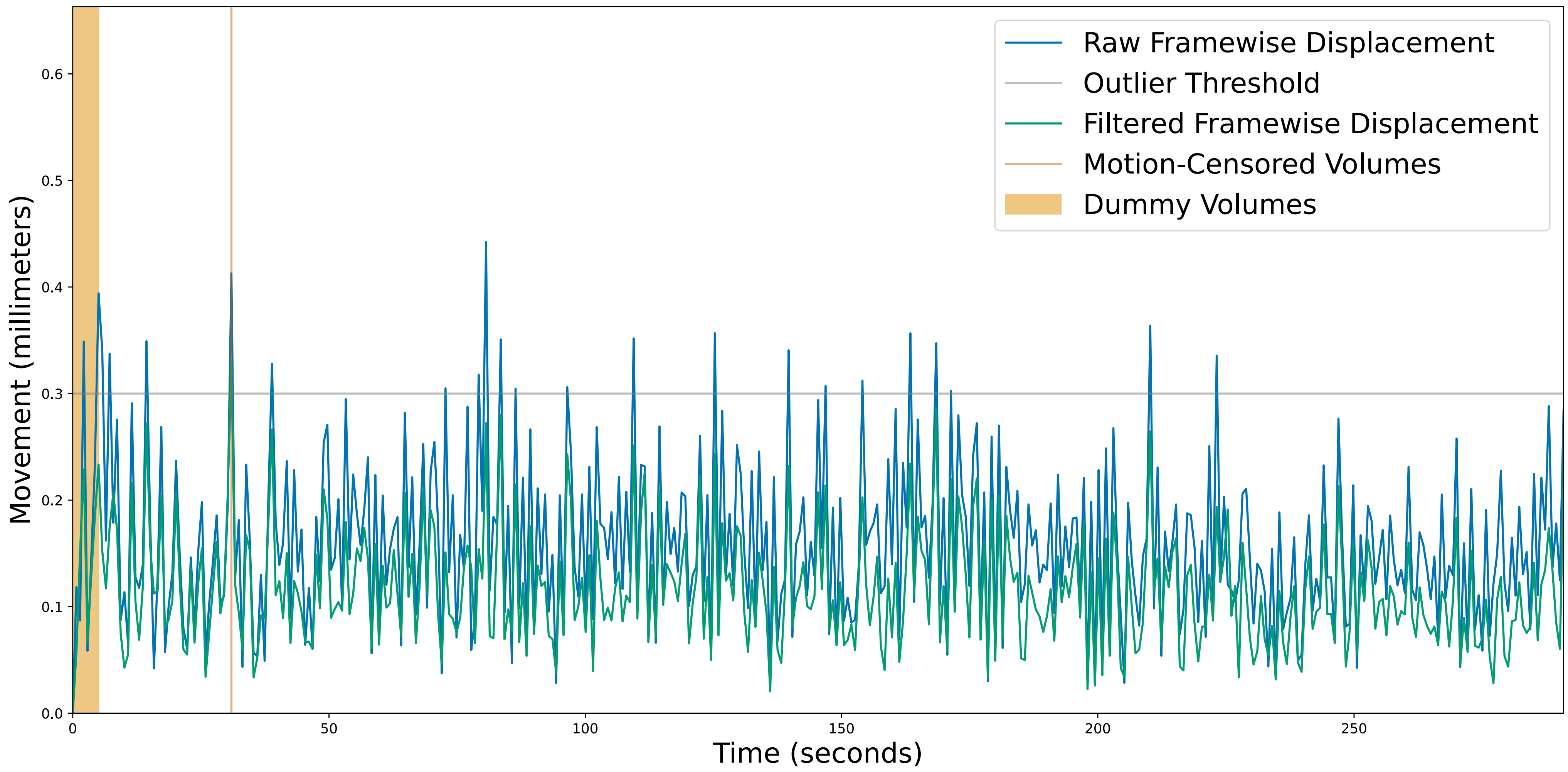


Get figure file: sub-113922/figures/sub-113922_task-WM_dir-LR_space-fsLR_desc-censoring_motion.svg

#### Design Matrix for Confound Regression

The "design matrix" represents the confounds that are used to denoise the BOLD data.

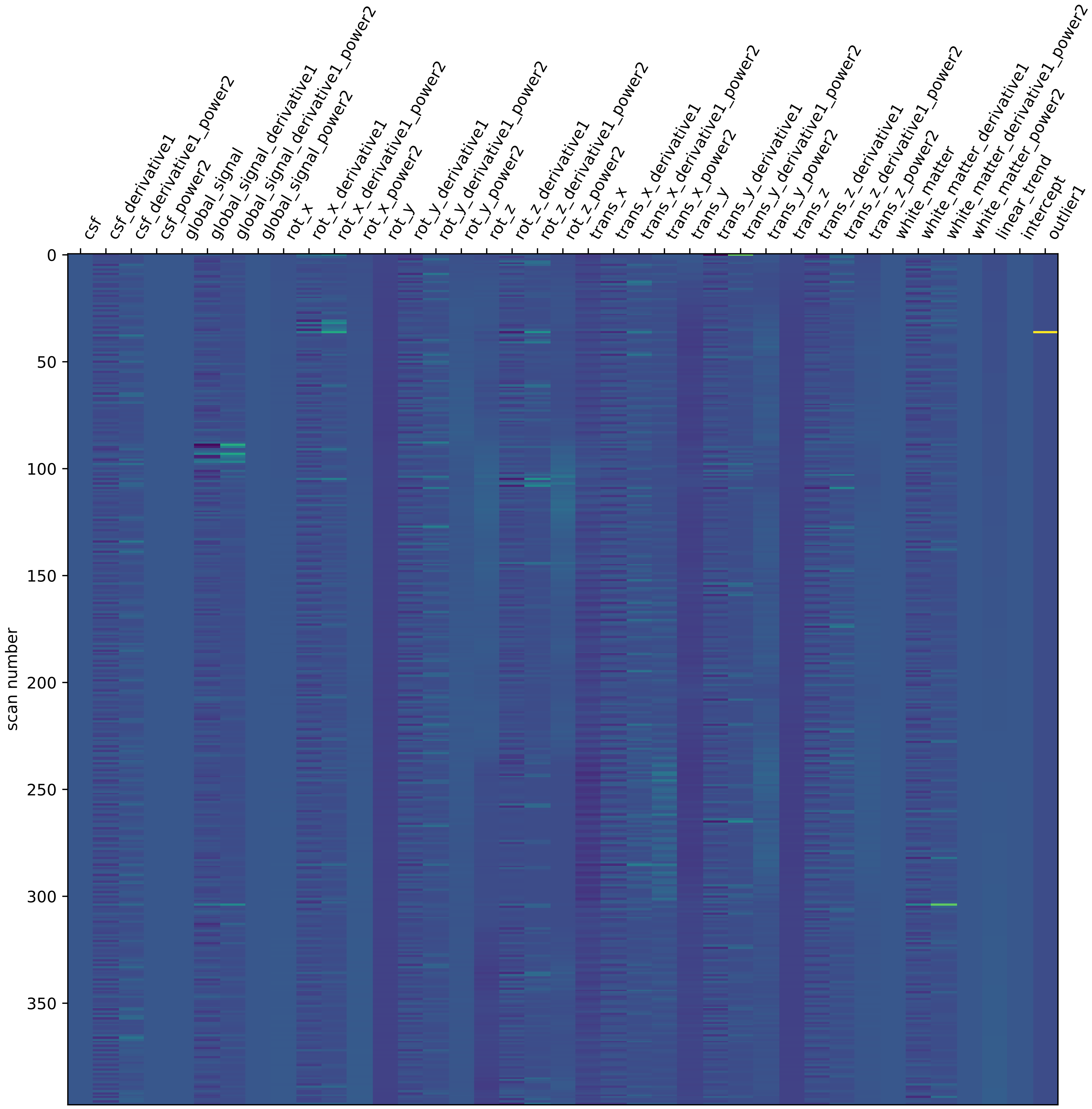


Get figure file: sub-113922/figures/sub-113922_task-WM_dir-LR_design.svg

#### Carpet Plot After Postprocessing

FD and DVARS are two measures of in-scanner motion. This plot shows standardized FD, DVARS, and then a carpet plot for the time series of each voxel/vertex's time series of activity.

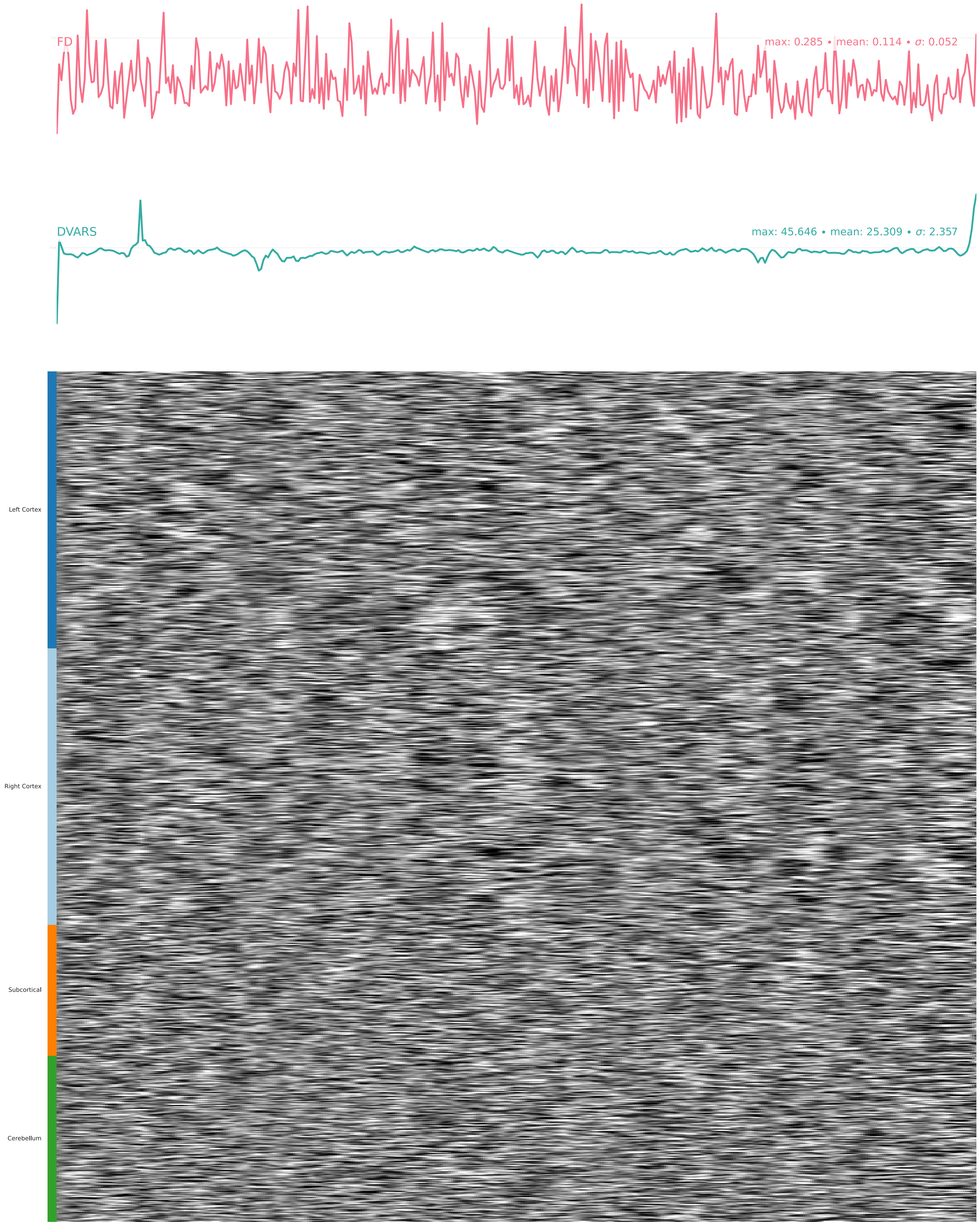


Get figure file: sub-113922/figures/sub-113922_task-WM_dir-LR_space-fsLR_desc-postprocessing_bold.svg

#### Correlation Heatmaps from Four Atlases

This plot shows heatmaps from ROI-to-ROI correlations from four atlases.

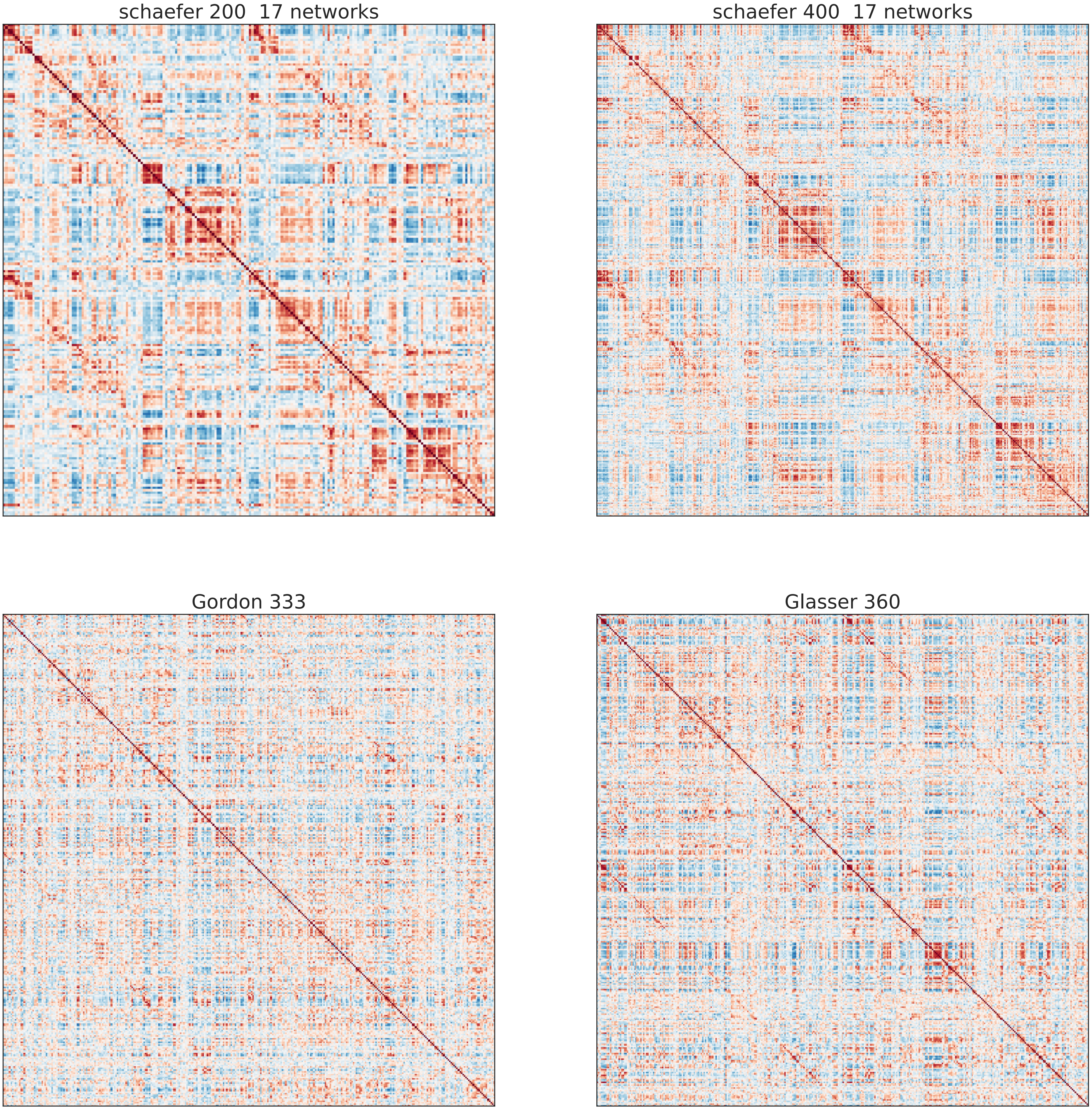


Get figure file: sub-113922/figures/sub-113922_task-WM_dir-LR_space-fsLR_desc-connectivityplot_bold.svg

### Reports for: task WM, direction RL.

#### Summary

- BOLD volume space: fsLR
- Repetition Time (TR): 0.72
- Mean Framewise Displacement: 0.1295
- Mean Relative RMS Motion: 0.7345
- Max Relative RMS Motion: 1.0183
- DVARS Before and After Processing : 469.1356, 25.6721
- Correlation between DVARS and FD Before and After Processing : 0.1558, 0.0943
- Number of Volumes Censored : 4

#### Alignment of functional and anatomical MRI data (surface driven)

bbregister was used to coregister functional and anatomical MRI data.

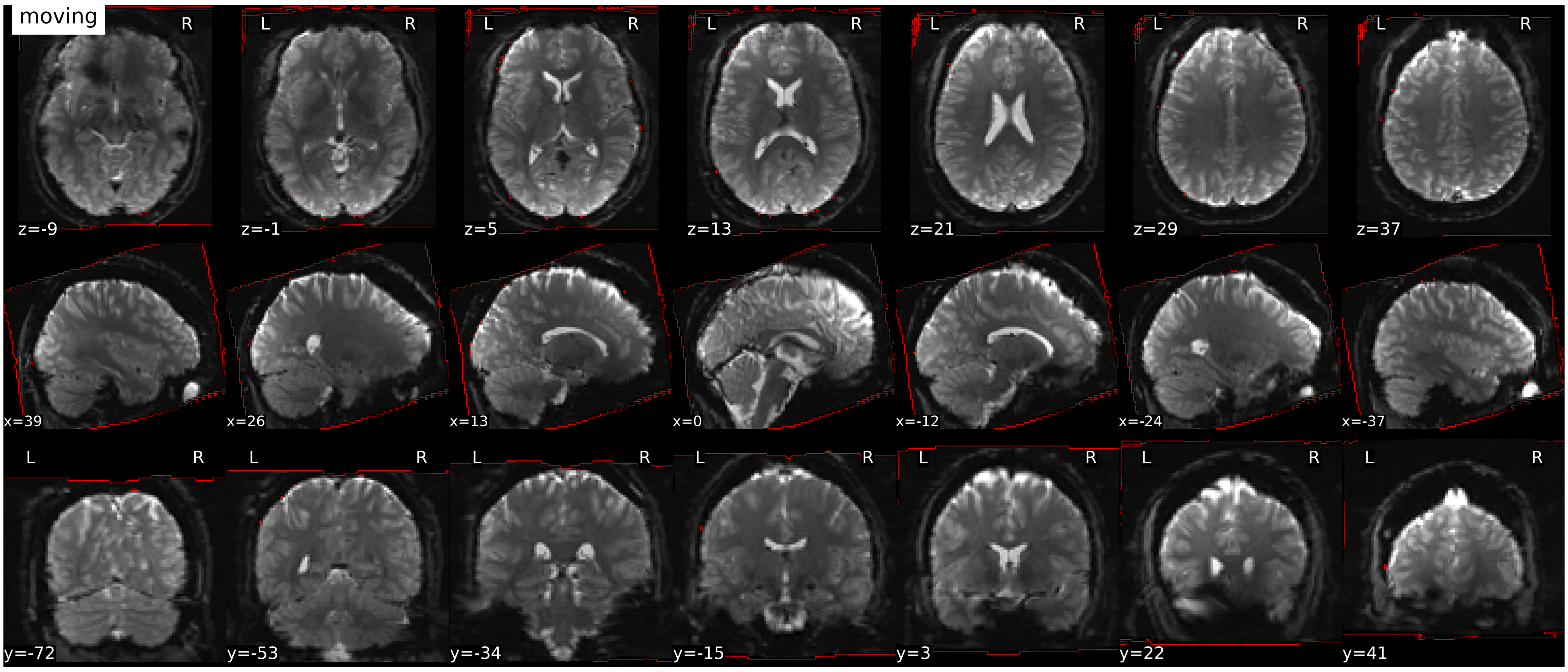


Get figure file: sub-113922/figures/sub-113922_task-WM_dir-RL_space-MNI152NLin6Asym_desc-bbregister_bold.svg

#### Carpet Plot Before Postprocessing

FD and DVARS are two measures of in-scanner motion. This plot shows standardized FD, DVARS, and then a carpet plot for the time series of each voxel/vertex's time series of activity.

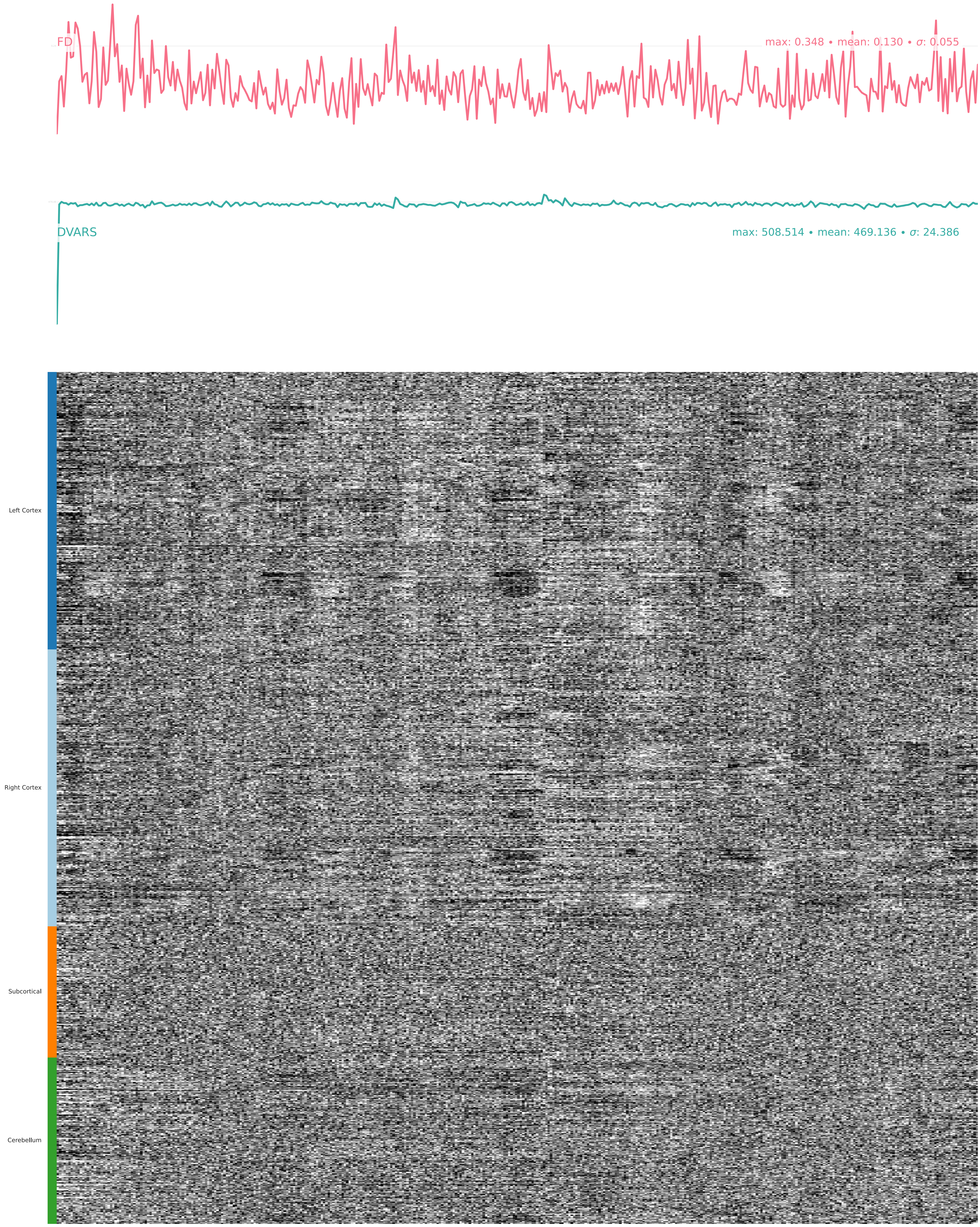


Get figure file: sub-113922/figures/sub-113922_task-WM_dir-RL_space-fsLR_desc-preprocessing_bold.svg

#### Framewise Displacement and Censored Volumes

Framewise displacement (FD) is used to flag high-motion volumes, which are then censored as part of the denoising procedure. If motion filtering is requested, then the six translation and rotation motion parameters are filtered to remove respiratory effects before FD is calculated and outlier volumes are identified.

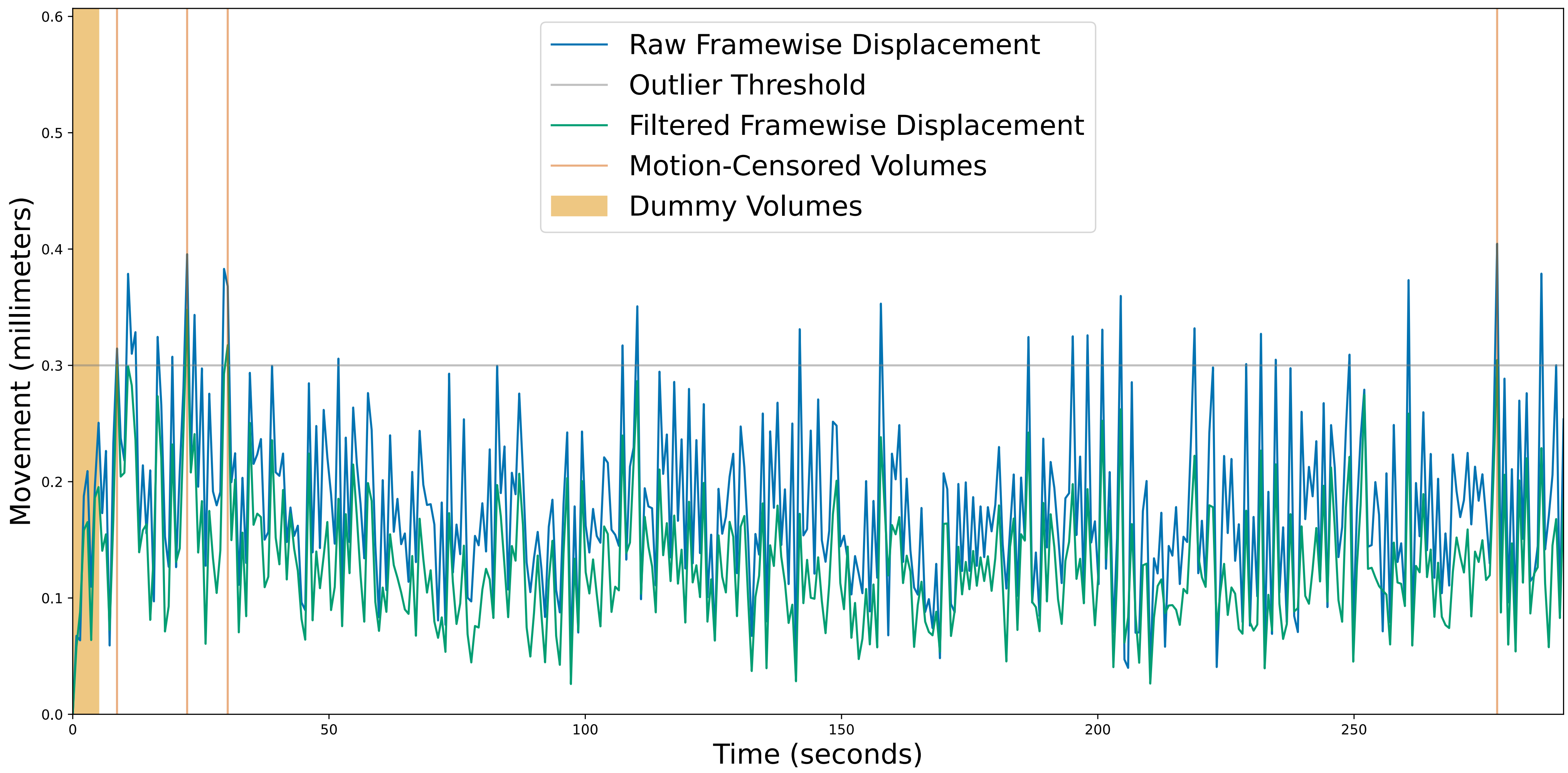


Get figure file: sub-113922/figures/sub-113922_task-WM_dir-RL_space-fsLR_desc-censoring_motion.svg

#### Design Matrix for Confound Regression

The "design matrix" represents the confounds that are used to denoise the BOLD data.

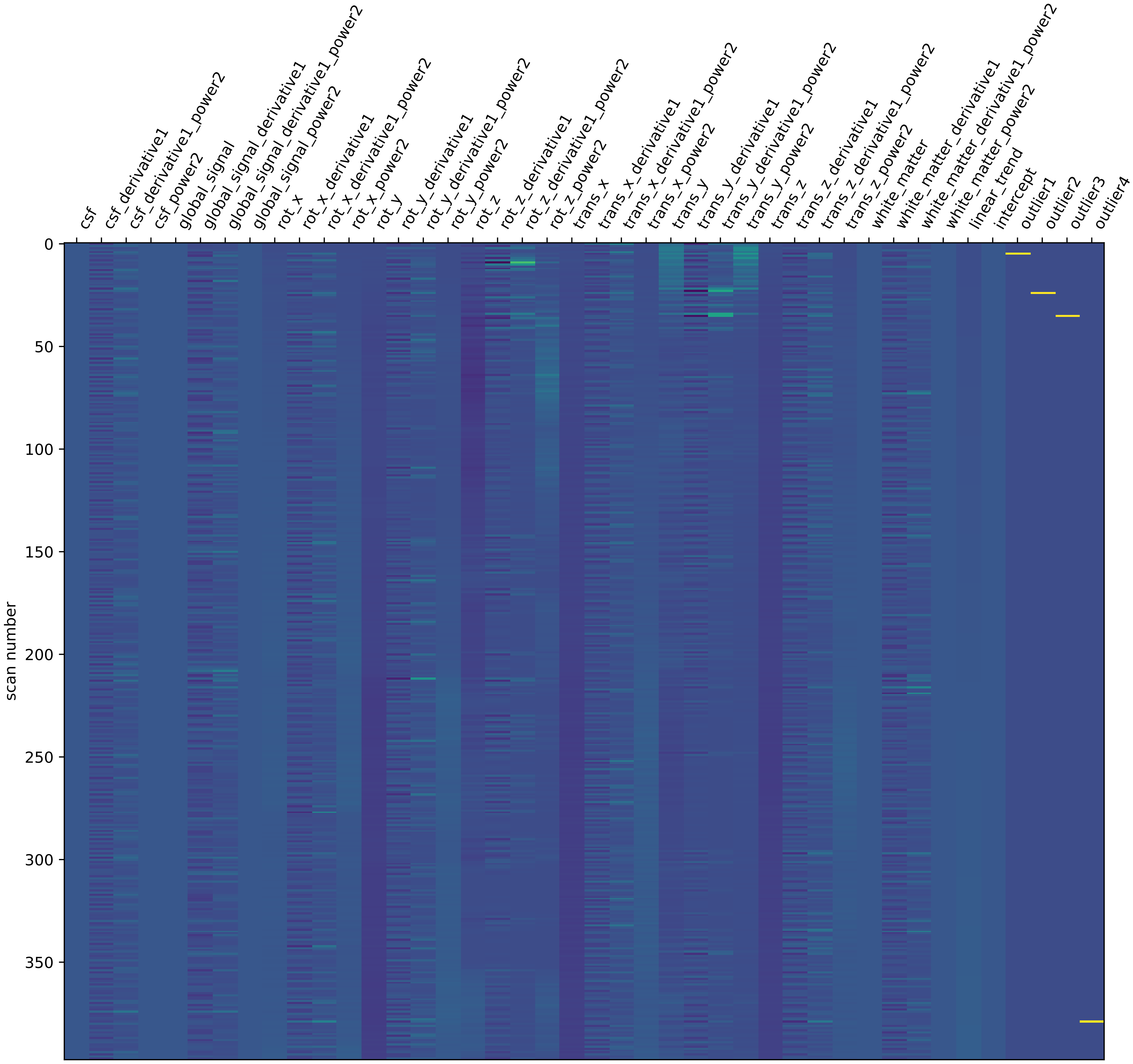


Get figure file: sub-113922/figures/sub-113922_task-WM_dir-RL_design.svg

#### Carpet Plot After Postprocessing

FD and DVARS are two measures of in-scanner motion. This plot shows standardized FD, DVARS, and then a carpet plot for the time series of each voxel/vertex's time series of activity.

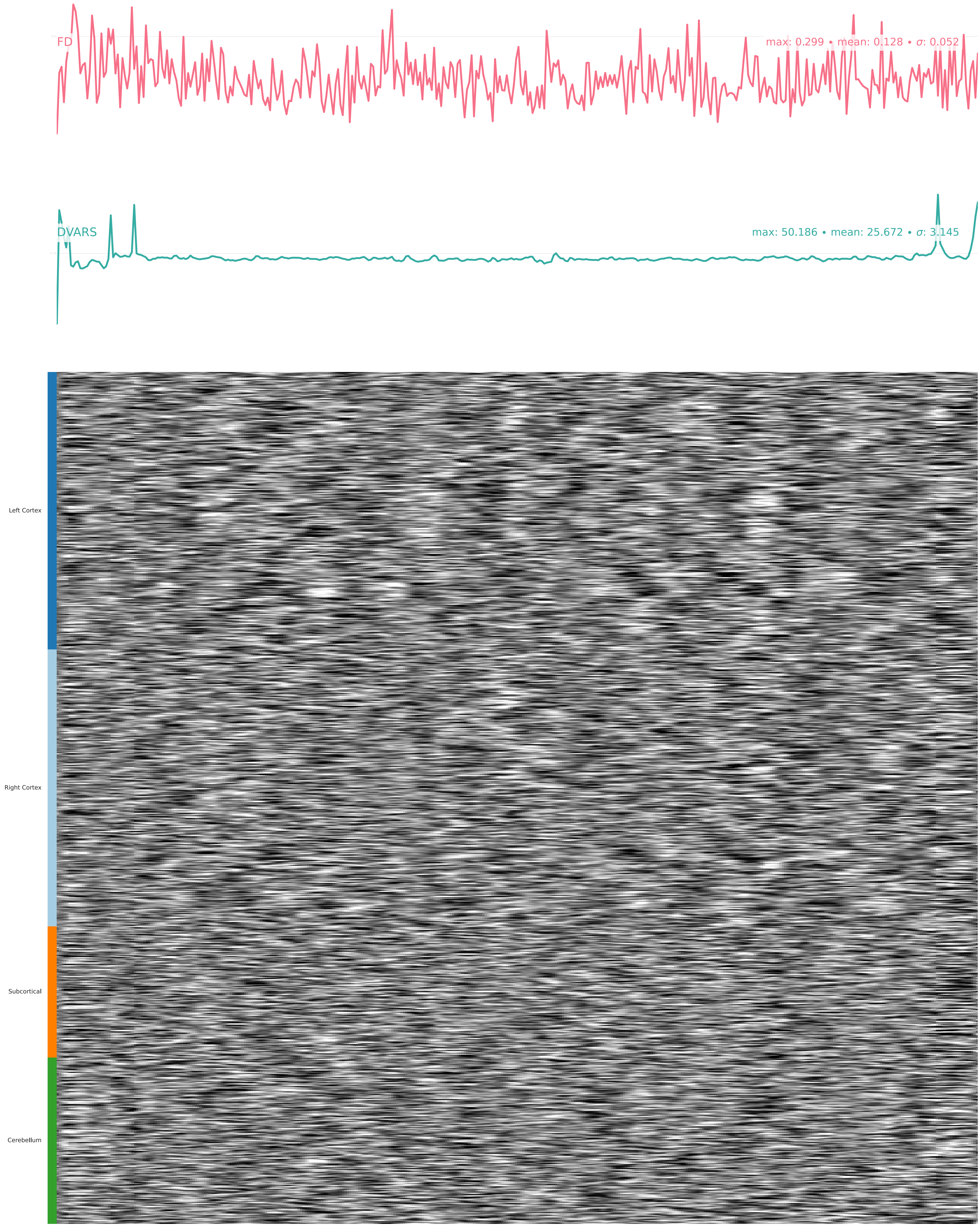


Get figure file: sub-113922/figures/sub-113922_task-WM_dir-RL_space-fsLR_desc-postprocessing_bold.svg

#### Correlation Heatmaps from Four Atlases

This plot shows heatmaps from ROI-to-ROI correlations from four atlases.

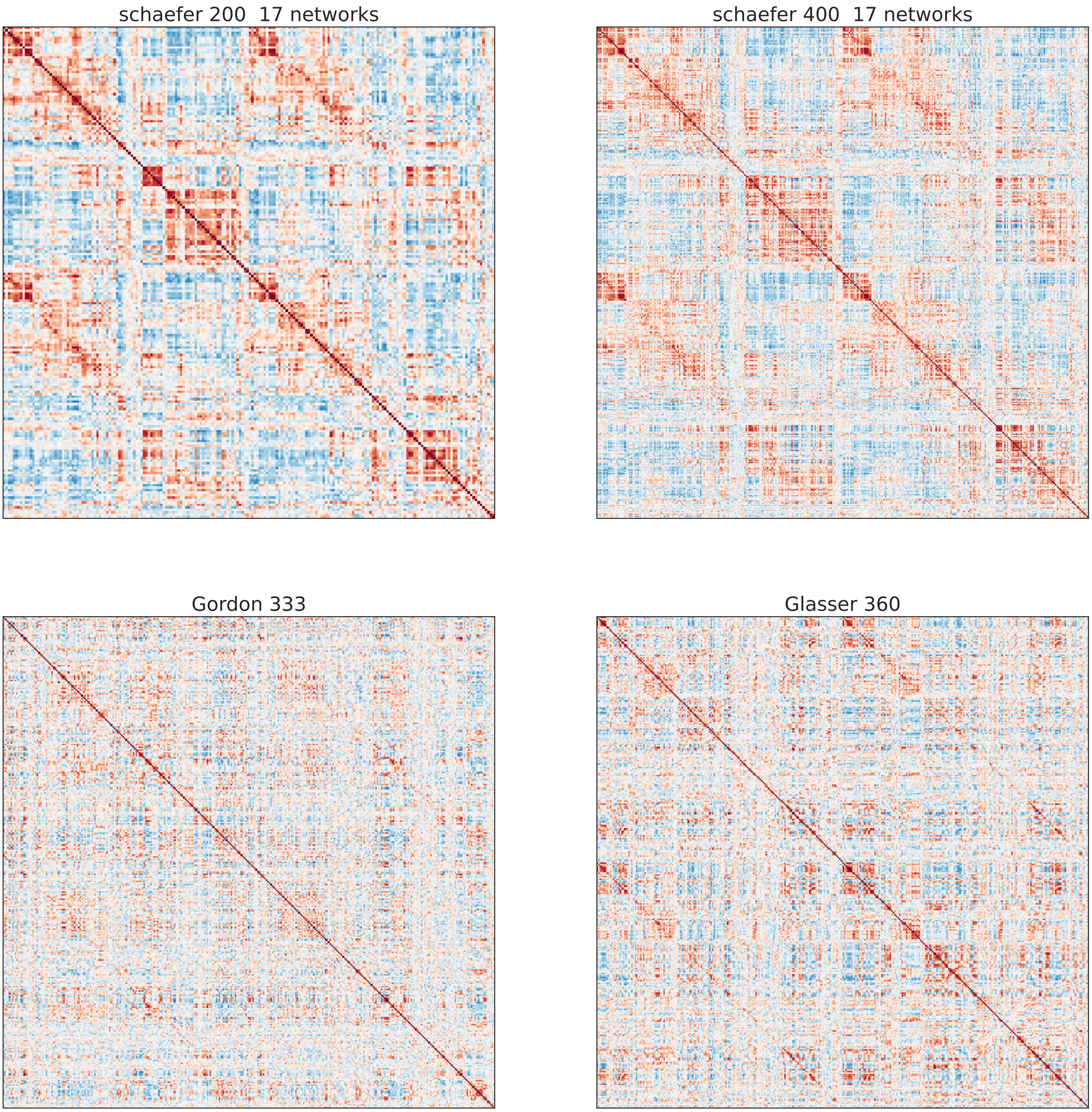


Get figure file: sub-113922/figures/sub-113922_task-WM_dir-RL_space-fsLR_desc-connectivityplot_bold.svg

### About

- xcp_d version: 0.5.0rc2
- xcp_d: `/usr/local/miniconda/bin/xcp_d inputs/data xcp participant --combineruns --nthreads 1 --omp-nthreads 1 --mem_gb 10 --smoothing 2 --min_coverage 0.5 --min_time 100 --dummy-scans 7 --random-seed 0 --bpf-order 2 --motion-filter-type notch --band-stop-min 12 --band-stop-max 18 --motion-filter-order 4 --head-radius auto --exact-time 300 480 600 --despike --lower-bpf 0.01 --upper-bpf 0.08 --participant_label 113922 -p 36P -f 0.3 --cifti --warp-surfaces-native2std --dcan-qc -w /scratch/mehtaka/SGE_8728772/job-8728772-113922/ds/.git/tmp/wkdir -v --input-type hcp`
- xcp_d preprocessed: 2023-09-18 02:14:48 -0400

### Methods

We kindly ask to report results preprocessed with this tool using the following boilerplate.

HTML Markdown LaTeX

### Post-processing of hcp outputs

The eXtensible Connectivity Pipeline- DCAN (XCP-D) (Ciric et al. 2018; Satterthwaite et al. 2013) was used to post-process the outputs of *HCP* version unknown (Glasser et al. 2013). XCP-D was built with *Nipype* version 1.8.6 (Gorgolewski et al. 2011, RRID:SCR_002502). For each of the eighteen BOLD runs found per subject (across all tasks and sessions), the following post-processing was performed. The first seven volumes of both the BOLD data and nuisance regressors were discarded as non-steady-state volumes, or 'dummy scans'. In order to identify high-motion outlier volumes, the six translation and rotation head motion traces were band-stop filtered to remove signals between 12.0 and 18.0 breaths-per-minute using a(n) fourth-order notch filter, based on Fair et al. (2020). Next, framewise displacement was calculated using the formula from Power et al. (2014), with a head radius of 76.0688323581881 mm. Volumes with filtered framewise displacement greater than 0.3 mm were flagged as high-motion outliers for the sake of later censoring (Power et al. 2014). In total, 36 nuisance regressors were selected from the preprocessing confounds, according to the '36P' strategy. These nuisance regressors included six filtered motion parameters, mean global signal, mean white matter signal, mean cerebrospinal fluid signal with their temporal derivatives, and quadratic expansion of six motion parameters, tissue signals and their temporal derivatives (Ciric et al. 2017; Satterthwaite et al. 2013). Finally, linear trend and intercept terms were added to the regressors prior to denoising. The BOLD data were converted to NIfTI format, despiked with *AFNI*'s *3dDespike*, and converted back to CIFTI format. Nuisance regressors were regressed from the BOLD data using linear regression, as implemented in *Nilearn*. Any volumes censored earlier in the workflow were then interpolated in the residual time series produced by the regression. The interpolated timeseries were then band-pass filtered using a(n) second-order Butterworth filter, in order to retain signals between 0.01-0.08 Hz. The filtered, interpolated time series were then re-censored to remove high-motion outlier volumes. The denoised BOLD was then smoothed using *Connectome Workbench* with a Gaussian kernel (FWHM=2.0 mm). The amplitude of low-frequency fluctuation (ALFF) (Zou et al. 2008) was computed by transforming the processed BOLD timeseries to the frequency domain. The power spectrum was computed within the 0.01-0.08 Hz frequency band and the mean square root of the power spectrum was calculated at each voxel to yield voxel-wise ALFF measures. The ALFF maps were smoothed with the Connectome Workbench using a Gaussian kernel (FWHM=2.0 mm).

For each hemisphere, regional homogeneity (ReHo) (Jiang and Zuo 2016) was computed using surface-based *2dReHo* (Zhang et al. 2019). Specifically, for each vertex on the surface, the Kendall's coefficient of concordance (KCC) was computed with nearest-neighbor vertices to yield ReHo. For the subcortical, volumetric data, ReHo was computed with neighborhood voxels using *AFNI*'s *3dReHo* (Taylor and Saad 2013).

Processed functional timeseries were extracted from residual BOLD using Connectome Workbench (Glasser et al. 2013) for the following atlases: the Schaefer 17-network 100, 200, 300, 400, 500, 600, 700, 800, 900, and 1000 parcel atlas (Schaefer et al. 2018), the Glasser atlas (Glasser et al. 2016), the Gordon atlas (Gordon et al. 2016), and the Tian subcortical atlas (Tian et al. 2020). Corresponding pair-wise functional connectivity between all regions was computed for each atlas, which was operationalized as the Pearson's correlation of each parcel's unsmoothed timeseries with the Connectome Workbench. In cases of partial coverage, uncovered vertices (values of all zeros or NaNs) were either ignored (when the parcel had >50.0% coverage) or were set to zero (when the parcel had <50.0% coverage). For each of the eighteen BOLD runs found per subject (across all tasks and sessions), the following post-processing was performed. The first seven volumes of both the BOLD data and nuisance regressors were discarded as non-steady-state volumes, or 'dummy scans'. In order to identify high-motion outlier volumes, the six translation and rotation head motion traces were band-stop filtered to remove signals between 12.0 and 18.0 breaths-per-minute using a(n) fourth-order notch filter, based on Fair et al. (2020). Next, framewise displacement was calculated using the formula from Power et al. (2014), with a head radius of 76.0688323581881 mm. Volumes with filtered framewise displacement greater than 0.3 mm were flagged as high-motion outliers for the sake of later censoring (Power et al. 2014). Additional sets of censoring volumes were randomly selected to produce additional correlation matrices limited to 416, 666, and 833 volumes. In total, 36 nuisance regressors were selected from the preprocessing confounds, according to the '36P' strategy. These nuisance regressors included six filtered motion parameters, mean global signal, mean white matter signal, mean cerebrospinal fluid signal with their temporal derivatives, and quadratic expansion of six motion parameters, tissue signals and their temporal derivatives (Ciric et al. 2017; Satterthwaite et al. 2013). Finally, linear trend and intercept terms were added to the regressors prior to denoising. The BOLD data were converted to NIfTI format, despiked with *AFNI*'s *3dDespike*, and converted back to CIFTI format. Nuisance regressors were regressed from the BOLD data using linear regression, as implemented in *Nilearn*. Any volumes censored earlier in the workflow were then interpolated in the residual time series produced by the regression. The interpolated timeseries were then band-pass filtered using a(n) second-order Butterworth filter, in order to retain signals between 0.01-0.08 Hz. The filtered, interpolated time series were then re-censored to remove high-motion outlier volumes. The denoised BOLD was then smoothed using *Connectome Workbench* with a Gaussian kernel (FWHM=2.0 mm). The amplitude of low-frequency fluctuation (ALFF) (Zou et al. 2008) was computed by transforming the processed BOLD timeseries to the frequency domain. The power spectrum was computed within the 0.01-0.08 Hz frequency band and the mean square root of the power spectrum was calculated at each voxel to yield voxel-wise ALFF measures. The ALFF maps were smoothed with the Connectome Workbench using a Gaussian kernel (FWHM=2.0 mm).

For each hemisphere, regional homogeneity (ReHo) (Jiang and Zuo 2016) was computed using surface-based *2dReHo* (Zhang et al. 2019). Specifically, for each vertex on the surface, the Kendall's coefficient of concordance (KCC) was computed with nearest-neighbor vertices to yield ReHo. For the subcortical, volumetric data, ReHo was computed with neighborhood voxels using *AFNI*'s *3dReHo* (Taylor and Saad 2013).

Processed functional timeseries were extracted from residual BOLD using Connectome Workbench (Glasser et al. 2013) for the following atlases: the Schaefer 17-network 100, 200, 300, 400, 500, 600, 700, 800, 900, and 1000 parcel atlas (Schaefer et al. 2018), the Glasser atlas (Glasser et al. 2016), the Gordon atlas (Gordon et al. 2016), and the Tian subcortical atlas (Tian et al. 2020). Corresponding pair-wise functional connectivity between all regions was computed for each atlas, which was operationalized as the Pearson's correlation of each parcel's unsmoothed timeseries with the Connectome Workbench. In cases of partial coverage, uncovered vertices (values of all zeros or NaNs) were either ignored (when the parcel had >50.0% coverage) or were set to zero (when the parcel had <50.0% coverage).

Many internal operations of *XCP-D* use *AFNI* (Cox 1996; Cox and Hyde 1997), *Connectome Workbench* (Marcus et al. 2011), *ANTS* (Avants et al. 2009), *TemplateFlow* version 0.8.1 (Ciric et al. 2022), *matplotlib* version 3.4.3 (Hunter 2007), *Nibabel* version 5.0.1 (Brett et al. 2022), *Nilearn* version 0.10.1 (Abraham et al. 2014), *numpy* version 1.22.4 (Harris et al. 2020), *pybids* version 0.15.5 (Yarkoni et al. 2019), and *scipy* version 1.10.1 (Virtanen et al. 2020). For more details, see the *XCP-D* website (https://xcp-d.readthedocs.io).

#### Copyright Waiver

The above methods description text was automatically generated by *XCP-D* with the express intention that users should copy and paste this text into their manuscripts *unchanged*. It is released under the CC0 license.

#### References

Abraham, Alexandre, Fabian Pedregosa, Michael Eickenberg, Philippe Gervais, Andreas Mueller, Jean Kossaifi, Alexandre Gramfort, Bertrand Thirion, and Gaël Varoquaux. 2014. "Machine Learning for Neuroimaging with Scikit-Learn." *Frontiers in Neuroinformatics*. Frontiers, 14.

Avants, Brian B, Nick Tustison, Gang Song, and others. 2009. "Advanced Normalization Tools (Ants)." *Insight J* 2 (365): 1–35.

Brett, Matthew, Christopher J. Markiewicz, Michael Hanke, Marc-Alexandre Côté, Ben Cipollini, Paul McCarthy, Dorota Jarecka, et al. 2022. *Nipy/Nibabel:* (version 4.0.0). Zenodo. https://doi.org/10.5281/zenodo.591597.

Ciric, Rastko, Adon F. G. Rosen, Guray Erus, Matthew Cieslak, Azeez Adebimpe, Philip A. Cook, Danielle S. Bassett, Christos Davatzikos, Daniel H. Wolf, and Theodore D. Satterthwaite. 2018. "Mitigating Head Motion Artifact in Functional Connectivity MRI." *Nature Protocols* 13 (12): 2801–26. https://doi.org/10.1038/s41596-018-0065-y.

Ciric, Rastko, William H Thompson, Romy Lorenz, Mathias Goncalves, Eilidh MacNicol, Christopher J Markiewicz, Yaroslav O Halchenko, et al. 2022. "TemplateFlow: FAIR-Sharing of Multi-Scale, Multi-Species Brain Models." *bioRxiv*. Cold Spring Harbor Laboratory, 2021–02. https://doi.org/10.1101/2021.02.10.430678.

Ciric, Rastko, Daniel H. Wolf, Jonathan D. Power, David R. Roalf, Graham Baum, Kosha Ruparel, Russell T. Shinohara, et al. 2017. "Benchmarking of Participant-Level Confound Regression Strategies for the Control of Motion Artifact in Studies of Functional Connectivity." *NeuroImage* 154 (July): 174–87. https://doi.org/10.1016/j.neuroimage.2017.03.020.

Cox, Robert W. 1996. "AFNI: Software for Analysis and Visualization of Functional Magnetic Resonance Neuroimages." *Computers and Biomedical Research* 29 (3). Elsevier: 162–73.

Cox, Robert W, and James S Hyde. 1997. "Software Tools for Analysis and Visualization of fMRI Data." *NMR in Biomedicine: An International Journal Devoted to the Development and Application of Magnetic Resonance in Vivo* 10 (4-5). Wiley Online Library: 171–78.

Fair, Damien A, Oscar Miranda-Dominguez, Abraham Z Snyder, Anders Perrone, Eric A Earl, Andrew N Van, Jonathan M Koller, et al. 2020. "Correction of Respiratory Artifacts in Mri Head Motion Estimates." *Neuroimage* 208. Elsevier: 116400. https://doi.org/10.1016/j.neuroimage.2019.116400.

Glasser, Matthew F., Timothy S. Coalson, Emma C. Robinson, Carl D. Hacker, John Harwell, Essa Yacoub, Kamil Ugurbil, et al. 2016. "A Multi-Modal Parcellation of Human Cerebral Cortex." *Nature* 536 (7615): 171–78. https://doi.org/10.1038/nature18933.

Glasser, Matthew F., Stamatios N. Sotiropoulos, J. Anthony Wilson, Timothy S. Coalson, Bruce Fischl, Jesper L. Andersson, Junqian Xu, et al. 2013. "The Minimal Preprocessing Pipelines for the Human Connectome Project." *NeuroImage* 80 (October): 105–24. https://doi.org/10.1016/j.neuroimage.2013.04.127.

Gordon, Evan M., Timothy O. Laumann, Babatunde Adeyemo, Jeremy F. Huckins, William M. Kelley, and Steven E. Petersen. 2016. "Generation and Evaluation of a Cortical Area Parcellation from Resting-State Correlations." *Cerebral Cortex* 26 (1): 288–303. https://doi.org/10.1093/cercor/bhu239.

Gorgolewski, Krzysztof, Christopher D. Burns, Cindee Madison, Dav Clark, Yaroslav O. Halchenko, Michael L. Waskom, and Satrajit S. Ghosh. 2011. "Nipype: A Flexible, Lightweight and Extensible Neuroimaging Data Processing Framework in Python." *Frontiers in Neuroinformatics* 5. https://doi.org/10.3389/fninf.2011.00013.

Harris, Charles R., Jarrod K. Millman, Stéfan J. van der Walt, Ralf Gommers, Pauli Virtanen, David Cournapeau, Eric Wieser, et al. 2020. "Array Programming with NumPy." *Nature* 585 (7825): 357–62. https://doi.org/10.1038/s41586-020-2649-2.

Hunter, John D. 2007. "Matplotlib: A 2D Graphics Environment." *Computing in Science & Engineering* 9 (03). IEEE Computer Society: 90–95.

Jiang, Lili, and Xi-Nian Zuo. 2016. "Regional Homogeneity: A Multimodal, Multiscale Neuroimaging Marker of the Human Connectome." *The Neuroscientist* 22 (5). Sage Publications Sage CA: Los Angeles, CA: 486–505.

Marcus, Daniel S, John Harwell, Timothy Olsen, Michael Hodge, Matthew F Glasser, Fred Prior, Mark Jenkinson, Timothy Laumann, Sandra W Curtiss, and David C Van Essen. 2011. "Informatics and Data Mining Tools and Strategies for the Human Connectome Project." *Frontiers in Neuroinformatics* 5. Frontiers Research Foundation: 4.

Power, Jonathan D., Anish Mitra, Timothy O. Laumann, Abraham Z. Snyder, Bradley L. Schlaggar, and Steven E. Petersen. 2014. "Methods to Detect, Characterize, and Remove Motion Artifact in Resting State fMRI." *NeuroImage* 84 (January): 320–41. https://doi.org/10.1016/j.neuroimage.2013.08.048.

Satterthwaite, Theodore D., Mark A. Elliott, Raphael T. Gerraty, Kosha Ruparel, James Loughead, Monica E. Calkins, Simon B. Eickhoff, et al. 2013. "An Improved Framework for Confound Regression and Filtering for Control of Motion Artifact in the Preprocessing of Resting-State Functional Connectivity Data." *NeuroImage* 64 (January): 240–56. https://doi.org/10.1016/j.neuroimage.2012.08.052.

Schaefer, Alexander, Ru Kong, Evan M. Gordon, Timothy O. Laumann, Xi-Nian Zuo, Avram J. Holmes, Simon B. Eickhoff, and B. T. Thomas Yeo. 2018. "Local-Global Parcellation of the Human Cerebral Cortex from Intrinsic Functional Connectivity MRI." *Cerebral Cortex (New York, N.Y.: 1991)* 28 (9): 3095–3114. https://doi.org/10.1093/cercor/bhx179.

Taylor, Paul A, and Ziad S Saad. 2013. "FATCAT:(An Efficient) Functional and Tractographic Connectivity Analysis Toolbox." *Brain Connectivity* 3 (5). Mary Ann Liebert, Inc. 140 Huguenot Street, 3rd Floor New Rochelle, NY 10801 USA: 523–35.

Tian, Ye, Daniel S Margulies, Michael Breakspear, and Andrew Zalesky. 2020. "Topographic Organization of the Human Subcortex Unveiled with Functional Connectivity Gradients." *Nature Neuroscience* 23 (11). Nature Publishing Group: 1421–32. https://doi.org/10.1038/s41593-020-00711-6.

Virtanen, Pauli, Ralf Gommers, Travis E. Oliphant, Matt Haberland, Tyler Reddy, David Cournapeau, Evgeni Burovski, et al. 2020. "SciPy 1.0: Fundamental Algorithms for Scientific Computing in Python." *Nature Methods* 17 (3): 261–72. https://doi.org/10.1038/s41592-019-0686-2.

Yarkoni, Tal, Christopher J Markiewicz, Alejandro de la Vega, Krzysztof J Gorgolewski, Taylor Salo, Yaroslav O Halchenko, Quinten McNamara, et al. 2019. "PyBIDS: Python Tools for Bids Datasets." *Journal of Open Source Software* 4 (40). NIH Public Access.

Zhang, Bo, Fei Wang, Hao-Ming Dong, Xiao-Wei Jiang, Sheng-Nan Wei, Miao Chang, Zhi-Yang Yin, et al. 2019. "Surface-Based Regional Homogeneity in Bipolar Disorder: A Resting-State fMRI Study." *Psychiatry Research* 278 (August): 199–204. https://doi.org/10.1016/j.psychres.2019.05.045.

Zou, Qi-Hong, Chao-Zhe Zhu, Yihong Yang, Xi-Nian Zuo, Xiang-Yu Long, Qing-Jiu Cao, Yu-Feng Wang, and Yu-Feng Zang. 2008. "An Improved Approach to Detection of Amplitude of Low-Frequency Fluctuation (ALFF) for Resting-State fMRI: Fractional ALFF." *Journal of Neuroscience Methods* 172 (1): 137–41. https://doi.org/10.1016/j.jneumeth.2008.04.012.

### Errors

No errors to report!
