## Supplemental Figure 3 for "XCP-D: A Robust Pipeline for the post-processing of fMRI data"

```
xcp_d
├──dataset_description.json
├──desc-linc_qc.json
├──sub-20712.html
├──space-MNI152NLin6Asym_atlas-Glasser_res-2_dseg.nii.gz
├──space-MNI152NLin6Asym_atlas-Gordon_res-2_dseg.nii.gz
├──space-MNI152NLin6Asym_atlas-HCP_res-2_dseg.nii.gz
├──space-MNI152NLin6Asym_atlas-Schaefer1017_res-2_dseg.nii.gz
├──space-MNI152NLin6Asym_atlas-Schaefer117_res-2_dseg.nii.gz
├──space-MNI152NLin6Asym_atlas-Schaefer217_res-2_dseg.nii.gz
├──space-MNI152NLin6Asym_atlas-Schaefer317_res-2_dseg.nii.gz
├──space-MNI152NLin6Asym_atlas-Schaefer417_res-2_dseg.nii.gz
├──space-MNI152NLin6Asym_atlas-Schaefer517_res-2_dseg.nii.gz
├──space-MNI152NLin6Asym_atlas-Schaefer617_res-2_dseg.nii.gz
├──space-MNI152NLin6Asym_atlas-Schaefer717_res-2_dseg.nii.gz
├──space-MNI152NLin6Asym_atlas-Schaefer817_res-2_dseg.nii.gz
├──space-MNI152NLin6Asym_atlas-Schaefer917_res-2_dseg.nii.gz
├──space-MNI152NLin6Asym_atlas-Tian_res-2_dseg.nii.gz
├──sub-20712_ses-12367_executive_summary.html
├──logs
       ├──CITATION.bib
       ├──CITATION.html
       ├──CITATION.md
       └──CITATION.tex
└──sub-20712
       └──ses-12367
       ├──anat
              ├──sub-20712_ses-12367_rec-defaced_space-MNI152NLin6Asym_desc-preproc_T1w.nii.g
              z
              └──sub-20712_ses-12367_rec-defaced_space-MNI152NLin6Asym_dseg.nii.gz
       └──func
              ├──sub-20712_ses-12367_task-restbold_run-1_desc-preproc_design.tsv
              ├──sub-20712_ses-12367_task-restbold_run-1_motion.json
              ├──sub-20712_ses-12367_task-restbold_run-1_motion.tsv
              ├──sub-20712_ses-12367_task-restbold_run-1_outliers.json
              ├──sub-20712_ses-12367_task-restbold_run-1_outliers.tsv
              ├──sub-20712_ses-12367_task-restbold_run-1_space-MNI152NLin6Asym_atlas-Glasser_
              coverage.tsv
              ├──sub-20712_ses-12367_task-restbold_run-1_space-MNI152NLin6Asym_atlas-Glasser_
              measure-pearsoncorrelation_conmat.tsv
              ├──sub-20712_ses-12367_task-restbold_run-1_space-MNI152NLin6Asym_atlas-Glasser_
              reho.tsv
              ├──sub-20712_ses-12367_task-restbold_run-1_space-MNI152NLin6Asym_atlas-Glasser_
              timeseries.tsv
              ├──sub-20712_ses-12367_task-restbold_run-1_space-MNI152NLin6Asym_atlas-Gordon_c
              overage.tsv
              ├──sub-20712_ses-12367_task-restbold_run-1_space-MNI152NLin6Asym_atlas-Gordon_m
              easure-pearsoncorrelation_conmat.tsv
              ├──sub-20712_ses-12367_task-restbold_run-1_space-MNI152NLin6Asym_atlas-Gordon_r
              eho.tsv
              ├──sub-20712_ses-12367_task-restbold_run-1_space-MNI152NLin6Asym_atlas-Gordon_t
              imeseries.tsv
              ├──sub-20712_ses-12367_task-restbold_run-1_space-MNI152NLin6Asym_atlas-HCP_cove
              rage.tsv
              ├──sub-20712_ses-12367_task-restbold_run-1_space-MNI152NLin6Asym_atlas-HCP_meas
              ure-pearsoncorrelation_conmat.tsv
              ├──sub-20712_ses-12367_task-restbold_run-1_space-MNI152NLin6Asym_atlas-HCP_reho
              .tsv
```

```
├──sub-20712_ses-12367_task-restbold_run-1_space-MNI152NLin6Asym_atlas-HCP_time
series.tsv
├──sub-20712_ses-12367_task-restbold_run-1_space-MNI152NLin6Asym_atlas-Schaefer
1017_coverage.tsv
├──sub-20712_ses-12367_task-restbold_run-1_space-MNI152NLin6Asym_atlas-Schaefer
1017_measure-pearsoncorrelation_conmat.tsv
├──sub-20712_ses-12367_task-restbold_run-1_space-MNI152NLin6Asym_atlas-Schaefer
1017_reho.tsv
├──sub-20712_ses-12367_task-restbold_run-1_space-MNI152NLin6Asym_atlas-Schaefer
1017_timeseries.tsv
├──sub-20712_ses-12367_task-restbold_run-1_space-MNI152NLin6Asym_atlas-Schaefer
117_coverage.tsv
├──sub-20712_ses-12367_task-restbold_run-1_space-MNI152NLin6Asym_atlas-Schaefer
117_measure-pearsoncorrelation_conmat.tsv
├──sub-20712_ses-12367_task-restbold_run-1_space-MNI152NLin6Asym_atlas-Schaefer
117_reho.tsv
├──sub-20712_ses-12367_task-restbold_run-1_space-MNI152NLin6Asym_atlas-Schaefer
117_timeseries.tsv
├──sub-20712_ses-12367_task-restbold_run-1_space-MNI152NLin6Asym_atlas-Schaefer
217_coverage.tsv
├──sub-20712_ses-12367_task-restbold_run-1_space-MNI152NLin6Asym_atlas-Schaefer
217_measure-pearsoncorrelation_conmat.tsv
├──sub-20712_ses-12367_task-restbold_run-1_space-MNI152NLin6Asym_atlas-Schaefer
217_reho.tsv
├──sub-20712_ses-12367_task-restbold_run-1_space-MNI152NLin6Asym_atlas-Schaefer
217_timeseries.tsv
├──sub-20712_ses-12367_task-restbold_run-1_space-MNI152NLin6Asym_atlas-Schaefer
317_coverage.tsv
├──sub-20712_ses-12367_task-restbold_run-1_space-MNI152NLin6Asym_atlas-Schaefer
317_measure-pearsoncorrelation_conmat.tsv
├──sub-20712_ses-12367_task-restbold_run-1_space-MNI152NLin6Asym_atlas-Schaefer
317_reho.tsv
├──sub-20712_ses-12367_task-restbold_run-1_space-MNI152NLin6Asym_atlas-Schaefer
317_timeseries.tsv
├──sub-20712_ses-12367_task-restbold_run-1_space-MNI152NLin6Asym_atlas-Schaefer
417_coverage.tsv
├──sub-20712_ses-12367_task-restbold_run-1_space-MNI152NLin6Asym_atlas-Schaefer
417_measure-pearsoncorrelation_conmat.tsv
├──sub-20712_ses-12367_task-restbold_run-1_space-MNI152NLin6Asym_atlas-Schaefer
417_reho.tsv
├──sub-20712_ses-12367_task-restbold_run-1_space-MNI152NLin6Asym_atlas-Schaefer
417_timeseries.tsv
├──sub-20712_ses-12367_task-restbold_run-1_space-MNI152NLin6Asym_atlas-Schaefer
517_coverage.tsv
├──sub-20712_ses-12367_task-restbold_run-1_space-MNI152NLin6Asym_atlas-Schaefer
517_measure-pearsoncorrelation_conmat.tsv
├──sub-20712_ses-12367_task-restbold_run-1_space-MNI152NLin6Asym_atlas-Schaefer
517_reho.tsv
├──sub-20712_ses-12367_task-restbold_run-1_space-MNI152NLin6Asym_atlas-Schaefer
517_timeseries.tsv
├──sub-20712_ses-12367_task-restbold_run-1_space-MNI152NLin6Asym_atlas-Schaefer
617_coverage.tsv
├──sub-20712_ses-12367_task-restbold_run-1_space-MNI152NLin6Asym_atlas-Schaefer
617_measure-pearsoncorrelation_conmat.tsv
├──sub-20712_ses-12367_task-restbold_run-1_space-MNI152NLin6Asym_atlas-Schaefer
617_reho.tsv
├──sub-20712_ses-12367_task-restbold_run-1_space-MNI152NLin6Asym_atlas-Schaefer
617_timeseries.tsv
├──sub-20712_ses-12367_task-restbold_run-1_space-MNI152NLin6Asym_atlas-Schaefer
717_coverage.tsv
├──sub-20712_ses-12367_task-restbold_run-1_space-MNI152NLin6Asym_atlas-Schaefer
717_measure-pearsoncorrelation_conmat.tsv
```

```
├──sub-20712_ses-12367_task-restbold_run-1_space-MNI152NLin6Asym_atlas-Schaefer717_reho.tsv
├──sub-20712_ses-12367_task-restbold_run-1_space-MNI152NLin6Asym_atlas-Schaefer717_timeseries.tsv
├──sub-20712_ses-12367_task-restbold_run-1_space-MNI152NLin6Asym_atlas-Schaefer817_coverage.tsv
├──sub-20712_ses-12367_task-restbold_run-1_space-MNI152NLin6Asym_atlas-Schaefer817_measure-pearsoncorrelation_conmat.tsv
├──sub-20712_ses-12367_task-restbold_run-1_space-MNI152NLin6Asym_atlas-Schaefer817_reho.tsv
├──sub-20712_ses-12367_task-restbold_run-1_space-MNI152NLin6Asym_atlas-Schaefer817_timeseries.tsv
├──sub-20712_ses-12367_task-restbold_run-1_space-MNI152NLin6Asym_atlas-Schaefer917_coverage.tsv
├──sub-20712_ses-12367_task-restbold_run-1_space-MNI152NLin6Asym_atlas-Schaefer917_measure-pearsoncorrelation_conmat.tsv
├──sub-20712_ses-12367_task-restbold_run-1_space-MNI152NLin6Asym_atlas-Schaefer917_reho.tsv
├──sub-20712_ses-12367_task-restbold_run-1_space-MNI152NLin6Asym_atlas-Schaefer917_timeseries.tsv
├──sub-20712_ses-12367_task-restbold_run-1_space-MNI152NLin6Asym_atlas-Tian_coverage.tsv
├──sub-20712_ses-12367_task-restbold_run-1_space-MNI152NLin6Asym_atlas-Tian_measure-pearsoncorrelation_conmat.tsv
├──sub-20712_ses-12367_task-restbold_run-1_space-MNI152NLin6Asym_atlas-Tian_reho.tsv
├──sub-20712_ses-12367_task-restbold_run-1_space-MNI152NLin6Asym_atlas-Tian_timeseries.tsv
├──sub-20712_ses-12367_task-restbold_run-1_space-MNI152NLin6Asym_desc-linc_qc.csv
├──sub-20712_ses-12367_task-restbold_run-1_space-MNI152NLin6Asym_res-2_desc-denoised_bold.json
├──sub-20712_ses-12367_task-restbold_run-1_space-MNI152NLin6Asym_res-2_desc-denoised_bold.nii.gz
├──sub-20712_ses-12367_task-restbold_run-1_space-MNI152NLin6Asym_res-2_desc-denoisedSmoothed_bold.json
├──sub-20712_ses-12367_task-restbold_run-1_space-MNI152NLin6Asym_res-2_desc-denoisedSmoothed_bold.nii.gz
├──sub-20712_ses-12367_task-restbold_run-1_space-MNI152NLin6Asym_res-2_reho.json
└──sub-20712_ses-12367_task-restbold_run-1_space-MNI152NLin6Asym_res-2_reho.nii.gz
```

**Supplemental Figure 3**: Example outputs of the XCP-D walkthrough for one run
